# Post-Translational Modifications Control Phase Transitions of Tau

**DOI:** 10.1101/2024.03.08.583040

**Authors:** Wyatt C. Powell, McKinley Nahum, Karl Pankratz, Morgane Herlory, James Greenwood, Darya Poliyenko, Patrick Holland, Ruiheng Jing, Luke Biggerstaff, Michael H. B. Stowell, Maciej A. Walczak

## Abstract

The self-assembly of Tau(297-391) into filaments, which mirror the structures observed in Alzheimer’s disease (AD) brains, raises questions about the role of AD-specific post-translational modifications (PTMs) in the formation of paired helical filaments (PHFs). To investigate this, we developed a synthetic approach to produce Tau(291-391) featuring N-acetyllysine, phosphoserine, phosphotyrosine, and N-glycosylation at positions commonly modified in post-mortem AD brains, thus facilitating the study of their roles in Tau pathology. Using transmission electron microscopy (TEM), cryo-electron microscopy (cryo-EM), and a range of optical microscopy techniques, we discovered that these modifications generally hinder the *in vitro* assembly of Tau into PHFs. Interestingly, while acetylation’s effect on Tau assembly displayed variability, either promoting or inhibiting phase transitions in the context of cofactor free aggregation, heparin-induced aggregation, and RNA-mediated liquid-liquid phase separation (LLPS), phosphorylation uniformly mitigated these processes. Our observations suggest that PTMs, particularly those situated outside the fibril’s rigid core are pivotal in the nucleation of PHFs. Moreover, in scenarios involving heparin-induced aggregation leading to the formation of heterogeneous aggregates, most AD-specific PTMs, except for K311, appeared to decelerate the aggregation process. The impact of acetylation on RNA-induced LLPS was notably site-dependent, exhibiting both facilitative and inhibitory effects, whereas phosphorylation consistently reduced LLPS across all proteoforms examined. These insights underscore the complex interplay between site-specific PTMs and environmental factors in modulating Tau aggregation kinetics, enhancing our understanding of the molecular underpinnings of Tau pathology in AD and highlighting the critical role of PTMs located outside the ordered filament core in driving the self-assembly of Tau into PHF structures.

## INTRODUCTION

Deposits of microtubule associated protein Tau are implicated in the onset and progression of several neurodegenerative diseases known as tauopathies.^1^ Primary tauopathies such as corticobasal degeneration (CBD) are characterized by neuronal and glial inclusions. In secondary tauopathies including Alzheimer’s disease (AD), Tau in neurofibrillary tangles (NFTs) is found together with amyloid-β plaques. Tau filaments from different patients with the same disease including AD, chronic traumatic encephalopathy (CTE), CBD, and Pick’s disease reproducibly adopt the same unique molecular structures.^2–5^ These disease-specific polymorphs may represent low energy conformations of Tau, and, together with environmental factors, likely play a role in the structural diversity and filament propagation.^6–9^ However, many *in vitro* and mouse model filaments create multiple Tau polymorphs and do not resemble the structures observed in human tauopathies.^10–12^ Only recently aided by high-throughput cryo-EM, short fragments of Tau have been shown to self-assemble into the AD and CTE filaments, but these peptides also form many other structures under similar conditions (Figure 1).^13–17^

**Figure 1.**
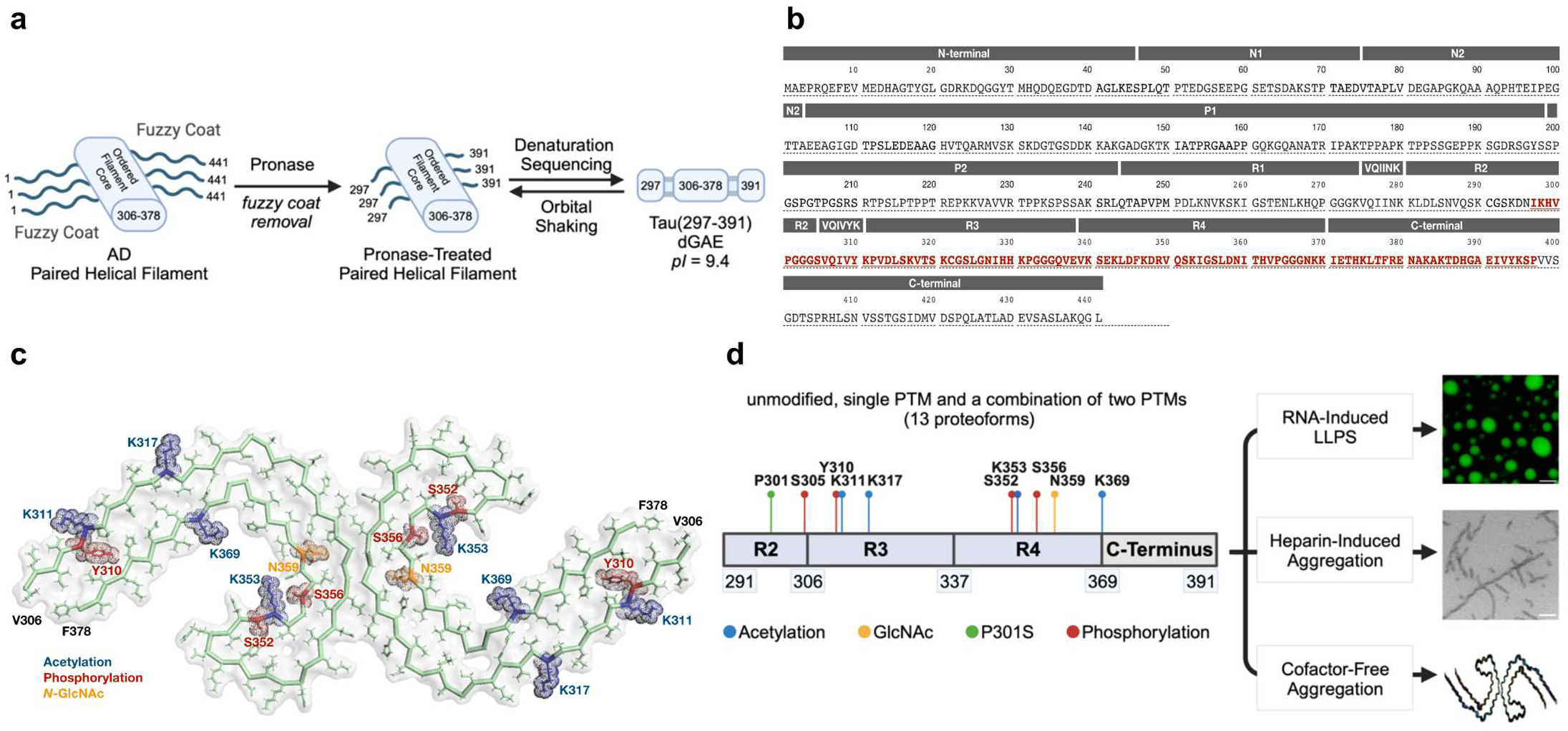
(a) Identification of the PHF core sequence from AD patients. Alzheimer’s disease paired helical filaments consist of a rigid filament core and a fuzzy coat that contains the N- and C-terminal regions. Cleavage of the fuzzy coat from AD paired helical filaments with pronase releases the proteolytically stable filament core.^55,56^ (b) Primary sequence of 2N4R Tau with the 297-391 peptide region highlighted in red. (c) Mapping of selected PTMs onto the AD core structure (PDB: 5O3L). (d) An overview of the study plan.

The factors that assist or control folding of full-length Tau into single disease specific polymorphs remain unclear. In understanding the structural polymorphism of Tau, post-translational modifications (PTMs) that alter the charge and hydrophobicity of amino acid side chains need to be considered.^18^ Tau filaments from AD and other tauopathies are heavily modified with PTMs, and site-specific modifications unique to the disease-associated filaments are also observed.^19,20^ It has been suggested that depending on the location of the modification relative to the fibril core, PTMs can alter the primary structure of Tau and drive folding into specific structures.^21–24^ Alternatively, Tau filaments may be nucleated by environmental factors, and site specific PTMs occur due to the pre-established conformation of the filament. Hydrophobic and non-proteinaceous cofactor densities are observed in several types of Tau filaments indicating strong interactions between various cellular components and Tau aggregates.^25–27^ Seeding of Tau in cells with AD- and CBD-derived brain seeds does not recapitulate the structures of the added brain seeds, and a similar effect is observed for α-synuclein filaments from multiple systems atrophy brain seeds.^28–30^ Understanding the role of these factors may yield a better insight into the mechanism of the disease, and potentially provide a platform to fine-tune diagnostic tools (e.g., antibodies) or even novel therapeutic strategies that capitalize on aberrant PTMs. Here, we describe synthetic, structural, and biophysical studies that comprehensively evaluate the role of each PTM in the ordered AD region, and we demonstrate the first example of a fully synthetic AD fibril produced by chemical means. This study also establishes a relationship between the position of the PTM and its impact on phase transitions. We demonstrate the synergy between two complementary PTMs that can be challenging to decipher with other methods, we show that some PTMs dominate the biophysical behavior of Tau.

## RESULTS

To probe the role of post-translational modifications in Tau phase transitions, we used chemical protein synthesis (Figure 1). We aimed to understand how site-specific modifications (acetylation, phosphorylation, and glycosylation) modulate AD filament structure, regulate aggregation, and liquid-liquid phase separation (LLPS). We hypothesized that AD-specific PTMs can facilitate the nucleation of Tau paired helical filaments *in vitro*. The Tau region of interest was selected based on the published study that identified Tau(297-391) as the sequence that recapitulates the AD fold in vitro,^13–17^ and our prior synthetic work which indicated a strategically advantageous synthetic disconnection at C291 to produce a peptide fragment Tau(291-391).^45^ We selected three PTMs that are linked to Tau physiology and disfunction:

A. Acetylation of Tau filaments occurs in the region that forms the rigid fibril core and usually overlaps with ubiquitination sites.^22,31^ For this study, we selected four individual acetylated positions (AcK311, AcK317, AcK353, and AcK369), which have been observed with high patient frequency in AD, globular glial tauopathy (GGT), frontotemporal dementia with parkinsonism-17 (FTDP-17), CBD, progressive supranuclear palsy (PSP), and Pick’s disease.^22,31^
B. Phosphorylation of Tau filaments is typically observed outside of the ordered fibril core, and the phosphorylation sites within the selected region are likely important for regulating microtubule binding. In our study, we selected four phosphorylation sites (pS305, pY310, pS352, and pS356) that all have been observed with various frequencies in AD, with pS305 being the most common AD-specific phosphorylation in this region.^31^ Additionally, two positions (pS356 and pS352) are also important for regulating microtubules binding. Three of these sites are modified serine sites and one unique position is phosphorylated tyrosine.
C. N-Glycosylation of Tau only occurs in AD patients but not in healthy controls, and it might be responsible for *in vivo* neurofibrillary tangle formation.^32–34^ We chose to investigate N-acetyl glucosamine at GlcNAcN359 as one of the two potential glycosylation sites.
D. To better understand the interplay between two orthogonal PTMs (phosphorylation or acetylation) and N-glycosylation,^35^ we included two additional proteoforms designated as GlcNAcK359+AcK353 and GlcNAcN359+pS305.
E. Finally, to complement these studies, we selected another relevant modification (P301S) that is linked to inherited FTDP-17 and CBD.^36^ In this mutation, we used only unmodified protein chain which served as the reference point and a benchmark together with the unmodified (WT) Tau(291-391) peptide. Together, the described study includes thirteen unique proteoforms.

### A. Synthesis of Tau(291-391)

To access the selected modifications of Tau, we used chemical protein synthesis (Figure 2). In our approach we divided the 101-mer peptide into three segments and merged them using the Native Chemical Ligation (NCL) reaction.^37^ Because the Tau(291-391) region contains an internal cysteine residue, we selected the K322-C321 junction for the last ligation step. This synthetic step would also be conducted using the tandem NCL-Thz removal protocol.^38^ For the first ligation site, we selected the L357-D358 junction using the diselenide-selenoester ligation (DSL)-deselenization protocol.^39–41^ Both the K322-C321 and L357-D358 ligation sites have been utilized in the previous syntheses.^42–45^ Using these disconnections, three peptide segments were envisioned: Tau(291-321), Tau(322-357), and Tau(358-391). These segments were designed to contain the relevant PTMs and could be assembled in a combinatorial fashion.

**Figure 2.**
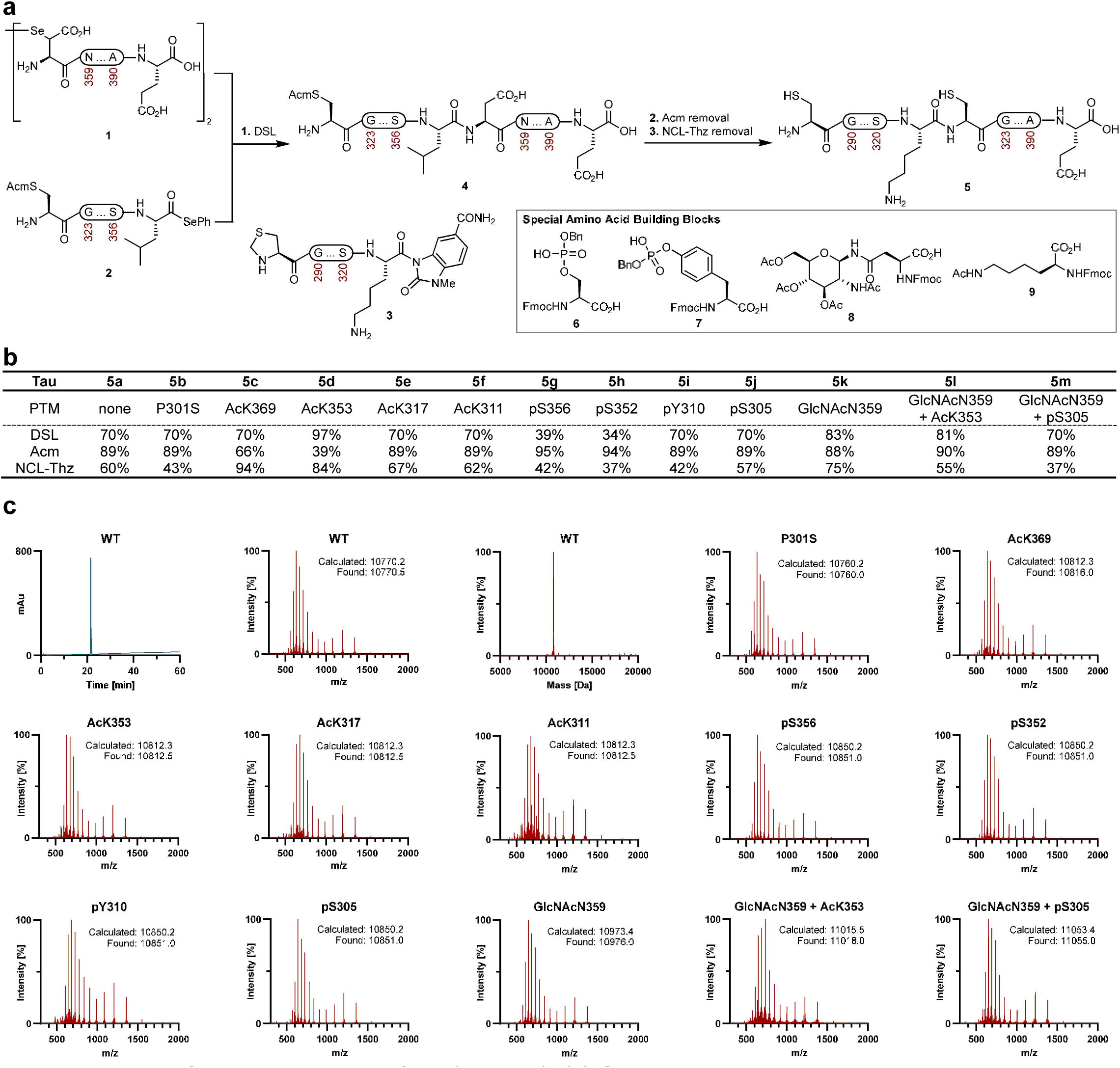
Chemical synthesis of Tau(291-391). (a) General synthetic scheme outlining the key synthetic peptide fragments used to assemble Tau(291-391). Reaction conditions: **1.** DSL: 1a. 6 M Gnd·HCl, 100 mM Na_2_HPO_4_, pH 6.2, 23 ⁰C, 1 h; 1b. extraction with hexane; 1c. adjust to 2% (v/v) hydrazine hydrate, pH 7.4, 23 ⁰C, 10 min; 1d. adjust to 83 mM TCEP·HCl, 8 mM DTT, pH 5.3, 23 ⁰C, 30 min. **2.** Acm removal: AgOAc (30 equiv.), AcOH:H_2_O (1:1), 23 ⁰C, 6 h; or PdCl_2_ (5 equiv.), MgCl_2_ (15 equiv.), 6 M Gnd·HCl, 0.1 M Na_2_HPO_4_, pH 7.2, 37 ⁰C, 2 h. **3.** NCL-Thz removal: 200 mM MPAA, 6 M Gnd·HCl, 200 mM Na_2_HPO_4_, 20 mM TCEP·HCl, pH 7.0, 23 ⁰C, 4 h; adjust to 200 mM MeONH_2_·HCl, pH 4.0, 23 ⁰C, 4 h. (b) Summary of reactions yields for each Tau(291-391) proteoform with individual PTMs. DSL refers to step 1 in Figure 2a and indicates isolated yield of peptide fragment **4** after deselenization. Acm refers to the yields of an isolated intermediate after Acm removal. NCL-Thz removal refers to the yields of purified **5** obtained after the NCL with fragments **3** followed by deprotection of the final product with MeONH_2_. (c) LC-MS data for unmodified Tau(291-391) (first three entries) and ESI-MS data for the remaining Tau(291-391) proteoforms. LC gradient for WT: 5-65% MeCN/H_2_O/0.05% TFA over 60 min, 4.6x100 mm, 4 µm, 100 Å, InfinityLab Poroshell 120 EC-C18 column (214 nm UV detection). Abbreviations: Ac – acetyl; Acm – acetamidomethyl; Bn – benzyl; DSL – diselenide selenoester ligation; DTT – dithiothreitol; ESI – electrospray ionization; Fmoc – fluorenylmethyloxycarbonyl; Gnd – guanidine; LC – liquid chromatography; MS – mass spectroscopy; MPAA – 4-mercaptophenylacetic acid; NCL – native chemical ligation; TCEP – tris(2-carboxyethyl)phosphine; Thz – thiazolidine.

Several solid-phase peptide synthesis (SPPS) methods are crucial to prepare the peptide segments with post-translational modifications. Segment **1** corresponding to Tau(358-391) was prepared on the Wang-peg resin to generate the C-terminal carboxylic acid, and the diselenide intermediate was formed from a PMB-protected selenoether under oxidative conditions (TFA, DMSO).^41^ The unmodified and AcK369 diselenides were relatively stable to the typical synthetic manipulations, but the GlcNAcN359 analog was prone to deselenization and was lyophilized immediately after HPLC purification, and protected from light to minimize material loss. Segment **2**, which corresponds to Tau(322-357), contains an N-terminal Acm cysteine and a C-terminal selenoester. Segment **2** was prepared on HMPB Chem-Matrix resin, and after SPPS, the protected peptide acid was selectively detached from the resin, and a phenyl selenoester was installed at the C-terminus using DPDS/P(*n*-Bu)_3_.^46^ Segment **3**, which corresponds to Tau(291-321), contained an N-terminal Thz protective group and has a C-terminal Nbz linker, which behaves like a thioester during the NCL step.^47^ We note that the phosphorylated amino acids building blocks were quantitatively coupled onto the growing chain using 2.5 equivalents of the reagent and a combination of HATU/Oxyma/DIPEA for 16 h.^48^ Selenoaspartic acid building block used in the preparation of fragment **1** was coupled overnight using 2 equivalents of the reagent, but with DIC/Oxyma for 16 h. The SPPS methods were effective for the preparation of all the peptide segments with PTMs in similar yield and purity (for details, see the SI).

The forward synthesis of the Tau(291-391) peptides was completed in a combinatorial manner. Segments **1** and **2** were ligated by the one-pot method using the diselenide-selenoester-ligation-deselenization, and oligopeptides **4** were prepared in high yields (70-97%), except for pS356 and pS352. The Acm group in **4** was removed using either PdCl_2_/MgCl_2_ or AgOAc in AcOH/H_2_O producing unprotected peptides in 39-95% yield.^49,50^ The last peptide union was performed by a ligation between the free N-terminal cysteine and **3** using MPAA as the catalyst, and the Thz group was removed with MeONH_2_.^38,51^ All of the Tau(291-391) peptides behaved similarly under these conditions and produced comparable yields (Figure 2b). However, Tau(291-322)-MeNbz fragment for the P301S mutant was poorly soluble in 6 M Gnd•HCl, and this fragment was solubilized with 8 M Gnd•HCl. The final peptides were purified by sample displacement mode chromatography with good yields achieved after LC.^52^ All the proteoforms were homogeneous by LC/HRMS and stable to the subsequent manipulations (Figure 2c).

### B. Characterization of Tau(291-391)

We characterized the synthetic Tau(291-391) proteins with gel electrophoresis to probe for their gel mobility (Figure 3a). Abnormal Tau from the AD brain has a visible gel shift when compared to the normal Tau due to the presence of PTMs.^53,54^ We found that with the exception of GlcNAcN359+pS305, which has a slightly increased gel shift, the electrophoretic mobilities of all Tau(291-391) constructs were similar (all proteoforms have an apparent molecular weight of ∼12 kDa). From this study, we conclude that site specific PTMs in this region of Tau do not cause a significant upward shift in the gel, supporting the observation that PTMs are present within proteolytically stable Tau PHFs from AD brains.^55,56^ To test if site specific PTMs induce a conformational change of Tau, we used temperature dependent circular dichroism (CD) spectroscopy (Figure 3b). The CD spectra of all Tau(291-391) proteoforms have a minimum around 200 nm, which is indicative of random coil and have a negative patch at 217 nm that is indicative of β-structure.^57,58^ A positive peak at 217 nm could also indicate the presence of a PPII helix. However, distinguishing PPII or β-structure may be difficult when working with proteins and not short peptides.^59,60^ Upon heating to 55 °C, the point at 200 nm becomes less pronounced, the patch at 217 nm becomes more negative, there is an isobesic point around 210 nm, and the minimum peak at 200 nm undergoes a red shift of about 2 nm (Figure 3b). All the Tau constructs have a similar CD spectrum as the unmodified WT Tau with some differences in the intensity at 200 nm and 217 nm at low temperatures for acetyl and phosphorylated forms (Figure 3c). To better understand how the folding changes in response to temperature, we plotted the Ɵ values for all proteoforms at 200 and 217 nm as a function of temperature (Figure 3d). This ratio would appear sigmoidal if cooperative folding occurs, and as a straight line if there was a structural change without cooperative folding.^58^ For the unmodified Tau(291-391), P301S, and GlcNAcN359, the plot shows an open curve (from ∼7 to 3) but the curve does not appear to have a sigmoidal shape.^58^ In contrast, the acetyl and phosphate modified constructs have a similar plots that decrease (6 to 3) with a more sigmoidal curve. The most striking differences were noted at 10 °C – the acetyl and phosphate PTMs had smaller ratios (∼5.5) than WT and P301S (∼7). Interestingly, glycans (∼6.5) also had larger ratios at 10 °C. For WT, P301S, and GlcNAc this could indicate that there is an additional contribution of PPII at the low temperature, which is consistent with previous findings for N-linked glycoproteins.^61^ Our findings also contrast with previous reports of phosphorylated tau having more PPII than WT, as the peak at 217 nm would be expected to be more positive.^62,63^ Taken together, the acetyl and phosphopeptides have more β-structure (or less PPII) at 10 °C than the GlcNAc and unmodified peptides, and all of the graphs converged upon heating. The 200/217 ratios suggest that WT and glycans undergo a structural transition that does not occur in a cooperative fashion, but the acetyl and phosphorylated peptides have a higher folding tendency at low temperature (although still not cooperative) because of the charge neutralization and/or increased hydrophobicity.

**Figure 3.**
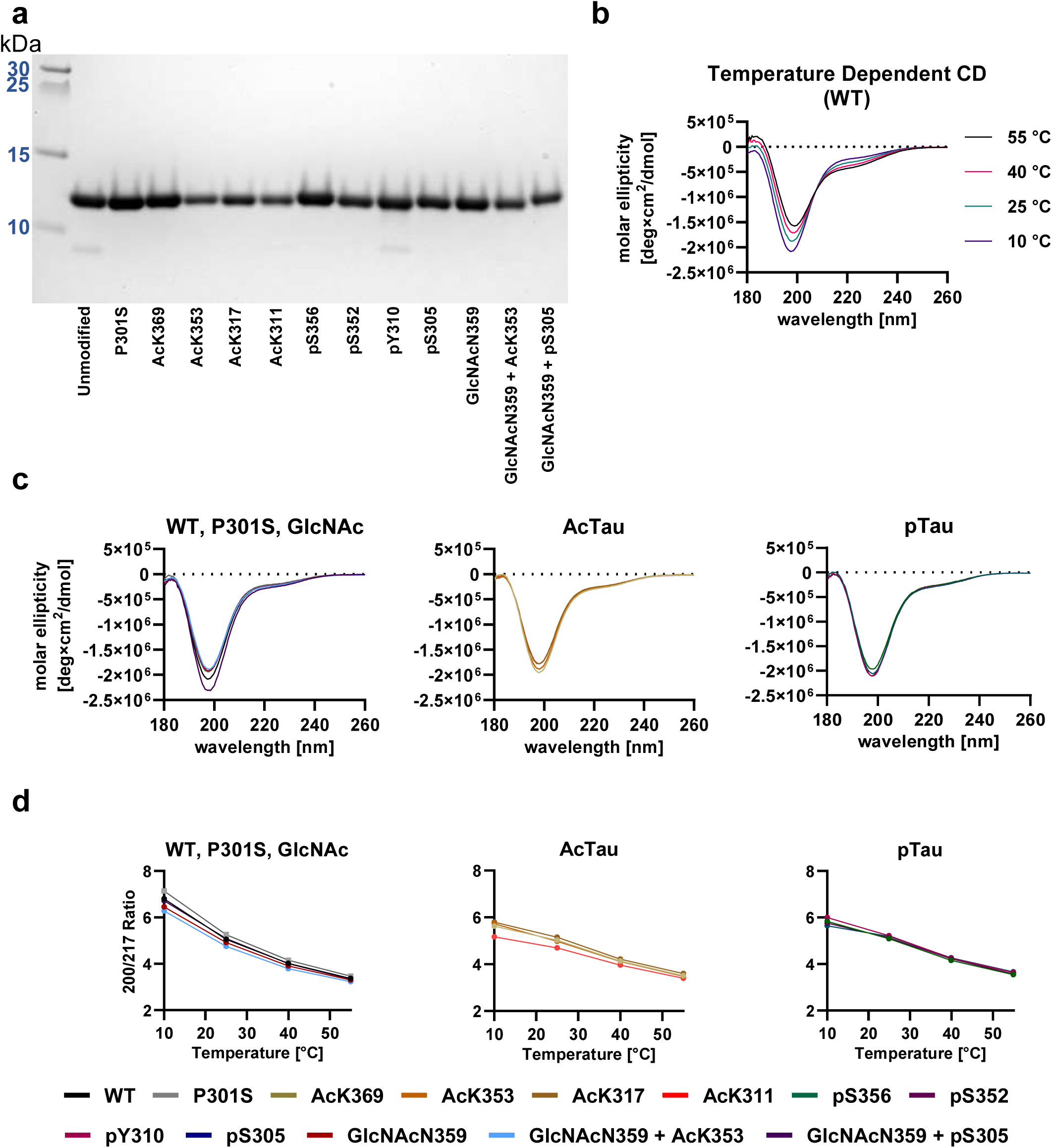
Characterization of Tau(291-391) proteoforms. The CD experiments were performed in triplicate. (a) 16% SDS-PAGE (Novex tricine) of the Tau(291-391) constructs with PTMs. (b) Temperature dependent CD spectra of 0.4 mg/mL Tau(291-391) WT at 10, 25, 40, and 55 ⁰C in 20 mM sodium phosphate buffer pH 7.4. (c) CD spectra of 0.4 mg/mL Tau(291-391) at 25 ⁰C, in 20 mM sodium phosphate buffer pH 7.4. (d) The ratio of Ɵ at 200/217 nm for the Tau(291-391) constructs. Abbreviations: CD – circular dichroism; PAGE – polyacrylamide gel electrophoresis; SDS – sodium dodecyl sulfate.

### C. Cofactor Free Self-Assembly

Next, we tested the capacity of Tau constructs to self-assemble without cofactors into PHFs or other filamentous structures, hypothesizing that AD-specific post-translational modifications might enhance the formation of PHFs, making the self-assembly process more selective and potentially more efficient. We initiated the self-assembly of the constructs at a concentration of 4 mg/mL in MgCl_2_ buffer with 300 rpm shaking, a condition previously identified to facilitate PHF formation.^14^ Subsequently, the mixtures were negatively stained for TEM analysis. Three Tau constructs (WT, AcK317, and AcK311) were observed to form fibrils at a significantly high density (Figure 4a-b). The AcK317 samples displayed short and amorphous fibrils, whereas AcK311 yielded long rod-like filaments. The WT fibrils were considered suitable for cryo-electron microscopy (cryo-EM) analysis, leading to the generation of a high-resolution 3D structure resembling PHFs found in AD brains, as well as recombinantly produced fibrils from Tau(297-391) (Figure 4c-f). However, attempts to produce high-resolution structures from the two acetyl derivatives were unsuccessful due to their fragile nature or their non-twisting topology. These findings suggest that (a) PTMs outside the ordered fibril core play a significant role in inducing Tau aggregation into PHFs, (b) the disease specific PTMs we investigated are introduced post-PHF formation, targeting specific residues influenced by the pre-established conformation of the PHF, or (c) PTMs have low occupancy, but have high patient frequency.

**Figure 4.**
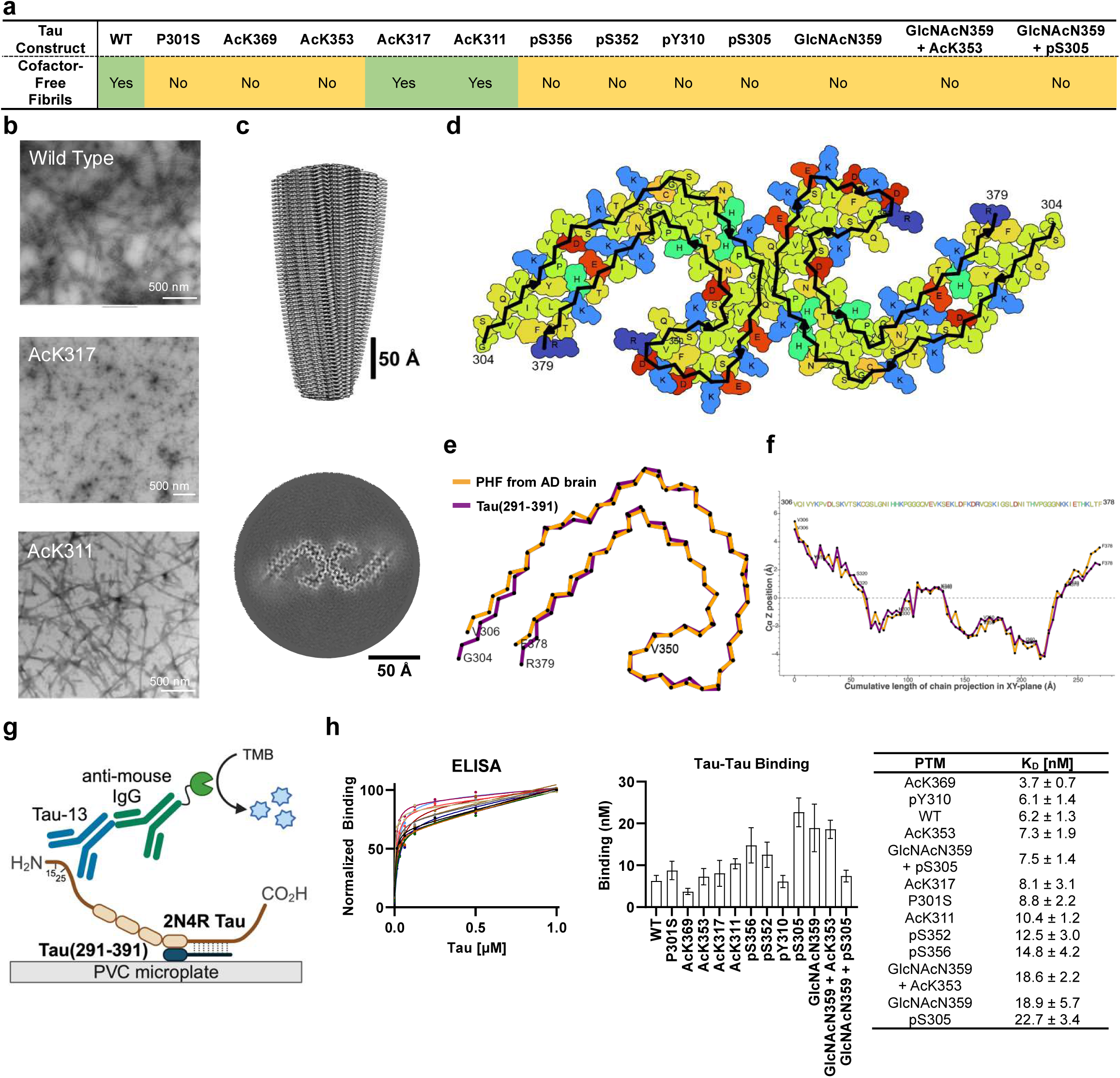
Cofactor-free self-assembly of Tau(291-391). (a) Summary of cofactor-free self-assembly of Tau(291-391) confirmed by negative stain TEM. Aggregation conditions: 4 mg/mL Tau(291-391), 200 mM MgCl_2_, 10 mM DTT, 20 mM Na_2_HPO_4_, pH 7.4, 37 ⁰C, 48 h. (b) Representative TEM micrographs of Tau(291-391) proteoforms obtained under the standard conditions. (c)-(d) Cryo-EM structure of unmodified WT Tau(291-391). (e)-(f) Comparison between Tau(291-391) and PHFs from AD brain (PDB: 5O3L). (g) Schematic of the ELISA to quantify interactions between Tau(291-391) and 2N4R Tau. In brief, Tau(291-391) proteoforms were immobilized on PVC plate and treated with recombinant 2N4R Tau. Binding affinities were quantified colorimetrically (HRP) with Tau-13 mAb recognizing residues 15-25 of 2N4R Tau. (h). Langmuir binding curves and tabulated K_D_ values for the ELISA. The apparent K_D_ values correspond to the average of three replicates and the error bars represent SD. Abbreviations: HRP – horseradish peroxidase; PVC – polyvinyl chloride; TMB – 3,3’,5,5’-tetramethylbenzidine.

The self-assembly of Tau(297-391) is facilitated by nanomolar Tau-Tau binding affinities.^64^ If the binding is primarily electrostatic, then PTMs that neutralize charge should enhance Tau-Tau binding by reducing charge-charge repulsion. To evaluate this hypothesis, Tau(291-391) constructs were anchored on PVC microplates, and their interactions with 2N4R Tau were assessed using an N-terminal specific monoclonal antibody, as shown in Figure 4g-h.^64,65^ Except for pY310 and AcK369, each proteoform exhibited a reduced binding affinity to 2N4R Tau compared to the WT. If Tau-Tau binding is electrostatic, phosphorylated PTMs would show stronger binding due to enhanced charge neutralization. Nevertheless, our observations revealed that phosphorylated Tau demonstrated inferior binding compared to acetylated constructs, with pY310 being an exception. Proteoforms modified with GlcNAcN359, pS305, and the combination of GlcNAcN359+AcK353 showed significantly diminished K_D_ compared to other PTMs, suggesting these modifications either introduce steric hindrance or destabilize crucial binding sites. Intriguingly, the combination of GlcNAcN359+pS305 exhibited stronger binding than each modification individually, indicating that multiple PTMs with weak binding can synergize, leading to non-linear effects on binding strength. Overall, our binding data implies that the Tau-Tau interaction may be governed more by hydrophobic and hydrogen bonding mechanisms rather than by electrostatic interactions, as charge-neutralizing PTMs appear to destabilize Tau-Tau binding.

### D. Aggregation Kinetics with Heparin

To better understand the role of PTMs in the heparin-induced fibrillation, we examined the aggregation kinetics of various Tau(291-391) constructs. The kinetics were assessed based on the time required to reach half of the maximum Thioflavin T (ThT) fluorescence, a measure of amyloid formation (Figure 5a). Due to the inherent differences in maximum ThT signals among Tau constructs, we normalized the aggregation assays to facilitate meaningful comparisons. The structures modified with acetylation displayed three distinct kinetic profiles: WT and AcK311 exhibited rapid aggregation, achieving half-maximum signal within an hour; AcK317 demonstrated slower aggregation kinetics compared to WT; both AcK353 and AcK369 also showed reduced kinetics similar to AcK317, but uniquely featured a second plateau, indicating a complex aggregation process. Phosphorylated constructs, in general, aggregated more slowly than the WT peptide. The pS356, pS352, and pY310 modifications all exhibited first and second kinetic plateaus akin to AcK353 and AcK369. Notably, pS305 and pY310, despite generating a ThT signal, failed to form filamentous structures under negative stain TEM, presenting only as amorphous aggregates. The GlcNAcN359 modification slightly decelerated the aggregation relative to WT, while the neutral P301S mutation expectedly sped up the process. The combined GlcNAcN359+AcK353 modification mirrored the kinetics of the AcK353 alone, with a slower aggregation rate and a secondary plateau. The GlcNAcN359+pS305 construct, however, did not produce a significant ThT signal, and filamentous structures were absent in TEM analyses, highlighting the profound impact of specific PTMs on Tau aggregation pathways. Collectively, our findings show PTMs generally reduce the heparin-induced aggregation kinetics of Tau, with some exceptions. Acetylation at specific sites, such as K311, accelerates aggregation, whereas acetylation at K369, K353, and K317 hinders the process, echoing similar findings for site-specific acetylation.^44,66–70^ Phosphorylation diminishes heparin induced aggregation, aligning with prior studies.^43,71,72^ The GlcNAc modification slows aggregation, and further contributes to our understanding of how N-glycosylation influences Tau pathology.^73^ Additionally, constructs with dual modifications tend to reflect the characteristics of their charge-neutralizing modifications. The P301S mutation’s pronounced effect in accelerating aggregation emphasizes the importance of genetic variations in tauopathy manifestations.^74,75^

**Figure 5.**
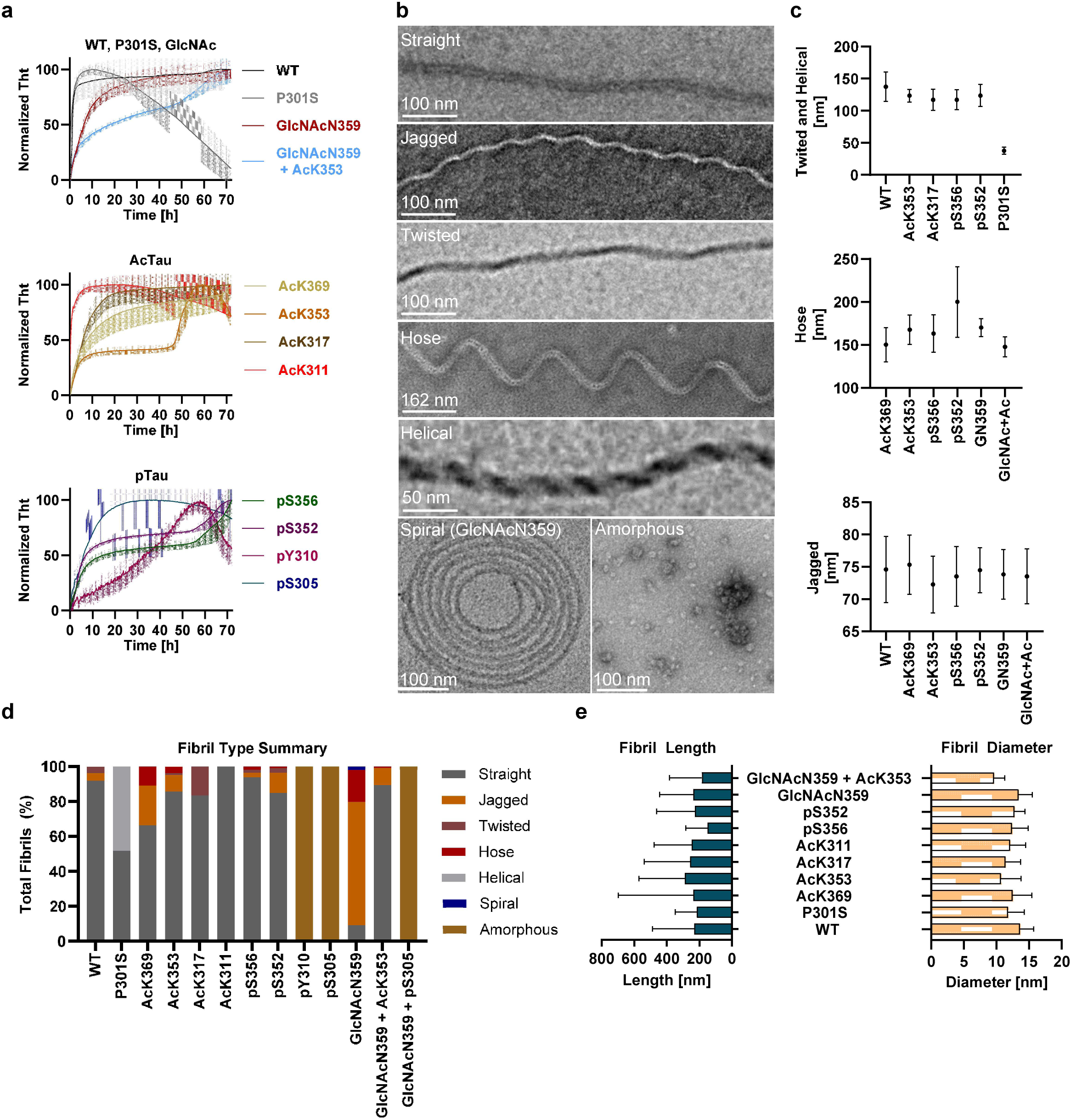
Heparin induced aggregation of Tau(291-391). All experiments were performed in triplicate. (a) ThT aggregation kinetics of Tau(291-391) proteoforms in the presence of heparin. Aggregation assay conditions: 100 µM Tau(291-391), 40 µM heparin, 50 mM NaCl, 10 mM DTT, 10 µM Tht, 25 mM HEPES, pH 7.4, 37 ⁰C, no shaking. The error bars represent SEM. (b) Representative negative stain images of various fibrils produced under conditions from Figure 5a. (c) Cross-over distances of fibril morphology, measured by negative stain TEM, for proteoforms that form stable fibrils with heparin. The error bars represent SD. (d) Sorting of Tau(291-391) fibrils based on the general structural features depicted in Figure 5b. (e) Length and diameter measurements of heparin-induced fibrils, determined with FibrilJ, and the error bars represent SD.

### E. TEM Analysis of Fibril Types

Given the distinct reaction kinetics presented by various PTMs, we examined the morphology of heparin-induced fibrils using negative stain TEM. This analysis revealed heterogeneous fibril populations including helical, jagged, twisted, hose-like, and straight morphologies (Figure 5b and 5c). Notably, straight fibrils emerged as the predominant form across the samples (Figure 5d). However, the GlcNAcN359 modification predominantly yielded jagged fibrils, while P301S was unique in forming a significant proportion of helical structures. Jagged fibrils were also common, with WT, AcK369, AcK353, pS356, pS352, and GlcNAcN359+AcK353 samples exhibiting 5-10% of this type. Hose-like fibrils were observed in several proteoforms including AcK369, AcK353, pS352, pS356, and GlcNAc, while twisted fibrils were identified in WT, AcK317, and pS352 samples, albeit in smaller proportions. This analysis concluded that PTMs significantly influence the distribution of fibril morphologies in heparin-induced aggregates, with certain modifications exerting dominant effects, and those in proximity showing similar morphological distributions.

In examining the crossover distances of the various fibril types, we found uniformity across the morphologies, with each type displaying consistent crossover distances indicative of structural similarities (Figure 5c). The consistent crossover distances among different fibril types further imply that despite the varied morphologies, there is a structural resemblance within each type.^10^ This comprehensive analysis concludes that PTMs significantly influence the distribution of fibril morphologies in heparin-induced aggregates, with certain modifications exerting dominant effects and those in proximity showing similar morphological distributions.

Further analysis with FibrilJ underscored small variations in fibril diameters and lengths (Figure 5e). Acetylation consistently reduced the average fibril diameter by 1-3 nm, whereas glycosylation alone had a lesser impact. Interestingly, phosphorylation mirrored the diameter reductions seen with acetylation. Notably, the P301S modification led to a broader diameter distribution, with an average reduction of 1.5 nm compared to WT. Kolmogorov-Smirnov tests confirmed the statistical significance (p < 0.0001) of these diameter variations, highlighting the impactful role of PTMs and mutations on fibril width distribution.

Fibril length analysis revealed that most modifications, except for AcK353, did not produce fibrils exceeding the lengths observed in WT samples. P301S fibrils exhibited the most constrained length distribution, with the majority measuring between 100-200 nm. This contrast in fibril lengths was also statistically significant across all groups (p < 0.0001), suggesting that while modifications influence fibril morphology and diameter, their impact on length is less consistent.

### F. RNA Induced Liquid-Liquid Phase Separation

LLPS of Tau may be an important intermediate phase that helps in the nucleation or oligomerization of Tau before fibrils are fully formed.^76^ This phase transition may also aid in concentrating Tau from a dilute solution to a more concentrated phase where aggregation occurs. Therefore, we wanted to know if LLPS of Tau(291-391) is a relevant phase transformation that aids in the aggregation of Tau. Like amyloid aggregation, there are two types of LLPS mechanisms: complex coacervation and self-coacervation. Electrostatically driven complex coacervation of Tau occurs when the positively charged lysine and arginine residues of Tau bind to negatively charged RNA, or other types of polyanions such as ssDNA, heparin, hyaluronic acid, and α-synuclein.^77–81^ Full-length Tau also undergoes electrostatically driven self-coacervation upon addition of molecular crowding agents (PEG, dextran) through attractive intramolecular electrostatic interactions between the negatively charged N-terminus and the positively charged MTBR.^82–85^ Hydrophobically driven self-coacervation of Tau occurs under high salt conditions or hyperphosphorylation, which weaken electrostatic interactions.^86,87^ However, Tau(291-391) did not undergo *in vitro* self-coacervation, as no droplets were observed even at 500 µM of Tau(291-391) under molecular crowding (15% PEG-10) or high salt (4.75 M NaCl) conditions. Nevertheless, Tau(291-391) underwent complex coacervation with RNA – treatment of Tau(291-391) with poly(U) immediately formed a visibly turbid solution, and liquid droplets were observed by fluorescence microscopy (Figure 6).

**Figure 6.**
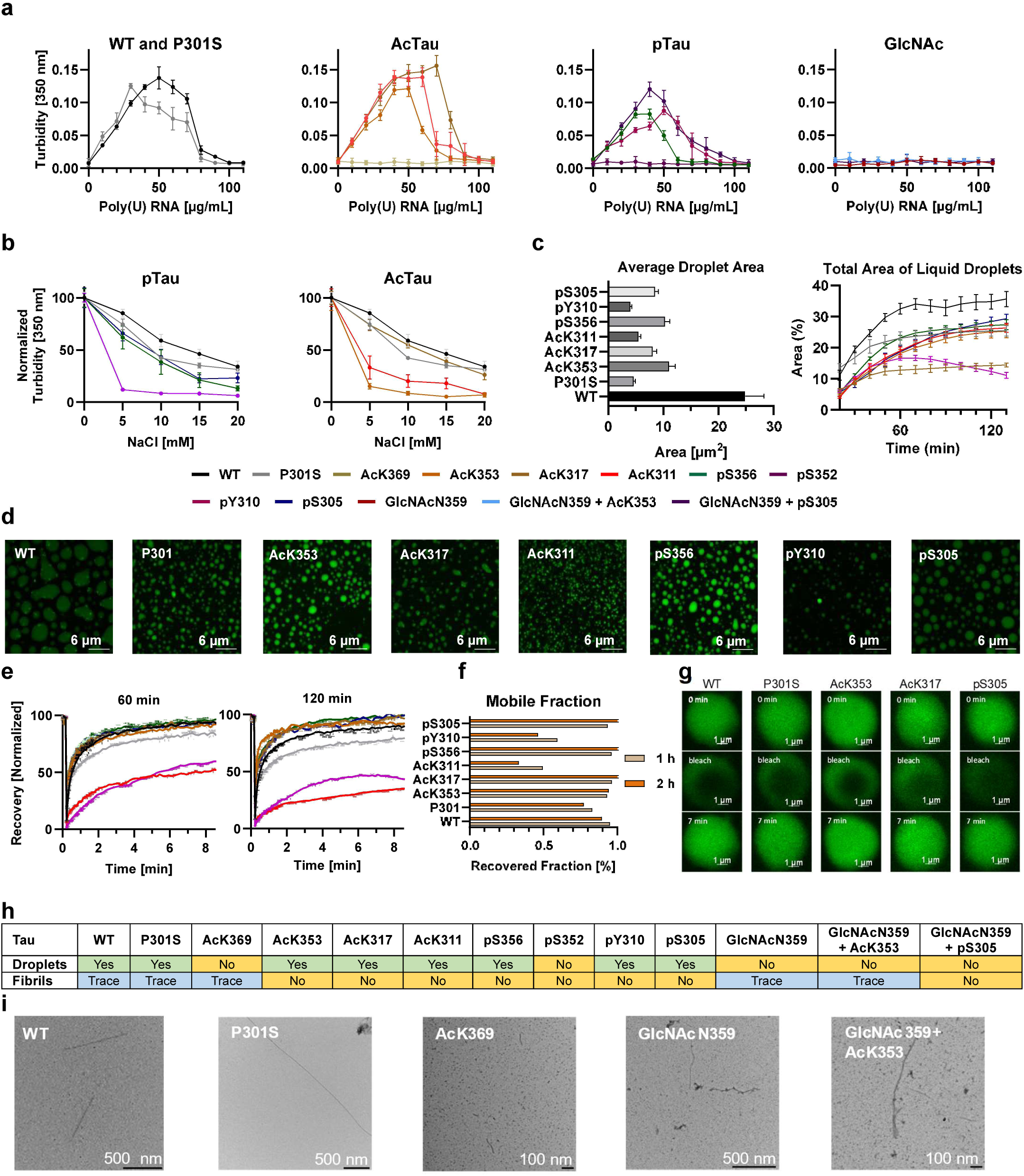
RNA-induced liquid-liquid phase separation of Tau(291-391). The turbidity and microscopy experiments were performed in triplicate and the error bars represent SEM. (a) Phase diagram of poly(U) RNA induced droplets. Turbidity was measure at 350 nm, 2 minutes after mixing with 30 µM Tau(291-391), variable µg/mL poly(U) RNA, 10 mM DTT, 25 mM HEPES, pH 7.4, 25 ⁰C. (b) Salt resistance in response of varying NaCl concentration measured by absorbance at 350 nm. Salt resistance assay diagram with 30 µM Tau(291-391), 40 µg/mL poly(U) RNA, 10 mM DTT, 25 mM HEPES, pH 7.4, 25 ⁰C, and variable mM NaCl. (c) Average droplet area, 2 h after preparing the droplets (left) and the total area of droplets that cover the microscope slide over time (right). (d) Images of Tau(291-391)-poly(U) RNA droplets, after 2 h, with 30 µM Tau(291-391), 0.3 µM AF 488 (adjusted to 1% labeling with AF488 labeled Tau(291-391)), 40 µg/mL poly(U) RNA, 10 mM DTT, 25 mM HEPES, pH 7.4, 25 ⁰C. Scale bar = 10 µm^2^ (e) FRAP kinetics for Tau(291-391) constructs with PTMS after 1 h and 2 h: 30 µM Tau(291-391), 0.3 µM AF488, 40 µg/mL poly(U) RNA, 10 mM DTT, 25 mM HEPES, pH 7.4, 23 ⁰C. (f) Calculated mobile fraction of the droplets, 1 and 2 h after preparing the droplets. (g) Representative images before, during, and after photobleaching, taken 1 h after preparing the droplets. (h) Summary of droplet and fibril formation under LLPS conditions: 30 µM Tau(291-391), 40 µg/mL poly(U) RNA, 10 mM DTT, 25 mM HEPES, pH 7.4, 23 ⁰C. The droplets are detected by turbidity after 2 h, and the fibrils are detected by negative stain TEM after 3 days. (h) Negative stain images of fibrils formed under LLPS conditions after 3 days.

Next, we wanted to test the effect of PTMs on the phase diagram of Tau(291-391). LLPS with RNA has a peak turbidity when the two components are under charge matching conditions, and the turbidity values decrease when the positive and negative charge concentrations are unequal until only a singular phase system exists.^79,88^ The intensity of turbidity values are semi-quantitative that can depend on the shape, size, and density of the droplets. However, the range of RNA concentrations and the RNA concentration at maximum turbidity provides valuable insight.^89^ Tau(291-391) contains 15 lysine and 2 arginine residues which undergo multi-valent binding with the phosphates of poly(U), driving complex coaceravation.^78^ We expected that P301S and N-glycosylation, two neutral PTMs, would have no effect on LLPS. In contrast, acetylation and phosphorylation were expected to alter the peak turbidity and decrease the range of RNA concentrations for LLPS because they (a) decrease the number of binding sites (acetylation) or weaken the interaction strength (both phosphorylation and acetylation), (b) lower the isoelectric point, and (c) have less net positive charge.^78,88,90^

To validate these hypotheses, we generated a phase diagram of Tau with poly(U) (Figure 6a). We titrated a fixed concentration of Tau (30 µM) with poly(U) ranging from 0 to 110 µg/mL, and measured absorbance at 350 nm as an indicator of LLPS. The phase diagram revealed that WT Tau(291-391) has a maximum turbidity value around 50 µg/mL RNA, and it spans the range of 0-100 µg/mL poly(U). Acetylation at K311 and K317 has only small effects on the phase diagram compared to the WT protein, but AcK353 has a reduced two-phase window, and AcK369 has completely suppressed LLPS. Similar to our findings on monoacetylation at AcK311 and AcK317, mono-ubiquitination at K311 and K317 was reported to have minor effects on K18 LLPS with poly(U), based on droplet size vs time analysis.^91^ Phosphorylation of Y310 is similar to WT, while S305 and S356 shift the peak turbidity value to 40 µg/mL and 30 µg/mL, and narrows the phase diagram, while pS352 suppresses LLPS. Single N-glycosylation (GlcNAc) completely suppressed complex coacervation. Furthermore, the addition of AcK353 or pS305 on the same peptide chain also does not rescue droplet formation ability. The P301S mutation displays a similar two-phase window as the unmodified peptide, however, the peak turbidity value is shifted to 30 µg/mL poly(U). In conclusion, our findings demonstrate that phosphorylation reduces LLPS, while acetylation can have a positive or negative effect. This implies that certain residues are crucial for RNA binding, and PTMs can impair these LLPS interaction sites. While some PTMs, like AcK369, pS352, and GlcNAcN359, negatively affect RNA binding, others show moderate or no significant impact on the phase diagram.

### G. Salt Resistance

Tau displays a decreased tendency to undergo LLPS at increasing salt concentrations because the salt competes for the electrostatic attraction between ammonium or guanidinium sites to the phosphate groups of RNA.^77,79,84,92,93^ We hypothesized that PTMs exacerbate site-specific effects related to denaturation with salt, and some PTMs, even with the same charge, may exhibit unequal LLPS propensities. The turbidity of WT and 40 µg/mL poly(U) decreased by half in the presence of 20 mM NaCl, and a similar trend occurred for P301S and AcK317 (even though one of the lysine residues is unavailable for binding with RNA) (Figure 6B). Droplets produced from AcK311 and AcK353 were less stable towards salt, and the turbidity was diminished at 20 mM and 15 mM NaCl, respectively. Phosphorylation at S356 and S305 leads to less salt resistance than the WT peptide, and turbidity diminished to ∼25% at 20 mM NaCl. Among all the proteoforms studied, the pY310 was the least salt-resistant, and LLPS was diminished at 10 mM NaCl. These data suggest that PTMs have differential effects on the salt resistance of LLPS of Tau, and some regions are more important for maintaining the charge in high salt solutions. The droplets of AcK311 and AcK353 diminished quickly, suggesting that these residues may play a crucial role in maintaining RNA binding. The pY310 proteoform has a similar phase diagram as WT, but LLPS is rapidly diminished with salt, which shows the unique effect of this phosphorylated residue together with its aromatic character.

### H. Analysis of Droplet Size Over Time in Relation to PTMs

Our investigations also focused on examining how PTMs influence the morphology and dynamics of liquid phase-separated droplets associated with Tau. By employing time-lapse microscopy, we were able to track changes in the total area covered by these droplets over time, with the results illustrated in Figure 6c. The droplets demonstrated liquid-like properties, including droplet fusion and Ostwald ripening, and the area they covered on the microscopy slide plateaued after 1 hour. Interestingly, the coverage area was consistent across most constructs, with the notable exceptions being the WT, which covered a larger area, and the AcK317 and pY310 constructs, which covered less. This suggests variations in the efficiency of monomer to droplet conversion or differences in droplet density—with WT droplets being less dense than those formed by AcK317 and pY310. By the 2h mark, the droplets began to exhibit distinct morphologies: AcK353, pS305, pS356, and pY310 predominantly formed circular droplets; WT and P301S showed a mixture of circular and amorphous droplets; AcK317 and AcK311 mostly produced smaller, amorphous droplets, as depicted in Figure 6d. Further analysis at this time point focused on the average area of the droplets (Figure 6c), revealing that WT droplets were the largest, with an average size of 35 µm^2^, while AcK317 and pY310 formed the smallest droplets, averaging 15 µm^2^. Droplets from the other constructs had an intermediate average size of about 25 µm^2^. These observations underscore the significant role of PTMs in modulating both the morphology and the apparent density of Tau liquid droplets. Regardless of their shape—be it circular or amorphous—all droplets retained a liquid-like appearance. Their size steadily increased until a stable state was achieved approximately two hours into the observation period, as the droplets settled on the slide surface. An anomaly was noted with the pY310 droplets, which exhibited a tendency to compress rather than settle, highlighting the unique behaviors induced by specific PTMs.

### I. Droplet Dynamics

We next explored whether amyloid fibril nucleation occurs during the aging of Tau(291-391)–RNA droplets, and how PTMs might differentially affect this process. Liquid phase separated droplets of intrinsically disordered proteins can undergo aging into gel-like structures, Maxwell glasses, or amyloid fibrils.^76,79,94^ However, the specific trajectory of Tau(291-391) during LLPS with RNA, particularly whether it undergoes fibrillation, was unknown. Pathogenic Tau mutations are known to facilitate the aging of liquid droplets into higher molecular weight species, but amyloid fibril nucleation typically requires polyanionic cofactors.^82,85,95^ Conversely, LLPS driven by hydrophobic interactions, such as those induced by high salt or hyperphosphorylation, is known to trigger amyloid fibril formation.^86,87^

To investigate these aspects of LLPS, we employed Fluorescence Recovery After Photobleaching (FRAP) to probe the liquid-solid transition during complex coacervation.^96^ Additionally, we used negative stain TEM to check for the presence of amyloid fibrils (Figure 6e-i). FRAP assessments conducted one hour into the experiment showed that Tau-RNA droplets from WT, P301S, AcK353, AcK317, pS356, and pS305 predominantly maintained their liquid state, demonstrating near-complete fluorescence recovery. Conversely, droplets from AcK311 and pY310 exhibited diminished dynamics, as reflected by a reduced mobile fraction (Figure 6f). After two hours, a decrease in mobility was noted for WT, P301S, pY310, and AcK311 droplets, suggesting gradual solidification. Nonetheless, droplets from AcK353, AcK317, pS305, and pS356 retained similar mobility, suggesting persistent liquidity. Further analysis using negative stain TEM after two hours revealed no fibril formation at this stage (Figure 6h-i). However, sparse fibril density was observed in several constructs after three days, including WT, P301S, AcK369, GlcNAcN359, and GlcNAcN359+AcK353. This suggests that while WT, P301S, AcK311, and pY310 droplets show a decrease in mobility over time, indicative of gelation or Maxwell glass formation, this does not necessarily lead to amyloid fibril formation, as no fibrils were visible after three hours. This observation underscores the notion that droplet formation and fibrillation are distinct processes, offering insights into Tau behavior during phase transitions and the complex interplay between LLPS and protein aggregation.^79^

## DISCUSSION

In this study, we explored the mechanisms behind Tau filament formation by examining the behavior of Tau protein segments and their interaction with post-translational modifications. The Tau protein in AD brains exists in three distinct forms: a normal cytosolic form, a soluble hyperphosphorylated form, and an insoluble paired helical filament form.^53^ Furthermore, the Tau from AD patients is also known to contain other post-translational modifications, and it is not understood how PTMs regulate the (*in vitro*) formation of AD filaments, or which phase transformations are relevant for PHF formation on full-length Tau. The paired helical filaments from AD patients consist of a rigid core surrounded by a “fuzzy coat,” which can be removed to reveal the proteolytically stable segment Tau(297-391).^55,56^ This finding is critical as it suggests a specific region of Tau (I297-E391 and L266-E391, corresponding to 4R and 3R Tau, respectively) is capable of maintaining its assembly without any cofactors.^97–101^ The morphology of these self-assembled filaments closely resembles that of those extracted from brain tissue, as demonstrated by negative stain TEM and atomic force microscopy.^102–105^ The observation that Tau(297-391) can self-assemble into filaments that matches the ones isolated from AD and CTE patients was further confirmed with cryoEM; however, the structures are largely dependent on the shaking speed and buffer composition.^14^ ^16^ To investigate the role of PTMs in filament formation, we developed a three-segment chemical synthesis of Tau(291-391), a promising region for self-assembly. This approach allowed us to produce thirteen unique proteoforms with site-specific PTMs, including phosphorylation, acetylation, N-glycosylation, and a point mutation (Figures 1 and 2). The solution phase behavior of these proteoforms was largely identical. However, significant differences were observed as the proteoforms were tested under various phase transition conditions. Tau fibril formation involves a liquid to solid phase transition known as nucleation, followed by secondary processes that catalyze further fibril nucleation.^106^ Although Tau aggregation can be accomplished under cofactor-induced and cofactor-free conditions, the exact phase transitions, and the role of oligomeric species or liquid-phase separated condensates remain complex and not fully understood.^107–111^ The essential role of polyanionic cofactors in the formation, stabilization, and aggregation kinetics of poly-anion induced filaments has been established,^112–116^ yet in vitro filament preparations with polyanions do not replicate the cryo-EM structures of filaments from human patients.^10,11^ Our findings emphasize the complexity of Tau aggregation and the significant impact of PTMs on filament formation. The WT, AcK311, and AcK317 constructs demonstrated self-assembly, but only WT fibrils formed PHF filaments under previously optimized conditions (confirmed through a cryo-EM structure that matched those isolated from AD brains).^14,16^ The self-assembly is facilitated by a Tau-Tau binding, which exhibits nanomolar affinities and are correlated with positional specificity and PTM identity.^117^ Interestingly, the AcK311 and P301S mutations accelerated heparin-induced aggregation, whereas AcK353, AcK369, AcK317, pS356, pS352, GlcNAcN359, and GlcNAcN359+AcK353 inhibited aggregation. Moreover, pS305 and pY310 mutations completely suppressed self-assembly with heparin, underscoring the critical role of specific PTMs in regulating Tau filament formation.

LLPS may also be an important intermediate phase that concentrates Tau from dilute solution and enables nucleation or oligomerization. Phase separation is related to aggregation as it occurs under both polyanion-induced and cofactor-free conditions.^77,78,82–88^ We established that Tau(291-391) undergoes RNA-induced liquid-liquid phase separation, and the AcK369, pS352, and GlcNAcN359 modifications completely suppressed the LLPS. RNA-induced LLPS does not help promote the fibrilization of Tau, rather gelation occurs during this step, and RNA-promoted aggregation is a separate process. It remains under debate whether Tau phase separation can drive the formation of fibrils and oligomers, or if it is an off pathway phase transformation that occurs under similar conditions.^79,85,86^ Our work indicates that this process may be controlled by specific PTMs, as the pathway for the aggregation or unproductive LLPS is dependent on the PTM identity and even position on Tau. These observations add to the complexity of the mechanism of self-assembly into PHFs.

The strength of our study lies in its ability to investigate the effects of site-specific PTMs on Tau aggregation. We have demonstrated that the specific residue carrying a post-translational modification can influence Tau aggregation positively, as evidenced by AcK311. Conversely, the same type of modification at a different site, such as AcK369, can have a negative impact. This finding underscores the complex yet significant role of PTMs in Tau aggregation. One interpretation of this data is that if the PTMs are on the full-length protein, the PTMs within the ordered filament core may aid in the self-assembly of full length hyperphosphorylated Tau. An alternative interpretation is that AD specific PTMs within the ordered filament core are present on insoluble Tau PHFs because they are at conformationally accessible residues and become modified because of reduced Tau clearance. A limitation of our approach is the use of a Tau fragment rather than the full-length protein, which is more representative of clinical conditions. Notably, several PTMs are located within the ordered filament core in AD patients, but many are also found outside this region.^31^ Furthermore, our attempts to study the interplay of two PTMs represents only a fraction of possible (and likely) combinations of PTMs in an individual Tau chain. In the disease state, the filaments are heavily modified with PTMs,^31^ and the future studies on Tau constructs that contain higher modification stoichiometry may provide a more comprehensive analysis of the interplay between the PTMs.

In conclusion, our work has advanced the understanding of Tau protein aggregation by demonstrating that site-specific PTMs can have varying effects on the aggregation process, with some facilitating and others inhibiting the formation of PHFs. These insights into the RNA-induced LLPS indicate that it does not contribute to the nucleation of AD filaments, emphasizing the role of environmental factors and PTMs located outside the ordered filament core in Tau self-assembly. Our findings lay the groundwork for future studies on full-length and hyperphosphorylated Tau constructs and the influence of higher modification stoichiometries on Tau aggregation, which could further elucidate the pathogenesis of Alzheimer’s disease and related tauopathies.

## METHODS

### Peptide synthesis and purification

Peptide synthesis was accomplished with the standard Fmoc-based chemistries using automated peptide synthesizers (room temperature or microwave-assisted coupling). The crude peptides obtained after resin cleavage were purified by preparative HPLC and lyophilized. The identity and purity of the peptides were confirmed by LC/HRMS.

### CD spectroscopy

Lyophilized Tau(291-391) was dissolved in 20 mM sodium phosphate buffer (pH 7.4), and the residual TFA salts were removed using a 7,000 MWCO Slide-A-Lyzer MINI Dialysis Unit (Thermo Scientific, Cat. No. 69562). The proteins were adjusted to 0.4 mg/mL in 20 mM sodium phosphate buffer (pH 7.4) and analyzed in triplicate by CD spectroscopy. The triplicate CD data were averaged and reported as molar eplicity.

### Co-factor free self-assembly

This protocol was adapted from the literature.^14^ Lyophilized Tau(291-391) was dissolved in 10 mM sodium phosphate, 10 mM DTT (pH 7.4) and purified by SEC over a Superdex 75 10/300 GL into 10 mM sodium phosphate, 10 mM DTT (pH 7.4). The fractions containing protein were concentrated to 8 mg/mL using a Pierce concentrator (PES, 3K MWCO, 0.5 mL, 88512).

The cofactor free self-assembly was run in a 96 well plate (Greiner BioOne, 96 well, PS, F Bottom, Chimney well, black, medium binding, 655096). The wells were filled with 100 µL of the sample with 4 mg/mL Tau(291-391), 200 mM MgCl_2_, 10 mM DTT, 10 mM sodium phosphate (pH 7.4). The plate was sealed with an adhesive film (VWR, polyester foil, 89134-430) and mixed at 37 ⁰C at 300 RPM on a thermomixer (Ika Matrix Orbital). After 48 h, the aggregation mixtures were directly assayed for fibrils by negative stain TEM.

### Heparin protein aggregation

A 0.65 µL Eppendorf tube was charged with 50 µL of aggregation assay mixture at the final concentration listed. The buffers and salts were added first, followed by protein, and heparin was added last. Three wells on a black, 384-well, non-binding microplate (Greiner BioOne, PS, F Bottom, small volume, HiBase, 784900) were filled with 15 µL of the aggregation assay mixture, the plate was sealed with polyester adhesive film (VWR, 89134-430), and the plate was incubated at 37 ⁰C into the microplate reader, monitoring the fluorescence (Ex/Em: 440/480 nm). The data from the three wells were averaged, and the experiment was repeated for a total of three times. The data was averaged then normalized, and the error bars on the graphs are reported as the standard error of the mean.

### ELISA

This protocol was taken from literature with minor modifications.^117,118^ The difference with our protocol is that we used 2N4R Tau(2-441) in the solution phase instead of Tau(297-391), and we used anti-Tau, 15-25 mouse antibody (binds residues 15-25 on 2N4R Tau, BioLegend, 835201) as the primary antibody instead of mAb 423, and anti-mouse IgG (H+L), HRP conjugate (Promega, W4021) as the secondary antibody.

### Phase diagram and FRAP

Turbidity phase diagrams were measured on a NanoDrop2000 instrument. The absorbance was measured at 350 nm with a path length of 0.1 cm. A 0.65 µL Eppendorf tube was charged with 3 µL of 2x protein stock. Next, 3 µL of 2x poly(U) RNA stock solution was added, and the mixture was pipetted up and down several times. A minute after mixing, 2 µL of the sample was transferred to Nanodrop, and the absorbance was measured three times. Using the same LLPS mixture, this was repeated twice. This experiment was performed in triplicate, and the data from the three experiments was averaged and reported as the standard error of the mean.

### Salt resistance

The salt resistance assay was measured by turbidity on a NanoDrop2000 instrument. The absorbance was measured at 350 nm with a path length of 0.1 cm. A 0.65 µL Eppendorf tube was charged with 3 µL of 3x protein stock. The mixture was treated with 1.5 µL of 6x NaCl stock. Next, 4.5 µL of 2x RNA stock was added and pipetted up and down several times. A minute after mixing, 2 µL of the droplet solution was transferred to Nanodrop, and the absorbance was measured three times. Using the same LLPS mixture, this was repeated twice. This experiment was performed in triplicate, and the data from the three experiments was averaged and reported with error bars that represent the standard error of the mean.

### Droplet size measurements by fluorescence microscopy

A 0.65 µL Eppendorf tube was charged with 3 µL of 2x protein. Next, 3 µL of 2x poly(U) RNA stock was added, and the mixture was pipetted up and down several times. The sample was transferred to the center of the micro-well in a 35 mm glass bottom dish with 14 mm micro-well. The edge of the microwell was lined with 25 mM HEPES (∼12 µL) to provide an evaporation shield, the microwell was covered with a glass coverslip, and the chamber was sealed with nail polish. For each droplet preparation, five random locations were imaged in 10-minute intervals. The experiment was repeated for a total of 3 or 4 independent experiments.

### FRAP

A 0.65 µL Eppendorf tube was charged with 3 µL of 2x protein stock. Next, 3 µL of 2x poly(U) RNA stock was added, and the mixture was pipetted up and down several times. The sample was transferred to the center of the micro-well in a 35 mm glass bottom dish with 14 mm micro-well. The edge of the microwell was lined with 25 mM HEPES (∼12 µL) to provide an evaporation shield, and the microwell was covered with a glass coverslip, and the chamber was sealed with nail polish. For each droplet preparation, one droplet is photobleached after 1 h, and another droplet was photobleached after 2 h. The experiment was repeated for a total of 5 independent experiments.

### Assembly of 291-391 (cryo-EM)

Lyophilized samples were resuspended in 1 mL of 10 mM PB pH 7.4 10 mM DTT and left for 20 minutes at RT. Samples were then centrifuged (13 000 rpm for 5 minutes 20 °C) prior to size exclusion chromatography (HiLoad Superdex 200 pg, Cytivia). Fractions containing protein were verified by SDS-PAGE (4-20% Tris Glycine), pooled and concentrated to 8 mg/ml using molecular weight concentrators with a cutoff filter of 3 kDa. Protein concentrations were determined using a NanoDrop2000 (Thermo Fisher Scientific). 291-391 was diluted to 5 mg/ml containing 10 mM phosphate buffer at pH 7.2, 100 mM MgCl_2_ and 10 mM DTT and assembled in a 384-well microplate that was sealed and placed in a Fluostar Omega (BMB Labtech) with 200 rpm shaking at 37 °C for 24 hrs.

### Cryo-EM

After 24 hours, the sample was taken directly from the microplate and 3 μL of the reaction mixture were applied to glow-discharged R1.2/1.3, 300 mesh carbon Au grids. The grids were plunge-frozen in liquid ethane using a Vitrobot Mark IV (Thermo Fisher Scientific). Cryo-EM images were acquired on a Krios G2 (Thermo Fisher Scientific) electron microscope that was equipped with a Falcon-4 camera (Thermo Fisher Scientific). Images were recorded at a dose of 30 electrons per square ångström using EPU software (Thermo Fisher Scientific) and converted to tiff format using relion_convert_to_tiff.

### Cryo-EM data processing

Movie frames were gain corrected, aligned and dose weighted using RELION’s motion correction program. Contrast transfer function (CTF) parameters were estimated using CTFFIND-4.1. Relion helical reconstruction was carried out using RELION-4.0. Filaments were picked manually and extracted in a box size of 768 pixels down-sampled to 128 pixels for initial 2D classification. Good classes were selected and re-extracted in a particle box size of 384 pixels with no down-sampling for 3D auto-refinements. After several rounds of 3D auto-refinements optimising for helical parameters, Bayesian polishing and CTF refinement were used to further increase the resolution. Final maps were sharpened using standard post-processing procedures in RELION. Reported resolutions were estimated using a threshold of 0.143 in the Fourier shell correlation (FSC) between two independently refined half-maps.

### Negative stain TEM of Tau filaments

Filaments were negatively stained with 3% uranyl acetate and imaged on an FEI Tecnai T12 Spirit at 120 kV with a LaB6 filament). 3 replicates of each group were imaged and 10 random images at 30,000x (2.119 nm/pixel) were obtained per replicate. Data from these replicates was compared and then pooled per group for subsequent analysis by FibrilJ.

### Analysis of fibrillar length and diameter

Tau(291-391) fibrils were analyzed using the ImageJ fibril analysis plugin, FibrilJ.^119^ A modified version of FibrilJ, incorporating an enhanced skeleton pruning algorithm, was employed to streamline the fibrils into lines and reduce the occurrence of branches erroneously generated during skeletonization. Before analysis, images underwent binarization via custom segmentation algorithms. Among the three diameter algorithms provided by FibrilJ, results from the D2 (human-like) algorithm are highlighted in the results section, as the D1 (area/length) and D3 (distance map) algorithms yielded inconsistent calculations. FibrilJ’s ability to accurately determine fibril lengths is constrained by the inherent limitations of the ImageJ 2D/3D Skeleton program and the AnalyzeSkeletons-based pruning plugin. In the pruning process, FibrilJ trims the shortest branches, sometimes inadvertently dividing fibrils at arbitrary points along their length. This issue is particularly prevalent when fibrils intersect or overlap, or when staining is uneven, affecting the binary image quality. Consequently, FibrilJ may erroneously interpret a single fibril as multiple entities, reporting the lengths of various segments rather than the entire fibril. The fibril properties are then exported to an Excel sheet, listing IDs for each fibril detected and measured by FibrilJ. To enhance the accuracy of the length data, we manually reviewed the length data for the fibrils, identifying and correcting for falsely detected fibrils and merging different fibril IDs that constitute a single fibril.

To improve the precision of fibril length measurements, we opted for images with lower density and minimal fibril overlap, along with areas of optimal negative (or positive) staining. This approach facilitates more uniform image binarization and minimizes the risk of fibril bisection due to variations in stain intensity across the image.

Data from three fibril replicates within each group were aggregated for inter-group comparison. The Kolmogorov-Smirnov test was employed to assess the length and diameter distributions across different fibril types.

## ABBREVIATIONS

Ac – Acetyl; Acm – Acetamidomethyl; AD – Alzheimer’s Disease; Bn – Benzyl; CBD – Corticobasal Degeneration; CC – Complex Coacervation; CD – Circular Dichroism; cryo-EM – cryo-Electron Microscopy; CTE – Chronic Traumatic Encephalopathy; DIC – Diisopropylcarbodiimide; DIPEA – Diisopropylethylamine; DMSO – Dimethyl Sulfoxide; DPDS – Diphenyldiselenide; DSL – Diselenide Selenoester Ligation; ELISA – Enzyme-linked Immunosorbent Assay; Fmoc – Fluorenylmethyloxycarbonyl; FRAP – Fluorescence Recovery After Photobleaching; FTDP-17 – Frontotemporal Dementia with Parkinsonism-17; GGT – Globular Glial Tauopathy; GlcNAc – N-Acetylglucosamine; Gnd – Guanidine; HATU – O-(7-azabenzotriazol-1-yl)-N,N,N’,N’-tetramethyluronium hexafluorophosphate; HEPES – 4-(2-Hydroxyethyl)-1-piperazineethanesulfonic acid; HMPB – 4-(4-Hydroxymethyl-3-methoxyphenoxy)butyric acid, polymer-bound; HPLC – High-Performance Liquid Chromatography; LC/HRMS – Liquid Chromatography/High-Resolution Mass Spectrometry; LLPS – Liquid-liquid Phase Separation; Me – Methyl; MPAA – Mercaptophenylacetic Acid; NCL – Native Chemical Ligation; NFTs – Neurofibrillary Tangles; Oxyma – Ethyl cyanoglyoxylate-2-oxime; P(*n*-Bu)_3_ – Tri-n-butylphosphine; PEG – Polyethyleneglycol; PHFs – Paired Helical Filaments; PMB – Para-methoxybenzyl; PSP – Progressive Supranuclear Palsy; PTMs – Post-translational Modifications; RNA – Ribonucleic Acid; SPPS – Solid-phase Peptide Synthesis; TFA – Trifluoroacetic Acid; TEM – Transmission Electron Microscopy; Thz – Thiazolidine; WT – Wild Type.

## ACKNOWLEDGEMENTS

This work was supported by the National Institute on Aging (AG079294 to M.A.W. and AG061829 to M.H.B.S), the MCDB Neurodegenerative Disease Fund (to M.H.B.S.), and the University of Colorado at Boulder. Mass spectrometry analyses were performed at the Proteomics and Mass Spectrometry Core Facility in the department of Biochemistry at the University of Colorado Boulder. Negative stain electron microscopy was done at the University of Colorado, Boulder EM Services Core Facility in the MCDB Department, with the technical assistance of facility staff. Cryo-electron microscopy was done at the MRC Laboratory of Molecular Biology, with help from Sofia Lövestam and Sjors Scheres. The droplet imaging work was performed at the BioFrontiers Institute Advanced Light Microscopy Core (RRID: SCR_018302). Laser scanning confocal microscopy was performed on an Nikon A1R microscope supported by NIST-CU Cooperative Agreement award number 70NANB15H226. We thank the Shared Instruments Pool (RRID: SCR_018986) of the Department of Biochemistry at the University of Colorado Boulder for the use of the CD spectrometer. The CD is funded by NIH Shared Instrumentation Grant S10RR028036. Figures 1 and 4 were generated with the assistance of biorender.com, ProCart, and Chimera.

