## Supplementary Information for "Post-Translational Modifications Control Phase Transitions of Tau"

#### 1. GENERAL INFORMATION

All chemicals and solvents were purchased as reagent grade and used as received without further purification unless otherwise noted. All peptide assembly reactions were carried out in screw top vials, in $N_2$ purged buffers. All aqueous buffers were filtered with a 0.22 μM syringe filter prior to use. Peptide synthesis is performed on a CEM Liberty Prime microwave peptide synthesizer, on a Biotage Initiator+Alstra, or on an AAPPTEC Focus SC peptide synthesizer. Analytical HPLC was performed on an Agilent 1260 Infinity II series HPLC or Waters Aquity UPLC. Preparative HPLC was performed on a Teledyne ISCO ACCQ Prep HP125. Low resolution mass was recorded on an Advion Expression L with ESI+ ionization. High-resolution mass spectra (HRMS) were recorded on a Waters Synapt G2 HDMS q-TOF hybrid mass spectrometer. Absorbance at A280 and turbidity is measured on the Nanodrop 2000 (Thermo Scientific). Proteins were analyzed by CD with a Modular Applied Photophysics Chirascan Plus CD spectrometer. Light microscopy was collected on a Nikon Eclipse Ti laser scanning confocal microscope and 60x oil immersion objective. A SpectraMax iD5 plate reader (Molecular Devices) was used for all microplate assays. Negative stain TEM images were taken on a FEI Tecnai T12 Spirit at 120 kV with a LaB6 filament. Fibril J software was used for the analysis of fibril diameters and lengths. Fiji/Image J is used for all image analysis, and Prism/Graphpad was used for all graphical, and statistical analysis.

#### 2. D358-E391 (SPPS)

### WT

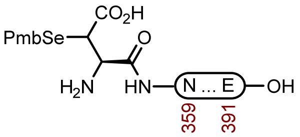

**S1a**

**Sequence**: H-Asp(SePmb)NITHVPGGGNKKIETHKLTFRENAKAKTDHGAE-OH

**Amino Acids**: Boc-L-(SePmb)Asp(O $^t$Bu)-OH,[1] Fmoc-L-Asn(Trt)-OH, Fmoc-L-Ile-OH, Fmoc-L-Thr( $^t$Bu)-OH, Fmoc-L-His(Trt)-OH, Fmoc-L-Val-OH, Fmoc-L-Pro-OH, Fmoc-Gly-OH, Fmoc-L-Lys(Boc)-OH, Fmoc-L-Glu(O $^t$Bu)-OH, Fmoc-L-Leu-OH, Fmoc-L-Phe-OH, Fmoc-L-Arg(Pbf)-OH, Fmoc-L-Ala-OH, Fmoc-L-Asp(O $^t$Bu)-OH

**Resin**: HMPB ChemMatrix resin from Biotage (100-200 mesh, 0.42 mmol/g, 238 mg, 0.1 mmol) was used on CEM. The resin was loaded by the symmetrical anhydride with Fmoc-L-Asp(OAllyl)-OH, DIC, and 4-DMAP.

**Coupling**: Amino acids were double coupled using 0.2 M amino acid in DMF (5 equiv.), 1 M DIC in DMF (10 equiv.), and 1 M oxyma pure in DMF (5 equiv.) at 90°C for 4 min. Histidine was double coupled at 50°C for 10 min.

**Deprotection**: 20% piperidine + 0.1 M oxyma pure in DMF (6 mL) at 23 ºC (1 x 5 min, 1 x 15 min)

**Global Deprotection**: The dry resin in a 15 mL polypropylene vial was treated with TFA:$H_2O$:thioanisole:phenol:TIPSH (85:5:5:2.5:2.5, 10 mL) and it was mixed on an overhead stirrer. The mixture was stirred at 23 ºC for 3 h before it was filtered, and the resin was washed with additional portion of TFA (3 x 1 mL). The pooled filtrate was cooled to 0°C and treated with $Et_2O$ to precipitate the peptide. The precipitate was centrifuged, and the resulting pellet was washed twice more with $Et_2O$. It was dissolved in 40% MeCN and it was lyophilized.

**Crude LCMS (SPPS)**: Measured on an Agilent EC-C18 Poroshell column (4.6 x 100 mm, 4 µm, 100 Å, 40 °C, 1 mL/min) 5%-65% MeCN/$H_2O$/0.05% TFA over 60 min.

**LC trace (214 nm) of unpurified peptide**

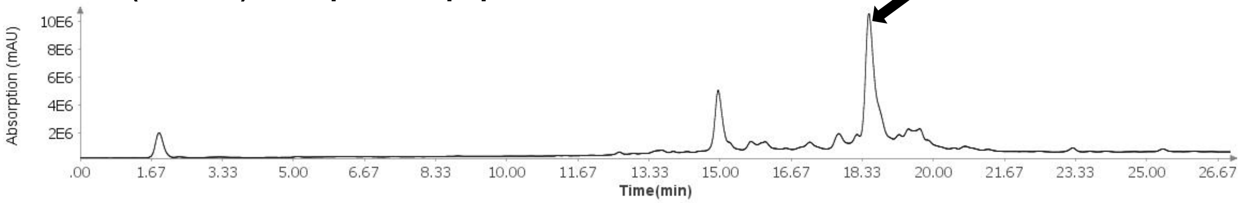

**Total Ion Current (TIC) of unpurified peptide**

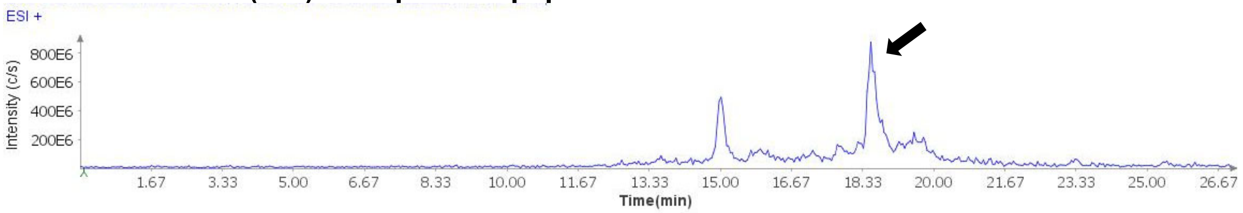

**Extracted Ion Current (XIC) of unpurified peptide**

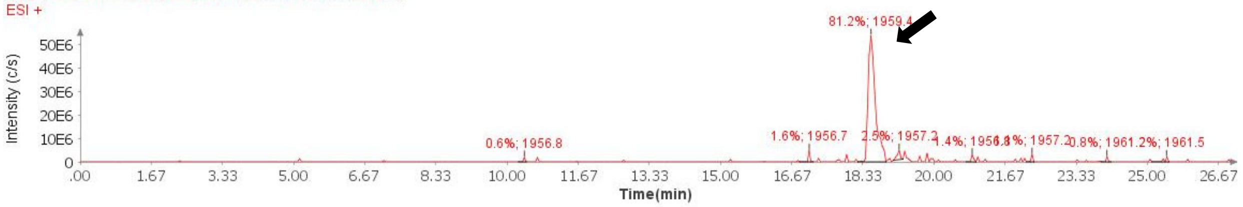

**Mass Spectrum (ESI) of the expected product (unpurified sample)**

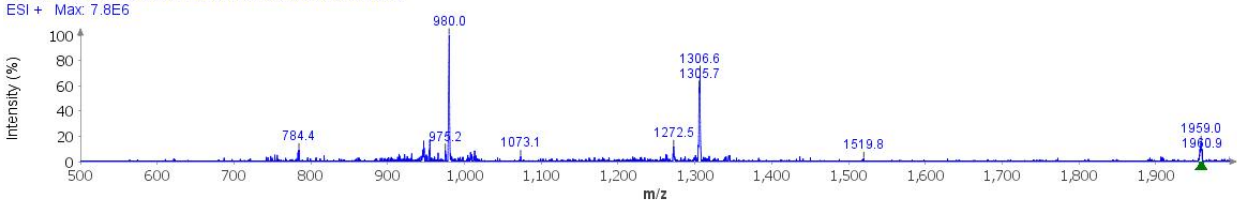

**Deconvoluted Mass Spectrum (ESI) of the expected product (unpurified sample)**

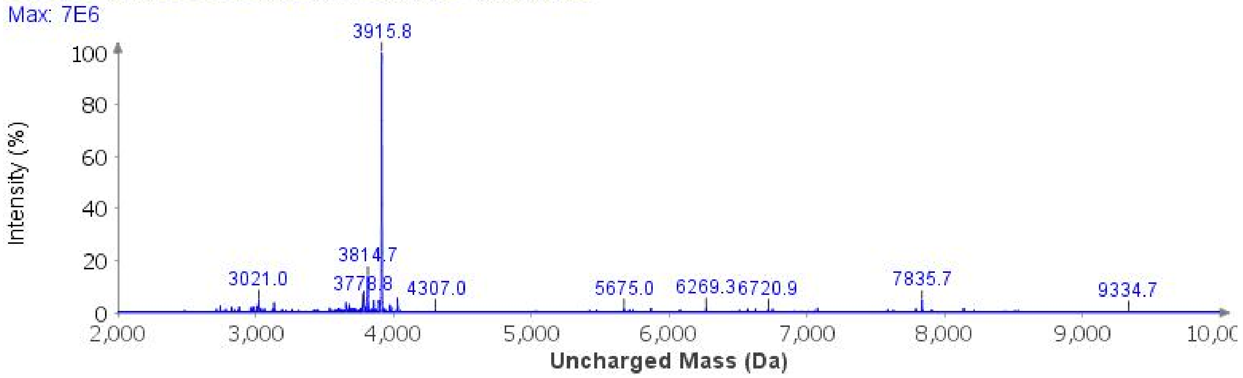

**Purification**: The material was purified over an ACQUITY Protein BEH C4 column (19 x 250 mm, 5 μm, 300 Å, 17.1 mL/min) 20%-40% MeCN/$H_2O$/0.1% TFA over 30 min. The fractions were lyophilized to afford **S1a** (41.1 mg, 11% yield).

**Analytical LCMS**: Measured on an Agilent EC-C18 Poroshell column (4.6 x 100 mm, 4 μm, 100 Å, 40 °C, 1 mL/min) 5%-65% MeCN/$H_2O$/0.05% TFA over 60 min.

**Chemical Formula**: $C_{167}H_{265}N_{51}O_{53}Se$
**Molecular Weight**: 3914.2320 g/mol

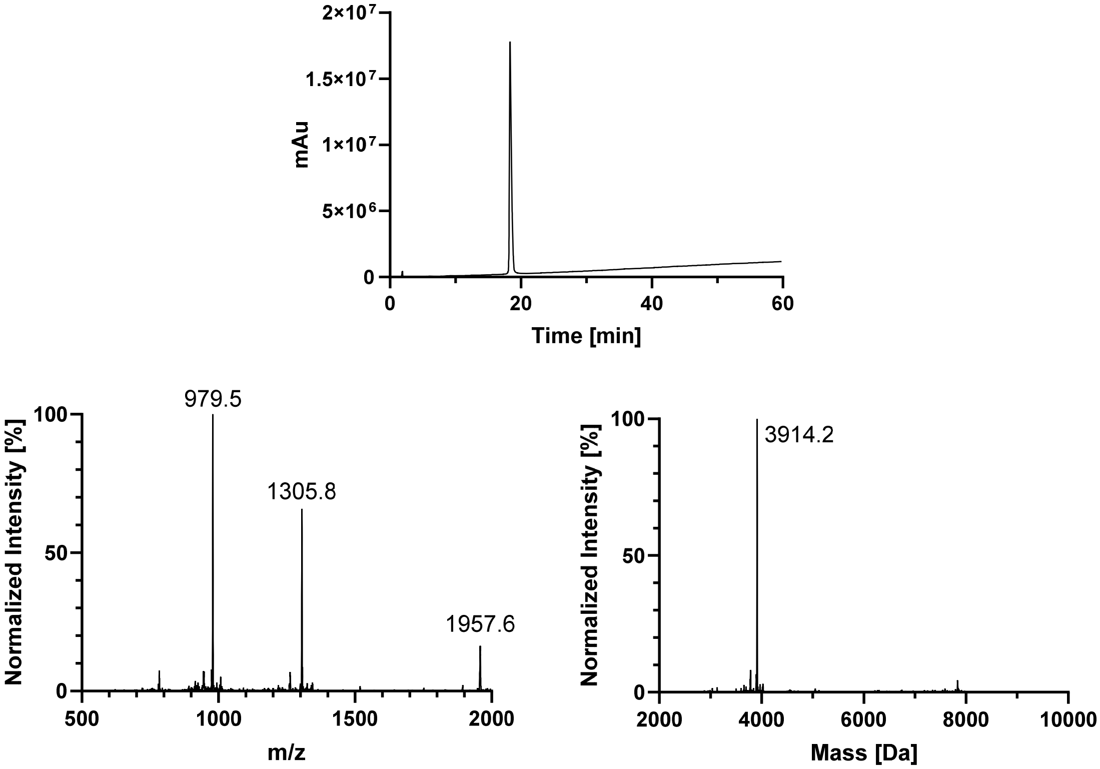
2×10^7
1.5×10^7
1×10^7
5×10^6
0
mAu
0
20
40
60
Time [min]
979.5
1305.8
1957.6
100
50
0
Normalized Intensity [%]
500
1000
1500
2000
m/z
3914.2
100
50
0
Normalized Intensity [%]
2000
4000
6000
8000
10000
Mass [Da]

#### AcK369

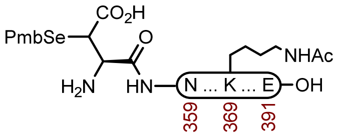

**S1c**

**Sequence**: H-Asp(SePmb)NITHVPGGGNK(Ac)KIETHKLTFRENAKAKTDHGAE-OH

**Amino Acids**: Boc-L-(SePmb)Asp(O $^t$Bu)-OH,[1] Fmoc-L-Asn(Trt)-OH, Fmoc-L-Ile-OH, Fmoc-L-Thr( $^t$Bu)-OH, Fmoc-L-His(Trt)-OH, Fmoc-L-Val-OH, Fmoc-L-Pro-OH, Fmoc-Gly-OH, Fmoc-L-Lys(Boc)-OH, Fmoc-L-Lys(Ac)-OH Fmoc-L-Glu(O $^t$Bu)-OH, Fmoc-L-Leu-OH, Fmoc-L-Phe-OH, Fmoc-L-Arg(Pbf)-OH, Fmoc-L-Ala-OH, Fmoc-L-Asp(O $^t$Bu)-OH

**Resin**: HMPB ChemMatrix resin from Biotage (100-200 mesh, 0.42 mmol/g, 238 mg, 0.1 mmol) was used on CEM. The resin was loaded by the symmetrical anhydride with Fmoc-L-Glu(O $^t$Bu)-OH, DIC, and 4-DMAP.

**Coupling**: Amino acids were double coupled using 0.2 M amino acid in DMF (5 equiv.), 1 M DIC in DMF (10 equiv.), and 1 M oxyma pure in DMF (5 equiv.) at 90°C for 4 min. Histidine was double coupled at 50°C for 10 min.

**Deprotection**: 20% piperidine + 0.1 M oxyma pure in DMF (6 mL) at 23 ºC (1 x 5 min, 1 x 15 min)

**Global Deprotection**: The dry resin in a 15 mL polypropylene vial was treated with TFA:$H_2O$:thioanisole:phenol:TIPSH (85:5:5:2.5:2.5, 10 mL) and it was mixed on an overhead stirrer. The mixture was stirred at 23 ºC for 3 h before it was filtered, and the resin was washed with additional portion of TFA (3 x 1 mL). The pooled filtrate was cooled to 0°C and treated with $Et_2O$ to precipitate the peptide. The precipitate was centrifuged, and the resulting pellet was washed twice more with $Et_2O$. It was dissolved in 40% MeCN and it was lyophilized.

**Crude LCMS (SPPS)**: Measured on an Agilent EC-C18 Poroshell column (4.6 x 100 mm, 4 µm, 100 Å, 40 °C, 1 mL/min) 5%-65% MeCN/$H_2O$/0.05% TFA over 60 min.

**LC trace (214 nm) of unpurified peptide**

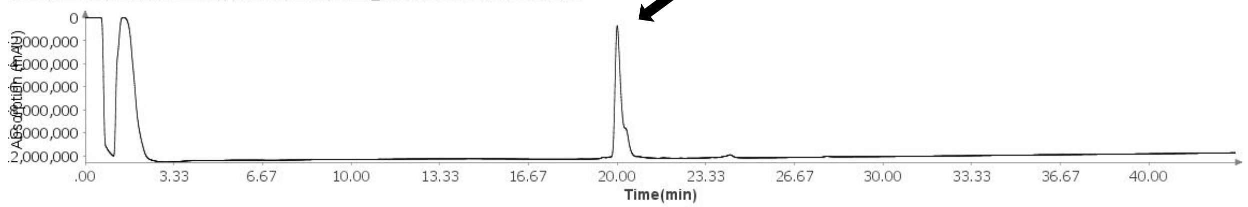

**Total Ion Current (TIC) of unpurified peptide**

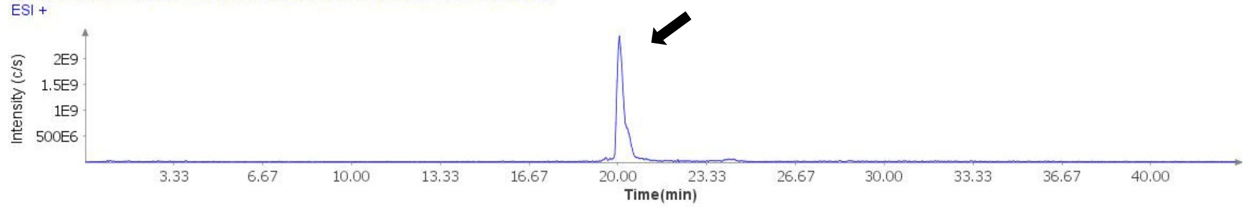

**Extracted Ion Current (XIC) of unpurified peptide**

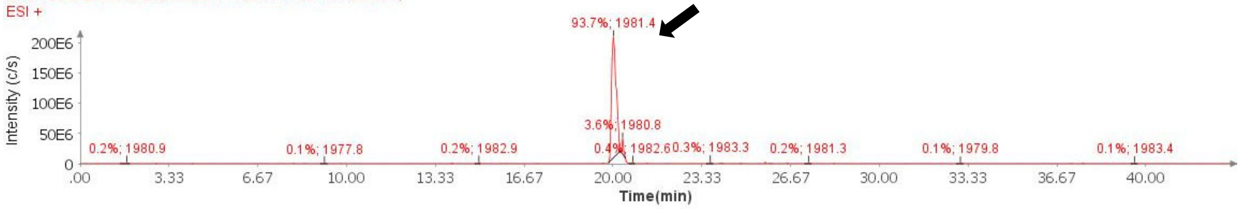

**Mass Spectrum (ESI) of the expected product (unpurified sample)**

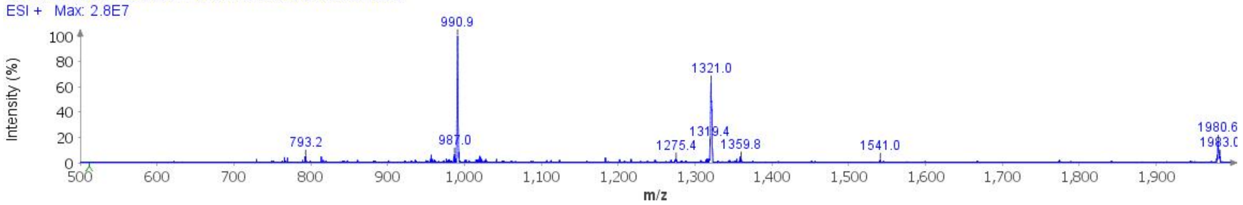

**Deconvoluted Mass Spectrum (ESI) of the expected product (unpurified sample)**

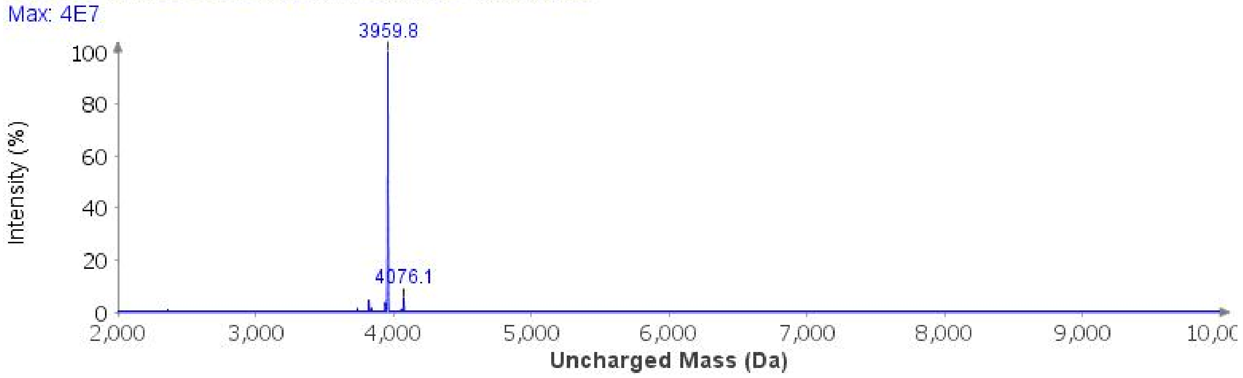

**Purification**: The material was purified over an ACQUITY Protein BEH C4 column (19 x 250 mm, 5 µm, 300 Å, 17.1 mL/min) 20%-40% MeCN/$H_2O$/0.1% TFA over 30 min. The fractions were lyophilized to afford **S1c** (53 mg, 13% yield).

**Analytical LCMS**: Measured on an Agilent EC-C18 Poroshell column (4.6 x 100 mm, 4 µm, 100 Å, 40 °C, 1 mL/min) 5%-65% MeCN/$H_2O$/0.05% TFA over 60 min.

**Chemical Formula**: $C_{169}H_{267}N_{51}O_{54}Se$
**Molecular Weight**: 3956.2690 g/mol

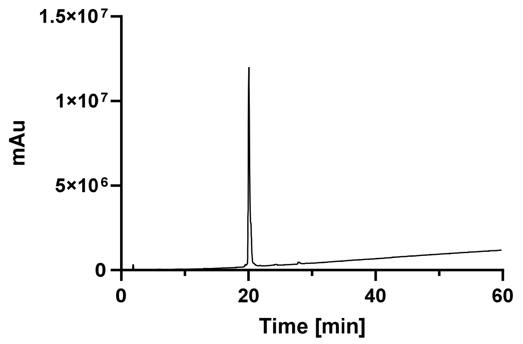

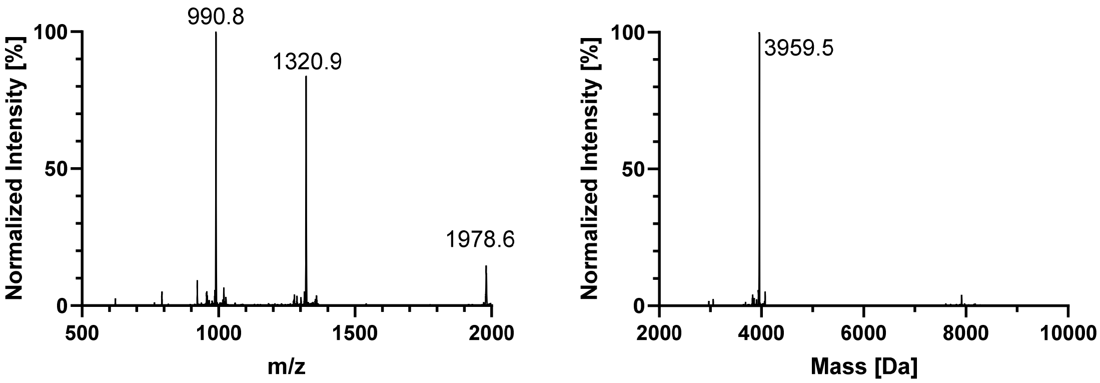
990.8
1320.9
1978.6
Normalized Intensity [%]
100
50
0
500
1000
1500
2000
m/z
3959.5
Normalized Intensity [%]
100
50
0
2000
4000
6000
8000
10000
Mass [Da]

#### GlcNAcN359

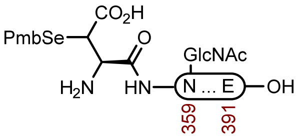

**S1k**

**Sequence**: H-Asp(SePmb)N(GlcNHAc)ITHVPGGGNKKIETHKLTFRENAKAKTDHGAE-OH

**Amino Acids**: Boc-L-(SePmb)Asp(O $^t$Bu)-OH,[1] Fmoc-L-Asn($Ac_3$GlcNHAc)-OH,[2] Fmoc-L-Ile-OH, Fmoc-L-Thr( $^t$Bu)-OH, Fmoc-L-His(Trt)-OH, Fmoc-L-Val-OH, Fmoc-L-Pro-OH, Fmoc-Gly-OH, Fmoc-L-Asn($Ac_3$GlcNHAc)-OH Fmoc-L-Lys(Boc)-OH, Fmoc-L-Glu(O $^t$Bu)-OH, Fmoc-L-Leu-OH, Fmoc-L-Phe-OH, Fmoc-L-Arg(Pbf)-OH, Fmoc-L-Ala-OH, Fmoc-L-Asp(O $^t$Bu)-OH

**Resin**: HMPB ChemMatrix resin from Biotage (100-200 mesh, 0.42 mmol/g, 238 mg, 0.1 mmol) was used on CEM. The resin was loaded by the symmetrical anhydride with Fmoc-L-Glu($^t$Bu)-OH, DIC, and 4-DMAP.

**Coupling**: Amino acids were double coupled using 0.2 M amino acid in DMF (5 equiv.), 1 M DIC in DMF (10 equiv.), and 1 M oxyma pure in DMF (5 equiv.) at 90 °C for 4 min. Histidine was double coupled at 50°C for 10 min.

**Deprotection**: 20% piperidine + 0.1 M oxyma pure in DMF (6 mL) at 23 ºC (1 x 5 min, 1 x 15 min)

**Global Deprotection**: The dry resin in a 15 mL polypropylene vial was treated with TFA:$H_2O$:thioanisole:phenol:TIPSH (85:5:5:2.5:2.5, 10 mL) and it was mixed on an overhead stirrer. The mixture was stirred at 23 ºC for 3 h before it was filtered, and the resin was washed with additional portion of TFA (3 x 1 mL). The pooled filtrate was cooled to 0°C and treated with $Et_2O$ to precipitate the peptide. The precipitate was centrifuged, and the resulting pellet was washed twice more with $Et_2O$. It was dissolved in 40% MeCN and it was lyophilized.

**Crude LCMS (SPPS)**: Measured on an Agilent EC-C18 Poroshell column (4.6 x 100 mm, 4 μm, 100 Å, 40 °C, 1 mL/min) 5%-65% MeCN/$H_2O$/0.05% TFA over 60 min.

**LC trace (214 nm) of unpurified peptide**

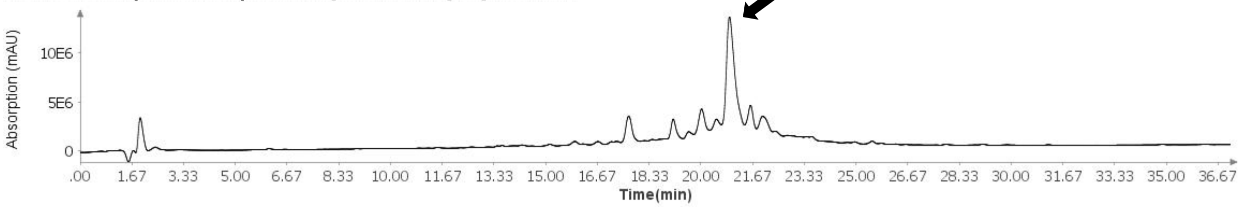

**Total Ion Current (TIC) of unpurified peptide**

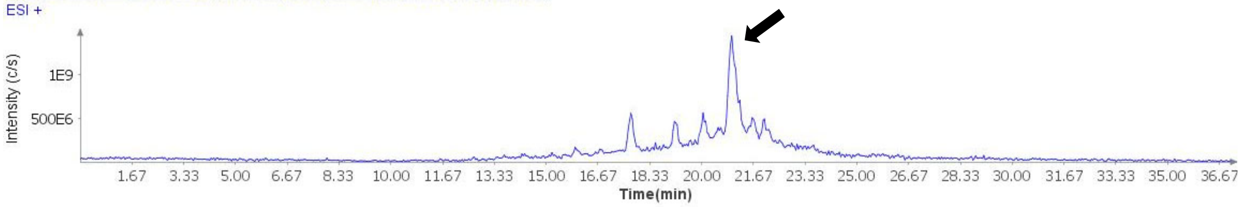

**Extracted Ion Current (XIC) of unpurified peptide**

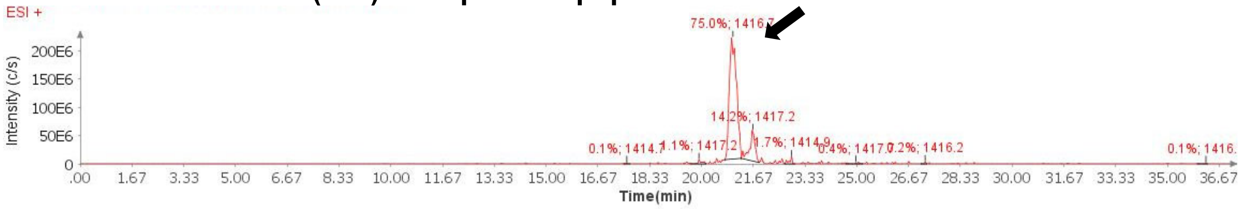

**Mass Spectrum (ESI) of the expected product (unpurified sample)**

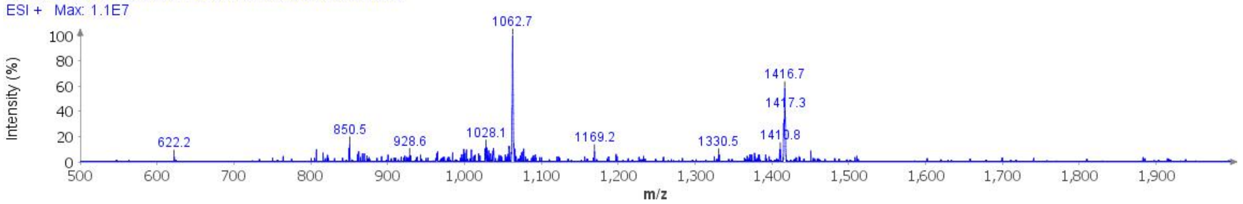

**Deconvoluted Mass Spectrum (ESI) of the expected product (unpurified sample)**

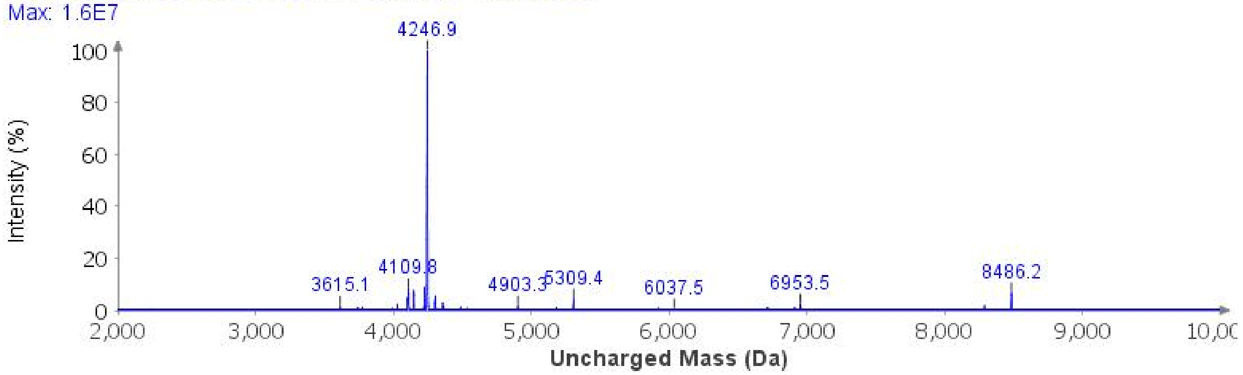

**Deacetylation**: The crude peptide was dissolved in hydrazine buffer (5 mL, 3% (v/v) hydrazine hydrate, 6 M GndHCl, 0.2 M $Na_2HPO_4$, pH 8.3). After 3 h, treated with an additional portion of hydrazine hydrate (20 µL) and after 4 h (40 µL). After 5 h, the reaction was purified over an ACQUITY Protein BEH C4 column (19 x 250 mm, 5 µm, 300 Å, 17.1 mL/min) 20%-40% MeCN/$H_2O$/0.1% TFA over 30 min. The fractions were lyophilized to afford **S1k** (76.2 mg, 18% yield).

**Crude LCMS (Deacetylation)**: Measured on an Agilent EC-C18 Poroshell column (4.6 x 100 mm, 4 µm, 100 Å, 40 °C, 1 mL/min) 5%-65% MeCN/$H_2O$/0.05% TFA over 60 min.

**LC trace (214 nm) of unpurified peptide**

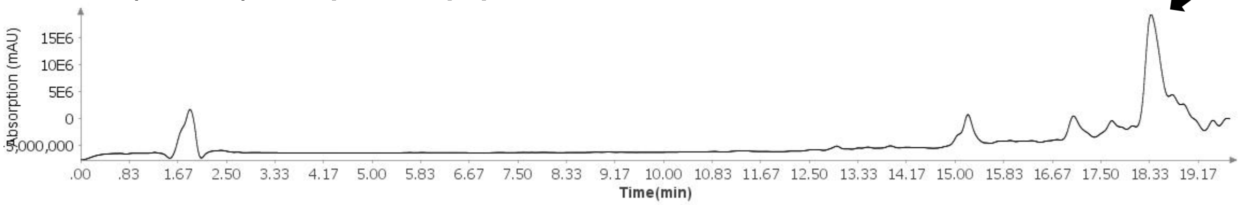

**Total Ion Current (TIC) of unpurified peptide**

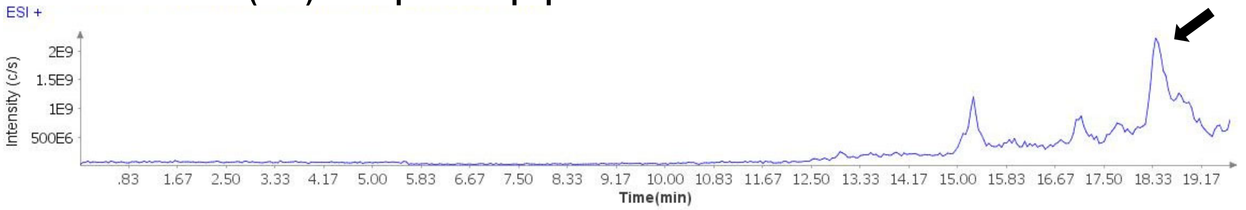

**Extracted Ion Current (XIC) of unpurified peptide**

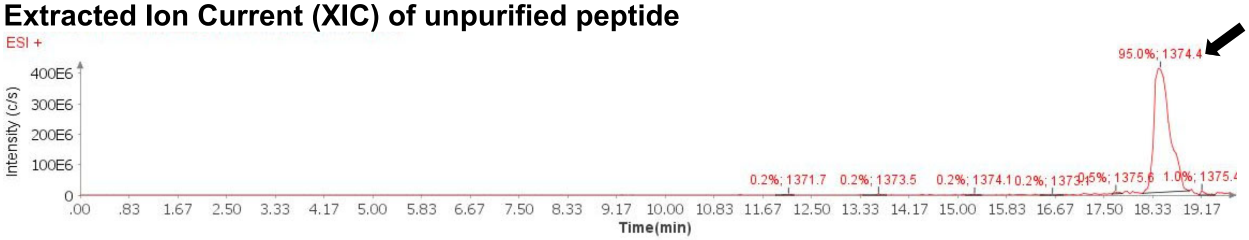

**Mass Spectrum (ESI) of the expected product (unpurified sample)**

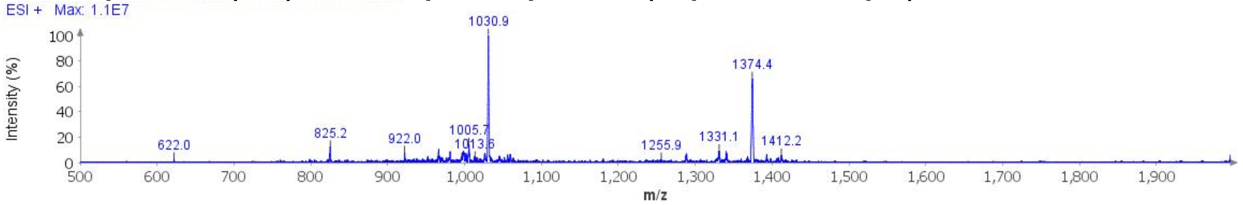

**Deconvoluted Mass Spectrum (ESI) of the expected product (unpurified sample)**

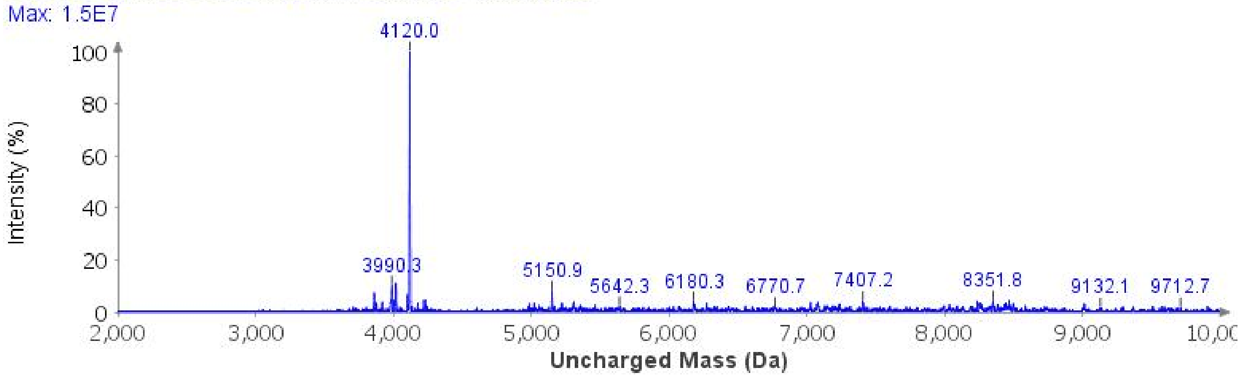

**Analytical LCMS**: Measured on an Agilent EC-C18 Poroshell column (4.6 x 100 mm, 4 μm, 100 Å, 40 °C, 1 mL/min) 5%-65% MeCN/$H_2O$/0.05% TFA over 60 min.

**Chemical Formula**: $C_{175}H_{278}N_{52}O_{58}Se$
**Molecular Weight**: 4117.4260 g/mol

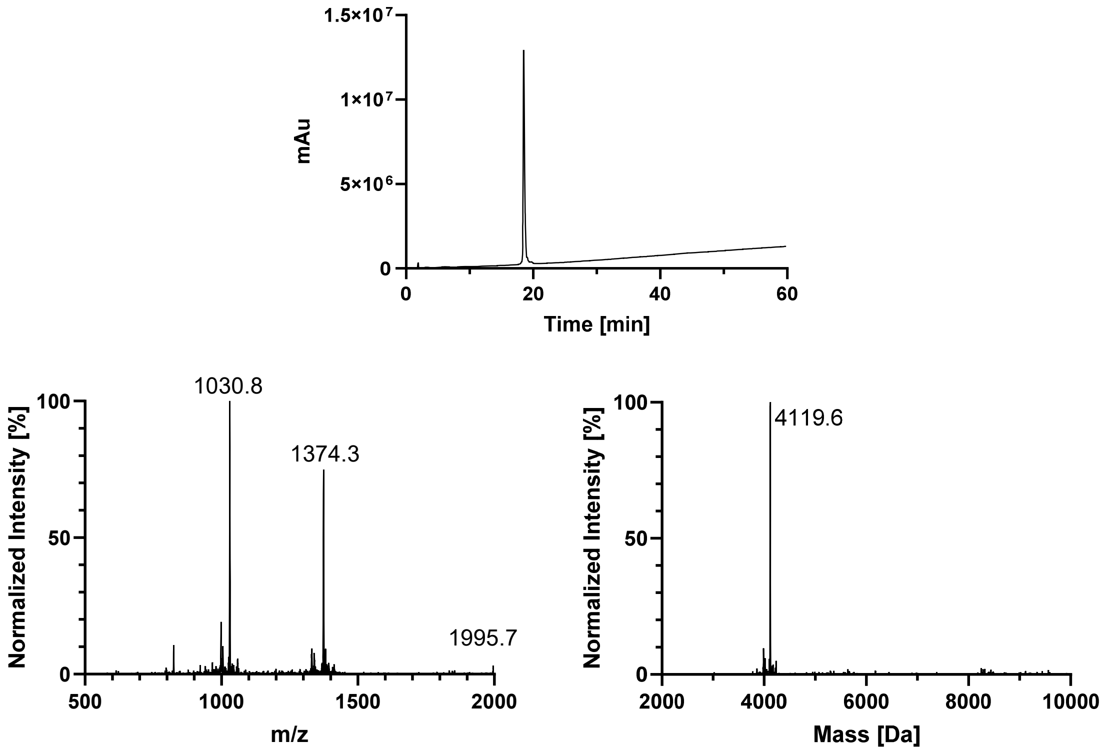

1.5×10⁷
1×10⁷
5×10⁶
0
mAu
0
20
40
60
Time [min]
1030.8
1374.3
1995.7
100
50
0
Normalized Intensity [%]
500
1000
1500
2000
m/z
4119.6
100
50
0
Normalized Intensity [%]
2000
4000
6000
8000
10000
Mass [Da]

#### 3. C322-L357 (SPPS)

### WT

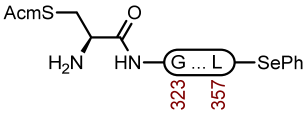

**2a**

**Sequence**: H-C(Acm)GSLGNIHHKPGGGQVEVKSEKLDFKDRVQSKIGSL-SePh

**Resin**: HMPB ChemMatrix resin from Biotage (100-200 mesh, 0.42 mmol/g, 595 mg, 0.25 mmol) was used. The resin was loaded by the symmetrical anhydride with Fmoc-L-Leu-OH, DIC, and 4-DMAP.

**Amino Acids**: Boc-L-Cys(Acm)-OH, Fmoc-Gly-OH, Fmoc-L-Ser($^t$Bu)-OH, Fmoc-L-Leu-OH, Fmoc-L-Asn(Trt)-OH, Fmoc-L-Ile-OH, Fmoc-L-His(Trt)-OH, Fmoc-L-Lys(Boc)-OH, Fmoc-L-Pro-OH, Fmoc-L-Gln(Trt)-OH, Fmoc-L-Val-OH, Fmoc-L-Glu(O $^t$Bu)-OH, Fmoc-L-Asp(O $^t$Bu)-OH, Fmoc-L-Phe-OH, Fmoc-L-Arg(Pbf)-OH,

**Coupling**: Amino acids were double coupled using 0.2 M amino acid in DMF (5 equiv.), 1 M DIC in DMF (10 equiv.), and 1 M oxyma pure in DMF (5 equiv.) at 90°C for 4 min. Histidine was double coupled at 50°C for 10 min.

**Deprotection**: 20% piperidine + 0.1 M oxyma pure in DMF (6 mL) at 23 ºC (1 x 5 min, 1 x 15 min).

**Analytical Cleavage (SPPS)**: A small portion of resin was treated with TFA:$H_2O$:TA:TIS (85:5:5:5) at 23 °C for 3 h, before it was filtered and precipitated with $E_2O$. The precipitate was centrifuged, and the resulting pellet was washed twice more with $Et_2O$, and the pellet was air dried.

**Crude LCMS (SPPS)**: Measured on an Agilent EC-C18 Poroshell column (4.6 x 100 mm, 4 µm, 100 Å, 40 °C, 1 mL/min) 5%-65% MeCN/$H_2O$/0.05% TFA over 60 min.

**LC trace (214 nm) of unpurified peptide**

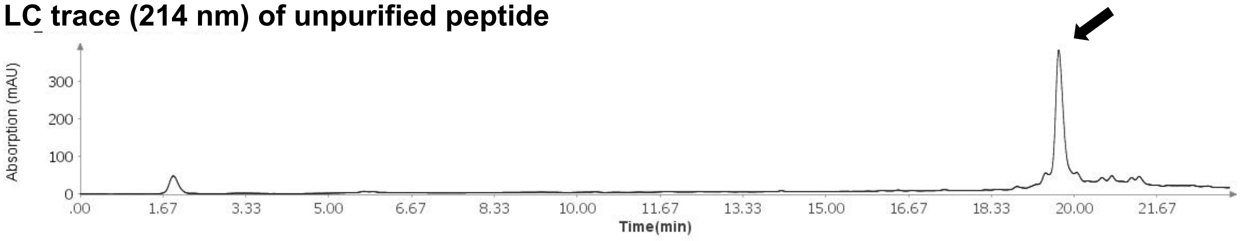

**Total Ion Current (TIC) of unpurified peptide**

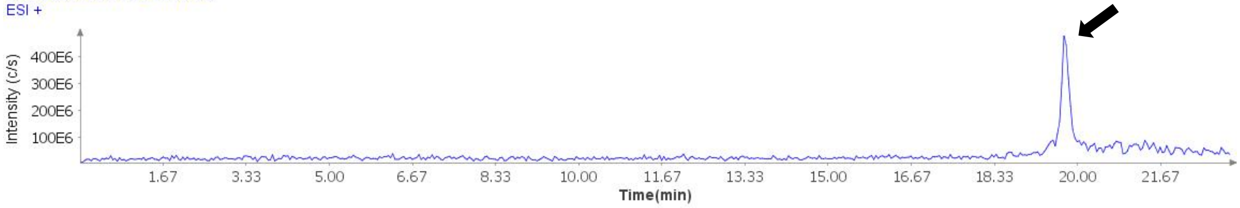

**Extracted Ion Current (XIC) of unpurified peptide**

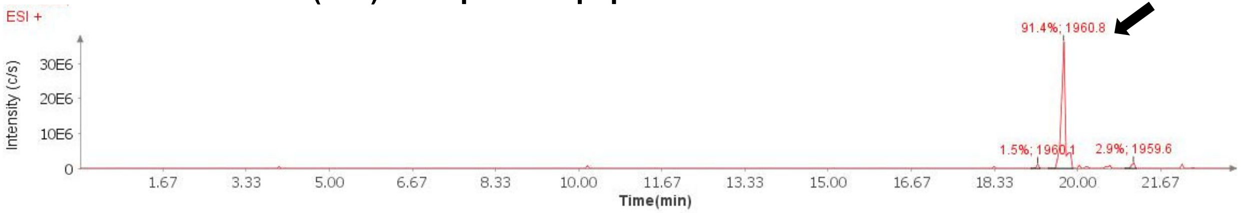

**Mass Spectrum (ESI) of the expected product (unpurified sample)**

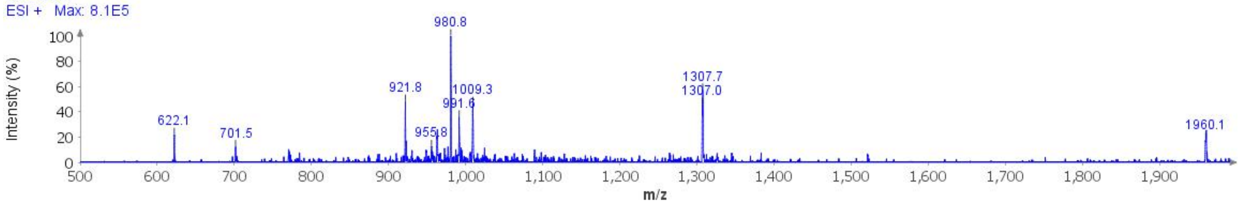

**Deconvoluted Mass Spectrum (ESI) of the expected product (unpurified sample)**

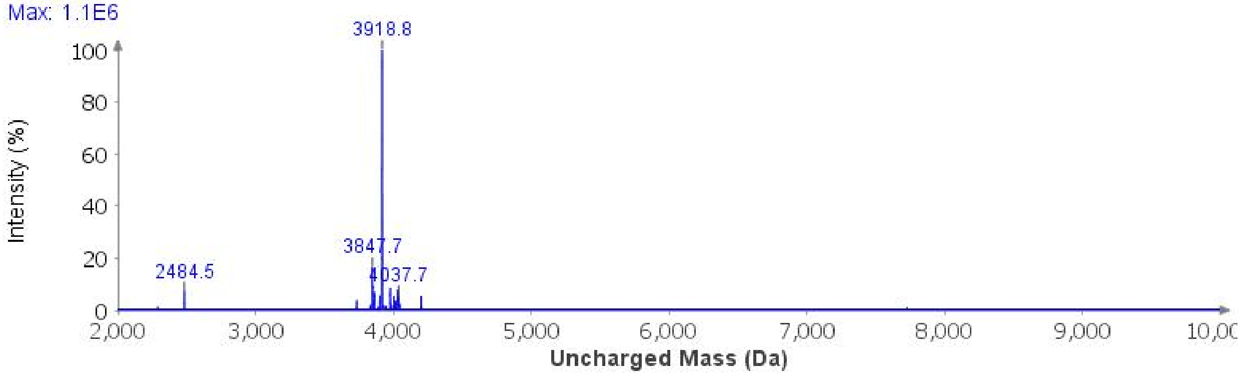

**Protected cleavage**: The resin was treated with 1% TFA in DCM (4 mL, 4 min x 10), and filtered into 10% pyridine in MeOH (60 mL). The pooled organic layers were concentrated, and protected peptide was precipitated with water, collected by centrifugation, and lyophilized to afford crude protected peptide acid (570 mg, 86.7 μmol).

**Selenoesterification**: A 100 mL RBF with the crude protected acid (570 mg, 86.7 μmol), DPDS (822 mg, 260 mmol), and dry DMF (15 mL) was cooled to 0°C in an ice bath and treated with P(*n*-Bu)$_3$ (642 μL, 2.60 mmol). The reaction was stirred at 0°C for 3 h before it was concentrated under a stream of compressed air. The residue was triturated with hexanes and taken forward to the next step. [1]

**Global Deprotection**: The crude selenoester was treated with TFA:$H_2O$:thioanisole:TIPSH (85:5:5:5, 20 mL) and it was stirred at 23 ºC for 3 h before it was cooled to 0°C and treated with $Et_2O$ to precipitate the peptide. The precipitate was centrifuged, and the resulting pellet was washed twice more with $Et_2O$. It was dissolved in 40% MeCN and it was lyophilized.

**Crude LCMS (Selenoesterification)**: Measured on an Agilent EC-C18 Poroshell column (4.6 x 100 mm, 4 μm, 100 Å, 40 °C, 1 mL/min) 5%-65% MeCN/$H_2O$/0.05% TFA over 60 min.

**LC trace (214 nm) of unpurified peptide**

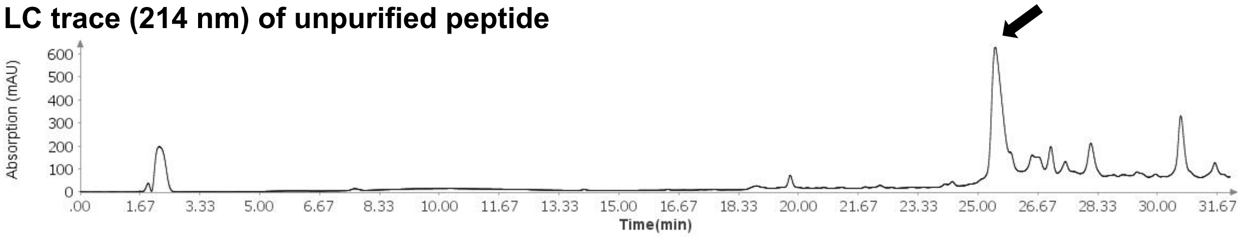

**Total Ion Current (TIC) of unpurified peptide**

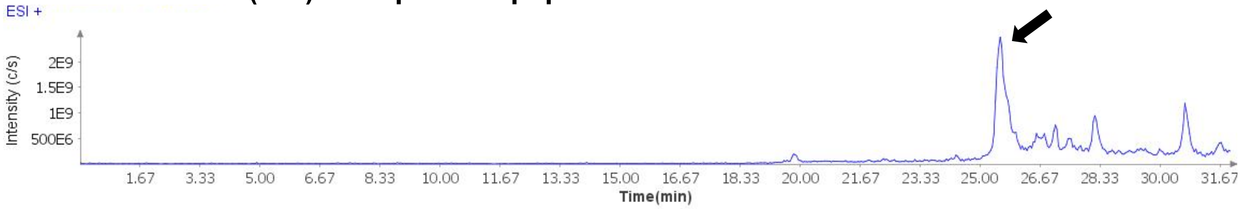

**Extracted Ion Current (XIC) of unpurified peptide**

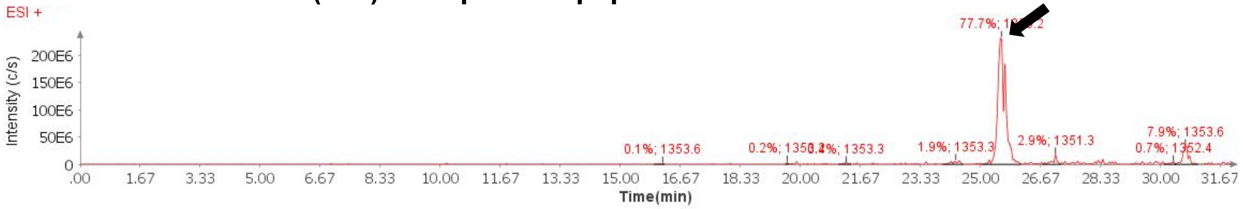

**Mass Spectrum (ESI) of the expected product (unpurified sample)**

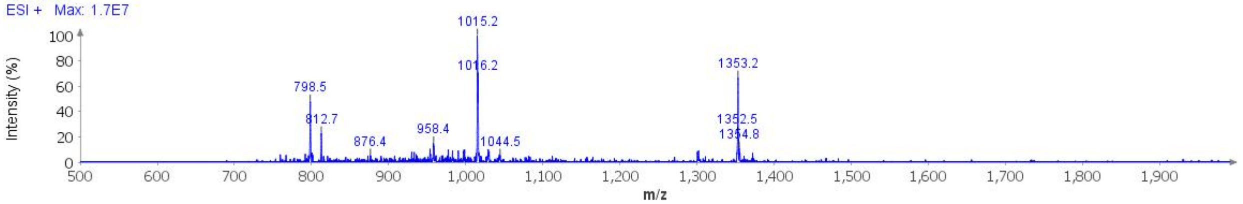

**Deconvoluted Mass Spectrum (ESI) of the expected product (unpurified sample)**

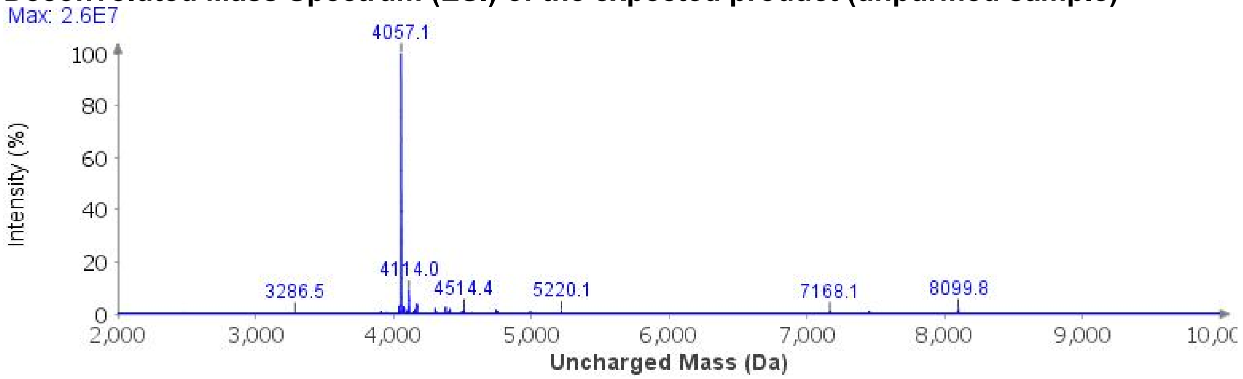

**Purification**: The material was purified over an ACQUITY Protein BEH C4 column (19 x 250 mm, 5 µm, 300 Å, 17.1 mL/min) 20%-45% MeCN/$H_2O$/0.1% TFA over 30 min. The fractions were lyophilized to afford **2a** (43.8 mg, 4% yield).[3]

**Analytical LCMS**: Measured on an Agilent EC-C18 Poroshell column (4.6 x 100 mm, 4 µm, 100 Å, 40 °C, 1 mL/min) 5%-40% MeCN/$H_2O$/0.1% FA over 10 min.

**Chemical Formula**: $C_{175}H_{284}N_{52}O_{52}SSe$
**Molecular Weight**: 4059.5400 g/mol

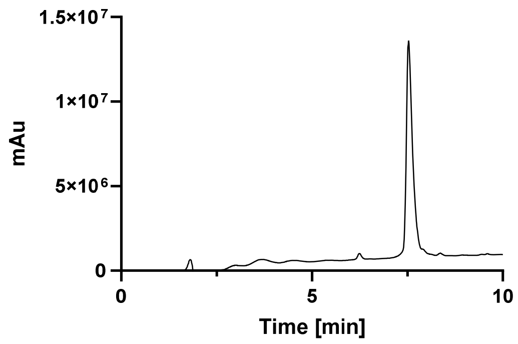

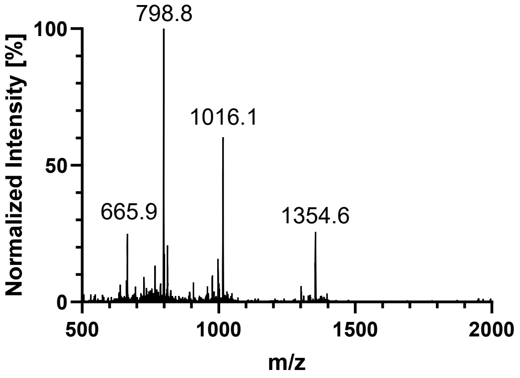

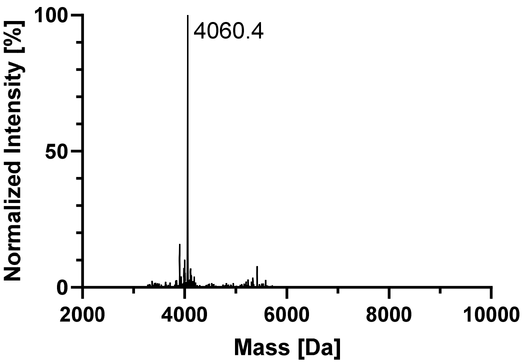

#### AcK353

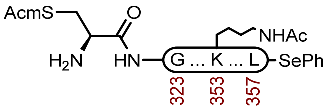

**2d**

**Sequence**: H-C(Acm)GSLGNIHHKPGGGQVEVKSEKLDFKDRVQSKIGSL-SePh

**Resin**: HMPB ChemMatrix resin from Biotage (100-200 mesh, 0.42 mmol/g, 238 mg, 0.1 mmol) was used. The resin was loaded by the symmetrical anhydride with Fmoc-L-Leu-OH, DIC, and 4-DMAP.

**Amino Acids**: Boc-L-Cys(Acm)-OH, Fmoc-Gly-OH, Fmoc-L-Ser($^t$Bu)-OH, Fmoc-L-Leu-OH, Fmoc-L-Asn(Trt)-OH, Fmoc-L-Ile-OH, Fmoc-L-His(Trt)-OH, Fmoc-L-Lys(Boc)-OH, Fmoc-L-Pro-OH, Fmoc-L-Gln(Trt)-OH, Fmoc-L-Val-OH, Fmoc-L-Glu(O $^t$Bu)-OH, Fmoc-L-Asp(O $^t$Bu)-OH, Fmoc-L-Phe-OH, Fmoc-L-Arg(Pbf)-OH, Fmoc-L-Lys(Ac)-OH

**Coupling**: Amino acids were double coupled using 0.2 M amino acid in DMF (5 equiv.), 1 M DIC in DMF (10 equiv.), and 1 M oxyma pure in DMF (5 equiv.) at 90°C for 4 min. Histidine was double coupled at 50°C for 10 min.

**Deprotection**: 20% piperidine + 0.1 M oxyma pure in DMF (6 mL) at 23 ºC (1 x 5 min, 1 x 15 min).

**Crude LCMS (SPPS)**: Measured on an Agilent EC-C18 Poroshell column (4.6 x 100 mm, 4 µm, 100 Å, 40 °C, 1 mL/min) 5%-100% MeCN/$H_2O$/0.05% TFA over 30 min.

**LC trace (214 nm) of unpurified peptide**

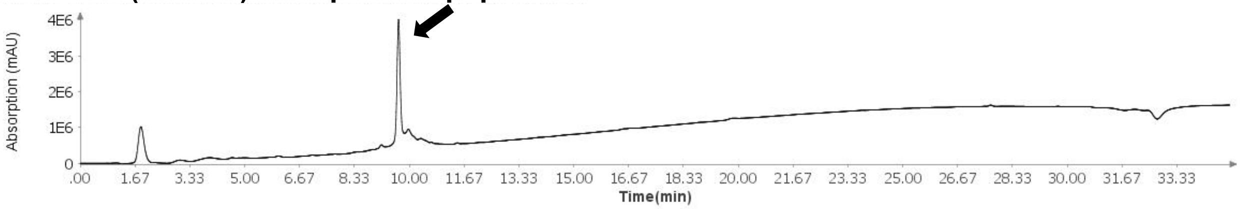

**Total Ion Current (TIC) of unpurified peptide**

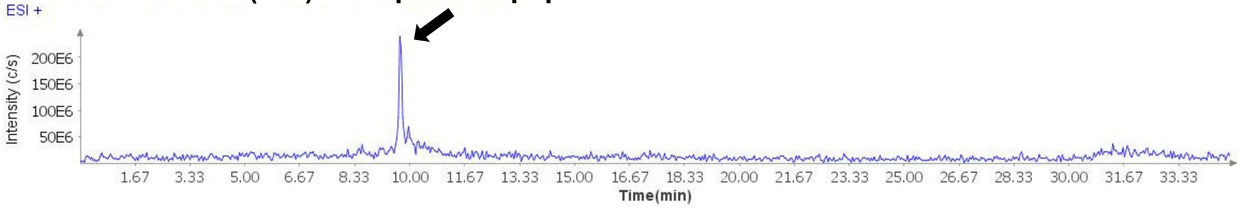

**Extracted Ion Current (XIC) of unpurified peptide**

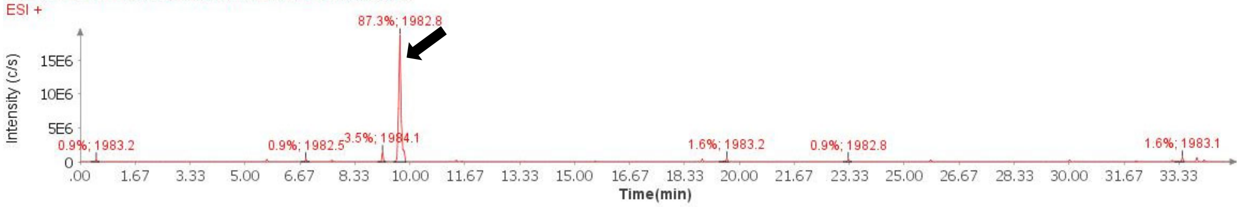

**Mass Spectrum (ESI) of the expected product (unpurified sample)**

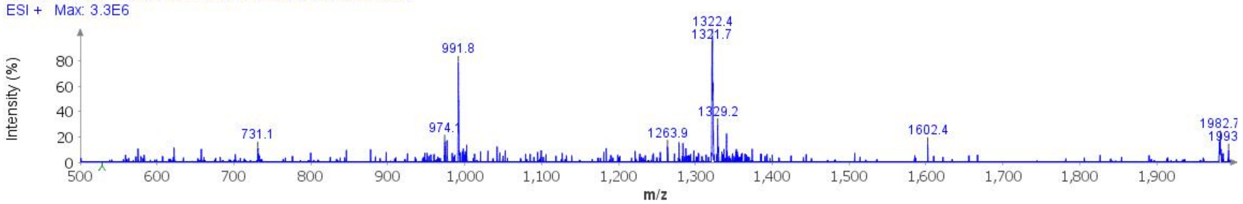

**Deconvoluted Mass Spectrum (ESI) of the expected product (unpurified sample)**

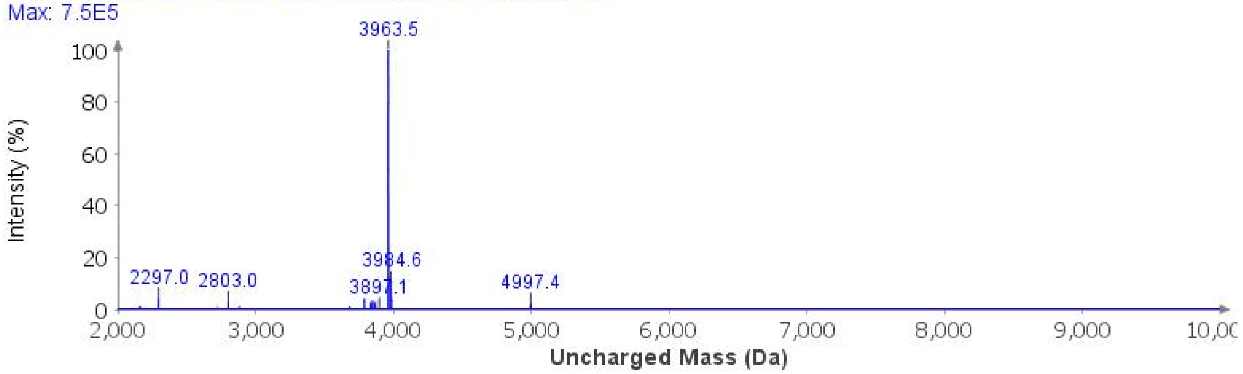

**Protected cleavage**: The resin was treated with 1% TFA in DCM (4 mL, 4 min x 10), and filtered into 10% pyridine in MeOH (60 mL). The pooled organic layers were concentrated, and protected peptide was precipitated with water, collected by centrifugation, and lyophilized to afford crude protected peptide acid (166.1 mg, 25.5 μmol).

**Selenoesterification**: A 100 mL RBF with the crude protected acid (166.1 mg, 25.5 μmol), DPDS (238.6 mg, 0.764 mmol), and dry DMF (3 mL) was cooled to 0°C in an ice bath and treated with P(*n*-Bu)$_3$ (188 μL, 0.764 mmol). The reaction was stirred at 0°C for 3 h before it was concentrated under a stream of compressed air. The residue was triturated with hexanes and taken forward to the next step.

**Global Deprotection**: The crude selenoester was treated with TFA:$H_2O$:thioanisole:TIPSH (85:5:5:5, 20 mL) and it was stirred at 23 ºC for 3 h before it was cooled to 0°C and treated with $Et_2O$ to precipitate the peptide. The precipitate was centrifuged, and the resulting pellet was washed twice more with $Et_2O$. It was dissolved in 40% MeCN and it was lyophilized.

**Crude LCMS (Selenoesterification)**: Measured on an Agilent EC-C18 Poroshell column (4.6 x 100 mm, 4 μm, 100 Å, 40 °C, 1 mL/min) 5%-65% MeCN/$H_2O$/0.05% TFA over 60 min.

**LC trace (214 nm) of unpurified peptide**

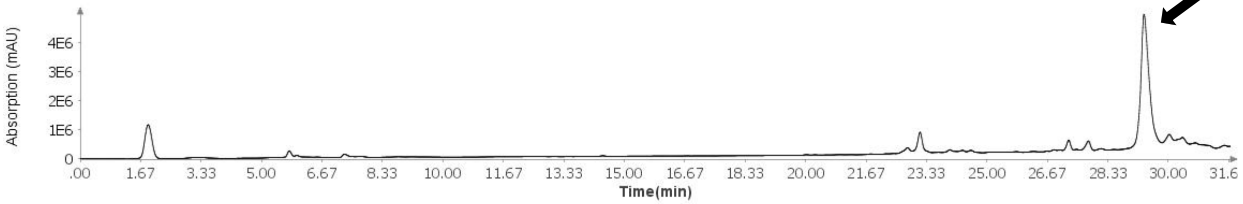

**Total Ion Current (TIC) of unpurified peptide**

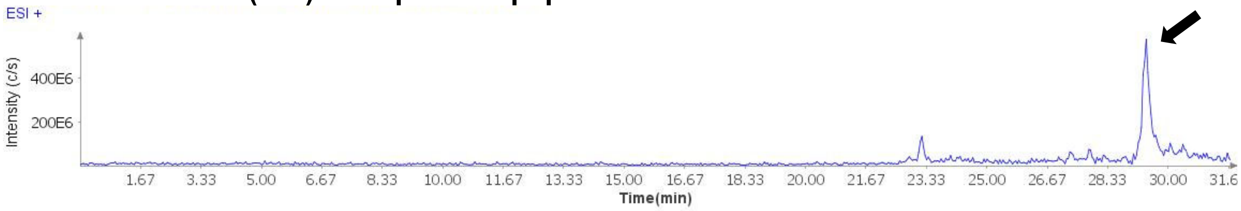

**Extracted Ion Current (XIC) of unpurified peptide**

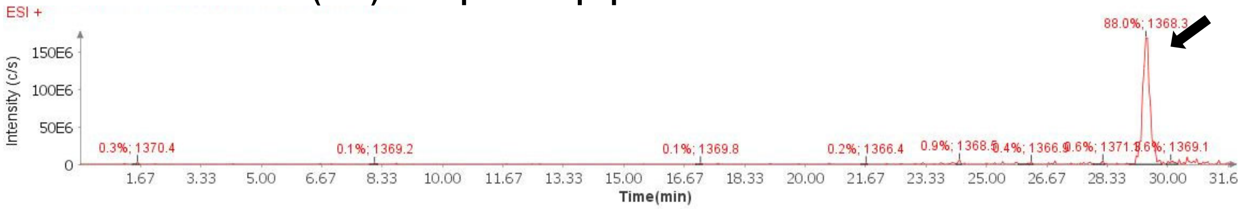

**Mass Spectrum (ESI) of the expected product (unpurified sample)**

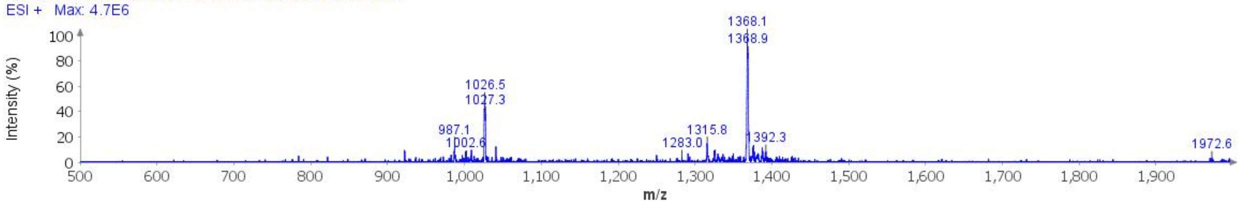

**Deconvoluted Mass Spectrum (ESI) of the expected product (unpurified sample)**

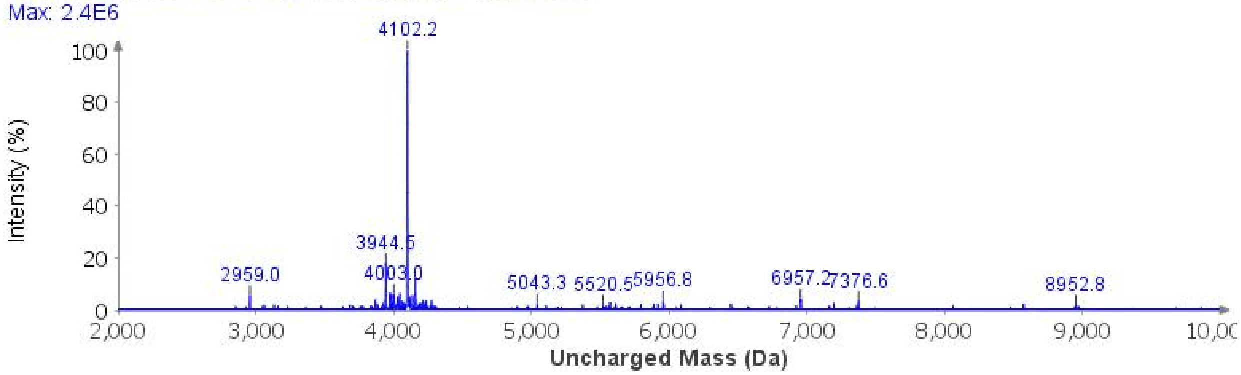

**Purification**: The material was purified over an ACQUITY Protein BEH C4 column (19 x 250 mm, 5 µm, 300 Å, 17.1 mL/min) 20%-45% MeCN/$H_2O$/0.1% TFA over 30 min. The fractions were lyophilized to afford **2d** (41.5 mg, 10% yield).

**Analytical LCMS**: Measured on an Agilent EC-C18 Poroshell column (4.6 x 100 mm, 4 µm, 100 Å, 40 °C, 1 mL/min) 5%-65% MeCN/$H_2O$/0.05% TFA over 60 min.

**Chemical Formula**: $C_{177}H_{286}N_{52}O_{53}SSe$
**Molecular Weight**: 4101.5770 g/mol

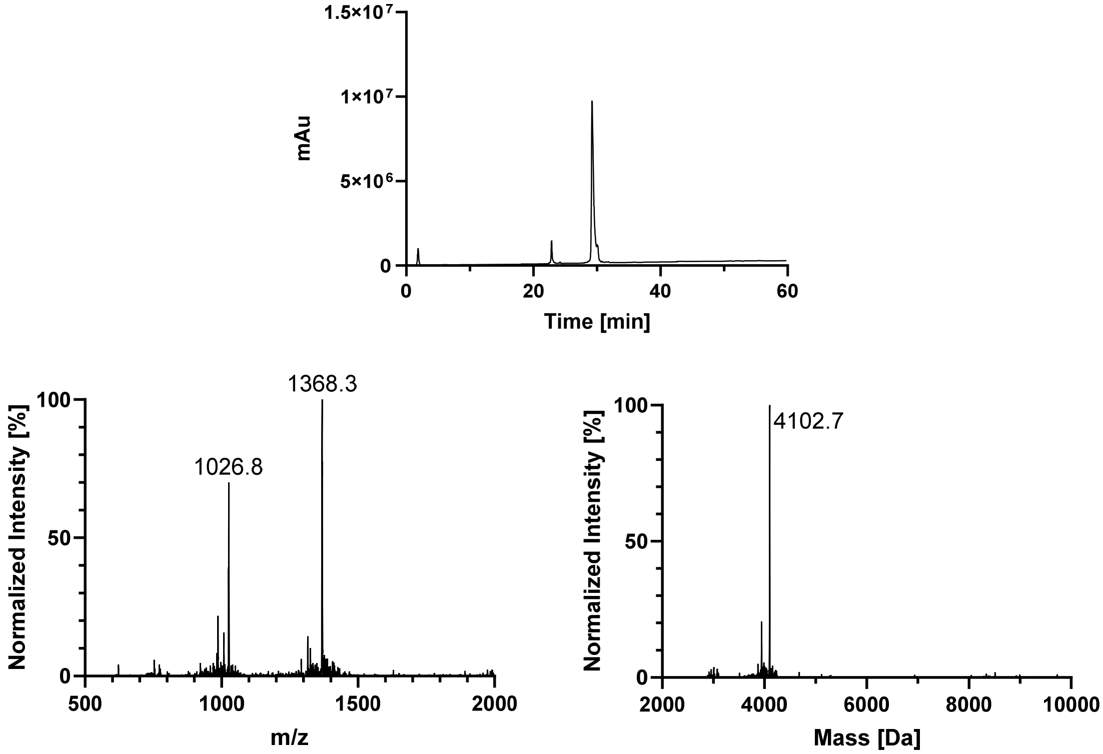
1.5×10⁷
1×10⁷
5×10⁶
0
mAu
0
20
40
60
Time [min]
1368.3
1026.8
Normalized Intensity [%]
100
50
0
500
1000
1500
2000
m/z
4102.7
Normalized Intensity [%]
100
50
0
2000
4000
6000
8000
10000
Mass [Da]

## pS356

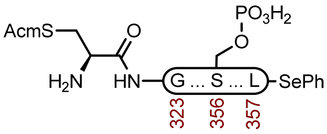

**2g**

**Sequence**: H-C(Acm)GSLGNIHHKPGGGQVEVKSEKLDFKDRVQSKIGS($OPO_3H_2$)L-SePh

**Resin**: HMPB ChemMatrix resin from Biotage (100-200 mesh, 0.42 mmol/g, 476 mg, 0.2 mmol) was used. The resin was loaded by the symmetrical anhydride with Fmoc-L-Leu-OH, DIC, and 4-DMAP.

**Amino Acids**: Boc-L-Cys(Acm)-OH, Fmoc-Gly-OH, Fmoc-L-Ser($^t$Bu)-OH, Fmoc-L-Leu-OH, Fmoc-L-Asn(Trt)-OH, Fmoc-L-Ile-OH, Fmoc-L-His(Trt)-OH, Fmoc-L-Lys(Boc)-OH, Fmoc-L-Pro-OH, Fmoc-L-Gln(Trt)-OH, Fmoc-L-Val-OH, Fmoc-L-Glu(O $^t$Bu)-OH, Fmoc-L-Asp(O $^t$Bu)-OH, Fmoc-L-Phe-OH, Fmoc-L-Arg(Pbf)-OH, Fmoc-L-Ser($HPO_3Bn$)-OH

**Coupling**: Amino acids were double coupled using 0.2 M amino acid in DMF (5 equiv.), 0.4 M HATU in DMF (5 equiv.), and 1 M DIPEA in NMP (5 equiv.) at 23 °C for 45 min.

**Coupling Fmoc-L-Ser($HPO_3Bn$)-OH**: Fmoc-L-Ser($HPO_3Bn$)-OH (2.5 equiv.), HATU (2.5 equiv.), Oxyma (2.5 equiv.), DIPEA (7 equiv.) was preactivated in DMF and added to the resin. Mixed at 25 °C for 16 h.

**Deprotection**: 20% piperidine + 0.1 M oxyma pure in DMF (6 mL) at 23 ºC (1 x 5 min, 1 x 15 min).

**Crude LCMS (SPPS)**: Measured on an Agilent EC-C18 Poroshell column (4.6 x 100 mm, 4 µm, 100 Å, 40 °C, 1 mL/min) 5%-65% MeCN/$H_2O$/0.05% TFA over 60 min.

##### LC trace (214 nm) of unpurified peptide

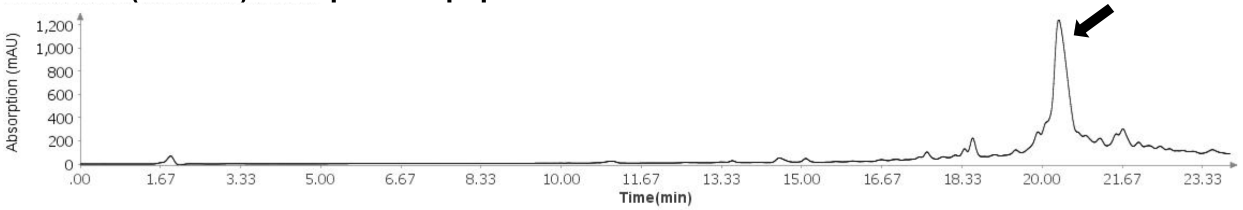

##### Total Ion Current (TIC) of unpurified peptide

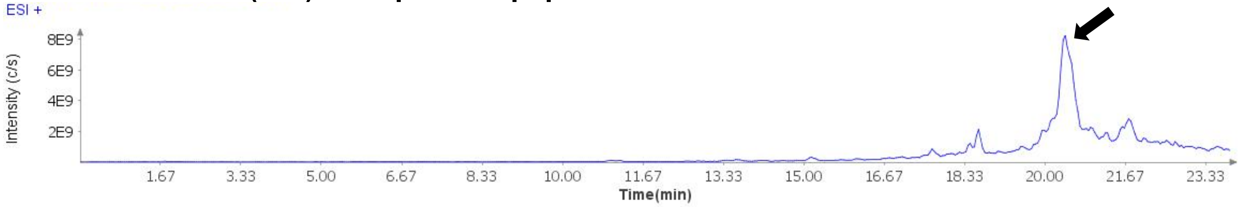

**Extracted Ion Current (XIC) of unpurified peptide**

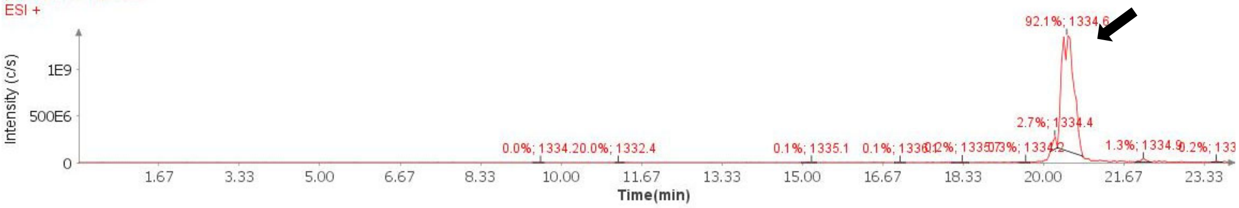

**Mass Spectrum (ESI) of the expected product (unpurified sample)**

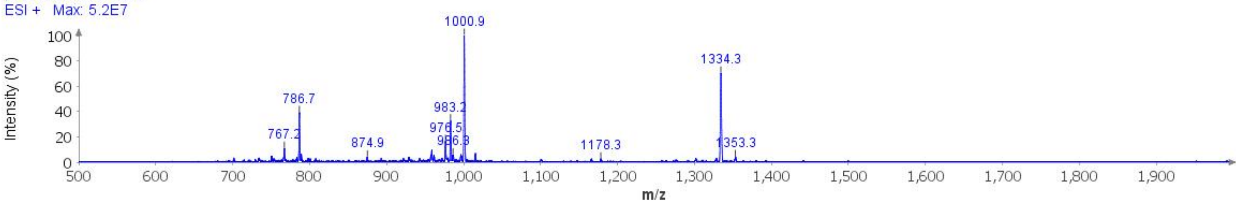

**Deconvoluted Mass Spectrum (ESI) of the expected product (unpurified sample)**

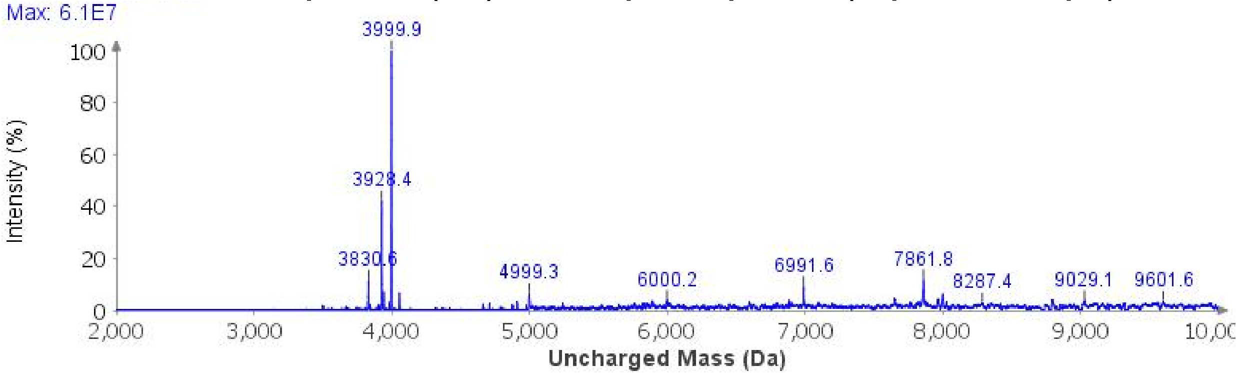

**Protected cleavage**: The resin was treated with 1% TFA in DCM (8 mL, 4 min x 10), and filtered into 10% pyridine in MeOH (120 mL). The pooled organic layers were concentrated, and protected peptide was precipitated with water, collected by centrifugation, and lyophilized to afford crude protected peptide acid (677.1 mg, 103.4 µmol).

**Selenoesterification**: A 100 mL RBF with the crud e protected acid (620.6 mg, 94.7 µmol), DPDS (887 mg, 2.84 mmol), and dry DMF (18.9 mL) was cooled to 0°C in an ice bath and treated with $P(n\text{-}Bu)_3$ (710 µL, 2.84 mmol). The reaction was stirred at 0°C for 3 h before it was concentrated under a stream of compressed air. The residue was triturated with hexanes and taken forward to the next step.

**Global Deprotection**: The crude selenoester was treated with TFA:$H_2O$:thioanisole:TIPSH (85:5:5:5, 20 mL) and it was stirred at 23 ºC for 3 h before it was cooled to 0°C and treated with $Et_2O$ to precipitate the peptide. The precipitate was centrifuged, and the resulting pellet was washed twice more with $Et_2O$. It was dissolved in 40% MeCN and it was lyophilized.

**Crude LCMS (Selenoesterification)**: Measured on an Agilent EC-C18 Poroshell column (4.6 x 100 mm, 4 µm, 100 Å, 40 °C, 1 mL/min) 5%-65% MeCN/$H_2O$/0.05% TFA over 60 min.

**LC trace (214 nm) of unpurified peptide**

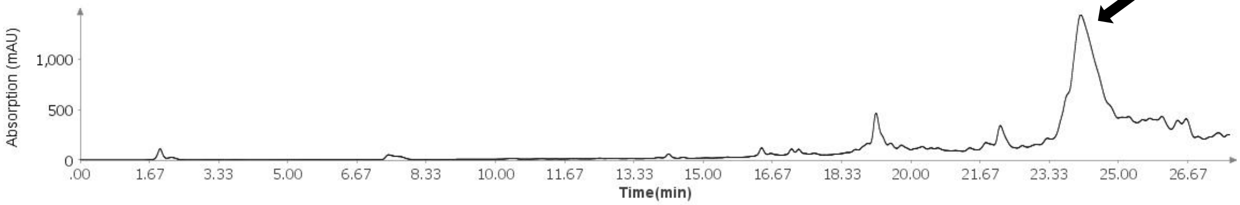

**Total Ion Current (TIC) of unpurified peptide**

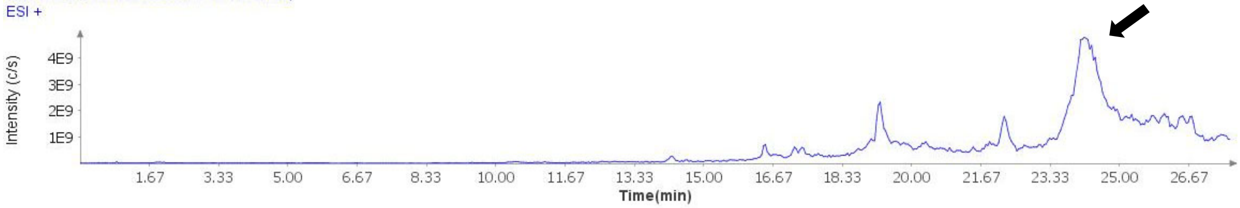

**Extracted Ion Current (XIC) of unpurified peptide**

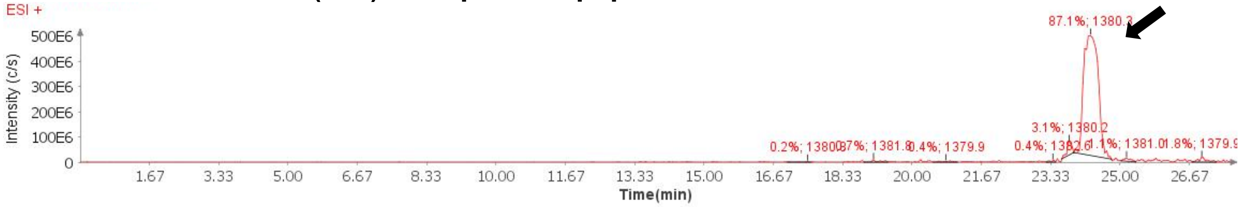

**Mass Spectrum (ESI) of the expected product (unpurified sample)**

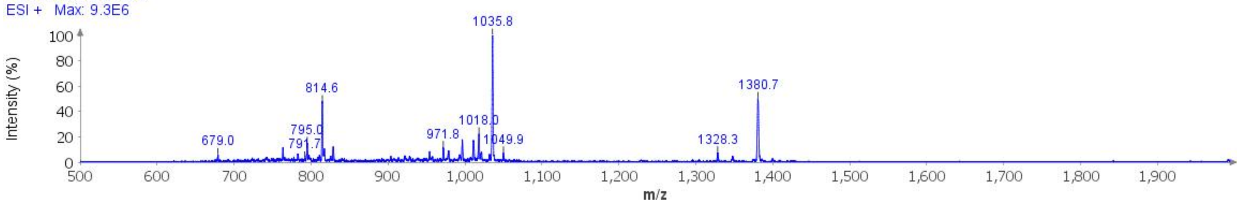

**Deconvoluted Mass Spectrum (ESI) of the expected product (unpurified sample)**

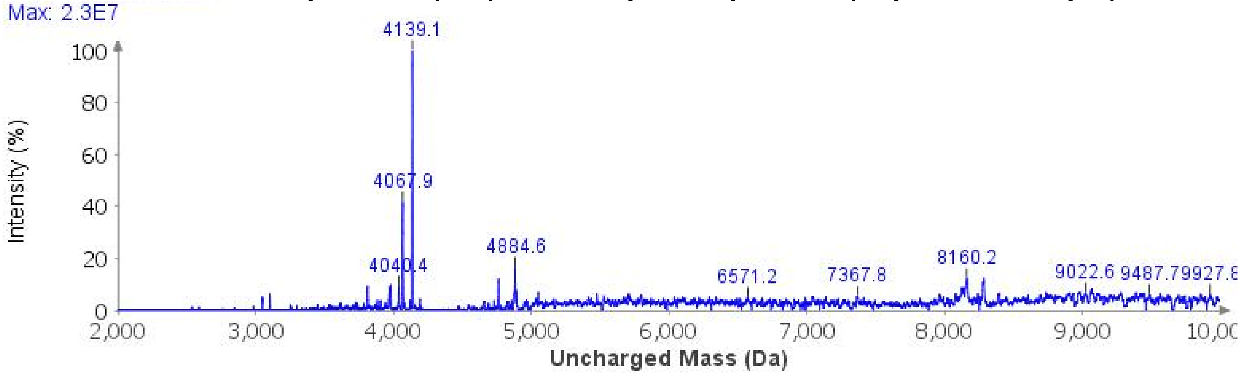

**Purification**: The material was purified over a Luna Prep C4 column (10 x 250 mm, 10 µm, 100 Å, 4.7 mL/min) 20%-35% MeCN/$H_2O$/0.1% TFA over 30 min. The fractions were lyophilized to afford **2g** (41.7 mg, 5% yield).

**Analytical LCMS**: Measured on an Agilent EC-C18 Poroshell column (4.6 x 100 mm, 4 µm, 100 Å, 40 °C, 1 mL/min) 5%-65% MeCN/$H_2O$/0.05% TFA over 60 min.

**Chemical Formula**: $C_{175}H_{285}N_{52}O_{55}PSSe$
**Molecular Weight**: 4139.5188 g/mol

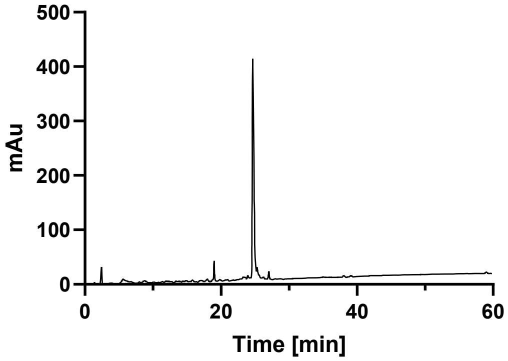

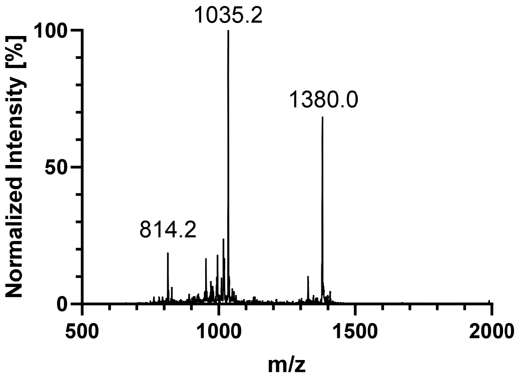

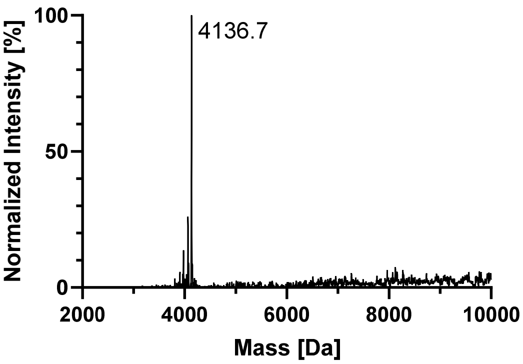

## pS352

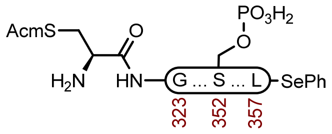

**2h**

**Sequence**: H-C(Acm)GSLGNIHHKPGGGQVEVKSEKLDFKDRVQS($OPO_3H_2$)KIGSL-SePh

**Resin**: HMPB ChemMatrix resin from Biotage (100-200 mesh, 0.42 mmol/g, 476 mg, 0.2 mmol) was used. The resin was loaded by the symmetrical anhydride with Fmoc-L-Leu-OH, DIC, and 4-DMAP.

**Amino Acids**: Boc-L-Cys(Acm)-OH, Fmoc-Gly-OH, Fmoc-L-Ser($^t$Bu)-OH, Fmoc-L-Leu-OH, Fmoc-L-Asn(Trt)-OH, Fmoc-L-Ile-OH, Fmoc-L-His(Trt)-OH, Fmoc-L-Lys(Boc)-OH, Fmoc-L-Pro-OH, Fmoc-L-Gln(Trt)-OH, Fmoc-L-Val-OH, Fmoc-L-Glu(O $^t$Bu)-OH, Fmoc-L-Asp(O $^t$Bu)-OH, Fmoc-L-Phe-OH, Fmoc-L-Arg(Pbf)-OH, Fmoc-L-Ser($HPO_3$Bn)-OH

**Coupling**: Amino acids were double coupled using 0.2 M amino acid in DMF (5 equiv.), 0.4 M HATU in DMF (5 equiv.), and 1 M DIPEA in NMP (5 equiv.) at 23 °C for 45 min.

**Coupling Fmoc-L-Ser($HPO_3$Bn)-OH**: Fmoc-L-Ser($HPO_3$Bn)-OH (2.5 equiv.), HATU (2.5 equiv.), Oxyma (2.5 equiv.), DIPEA (7 equiv.) was preactivated in DMF and added to the resin. Mixed at 25 °C for 16 h.

**Deprotection**: 20% piperidine + 0.1 M oxyma pure in DMF (6 mL) at 23 ºC (1 x 5 min, 1 x 15 min).

**Crude LCMS (SPPS)**: Measured on an Agilent EC-C18 Poroshell column (4.6 x 100 mm, 4 μm, 100 Å, 40 °C, 1 mL/min) 5%-65% MeCN/$H_2O$/0.05% TFA over 60 min.

**LC trace (214 nm) of unpurified peptide**

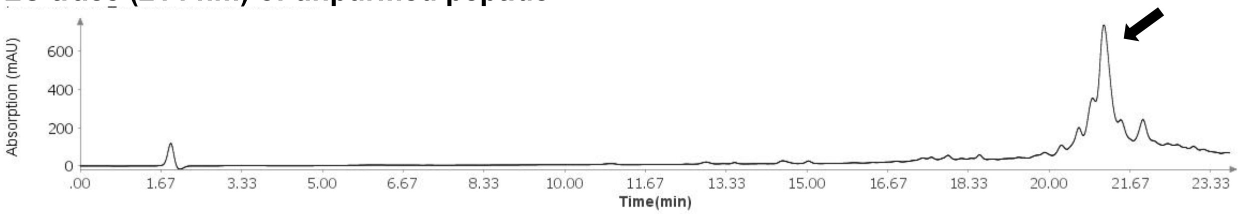

**Total Ion Current (TIC) of unpurified peptide**

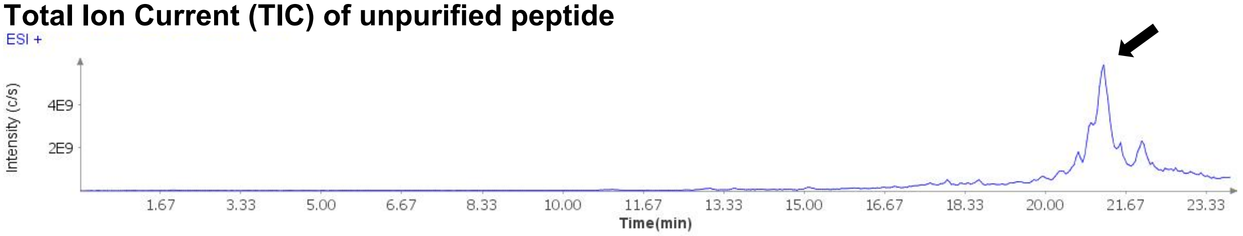

**Extracted Ion Current (XIC) of unpurified peptide**

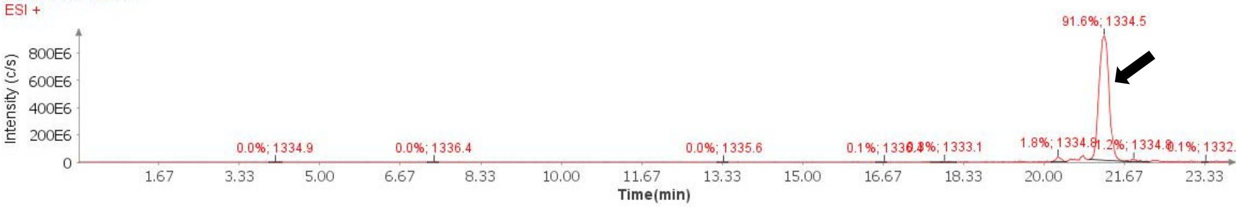

**Mass Spectrum (ESI) of the expected product (unpurified sample)**

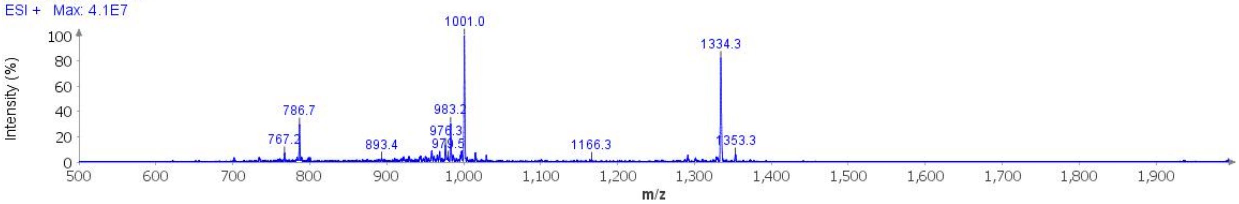

**Deconvoluted Mass Spectrum (ESI) of the expected product (unpurified sample)**

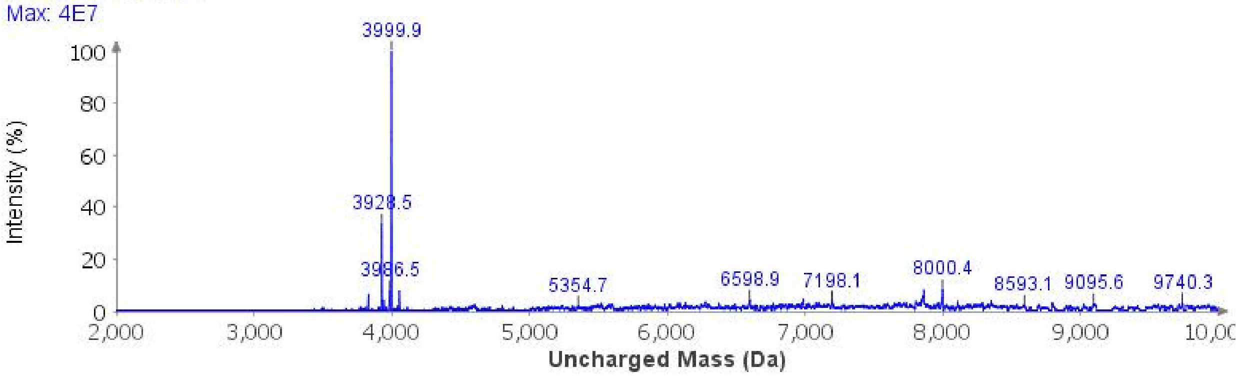

**Protected cleavage**: The resin was treated with 1% TFA in DCM (8 mL, 4 min x 10), and filtered into 10% pyridine in MeOH (120 mL). The pooled organic layers were concentrated, and protected peptide was precipitated with water, collected by centrifugation, and lyophilized to afford crude protected peptide acid (548.1 mg, 83.7 μmol).

**Selenoesterification**: A 100 mL RBF with the crude protected acid (508.2 mg, 77.6 μmol), DPDS (726.6 mg, 2.33 mmol), and dry DMF (18.9 mL) was cooled to 0°C in an ice bath and treated with P(*n*-Bu)$_3$ (471 μL, 2.33 mmol). The reaction was stirred at 0°C for 3 h before it was concentrated under a stream of compressed air. The residue was triturated with hexanes and taken forward to the next step.

**Global Deprotection**: The crude selenoester was treated with TFA:$H_2O$:thioanisole:TIPSH (85:5:5:5, 20 mL) and it was stirred at 23 ºC for 3 h before it was cooled to 0°C and treated with $Et_2O$ to precipitate the peptide. The precipitate was centrifuged, and the resulting pellet was washed twice more with $Et_2O$. It was dissolved in 40% MeCN and it was lyophilized.

**Crude LCMS (Selenoesterification)**: Measured on an Agilent EC-C18 Poroshell column (4.6 x 100 mm, 4 μm, 100 Å, 40 °C, 1 mL/min) 5%-65% MeCN/$H_2O$/0.05% TFA over 60 min.

**LC trace (214 nm) of unpurified peptide**

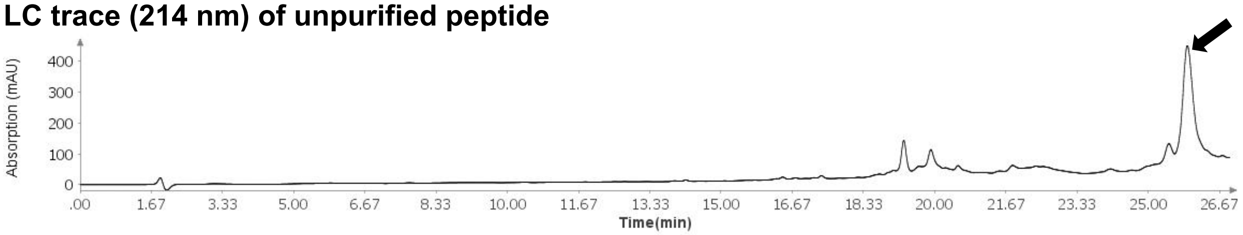

**Total Ion Current (TIC) of unpurified peptide**

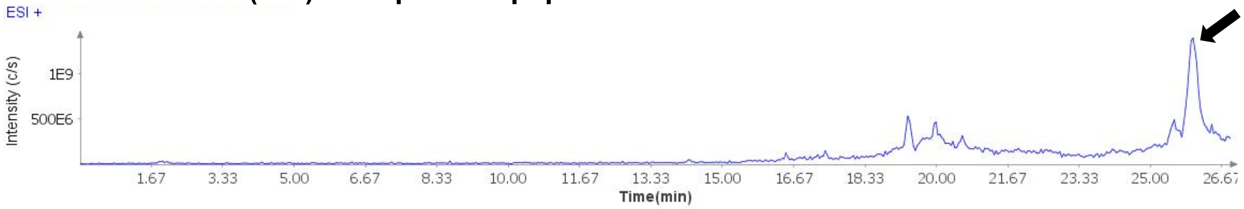

**Extracted Ion Current (XIC) of unpurified peptide**

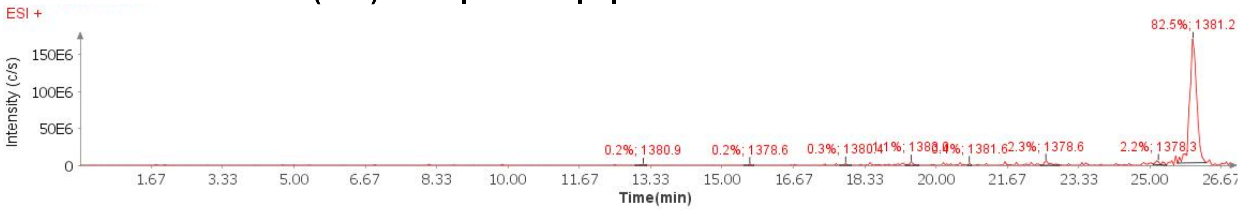

**Mass Spectrum (ESI) of the expected product (unpurified sample)**

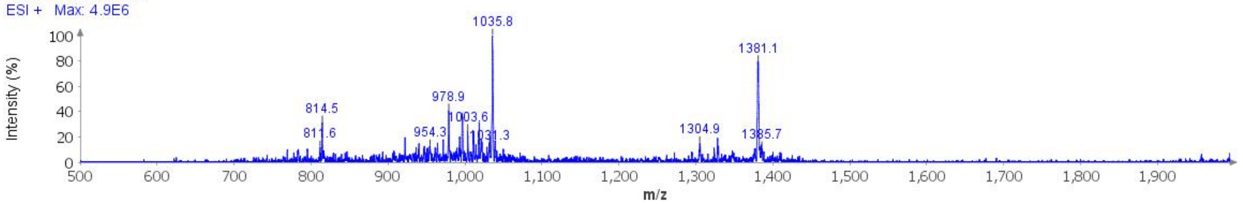

**Deconvoluted Mass Spectrum (ESI) of the expected product (unpurified sample)**

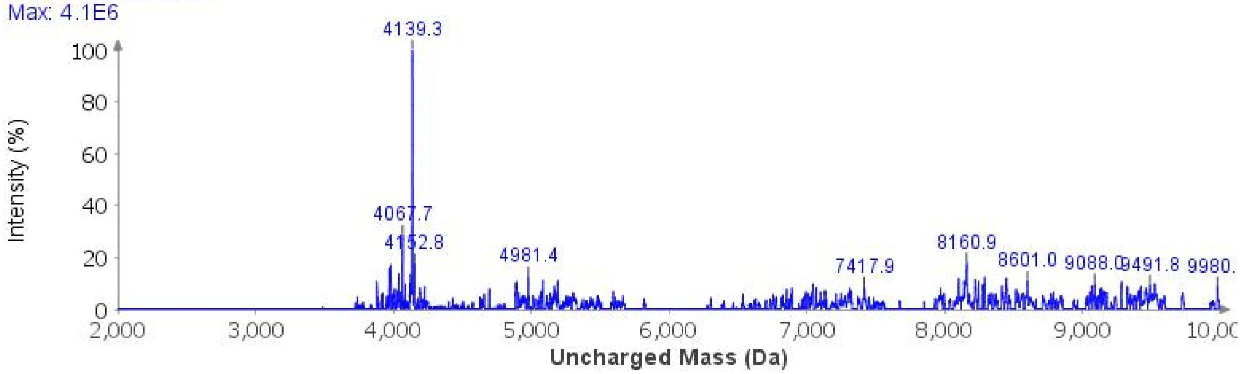

**Purification**: The material was purified over an ACQUITY Protein BEH C4 column (19 x 250 mm, 5 µm, 300 Å, 17.1 mL/min) 25%-31% MeCN/$H_2O$/0.1% TFA over 30 min. The fractions were lyophilized to afford **2h** (38 mg, 5% yield).

**Analytical LCMS**: Measured on an Agilent EC-C18 Poroshell column (4.6 x 100 mm, 4 µm, 100 Å, 40 °C, 1 mL/min) 5%-65% MeCN/$H_2O$/0.05% TFA over 60 min.

**Chemical Formula**: $C_{175}H_{285}N_{52}O_{55}PSSe$
**Molecular Weight**: 4139.5188 g/mol

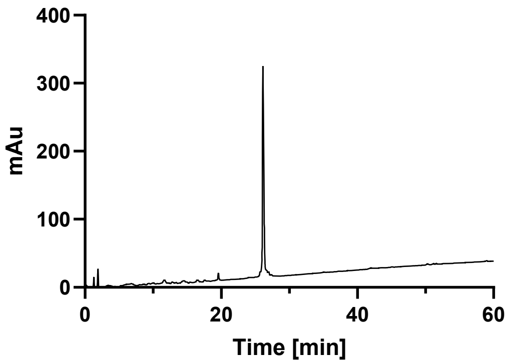

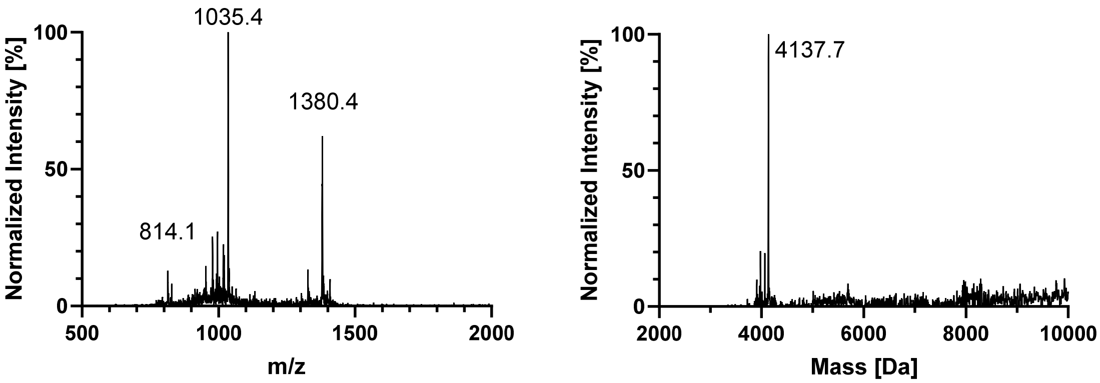

#### 4. C291-K321 (SPPS)

### WT

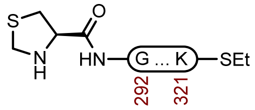

**3a**

**Sequence**: H-ThzGSKDNIKHVPGGGSVQIVYKPVDLSKVTSK-SEt

**Amino Acids**: Boc-L-Thz-OH, Fmoc-Gly-OH, Fmoc-L-Ser($^t$Bu)-OH, Fmoc-L-Lys(Boc)-OH, Fmoc-L-Asp(O$^t$Bu)-OH, Fmoc-L-Asn(Trt)-OH, Fmoc-L-Ile-OH, Fmoc-L-His(Trt)-OH, Fmoc-L-Val-OH, Fmoc-L-Pro-OH, Fmoc-L-Gln(Trt)-OH, Fmoc-L-Tyr(O$^t$Bu)-OH, Fmoc-L-Leu-OH, Fmoc-L-Thr($^t$Bu)-OH.

**Resin**: HMPB ChemMatrix resin from Biotage (100-200 mesh, 0.42 mmol/g, 357 mg, 0.15 mmol) was used on CEM. The resin was loaded by the symmetrical anhydride with Fmoc-L-Lys(Boc)-OH, DIC, and 4-DMAP.

**Coupling**: Amino acids were double coupled using 0.25 M amino acid in DMF (5 equiv.), 1 M DIC in DMF (10 equiv.), and 2 M oxyma pure in DMF (5 equiv.) at 90 °C for 4 min. Histidine was double coupled at 50 °C for 10 min.

**Capping:** The resin was capped after each coupling by treatment with a solution of 5 M $Ac_2O$ (30 equiv.) in DMF, and 5M DIPEA (30 equiv.) in NMP at 23 °C (10 min).

**Deprotection**: 20% piperidine + 0.1 M oxyma pure in DMF (9 mL) at 23 °C (1 x 5 min, 1 x 15 min).

**Analytical Cleavage (SPPS):** A small portion of resin was treated with TFA:H2O:TA:TIS (85:5:5:5) at 23 °C for 3 h, before it was filtered and precipitated with E2O. The precipitate was centrifuged, and the resulting pellet was washed twice more with Et2O, and the pellet was air dried.

**Protected cleavage**: The resin was treated with 1% TFA in DCM (6 mL, 3 min x 10), and filtered into 10% pyridine in MeOH (60 mL). The pooled organic layers were concentrated, and protected peptide was precipitated with water, collected by centrifugation, and lyophilized to afford crude protected peptide acid.

**Thioesterification**: A 100 mL RBF with crude protected acid, PyBOP (390 mg, 0.750 mmol) and dry DMSO (10 mL) was cooled in an ice bath and treated with EtSH (0.27 mL, 3.75 mmol) and DIPEA (0.65 mL, 3.75 mmol). The reaction was slowly warmed to 23 °C overnight, and precipitated with water after 12 h. The precipitate was collected by centrifugation and taken forward to the next step.[3]

**Global Deprotection**: The crude thioester was treated with TFA:$H_2O$:thioanisole:TIPSH (85:5:5:5, 10 mL) and it was stirred at 23 °C for 3 h before it was cooled to 0 °C and treated with $Et_2O$ to precipitate the peptide. The precipitate was centrifuged, and the resulting pellet was washed twice more with $Et_2O$. It was dissolved in 40% MeCN and it was lyophilized.

**Purification**: The material was purified over an ACQUITY Protein BEH C4 column (19 x 250 mm, 5 µm, 300 Å, 17.1 mL/min) 15%-30% MeCN/$H_2O$/0.1% TFA over 30 min. The fractions were lyophilized to afford **3a** (54.0 mg, 11% yield).[3]

**Analytical LCMS**: Measured on an Agilent EC-C18 Poroshell column (4.6 x 100 mm, 4 µm, 100 Å, 40 °C, 1 mL/min) 5%-65% MeCN/$H_2O$/0.05% TFA over 30 min.

**Chemical Formula:** $C_{145}H_{242}N_{40}O_{43}S_2$
**Molecular Weight:** 3297.8880 g/mol

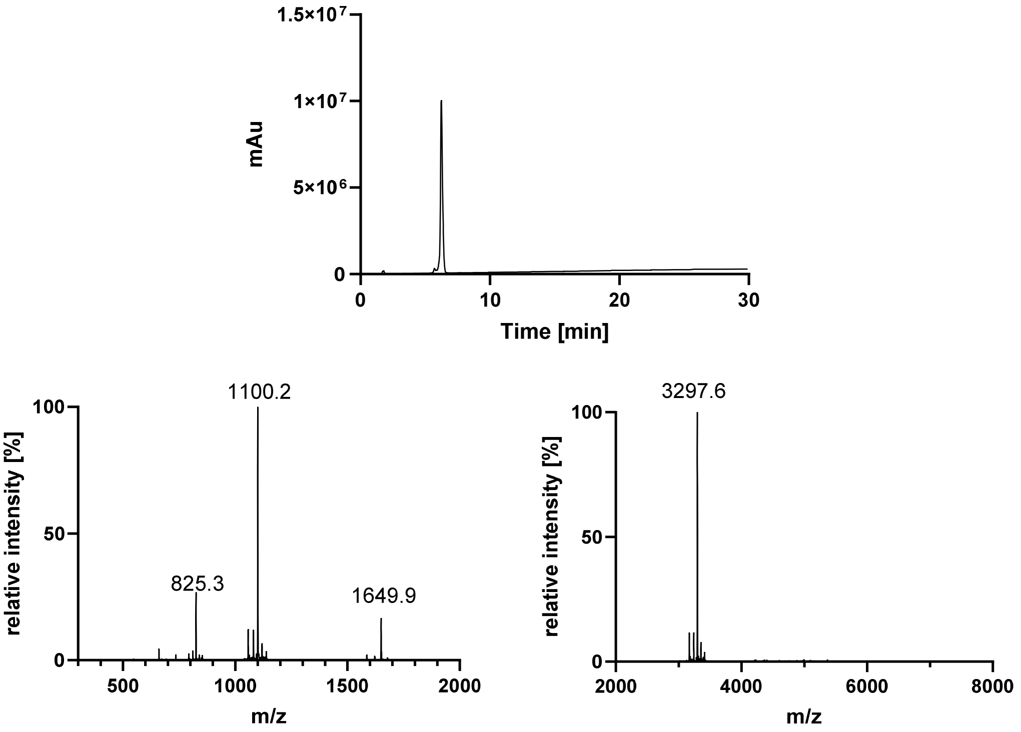

## P301S

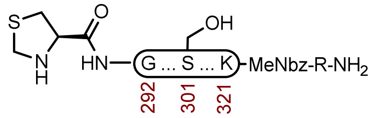

**3b**

**Sequence**: H-ThzGSKDNIKHVSGGGSVQIVYKPVDLSK(Ac)VTSK-MeNbz-R-$NH_2$

**Amino Acids**: Boc-L-Thz-OH, Fmoc-Gly-OH, Fmoc-L-Ser($^t$Bu)-OH, Fmoc-L-Lys(Boc)-OH, Fmoc-L-Asp(O$^t$Bu)-OH, Fmoc-L-Asn(Trt)-OH, Fmoc-L-Ile-OH, Fmoc-L-His(Trt)-OH, Fmoc-L-Val-OH, Fmoc-L-Pro-OH, Fmoc-L-Gln(Trt)-OH, Fmoc-L-Tyr(O$^t$Bu)-OH, Fmoc-L-Leu-OH, Fmoc-L-Lys(Ac)-OH, Fmoc-L-Thr($^t$Bu)-OH

**Resin**: Fmoc-RinkAmide Protide resin from CEM (100-200 mesh, 0.6 mmol/g, 416 mg, 0.25 mmol) was used. Fmoc-Arg(Pbf)-OH was double coupled manually: 0.2 M Fmoc-Arg(Pbf)-OH in DMF (5 equiv.), 0.5 M HATU in DMF (5 equiv.) and 1 M DIPEA (6 equiv.) was coupled at 23 ºC for 1 h Fmoc-MeDbz-OH was double coupled manually: 0.2 M Fmoc-MeDbz-OH in NMP (5 equiv.), 0.5 M HATU in DMF (5 equiv.) and 1 M DIPEA (6 equiv.) was coupled at 23 ºC for 1 h.

**Coupling**: Amino acids were double coupled using 0.25 M amino acid in DMF (5 equiv.), 1 M DIC in DMF (10 equiv.), and 2 M oxyma pure in DMF (5 equiv.) at 90°C for 4 min. Histidine was double coupled at 50°C for 10 min.

**Deprotection**: 20% piperidine + 0.1 M oxyma pure in DMF (6 mL) at 23 ºC (1 x 5 min, 1 x 30 min)

**Cyclization**: The dry peptide resin was treated with *p*-nitrophenyl chloroformate (250 mg) and a minimal amount of DCM (2 x 45 min). The resin was then treated with 1 M DIPEA in DMF (5 mL, 4 x 20 min).

**Global Deprotection**: The dry resin in a 15 mL polypropylene vial was treated with TFA:$H_2O$:thioanisole:TIPSH (85:5:5:5, 10 mL) and it was mixed on an overhead stirrer. The mixture was stirred at 23 ºC for 3 h before it was filtered, and the resin was washed with additional portion of TFA (3 x 1 mL). The pooled filtrate was cooled to 0°C and treated with $Et_2O$ to precipitate the peptide. The precipitate was centrifuged, and the resulting pellet was washed twice more with $Et_2O$. It was dissolved in 40% MeCN and it was lyophilized.

**Crude LCMS (Cyclization)**: Measured on an Agilent EC-C18 Poroshell column (4.6 x 100 mm, 4 μm, 100 Å, 40 °C, 1 mL/min) 5%-65% MeCN/$H_2O$/0.05% TFA over 60 min.

**LC trace (214 nm) of unpurified peptide**

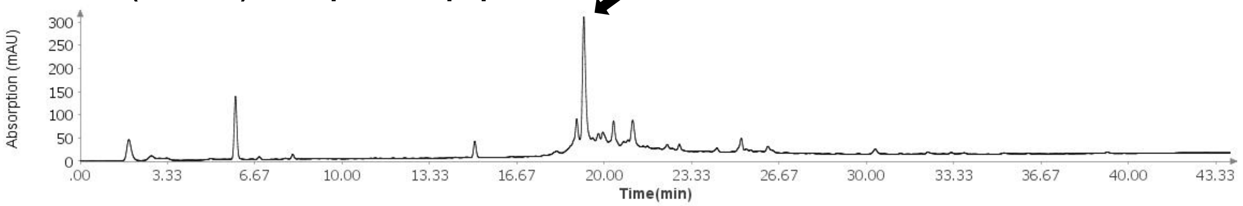

**Total Ion Current (TIC) of unpurified peptide**

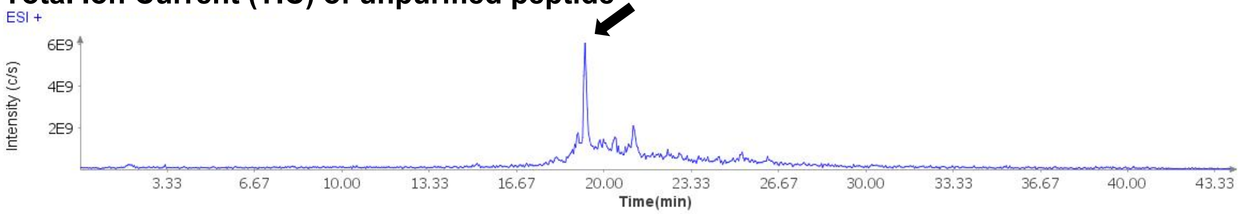

**Extracted Ion Current (XIC) of unpurified peptide**

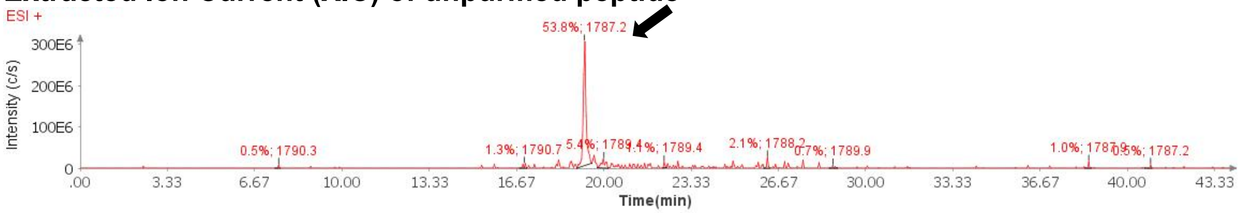

**Mass Spectrum (ESI) of the expected product (unpurified sample)**

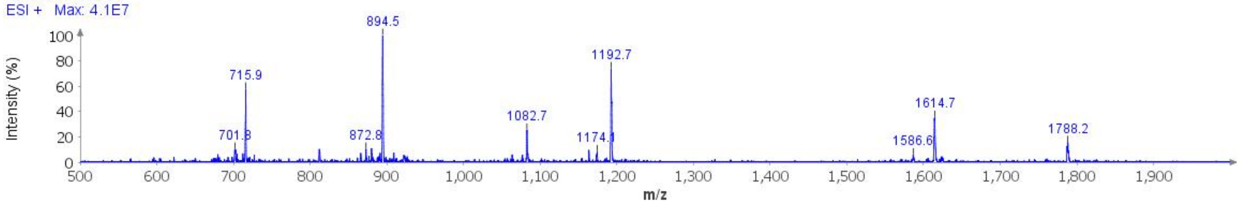

**Deconvoluted Mass Spectrum (ESI) of the expected product (unpurified sample)**

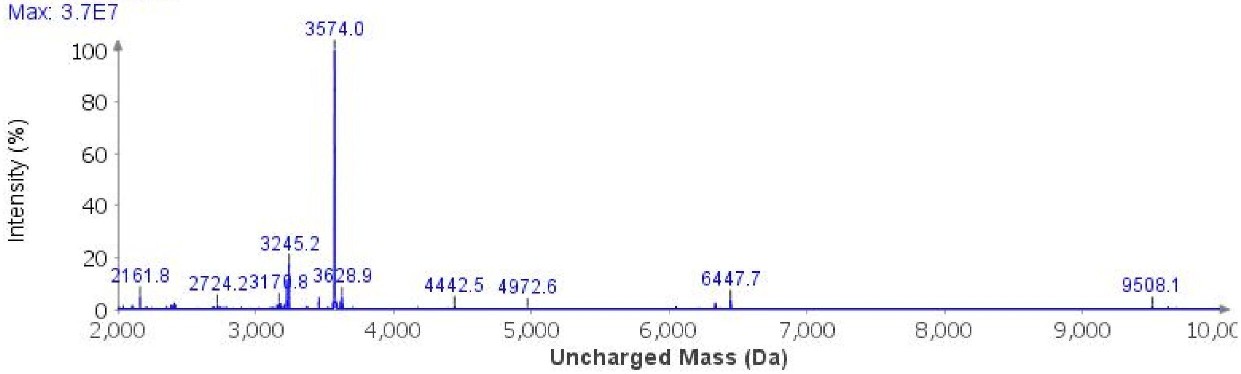

**Purification**: The material was purified over a Luna C4 column (10 x 250 mm, 10 µm, 100 Å, 4.7 mL/min) 15%-35% MeCN/$H_2O$/0.1% TFA over 30 min,. The fractions were lyophilized to afford **3b** (23.6 mg, 3% yield).

**Analytical LCMS**: Measured on an Agilent EC-C18 Poroshell column (4.6 x 100 mm, 4 µm, 100 Å, 40 °C, 1 mL/min) 5%-45% MeCN/$H_2O$/0.1% FA over 10 min.

**Chemical Formula**: $C_{156}H_{255}N_{47}O_{47}S$
**Molecular Weight**: 3573.0980 g/mol

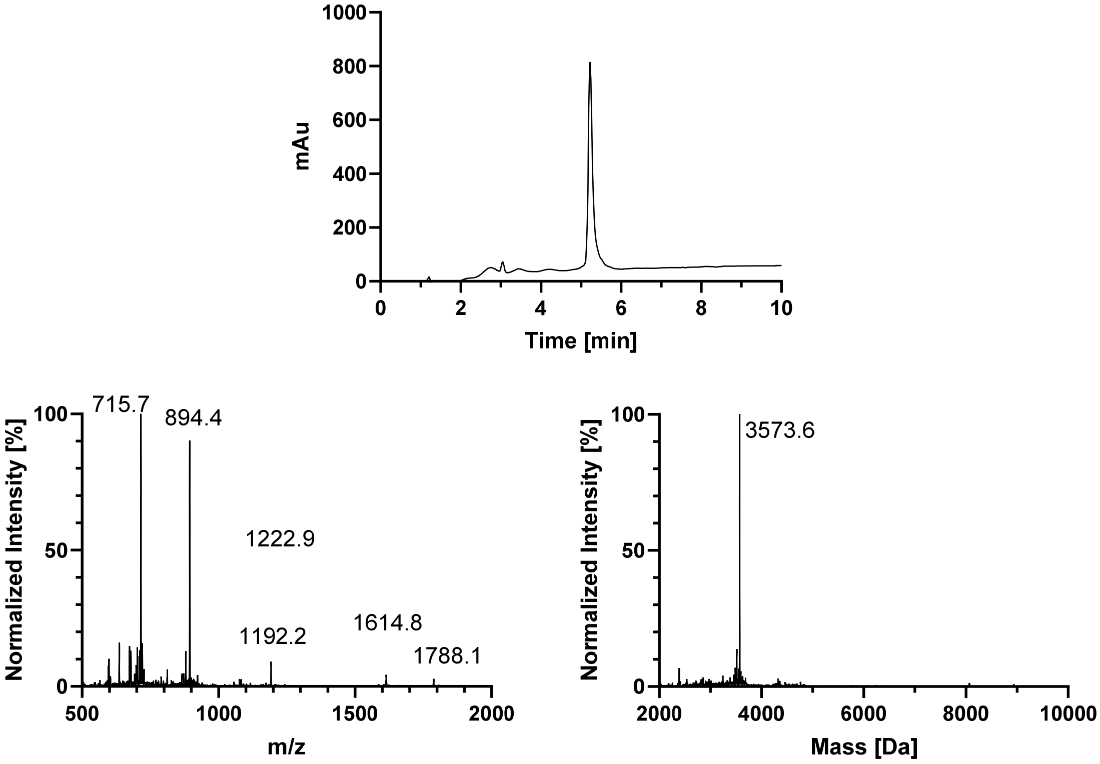
1000
800
600
400
200
0
mAu
0
2
4
6
8
10
Time [min]
715.7
894.4
1222.9
1192.2
1614.8
1788.1
100
50
0
Normalized Intensity [%]
500
1000
1500
2000
m/z
3573.6
100
50
0
Normalized Intensity [%]
2000
4000
6000
8000
10000
Mass [Da]

#### AcK317

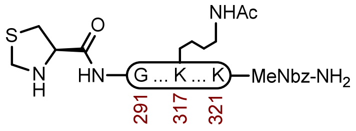

**3e**

**Sequence**: H-ThzGSKDNIKHVPGGGSVQIVYKPVDLSK(Ac)VTSK-MeNbz-R-$NH_2$

**Amino Acids**: Boc-L-Thz-OH, Fmoc-Gly-OH, Fmoc-L-Ser($^t$Bu)-OH, Fmoc-L-Lys(Boc)-OH, Fmoc-L-Asp(O$^t$Bu)-OH, Fmoc-L-Asn(Trt)-OH, Fmoc-L-Ile-OH, Fmoc-L-His(Trt)-OH, Fmoc-L-Val-OH, Fmoc-L-Pro-OH, Fmoc-L-Gln(Trt)-OH, Fmoc-L-Tyr(O$^t$Bu)-OH, Fmoc-L-Leu-OH, Fmoc-L-Lys(Ac)-OH, Fmoc-L-Thr($^t$Bu)-OH

**Resin**: H-RinkAmideChemMatrix resin from Biotage (100-200 mesh, 0.41 mmol/g, 244 mg, 0.1 mmol) was used. Fmoc-MeDbz-OH was double coupled manually: 0.2 M Fmoc-MeDbz-OH in NMP (5 equiv.), 0.5 M HATU in DMF (5 equiv.) and 1 M DIPEA (6 equiv.) was coupled at 23 ºC for 1 h. Fmoc-Lys(Boc)-OH was double coupled manually: 0.2 M Fmoc-Lys(Boc)-OH in DMF (5 equiv.), 0.5 M HATU in DMF (5 equiv.) and 1 M DIPEA (6 equiv.) was coupled at 23 ºC for 1 h.

**Coupling**: Amino acids were double coupled using 0.2 M amino acid in DMF (5 equiv.), 1 M DIC in DMF (10 equiv.), and 1 M oxyma pure in DMF (5 equiv.) at 90°C for 4 min. Histidine was double coupled at 50°C for 10 min.

**Deprotection**: 20% piperidine + 0.1 M oxyma pure in DMF (6 mL) at 23 ºC (1 x 5 min, 1 x 15 min).

**Analytical Cleavage (SPPS)**: A small portion of resin was treated with TFA:$H_2O$:TA:TIS (85:5:5:5) at 23 °C for 3 h, before it was filtered and precipitated with $E_2O$. The precipitate was centrifuged, and the resulting pellet was washed twice more with $Et_2O$, and the pellet was air dried.

**Crude LCMS (SPPS)**: Measured on an Agilent EC-C18 Poroshell column (4.6 x 100 mm, 4 µm, 100 Å, 40 °C, 1 mL/min) 5%-65% MeCN/$H_2O$/0.05% TFA over 30 min.

**LC trace (214 nm) of unpurified peptide**

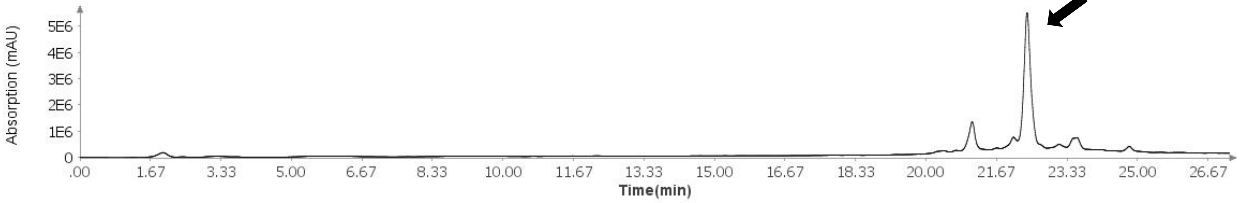

**Total Ion Current (TIC) of unpurified peptide**

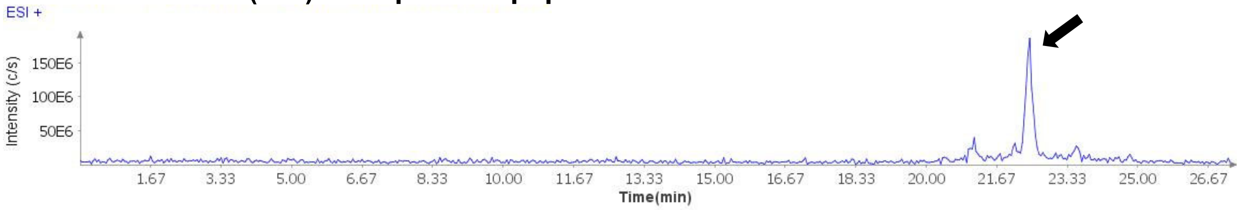

**Extracted Ion Current (XIC) of unpurified peptide**

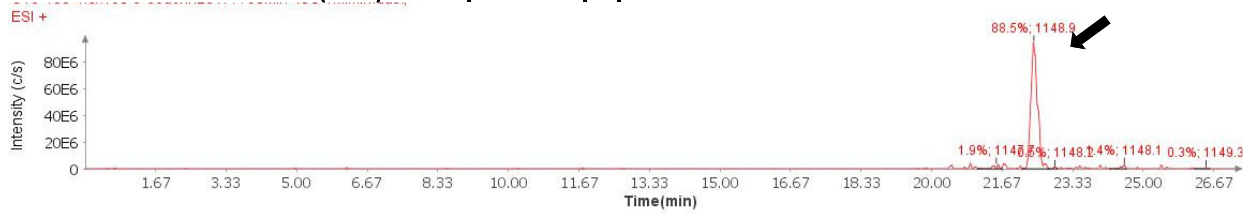

**Mass Spectrum (ESI) of the expected product (unpurified sample)**

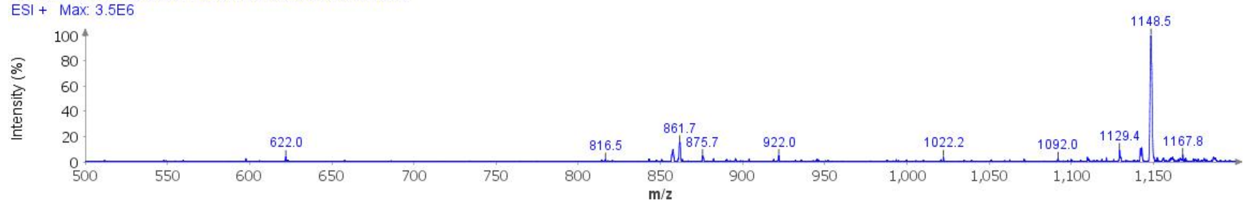

**Deconvoluted Mass Spectrum (ESI) of the expected product (unpurified sample)**

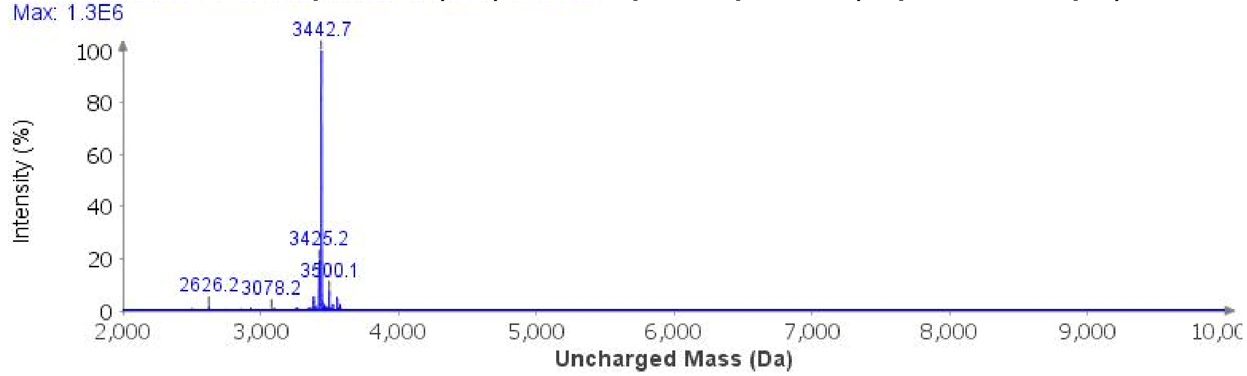

**Cyclization**: The dry peptide resin was treated with *p*-nitrophenyl chloroformate (100 mg) and a minimal amount of DCM (2 x 45 min). The resin was then treated with 1 M DIPEA in DMF (5 mL, 2 x 30 min).

**Global Deprotection**: The dry resin in a 15 mL polypropylene vial was treated with TFA:$H_2O$:thioanisole:TIPSH (85:5:5:5, 10 mL) and it was mixed on an overhead stirrer. The mixture was stirred at 23 ºC for 3 h before it was filtered, and the resin was washed with additional portion of TFA (3 x 1 mL). The pooled filtrate was cooled to 0°C and treated with $Et_2O$ to precipitate the peptide. The precipitate was centrifuged, and the resulting pellet was washed twice more with $Et_2O$. It was dissolved in 40% MeCN and it was lyophilized.

**Crude LCMS (Cyclization)**: Measured on an Agilent EC-C18 Poroshell column (4.6 x 100 mm, 4 µm, 100 Å, 40 °C, 1 mL/min) 5%-65% MeCN/$H_2O$/0.05% TFA over 30 min.

#### LC trace (214 nm) of unpurified peptide

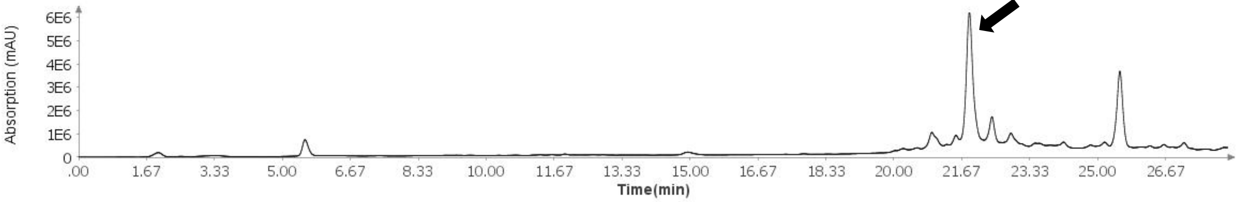

#### Total Ion Current (TIC) of unpurified peptide

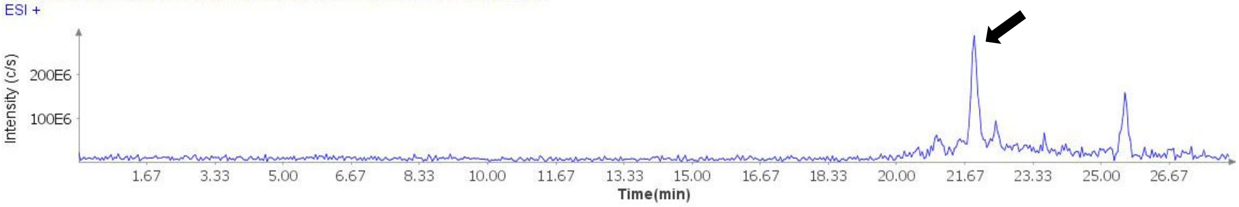

#### Extracted Ion Current (XIC) of unpurified peptide

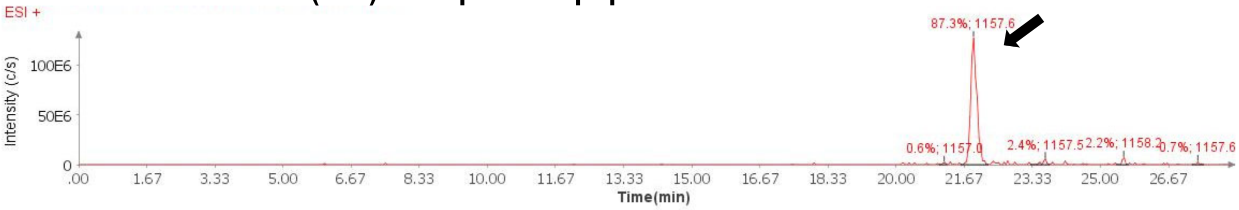

#### Mass Spectrum (ESI) of the expected product (unpurified sample)

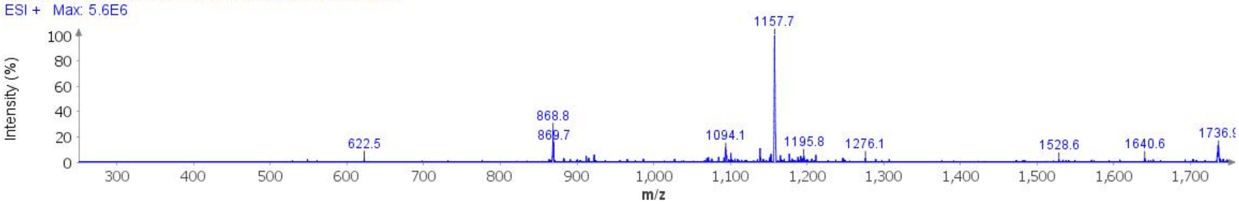

#### Deconvoluted Mass Spectrum (ESI) of the expected product (unpurified sample)

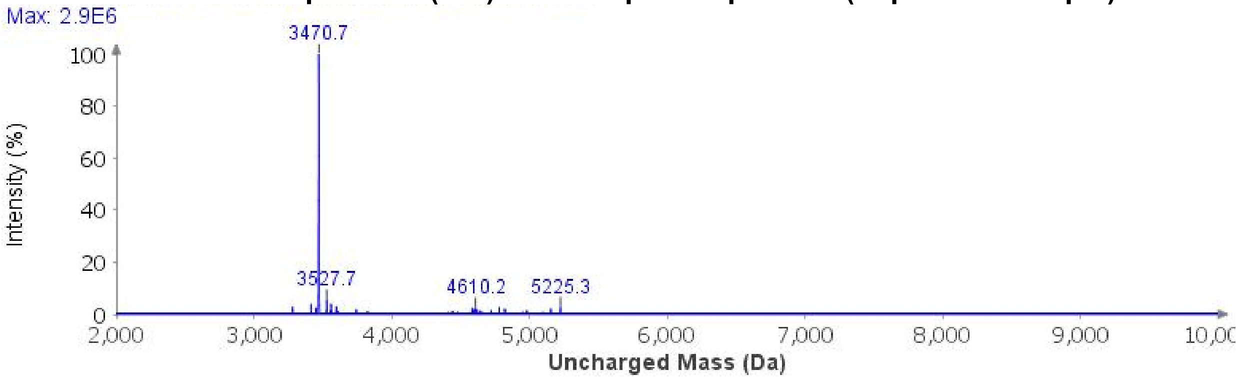

**Purification**: The material was purified over a Luna Phenyl-Hexyl column (10 x 250 mm, 5 µm, 100 Å, 4.7 mL/min) 35%-50% MeCN/$H_2O$/0.1% TFA over 30 min. The fractions were lyophilized to afford **3e** (12.6 mg, 4% yield).

**Analytical LCMS**: Measured on an Agilent EC-C18 Poroshell column (4.6 x 100 mm, 4 µm, 100 Å, 40 °C, 1 mL/min) 5%-65% MeCN/$H_2O$/0.05% TFA over 60 min.

**Chemical Formula**: $C_{154}H_{247}N_{43}O_{46}S$
**Molecular Weight**: 3468.9850 g/mol

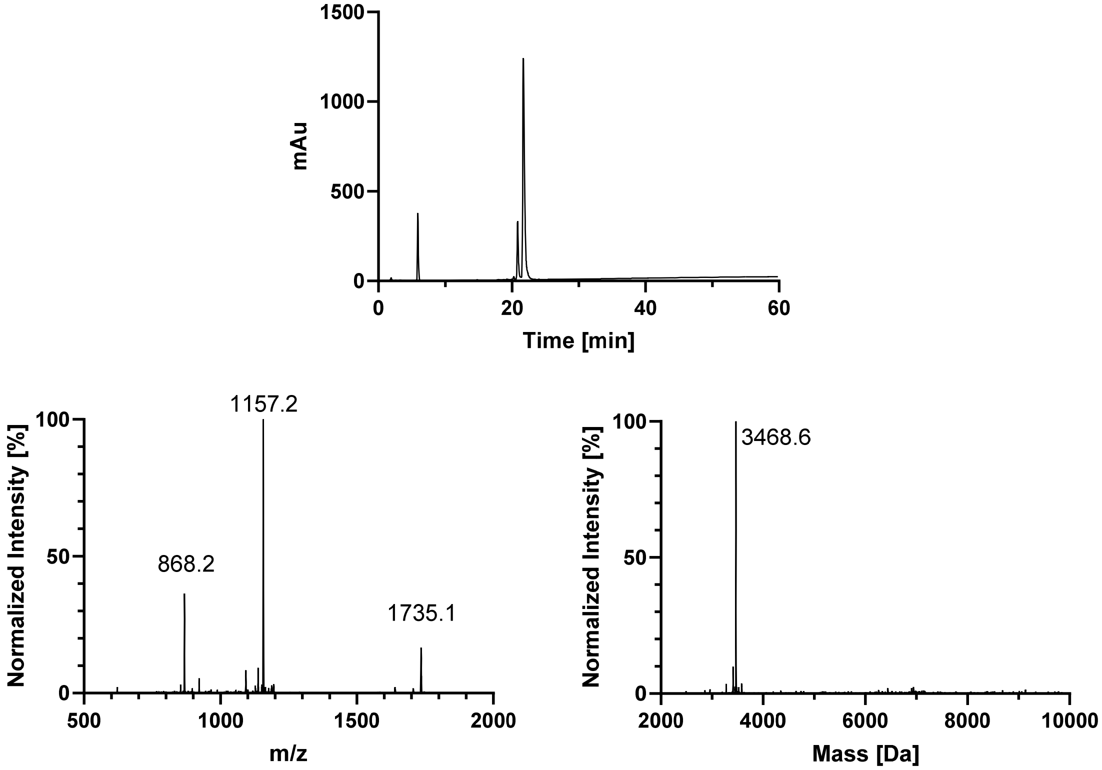

#### AcK311

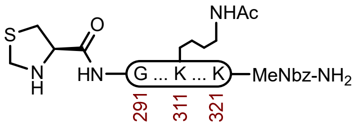

**3f**

**Sequence**: H-ThzGSKDNIKHVPGGGSVQIVYK(Ac)PVDLSKVTSK-MeNbz-R-$NH_2$

**Amino Acids**: Boc-L-Thz-OH, Fmoc-Gly-OH, Fmoc-L-Ser($^t$Bu)-OH, Fmoc-L-Lys(Boc)-OH, Fmoc-L-Asp(O$^t$Bu)-OH, Fmoc-L-Asn(Trt)-OH, Fmoc-L-Ile-OH, Fmoc-L-His(Trt)-OH, Fmoc-L-Val-OH, Fmoc-L-Pro-OH, Fmoc-L-Gln(Trt)-OH, Fmoc-L-Tyr(O$^t$Bu)-OH, Fmoc-L-Lys(Ac)-OH, Fmoc-L-Leu-OH, Fmoc-L-Thr($^t$Bu)-OH

**Resin**: H-RinkAmideChemMatrix resin from Biotage (100-200 mesh, 0.41 mmol/g, 244 mg, 0.1 mmol) was used. Fmoc-MeDbz-OH was double coupled manually: 0.2 M Fmoc-MeDbz-OH in NMP (5 equiv.), 0.5 M HATU in DMF (5 equiv.) and 1 M DIPEA (6 equiv.) was coupled at 23 ºC for 1 h. Fmoc-Lys(Boc)-OH was double coupled manually: 0.2 M Fmoc-Lys(Boc)-OH in DMF (5 equiv.), 0.5 M HATU in DMF (5 equiv.) and 1 M DIPEA (6 equiv.) was coupled at 23 ºC for 1 h.

**Coupling**: Amino acids were double coupled using 0.2 M amino acid in DMF (5 equiv.), 1 M DIC in DMF (10 equiv.), and 1 M oxyma pure in DMF (5 equiv.) at 90°C for 4 min. Histidine was double coupled at 50°C for 10 min.

**Deprotection**: 20% piperidine + 0.1 M oxyma pure in DMF (6 mL) at 23 ºC (1 x 5 min, 1 x 15 min).

**Analytical Cleavage (SPPS)**: A small portion of resin was treated with TFA:$H_2O$:TA:TIS (85:5:5:5) at 23 °C for 3 h, before it was filtered and precipitated with $E_2O$. The precipitate was centrifuged, and the resulting pellet was washed twice more with $Et_2O$, and the pellet was air dried.

**Crude LCMS (SPPS)**: Measured on an Agilent EC-C18 Poroshell column (4.6 x 100 mm, 4 µm, 100 Å, 40 °C, 1 mL/min) 5%-65% MeCN/$H_2O$/0.05% TFA over 30 min.

**LC trace (214 nm) of unpurified peptide**

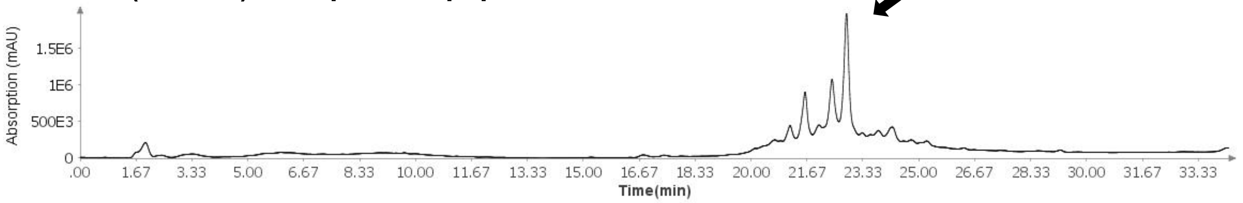

**Total Ion Current (TIC) of unpurified peptide**

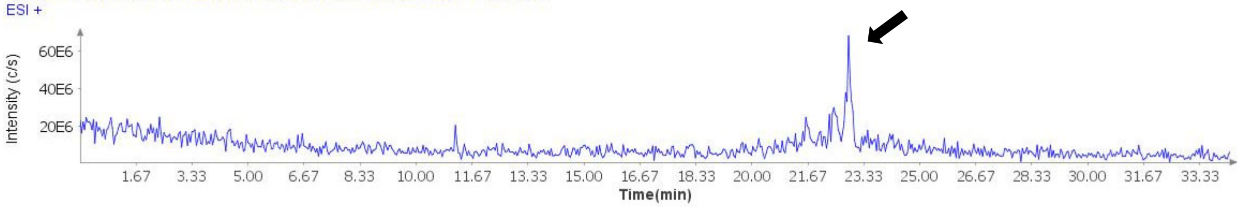

**Extracted Ion Current (XIC) of unpurified peptide**

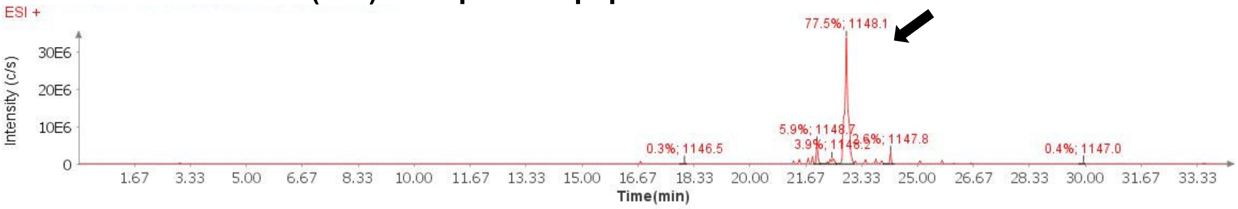

**Mass Spectrum (ESI) of the expected product (unpurified sample)**

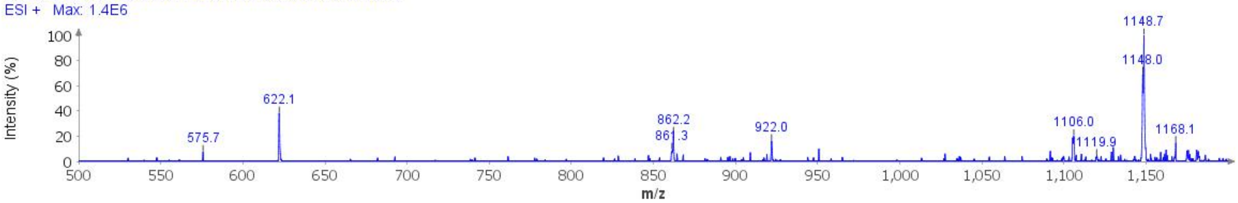

**Deconvoluted Mass Spectrum (ESI) of the expected product (unpurified sample)**

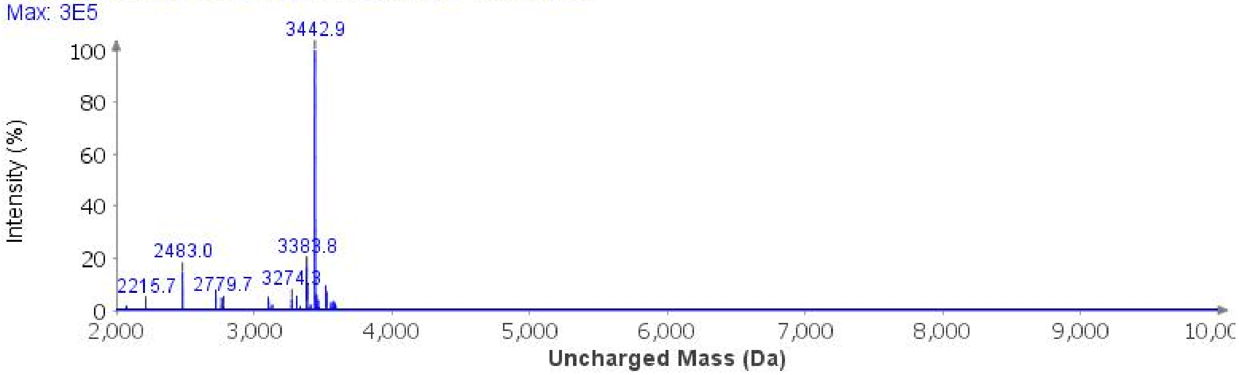

**Cyclization**: The dry peptide resin was treated with *p*-nitrophenyl chloroformate (100 mg) and a minimal amount of DCM (2 x 45 min). The resin was then treated with 1 M DIPEA in DMF (5 mL, 2 x 15 min).

**Global Deprotection**: The dry resin in a 15 mL polypropylene vial was treated with TFA:$H_2O$:thioanisole:TIPSH (85:5:5:5, 10 mL) and it was mixed on an overhead stirrer. The mixture was stirred at 23 ºC for 3 h before it was filtered, and the resin was washed with additional portion of TFA (3 x 1 mL). The pooled filtrate was cooled to 0°C and treated with $Et_2O$ to precipitate the peptide. The precipitate was centrifuged, and the resulting pellet was washed twice more with $Et_2O$. It was dissolved in 40% MeCN and it was lyophilized.

**Crude LCMS (Cyclization)**: Measured on an Agilent EC-C18 Poroshell column (4.6 x 100 mm, 4 µm, 100 Å, 40 °C, 1 mL/min) 5%-65% MeCN/$H_2O$/0.05% TFA over 30 min.

**LC trace (214 nm) of unpurified peptide**

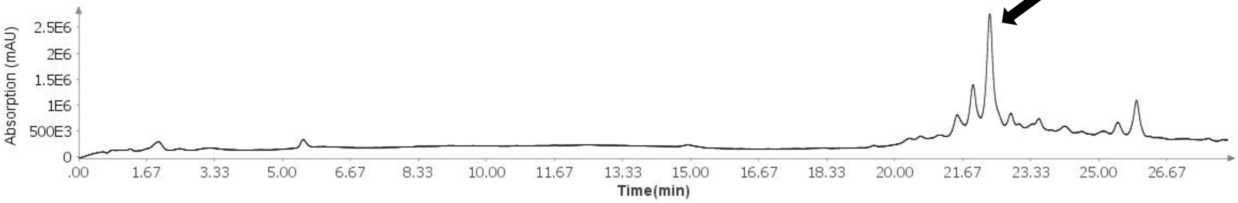

**Total Ion Current (TIC) of unpurified peptide**

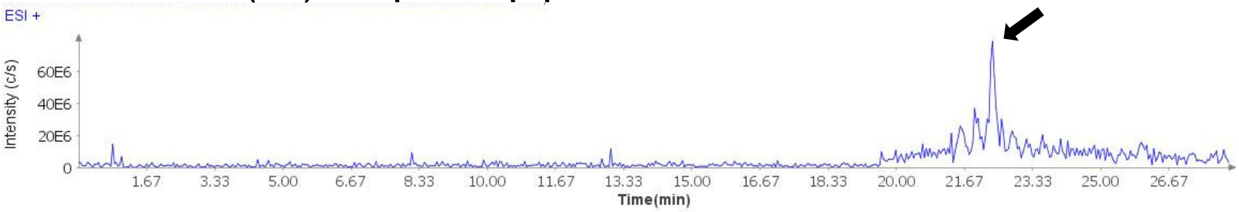

**Extracted Ion Current (XIC) of unpurified peptide**

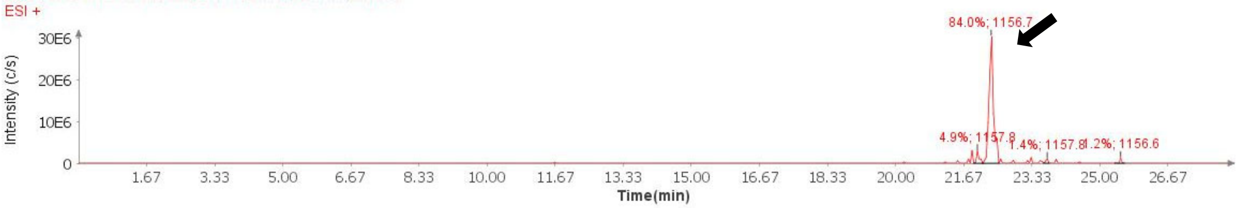

**Mass Spectrum (ESI) of the expected product (unpurified sample)**

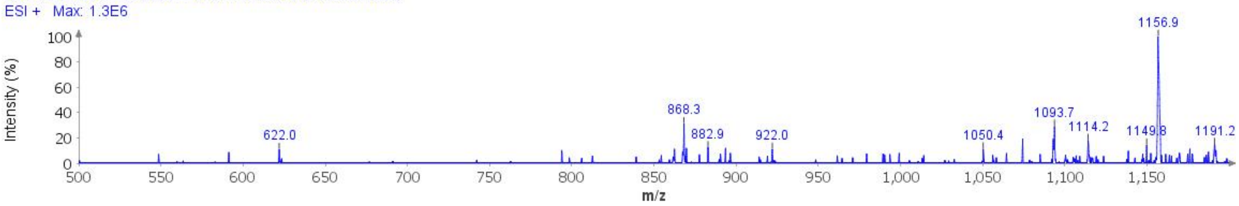

**Deconvoluted Mass Spectrum (ESI) of the expected product (unpurified sample)**

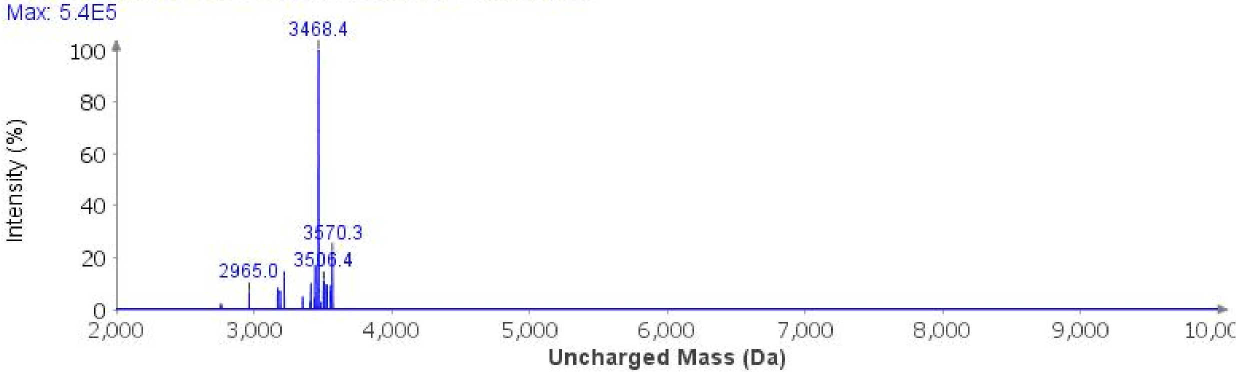

**Purification**: The material was purified over a Luna Phenyl-Hexyl column (10 x 250 mm, 5 µm, 100 Å, 4.7 mL/min) 30-55% MeCN/$H_2O$/0.1% TFA over 30 min. The fractions were lyophilized to afford **3f** (13.9 mg, 4% yield).

**Analytical LCMS**: Measured on an Agilent EC-C18 Poroshell column (4.6 x 100 mm, 4 µm, 100 Å, 40 °C, 1 mL/min) 5%-65% MeCN/$H_2O$/0.05% TFA over 60 min.

**Chemical Formula**: $C_{154}H_{247}N_{43}O_{46}S$
**Molecular Weight**: 3468.9850 g/mol

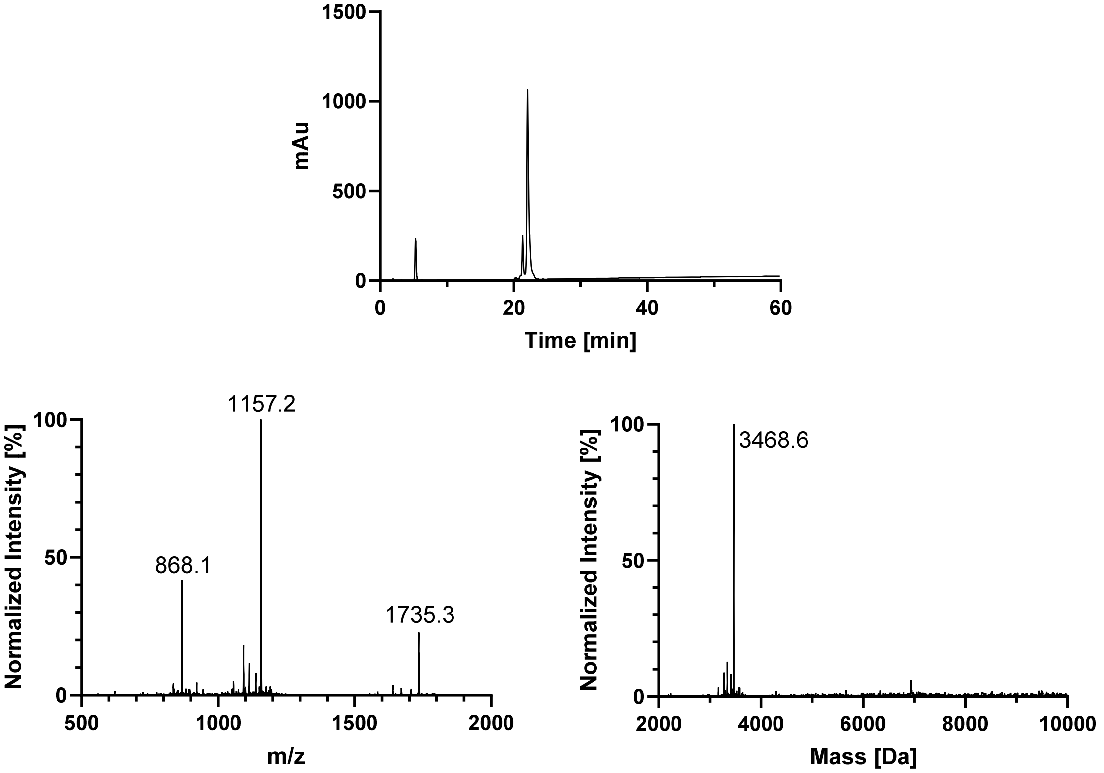

## pY310

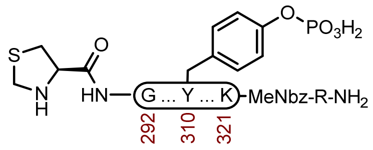

**3i**

**Sequence**: H-ThzGSKDNIKHVPGGGSVQIVY($OPO_3H_2$)KPVDLSKVTSK-MeNbz-R-$NH_2$

**Amino Acids**: Boc-L-Thz-OH, Fmoc-Gly-OH, Fmoc-L-Ser($^t$Bu)-OH, Fmoc-L-Lys(Boc)-OH, Fmoc-L-Asp(O$^t$Bu)-OH, Fmoc-L-Asn(Trt)-OH, Fmoc-L-Ile-OH, Fmoc-L-His(Trt)-OH, Fmoc-L-Val-OH, Fmoc-L-Pro-OH, Fmoc-L-Tyr($HPO_3$Bn)-OH, Fmoc-L-Gln(Trt)-OH, Fmoc-L-Tyr(O$^t$Bu)-OH, Fmoc-L-Leu-OH, Fmoc-L-Thr($^t$Bu)-OH

**Resin**: Fmoc-RinkAmide Protide resin from CEM (100-200 mesh, 0.59 mmol/g, 423 mg, 0.25 mmol) was used. Fmoc-Arg(Pbf)-OH was double coupled manually: 0.2 M Fmoc-Arg(Pbf)-OH in DMF (5 equiv.), 0.5 M HATU in DMF (5 equiv.) and 1 M DIPEA (6 equiv.) was coupled at 23 ºC for 1 h Fmoc-MeDbz-OH was double coupled manually: 0.2 M Fmoc-MeDbz-OH in NMP (5 equiv.), 0.5 M HATU in DMF (5 equiv.) and 1 M DIPEA (6 equiv.) was coupled at 23 ºC for 1 h. Fmoc-Lys(Boc)-OH was double coupled manually: 0.2 M Fmoc-Lys(Boc)-OH in DMF (5 equiv.), 0.5 M HATU in DMF (5 equiv.) and 1 M DIPEA (6 equiv.) was coupled at 23 ºC for 1 h.

**Coupling**: Amino acids were double coupled using 0.25 M amino acid in DMF (5 equiv.), 1 M DIC in DMF (10 equiv.), and 1 M oxyma pure in DMF (5 equiv.) at 90°C for 4 min. Histidine was double coupled at 50°C for 10 min.

**Coupling Fmoc-L-Tyr($HPO_3$Bn)-OH**: Fmoc-L-Tyr($HPO_3$Bn)-OH (2.5 equiv.), HATU (2.5 equiv.), Oxyma (2.5 equiv.), DIPEA (7 equiv.) was preactivated in DMF and added to the resin. Mixed at 25 °C for 16 h.

**Deprotection**: 20% piperidine + 0.1 M oxyma pure in DMF (6 mL) at 23 ºC (1 x 5 min, 1 x 15 min).

**Analytical Cleavage (SPPS)**: A small portion of resin was treated with TFA:$H_2O$:TA:TIS (85:5:5:5) at 23 °C for 3 h, before it was filtered and precipitated with $E_2O$. The precipitate was centrifuged, and the resulting pellet was washed twice more with $Et_2O$, and the pellet was air dried.

**Crude LCMS (SPPS)**: Measured on an Agilent EC-C18 Poroshell column (4.6 x 100 mm, 4 μm, 100 Å, 40 °C, 1 mL/min) 5%-65% MeCN/$H_2O$/0.05% TFA over 60 min.

**LC trace (214 nm) of unpurified peptide**

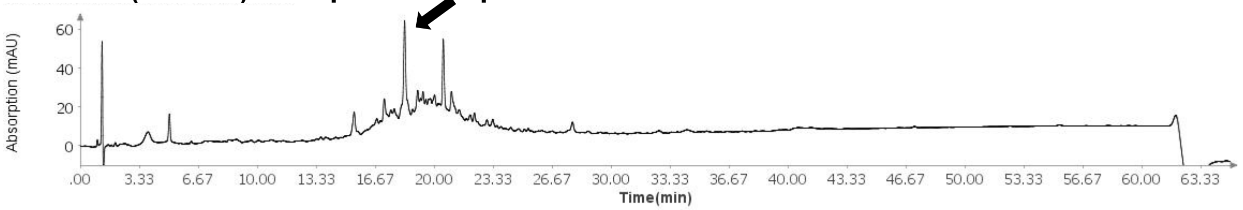

**Total Ion Current (TIC) of unpurified peptide**

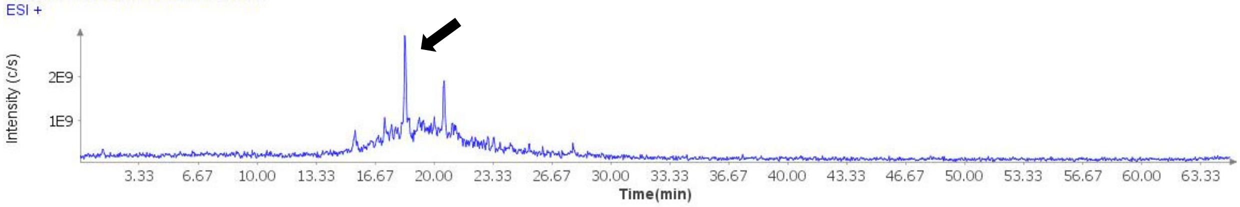

**Extracted Ion Current (XIC) of unpurified peptide**

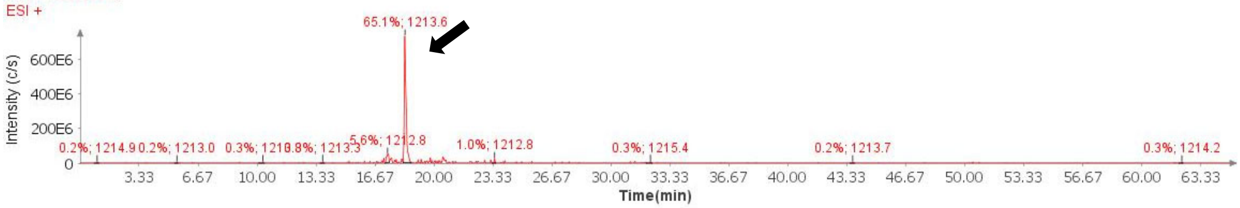

**Mass Spectrum (ESI) of the expected product (unpurified sample)**

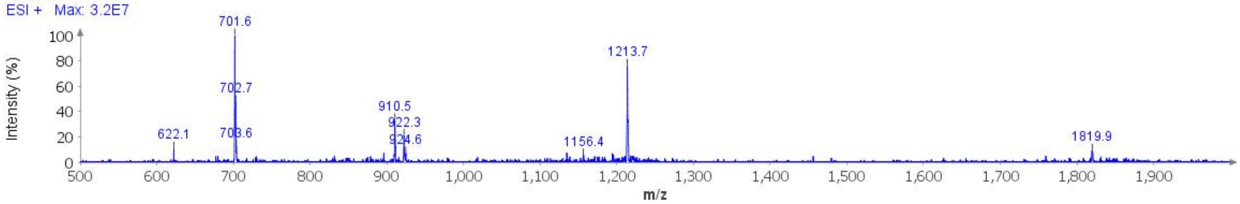

**Deconvoluted Mass Spectrum (ESI) of the expected product (unpurified sample)**

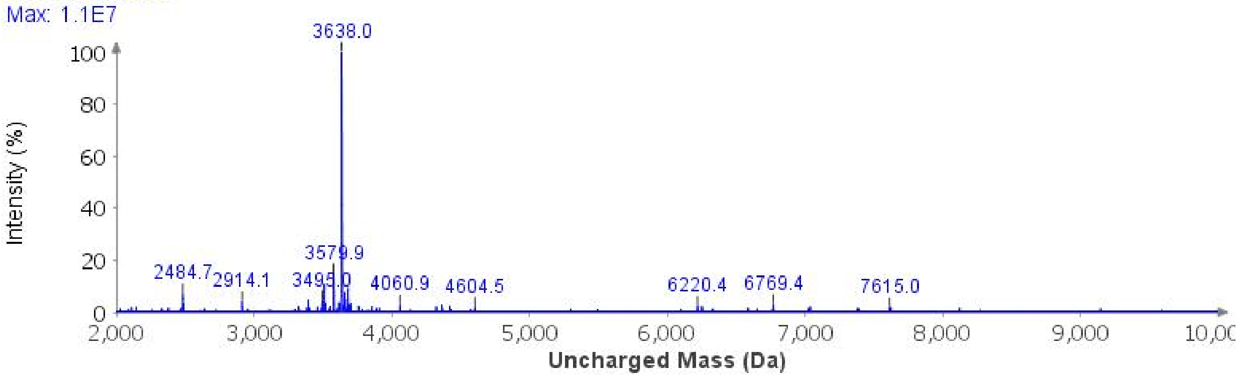

**Cyclization**: The dry peptide resin was treated with *p*-nitrophenyl chloroformate (250 mg) and a minimal amount of DCM (2 x 45 min). The resin was then treated with 1 M DIPEA in DMF (10 mL, 3 x 30 min).

**Global Deprotection**: The dry resin in a 25 mL polypropylene vial was treated with TFA:$H_2O$:thioanisole:TIPSH (85:5:5:5, 20 mL) and it was mixed on an overhead stirrer. The mixture was stirred at 23 ºC for 3 h before it was filtered, and the resin was washed with additional portion of TFA (3 x 1 mL). The pooled filtrate was cooled to 0°C and treated with $Et_2O$ to precipitate the peptide. The precipitate was centrifuged, and the resulting pellet was washed twice more with $Et_2O$. It was dissolved in 40% MeCN and it was lyophilized.

**Crude LCMS (Cyclization)**: Measured on an Agilent EC-C18 Poroshell column (4.6 x 100 mm, 4 µm, 100 Å, 40 °C, 1 mL/min) 5%-65% MeCN/$H_2O$/0.05% TFA over 60 min.

**LC trace (214 nm) of unpurified peptide**

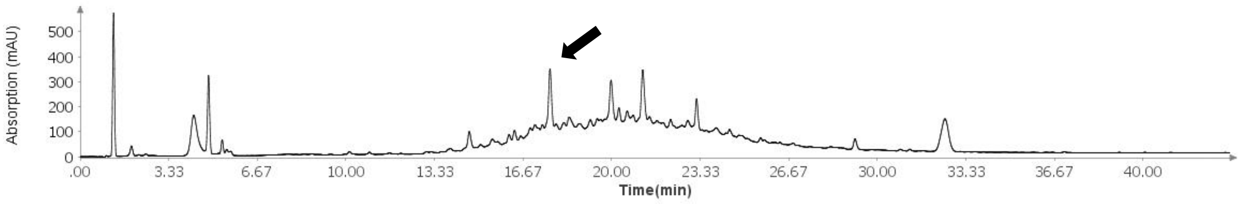

**Total Ion Current (TIC) of unpurified peptide**

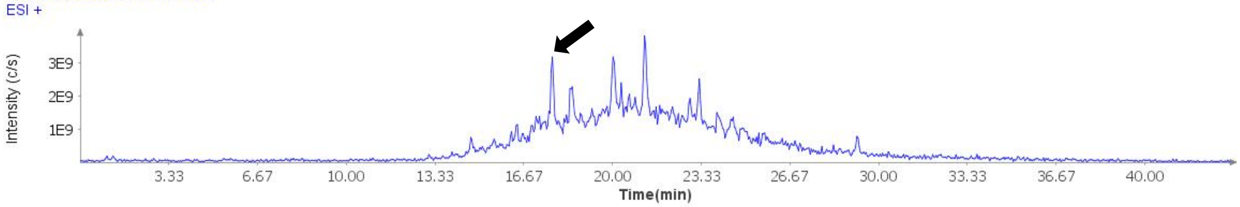

**Extracted Ion Current (XIC) of unpurified peptide**

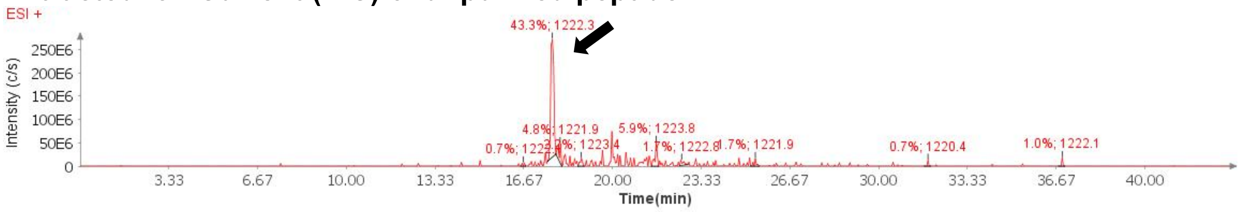

**Mass Spectrum (ESI) of the expected product (unpurified sample)**

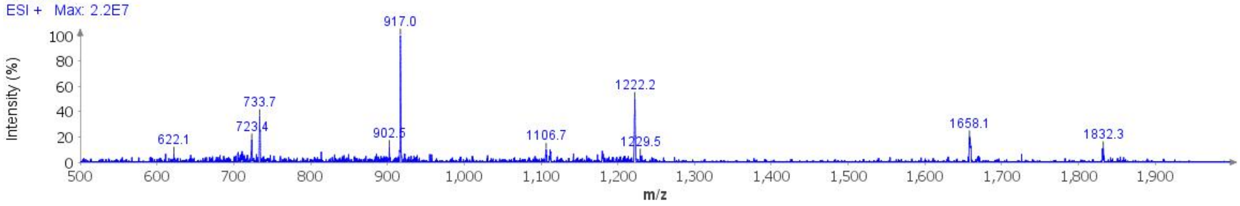

**Deconvoluted Mass Spectrum (ESI) of the expected product (unpurified sample)**

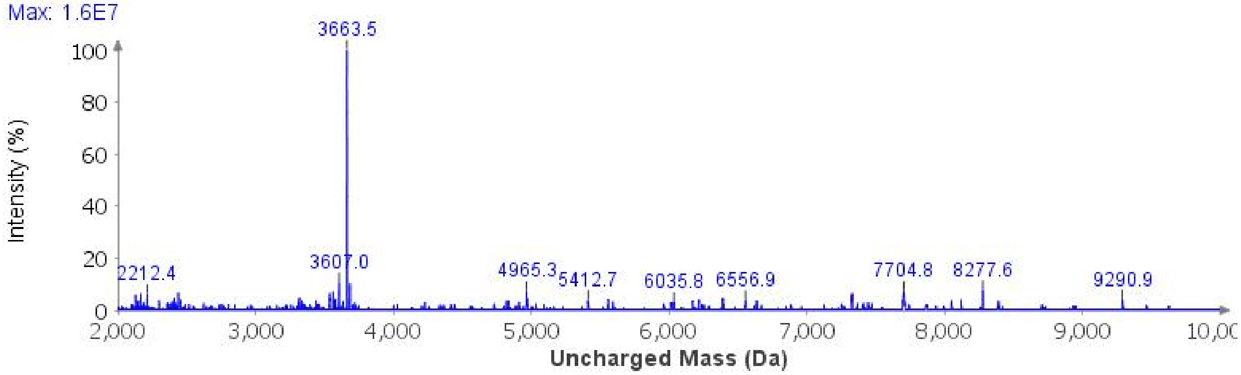

**Purification**: The material was purified over a Luna Phenyl-Hexyl column (10 x 250 mm, 5 µm, 100 Å, 4.7 mL/min) 15%-23% MeCN/$H_2O$/0.1% TFA over 30 min. The fractions were lyophilized to afford **3i** (20.7 mg, 2% yield).

**Analytical LCMS**: Measured on an Agilent EC-C18 Poroshell column (4.6 x 100 mm, 4 µm, 100 Å, 40 °C, 1 mL/min) 5%-40% MeCN/$H_2O$/0.1% FA over 10 min.

**Chemical Formula**: $C_{158}H_{258}N_{47}O_{49}PS$
**Molecular Weight**: 3663.1158 g/mol

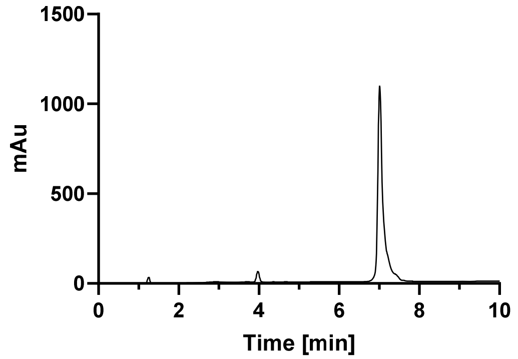

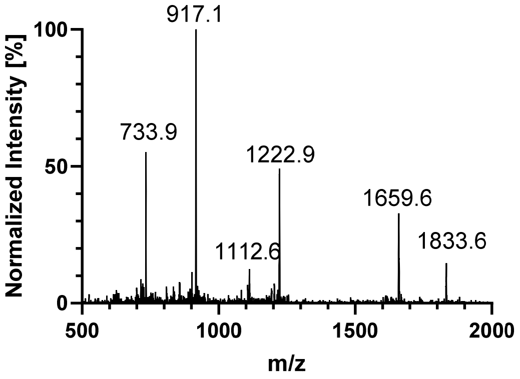

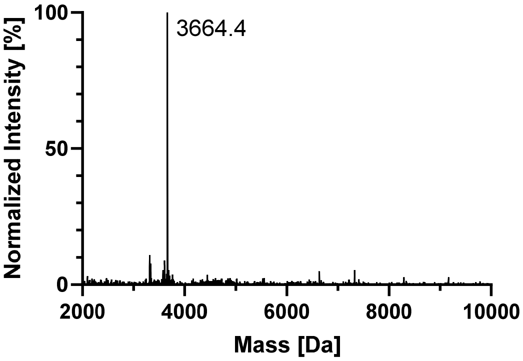

## pS305

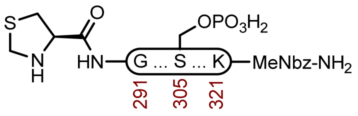

**3j**

**Sequence**: H-ThzGSKDNIKHVPGGGS($OPO_3H_2$)VQIVYKPVDLSKVTSK-MeNbz-R-$NH_2$

**Amino Acids**: Boc-L-Thz-OH, Fmoc-Gly-OH, Fmoc-L-Ser($^t$Bu)-OH, Fmoc-L-Lys(Boc)-OH, Fmoc-L-Asp(O$^t$Bu)-OH, Fmoc-L-Asn(Trt)-OH, Fmoc-L-Ile-OH, Fmoc-L-His(Trt)-OH, Fmoc-L-Val-OH, Fmoc-L-Pro-OH, Fmoc-L-Ser($HPO_3$Bn)-OH, Fmoc-L-Gln(Trt)-OH, Fmoc-L-Tyr(O$^t$Bu)-OH, Fmoc-L-Leu-OH, Fmoc-L-Thr($^t$Bu)-OH

**Resin**: H-RinkAmideChemMatrix resin from Biotage (100-200 mesh, 0.41 mmol/g, 244 mg, 0.1 mmol) was used. Fmoc-MeDbz-OH was double coupled manually: 0.2 M Fmoc-MeDbz-OH in NMP (5 equiv.), 0.5 M HATU in DMF (5 equiv.) and 1 M DIPEA (6 equiv.) was coupled at 23 ºC for 1 h. Fmoc-Lys(Boc)-OH was double coupled manually: 0.2 M Fmoc-Lys(Boc)-OH in DMF (5 equiv.), 0.5 M HATU in DMF (5 equiv.) and 1 M DIPEA (6 equiv.) was coupled at 23 ºC for 1 h.

**Coupling**: Amino acids were double coupled using 0.2 M amino acid in DMF (5 equiv.), 1 M DIC in DMF (10 equiv.), and 1 M oxyma pure in DMF (5 equiv.) at 90°C for 4 min. Histidine was double coupled at 50°C for 10 min.

**Coupling Fmoc-L-Ser($HPO_3$Bn)-OH**: Fmoc-L-Ser($HPO_3$Bn)-OH (2.5 equiv.), HATU (2.5 equiv.), Oxyma (2.5 equiv.), DIPEA (7 equiv.) was preactivated in DMF and added to the resin. Mixed at 25 °C for 16 h.

**Deprotection**: 20% piperidine + 0.1 M oxyma pure in DMF (6 mL) at 23 ºC (1 x 5 min, 1 x 15 min).

**Analytical Cleavage (SPPS)**: A small portion of resin was treated with TFA:$H_2O$:TA:TIS (85:5:5:5) at 23 °C for 3 h, before it was filtered and precipitated with $E_2O$. The precipitate was centrifuged, and the resulting pellet was washed twice more with $Et_2O$, and the pellet was air dried.

**Crude LCMS (SPPS)**: Measured on an Agilent EC-C18 Poroshell column (4.6 x 100 mm, 4 µm, 100 Å, 40 °C, 1 mL/min) 5%-65% MeCN/$H_2O$/0.05% TFA over 60 min.

**LC trace (214 nm) of unpurified peptide**

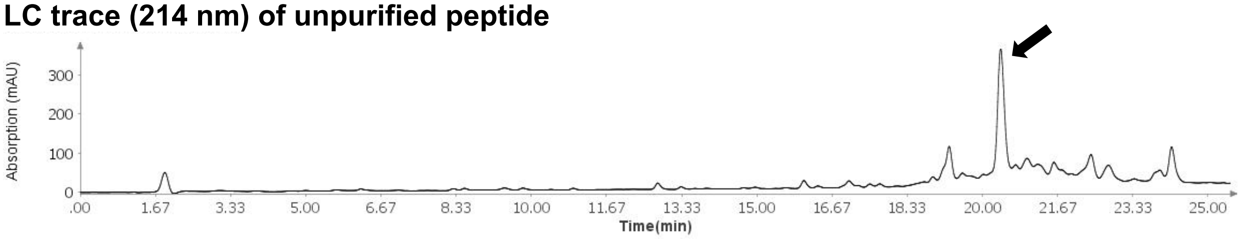

**Total Ion Current (TIC) of unpurified peptide**

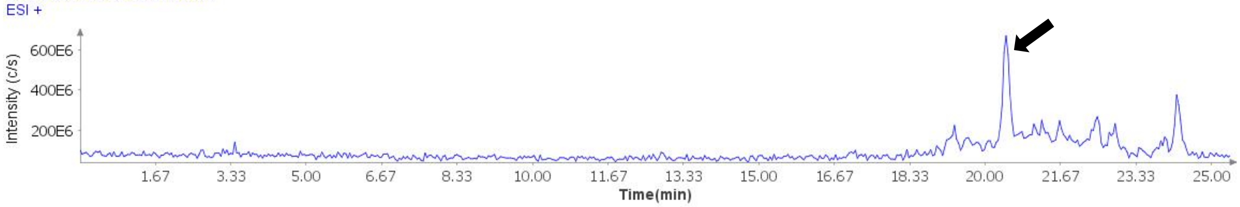

**Extracted Ion Current (XIC) of unpurified peptide**

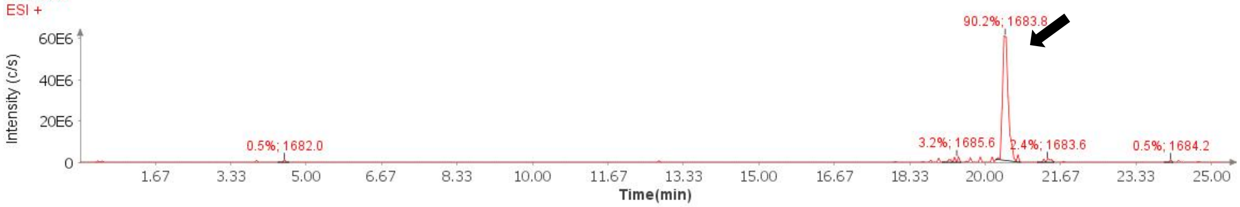

**Mass Spectrum (ESI) of the expected product (unpurified sample)**

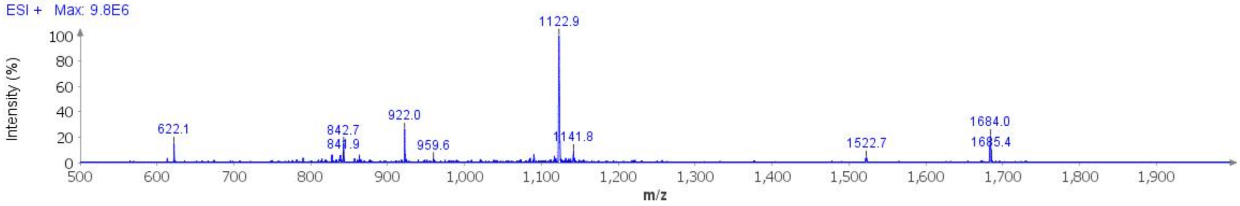

**Deconvoluted Mass Spectrum (ESI) of the expected product (unpurified sample)**

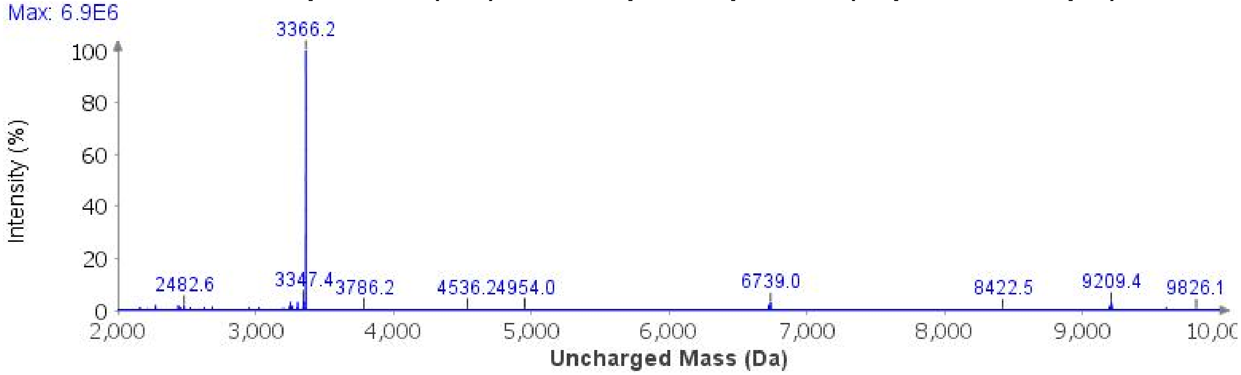

**Cyclization**: The dry peptide resin was treated with *p*-nitrophenyl chloroformate (100 mg) and a minimal amount of DCM (2 x 45 min). The resin was then treated with 1 M DIPEA in DMF (5 mL, 2 x 30 min).

**Global Deprotection**: The dry resin in a 15 mL polypropylene vial was treated with TFA:$H_2O$:thioanisole:TIPSH (85:5:5:5, 10 mL) and it was mixed on an overhead stirrer. The mixture was stirred at 23 ºC for 3 h before it was filtered, and the resin was washed with additional portion of TFA (3 x 1 mL). The pooled filtrate was cooled to 0°C and treated with $Et_2O$ to precipitate the peptide. The precipitate was centrifuged, and the resulting pellet was washed twice more with $Et_2O$. It was dissolved in 40% MeCN and it was lyophilized.

**Crude LCMS (Cyclization)**: Measured on an Agilent EC-C18 Poroshell column (4.6 x 100 mm, 4 µm, 100 Å, 40 °C, 1 mL/min) 5%-65% MeCN/$H_2O$/0.05% TFA over 60 min.

##### LC trace (214 nm) of unpurified peptide

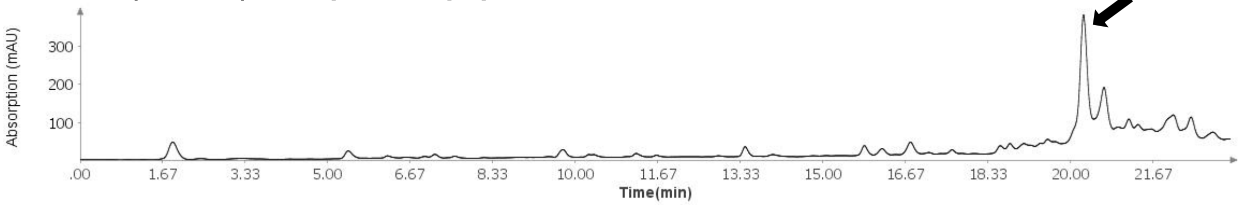

##### Total Ion Current (TIC) of unpurified peptide

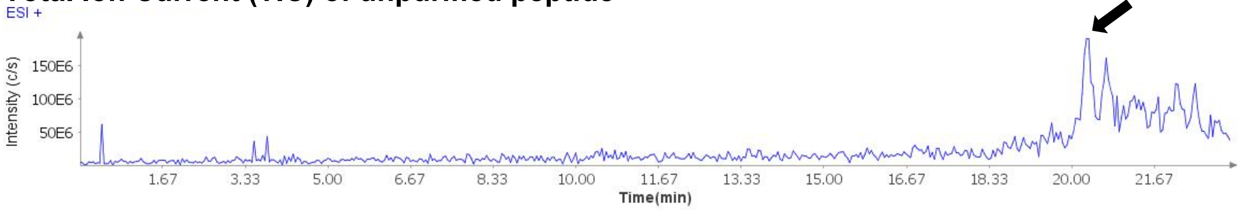

##### Extracted Ion Current (XIC) of unpurified peptide

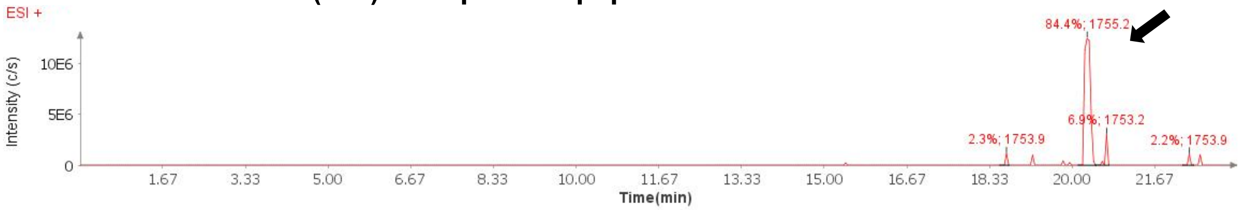

##### Mass Spectrum (ESI) of the expected product (unpurified sample)

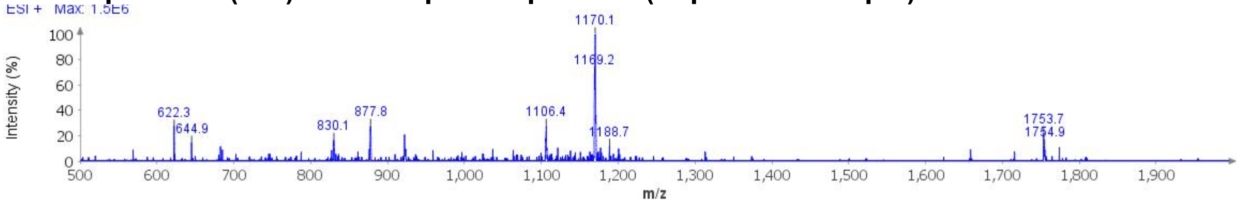

##### Deconvoluted Mass Spectrum (ESI) of the expected product (unpurified sample)

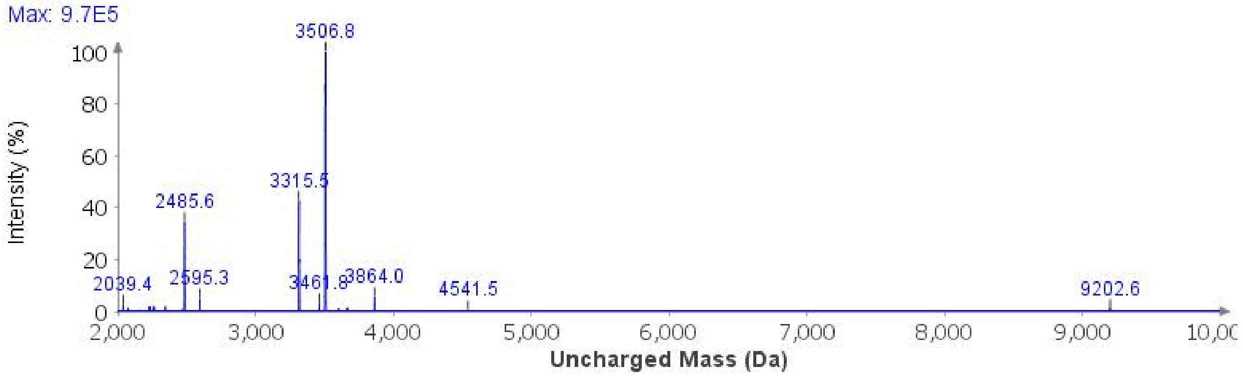

**Purification**: The material was purified over a Luna Phenyl-Hexyl column (10 x 250 mm, 5 µm, 100 Å, 4.7 mL/min) 15%-40% MeCN/$H_2O$/0.1% TFA over 30 min. The fractions were lyophilized to afford **3j** (13.7 mg, 4% yield).

**Analytical LCMS**: Measured on an Agilent EC-C18 Poroshell column (4.6 x 100 mm, 4 µm, 100 Å, 40 °C, 1 mL/min) 5%-35% MeCN/$H_2O$/0.1% FA over 10 min.

**Chemical Formula**: $C_{152}H_{246}N_{43}O_{48}PS$
**Molecular Weight**: 3506.9268 g/mol

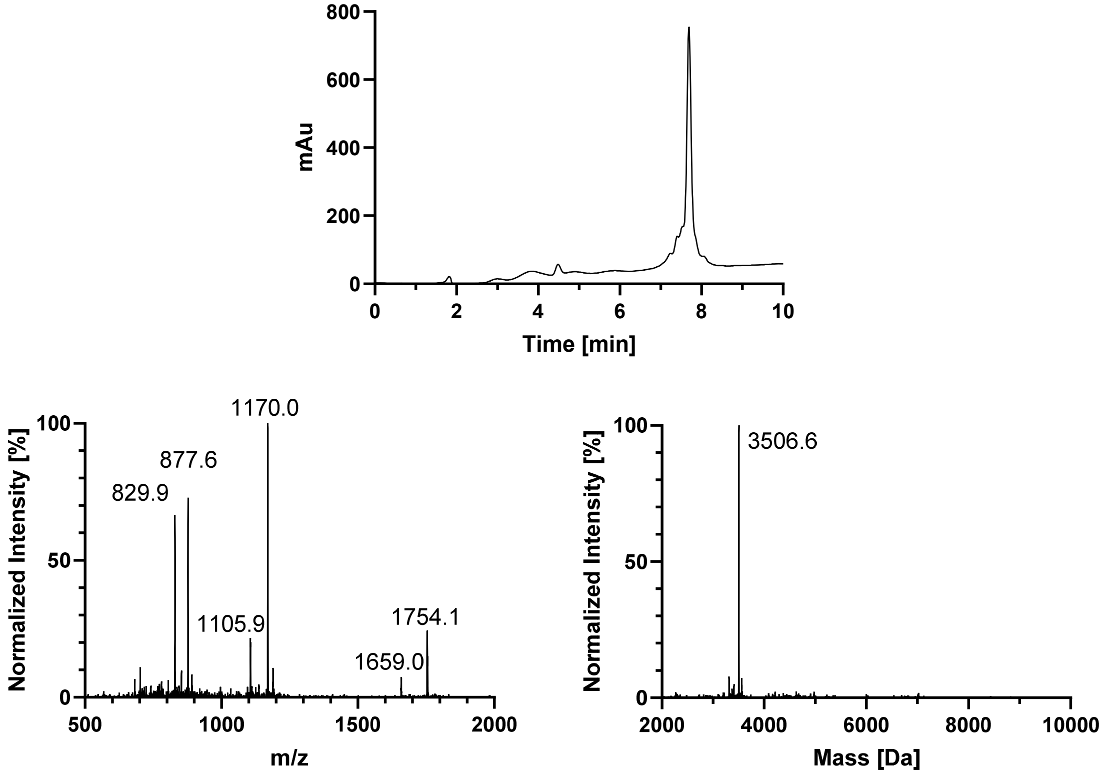

#### 5. D358-E391 (DISELENIDE SYNTHESIS)

### WT

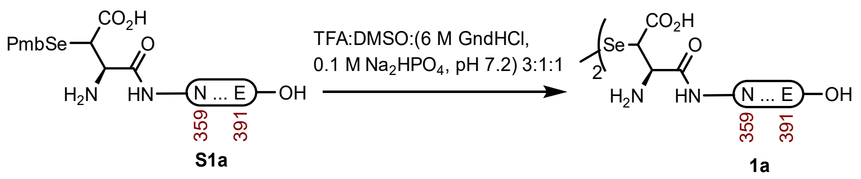

**Peptide 1a.** This diselenide synthesis method was modified from the literature.[1] A 2-dram vial containing **S1a** (45.5 mg, 11.6 µmol) was dissolved in buffer (400 µL, 6 M GndHCl, 0.1 M $Na_2HPO_4$, pH 7.2) and DMSO (400 µL). The mixture was cooled in an ice bath and treated with TFA (1.2 mL), wrapped with Al-Foil, and mixed on an overhead stirrer. The reaction was stirred at 23 °C for 45 min before it was purified over a Luna C4 column (10 x 250 mm, 10 µm, 100 Å, 4.7 mL/min) 15%-35% MeCN/$H_2O$/0.1% TFA over 30 min. The fractions were lyophilized* to afford **1a** (38.3 mg, 87% yield).

* The fractions were lyophilized immediately and protected from light to prevent deselenization (reduction of diselenide into alanine).

**Prep HPLC trace of diselenide**

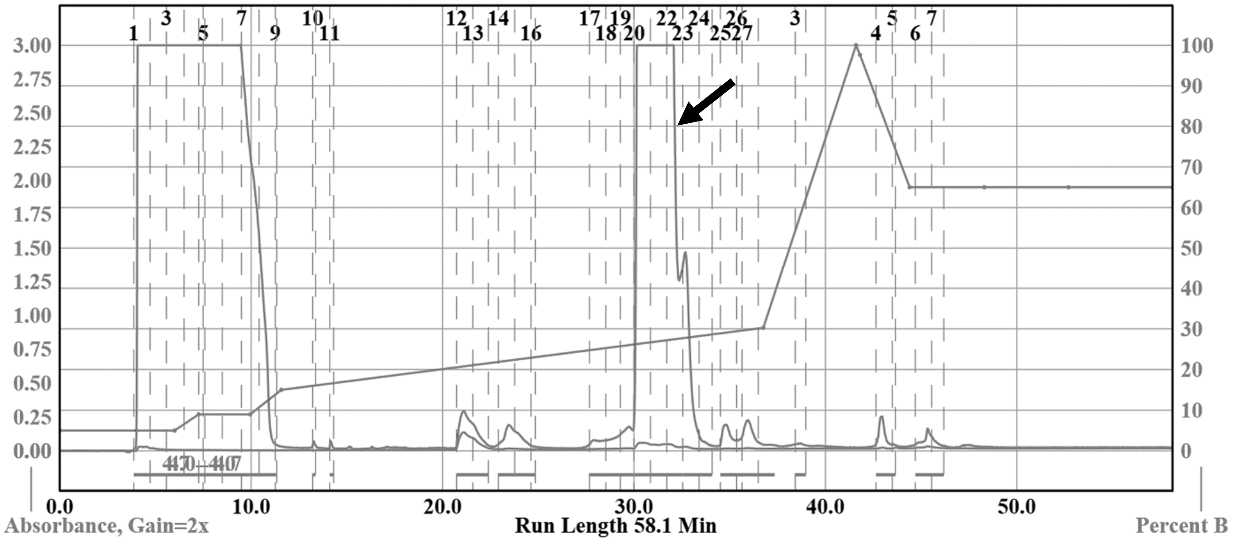

**Analytical LCMS**: Measured on an Agilent EC-C18 Poroshell column (4.6 x 100 mm, 4 µm, 100 Å, 40 °C, 1 mL/min) 5%-65% MeCN/$H_2O$/0.05% TFA over 60 min.

**Chemical Formula**: $C_{318}H_{512}N_{102}O_{104}Se_2$
**Molecular Weight**: 7586.1460 g/mol

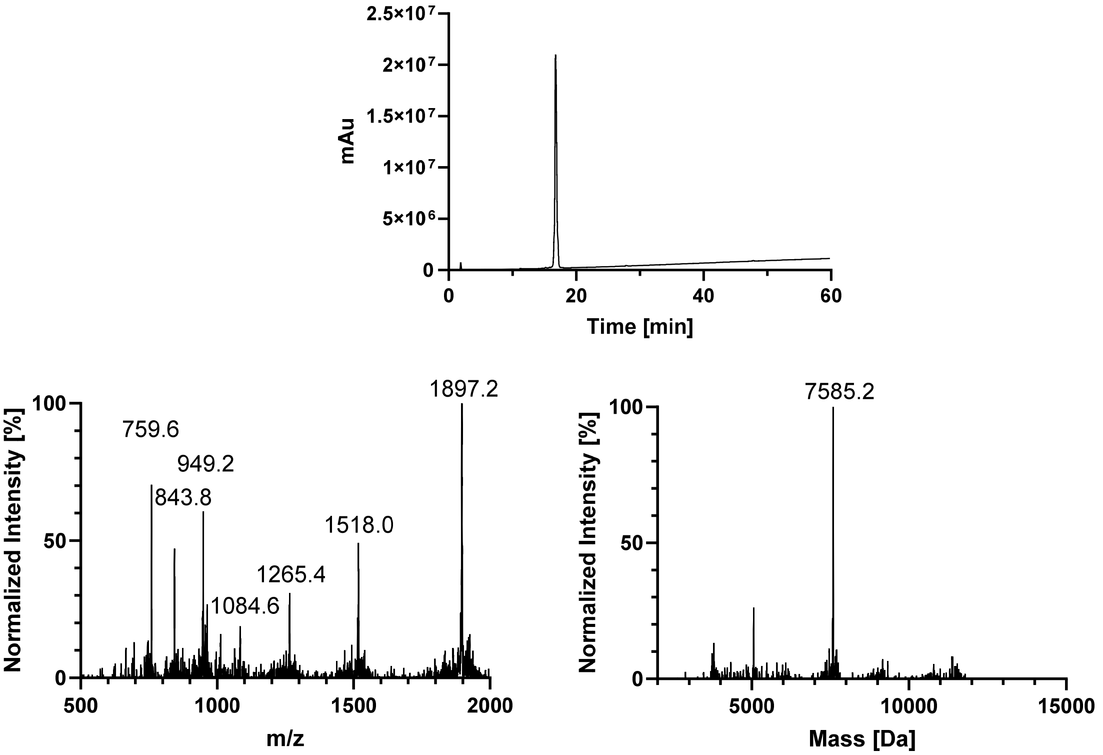
2.5×10⁷
2×10⁷
1.5×10⁷
1×10⁷
5×10⁶
0
mAu
0
20
40
60
Time [min]
Normalized Intensity [%]
100
50
0
759.6
949.2
843.8
1084.6
1265.4
1518.0
1897.2
500
1000
1500
2000
m/z
Normalized Intensity [%]
100
50
0
7585.2
5000
10000
15000
Mass [Da]

#### AcK369

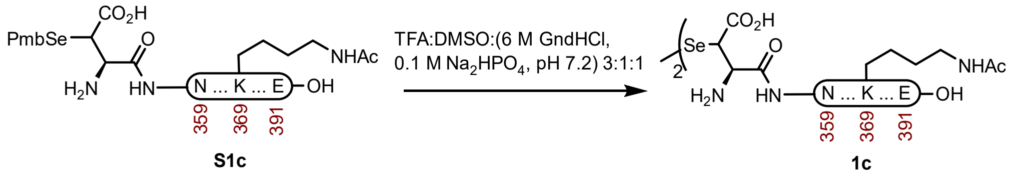

**Peptide 1c.** This diselenide synthesis method was modified from the literature.[1] A 2-dram vial containing **S1c** (53.0 mg, 13.3 µmol) was dissolved in buffer (400 µL, 6 M GndHCl, 0.1 M $Na_2HPO_4$, pH 7.2) and DMSO (400 µL). The mixture was cooled in an ice bath and treated with TFA (1.2 mL), wrapped with Al-Foil, and mixed on an overhead stirrer. The reaction was stirred at 23 ºC for 45 min before it was purified over a Luna C4 column (10 x 250 mm, 10 µm, 100 Å, 4.7 mL/min) 15%-35% MeCN/$H_2O$/0.1% TFA over 30 min. The fractions were lyophilized* to afford **1c** (47.0 mg, 92% yield).

* The fractions were lyophilized immediately and protected from light to prevent deselenization (reduction of diselenide into alanine).

**Prep HPLC trace of diselenide**

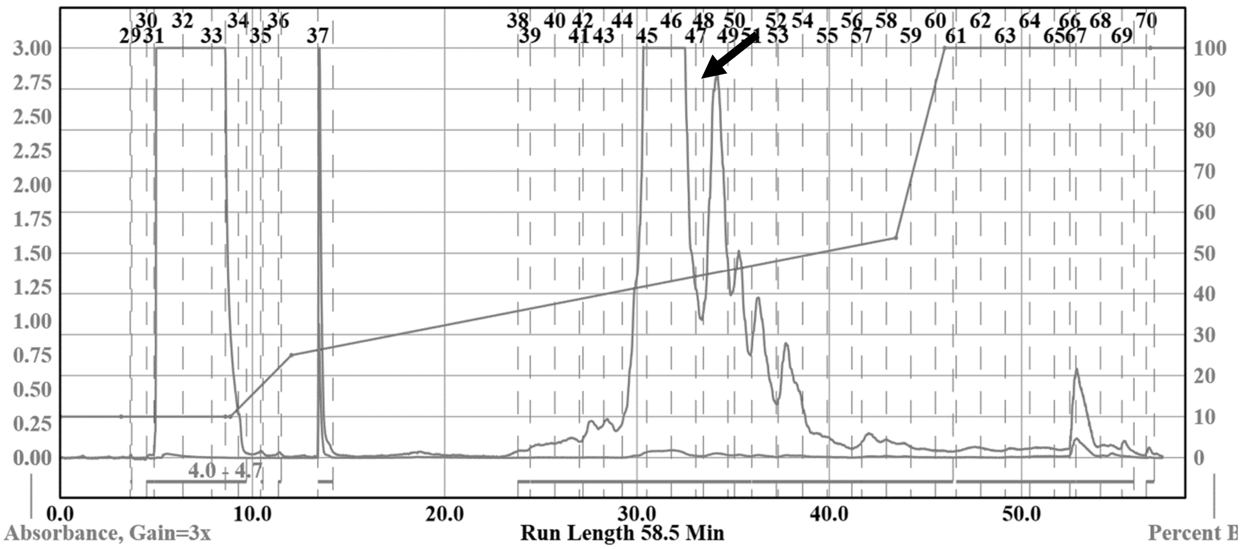

**Analytical LCMS**: Measured on an Agilent EC-C18 Poroshell column (4.6 x 100 mm, 4 µm, 100 Å, 40 °C, 1 mL/min) 5%-65% MeCN/$H_2O$/0.05% TFA over 60 min.

**Chemical Formula**: $C_{322}H_{516}N_{102}O_{106}Se_2$
**Molecular Weight**: 7670.2200 g/mol

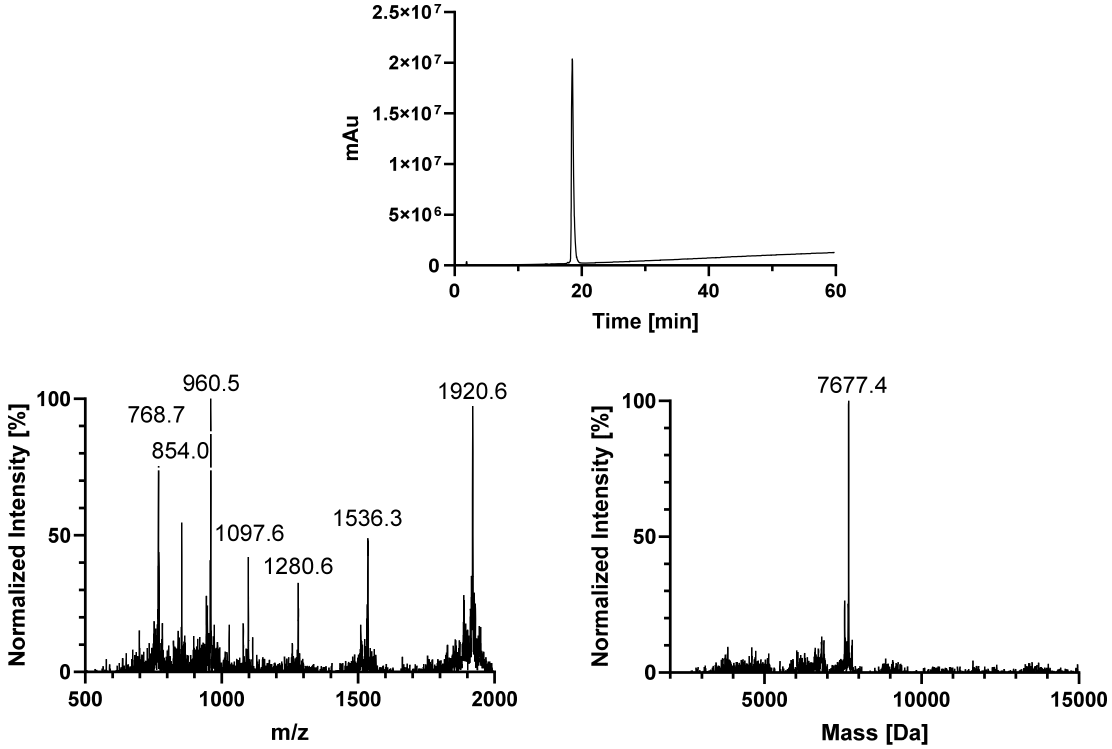
2.5×10⁷
2×10⁷
1.5×10⁷
1×10⁷
5×10⁶
0
mAu
0
20
40
60
Time [min]
960.5
768.7
854.0
1920.6
1097.6
1536.3
1280.6
Normalized Intensity [%]
100
50
0
500
1000
1500
2000
m/z
7677.4
Normalized Intensity [%]
100
50
0
5000
10000
15000
Mass [Da]

#### GlcNAcN359

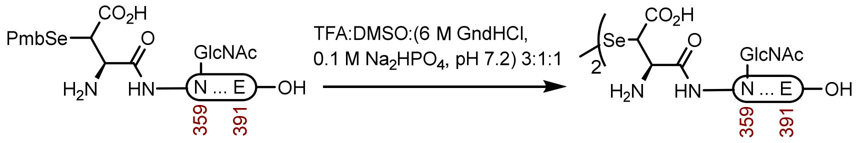

**S1k** **1k**

**Peptide 1k.** This diselenide synthesis method was modified from the literature.[1] A 2-dram vial containing **S1k** (24.9 mg, 6.05 µmol) was dissolved in buffer (300 µL, 6 M GndHCl, 0.1 M $Na_2HPO_4$, pH 7.2) and DMSO (300 µL). The mixture was cooled in an ice bath and treated with TFA (900 µL), wrapped with Al-Foil, and mixed on an overhead stirrer. The reaction was stirred at 23 ºC for 30 min before it was purified over a Luna C4 column (10 x 250 mm, 10 µm, 100 Å, 4.7 mL/min) 15%-35% MeCN/$H_2O$/0.1% TFA over 30 min. The fractions were lyophilized* to afford **1k** (19.0 mg, 79% yield).

* This diselenide was highly unstable in the HPLC buffer. The fractions that contain the peptides were lyophilized immediately, without checking LCMS and protected from light to prevent deselenization (reduction of diselenide into alanine). The LCMS data shows that the product is stable in the crude reaction mixture.

**Crude LCMS (Diselenide Synthesis)**: Measured on an Agilent EC-C18 Poroshell column (4.6 x 100 mm, 4 µm, 100 Å, 40 °C, 1 mL/min) 5%-65% MeCN/$H_2O$/0.05% TFA over 60 min.

**LC trace (214 nm) of unpurified peptide**

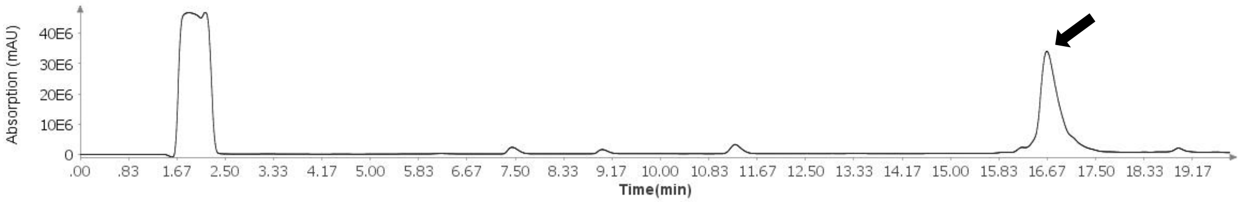

**Total Ion Current (TIC) of unpurified peptide**

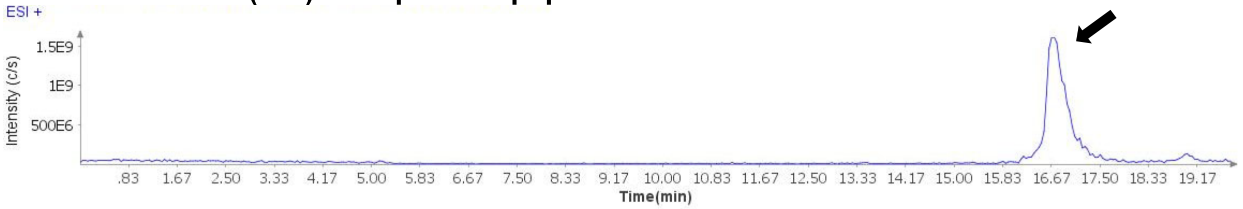

**Extracted Ion Current (XIC) of unpurified peptide**

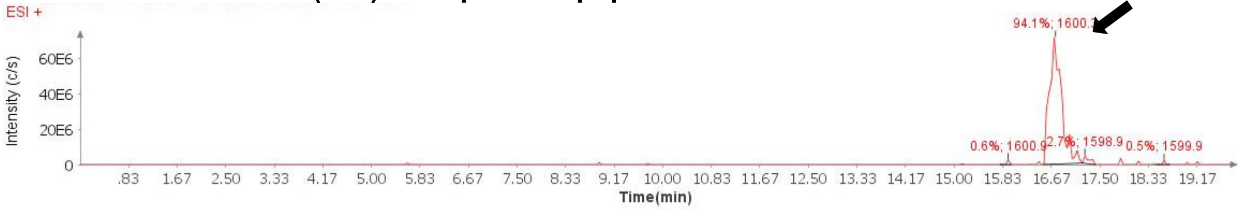

**Mass Spectrum (ESI) of the expected product (unpurified sample)**

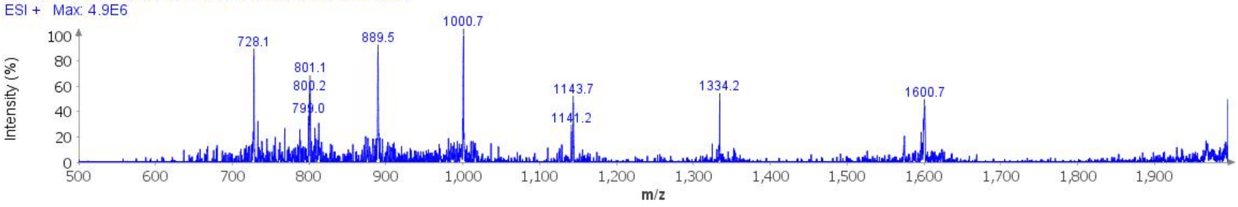

**Deconvoluted Mass Spectrum (ESI) of the expected product (unpurified sample)**

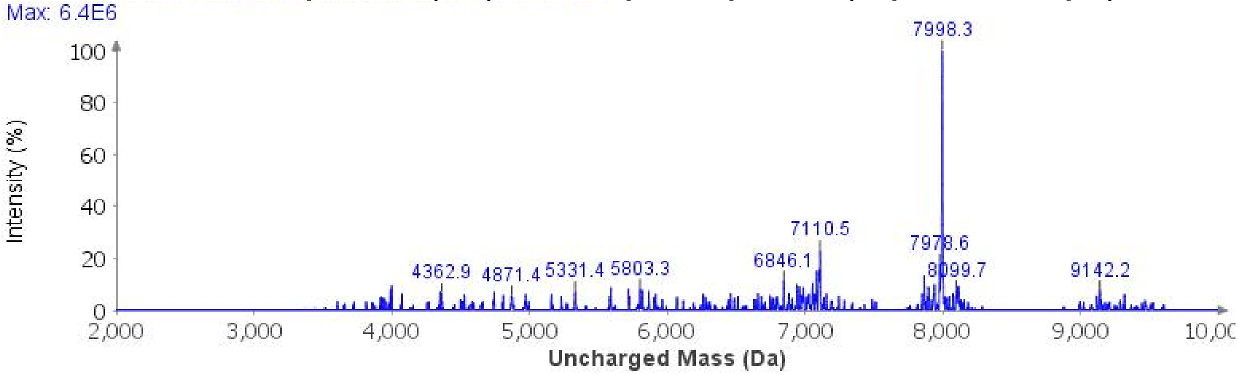

**Analytical LCMS**: Measured on an Agilent EC-C18 Poroshell column (4.6 x 100 mm, 4 µm, 100 Å, 40 °C, 1 mL/min) 5%-65% MeCN/$H_2O$/0.05% TFA over 20 min.

**Chemical Formula**: $C_{334}H_{538}N_{104}O_{114}Se_2$
**Molecular Weight**: 7992.5340 g/mol

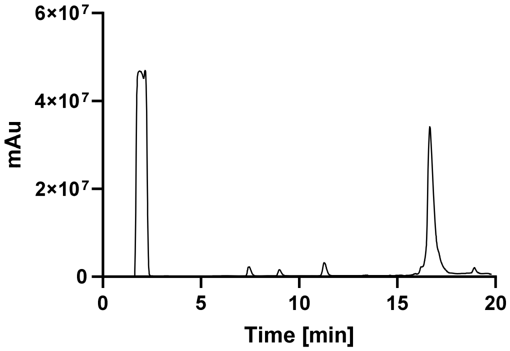

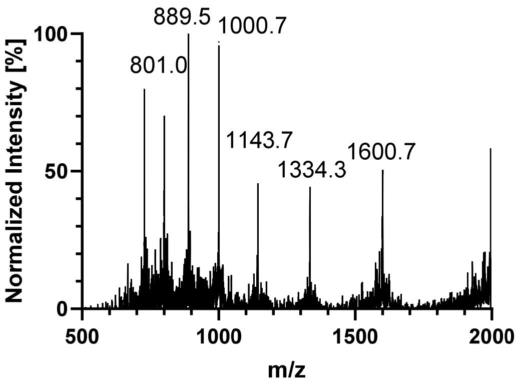

889.5
1000.7
801.0
1143.7
1334.3
1600.7
Normalized Intensity [%]
100
50
0
500
1000
1500
2000
m/z

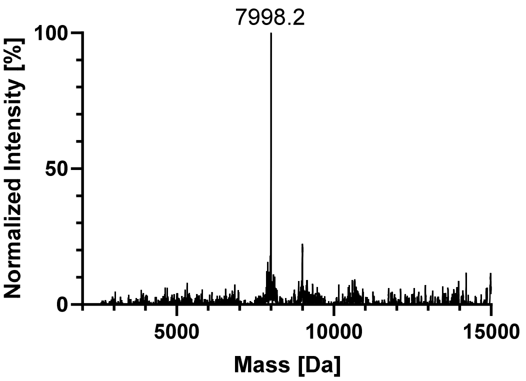

7998.2
Normalized Intensity [%]
100
50
0
5000
10000
15000
Mass [Da]

#### 6. C322-E391 (DSL)

### WT

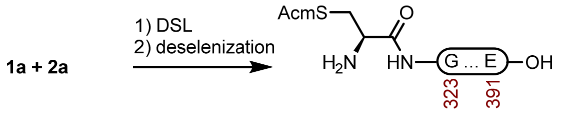

**4a**

**Peptide 4a.** This one pot DLS-deselenization method was modified from the literature.[1] A 2-dram vial containing **1a** (38.3 mg, 5.05 µmol) and **2a** (45.0 mg, 11.1 µmol) was dissolved in fresh DSL buffer (1.0 mL, 6 M GndHCl, 0.1 M $Na_2HPO_4$, pH 6.2) that had been sparged with $N_2$. The reaction was mixed at 23 ºC for 1 h before the residual DPDS was extracted with Hex (5 x 1 mL), and treated with hydrazine buffer (1.0 mL, 6 M GndHCl, 0.1 M $Na_2HPO_4$, 4% (v/v) hydrazine hydrate pH 7.4) that had been sparged with $N_2$. The reaction was stirred at 23 ºC for 10 min before it was treated with deselenization buffer (1.0 mL, 6 M GndHCl, 0.1 M $NaH_2PO_4$, 250 mM TCEP HCl, 25 mM DTT, pH 5.1). The reaction was stirred at 23 ºC for 10 min before it was purified over a Luna C4 column (10 x 250 mm, 10 µm, 100 Å, 4.7 mL/min) 15%-35% MeCN/$H_2O$/0.1% TFA over 30 min. The fractions were lyophilized to afford **4a** (54.2 mg, 70% yield).

**Crude LCMS (Deselenization)**: Measured on an Agilent EC-C18 Poroshell column (4.6 x 100 mm, 4 µm, 100 Å, 40 °C, 1 mL/min) 5%-65% MeCN/$H_2O$/0.05% TFA over 60 min.

**LC trace (214 nm) of unpurified peptide**

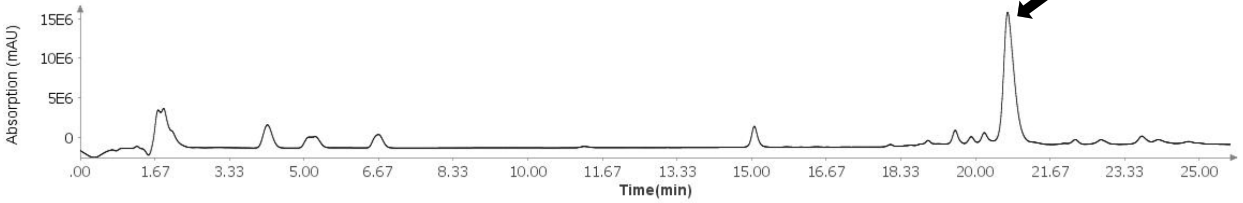

**Total Ion Current (TIC) of unpurified peptide**

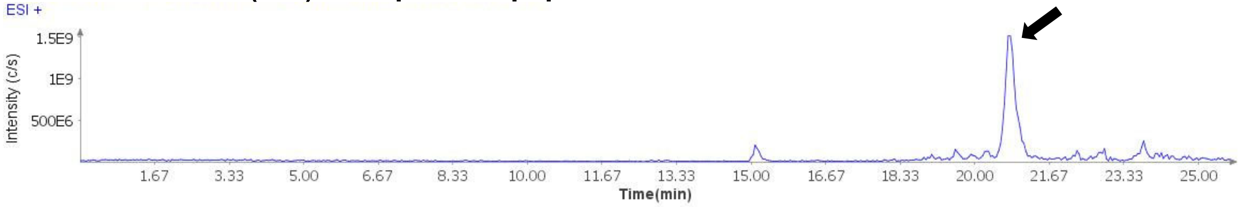

**Extracted Ion Current (XIC) of unpurified peptide**

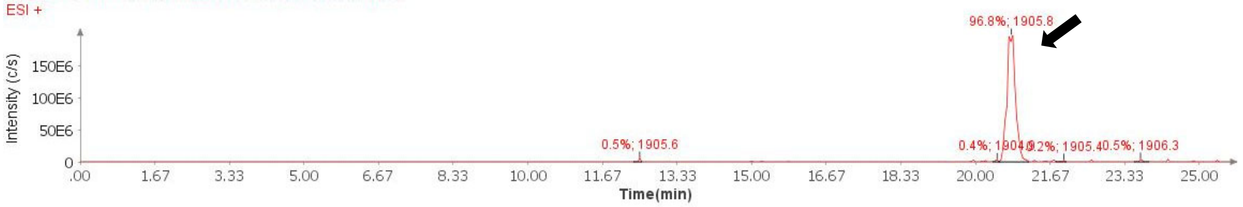

**Mass Spectrum (ESI) of the expected product (unpurified sample)**

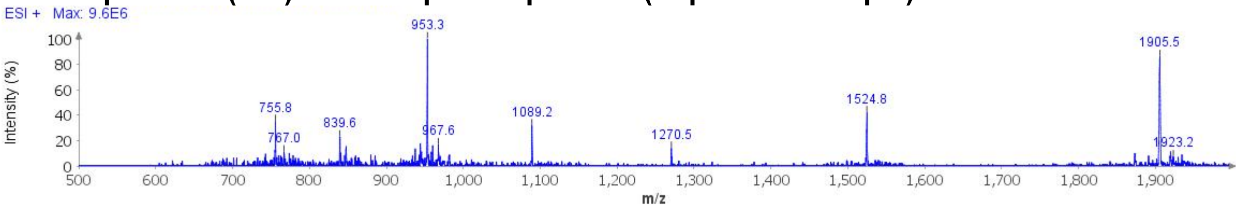

**Deconvoluted Mass Spectrum (ESI) of the expected product (unpurified sample)**

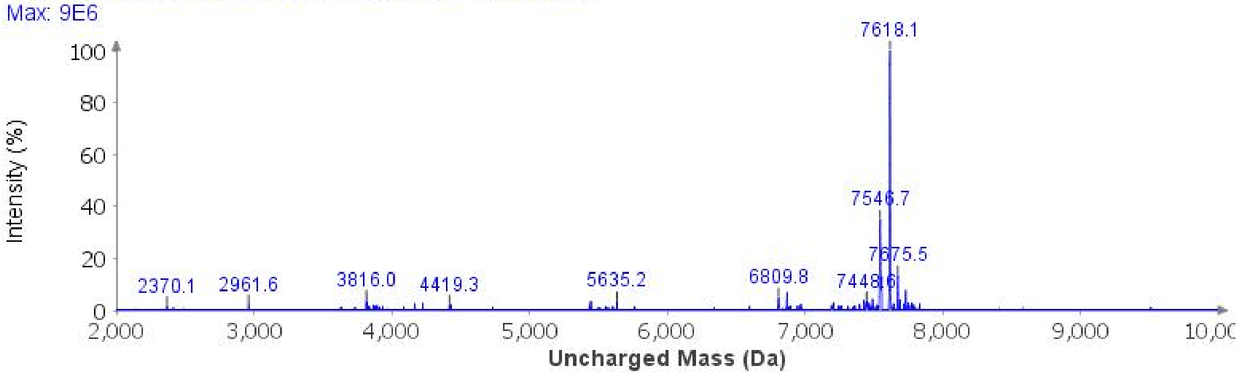

**Analytical LCMS**: Measured on an Agilent EC-C18 Poroshell column (4.6 x 100 mm, 4 µm, 100 Å, 40 °C, 1 mL/min) 5%-65% MeCN/$H_2O$/0.05% TFA over 60 min.

**Chemical Formula**: $C_{325}H_{530}N_{102}O_{103}S$
**Molecular Weight**: 7546.4860 g/mol

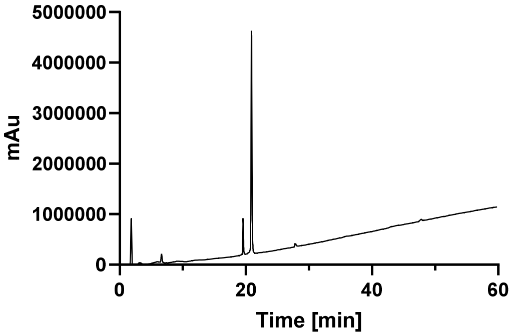

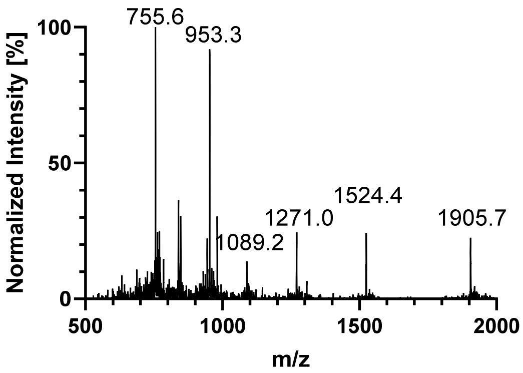
755.6
953.3
1089.2
1271.0
1524.4
1905.7
Normalized Intensity [%]
100
50
0
500
1000
1500
2000
m/z

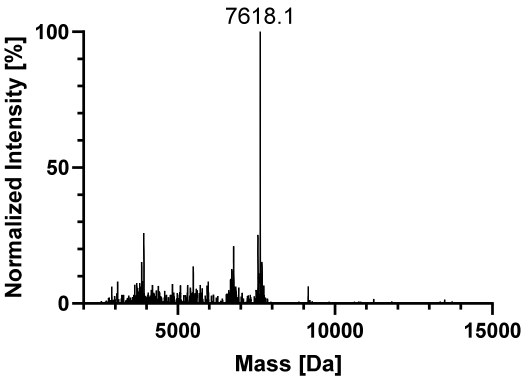
7618.1
Normalized Intensity [%]
100
50
0
5000
10000
15000
Mass [Da]

##### AcK369

1c + 2a

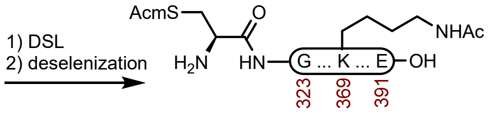

4c

**Peptide 4c.** This one pot DLS-deselenization method was modified from the literature.[1] A 1-dram vial containing **1c** (7.7 mg, 1.00 μmol) and **2a** (9.5 mg, 2.34 μmol) was dissolved in fresh DSL buffer (400 μL, 6 M GndHCl, 0.1 M $Na_2HPO_4$, pH 6.3) that had been sparged with $N_2$. The reaction was mixed at 23 ºC for 1 h before the residual DPDS was extracted with Hex (5 x 400 μL), and treated with hydrazine buffer (400 μL, 6 M GndHCl, 0.1 M $Na_2HPO_4$, 4% (v/v) hydrazine hydrate pH 7.4) that had been sparged with $N_2$. The reaction was stirred at 23 ºC for 15 min before it was treated with deselenization buffer (400 μL, 6 M GndHCl, 0.1 M $NaH_2PO_4$, 250 mM TCEP HCl, 25 mM DTT, pH 5.1). The reaction was stirred at 23 ºC for 5 min before it was purified over a Luna C4 column (10 x 250 mm, 10 μm, 100 Å, 4.7 mL/min) 15%-35% MeCN/$H_2O$/0.1% TFA over 30 min. The fractions were lyophilized to afford **4c** (14.2 mg, 93% yield).

**Crude LCMS (Deselenization)**: Measured on an Agilent EC-C18 Poroshell column (4.6 x 100 mm, 4 μm, 100 Å, 40 °C, 1 mL/min) 5%-65% MeCN/$H_2O$/0.05% TFA over 60 min.

**LC trace (214 nm) of unpurified peptide**

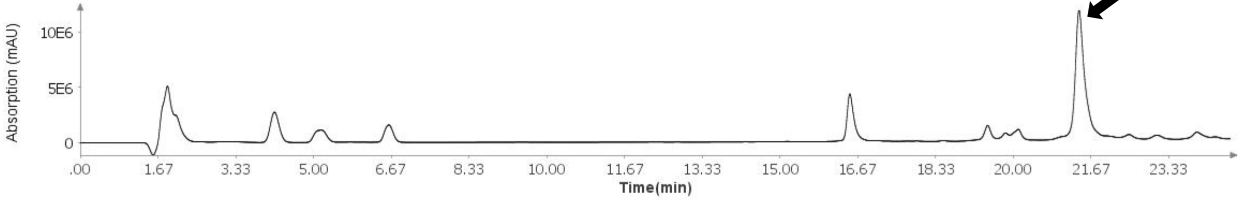

**Total Ion Current (TIC) of unpurified peptide**

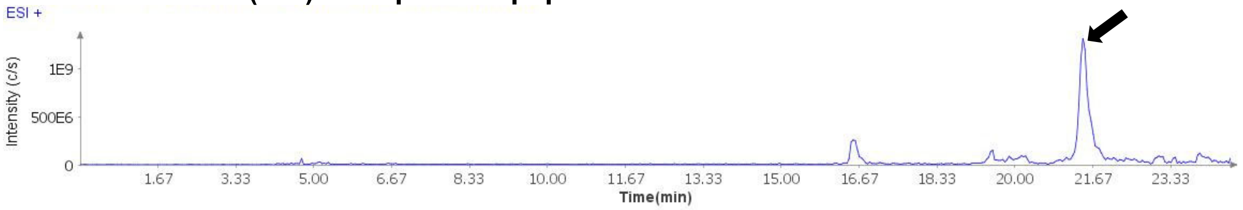

**Extracted Ion Current (XIC) of unpurified peptide**

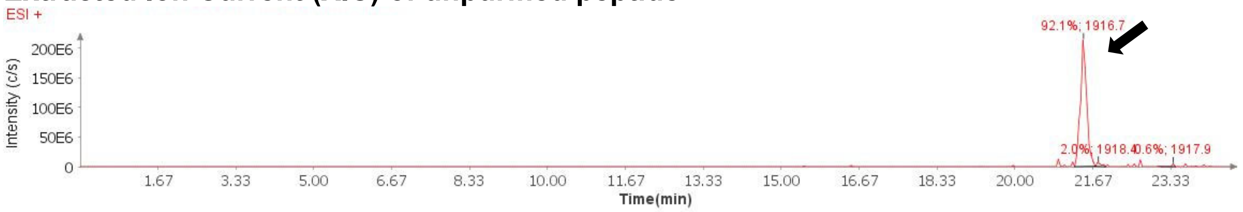

**Mass Spectrum (ESI) of the expected product (unpurified sample)**

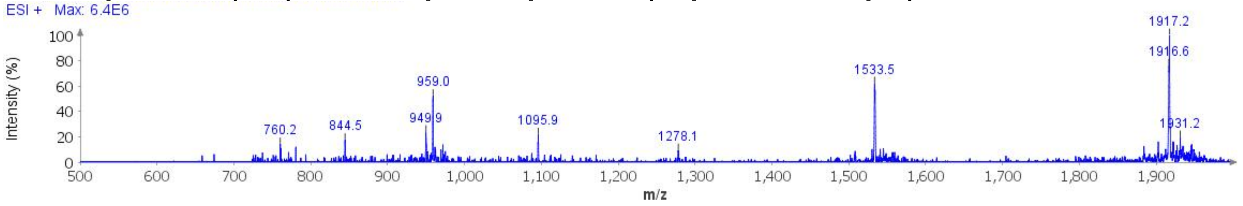

**Deconvoluted Mass Spectrum (ESI) of the expected product (unpurified sample)**

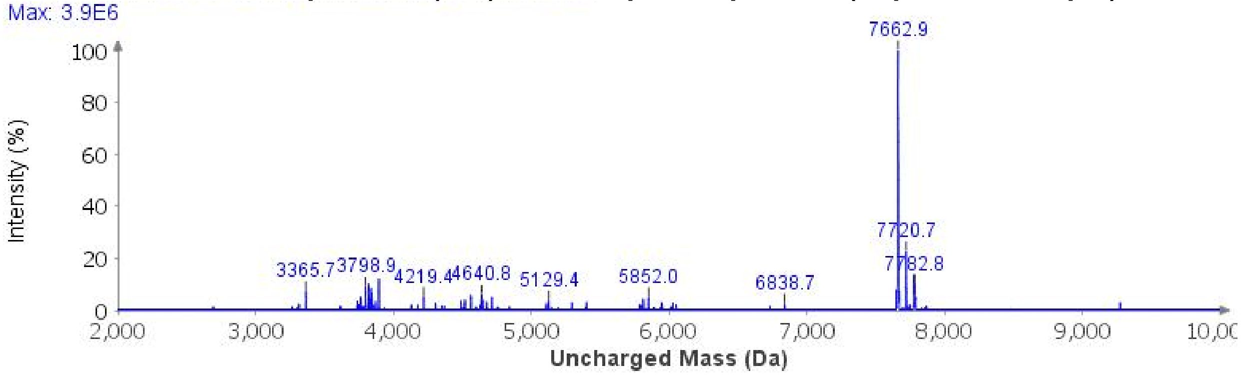

**Analytical LCMS**: Measured on an Agilent EC-C18 Poroshell column (4.6 x 100 mm, 4 µm, 100 Å, 40 °C, 1 mL/min) 5%-65% MeCN/$H_2O$/0.05% TFA over 60 min.

**Chemical Formula**: $C_{330}H_{537}N_{103}O_{105}S$
**Molecular Weight**: 7659.6020 g/mol

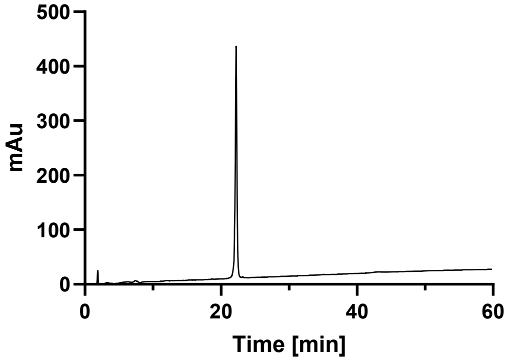

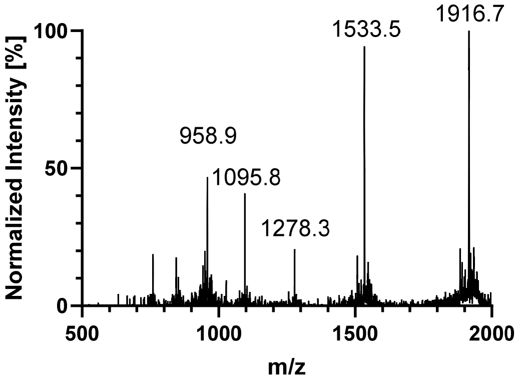
Normalized Intensity [%]
100
50
0
500
1000
1500
2000
m/z
958.9
1095.8
1278.3
1533.5
1916.7

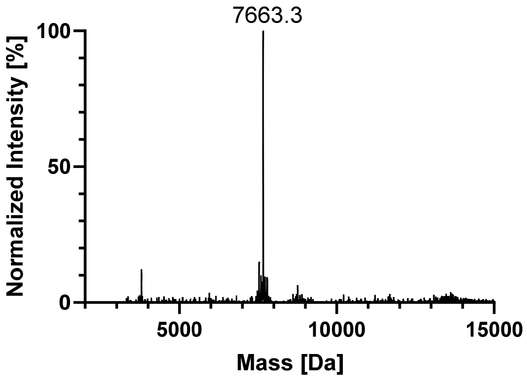
Normalized Intensity [%]
100
50
0
5000
10000
15000
Mass [Da]
7663.3

#### AcK353

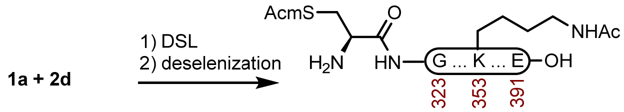

**4d**

**Peptide 4d.** This one pot DLS-deselenization method was modified from the literature.[1] A 1-dram vial containing **1a** (9.1 mg, 1.20 µmol) and **2d** (13.7 mg, 3.34 µmol) was dissolved in fresh DSL buffer (500 µL, 6 M GndHCl, 0.1 M $Na_2HPO_4$, pH 6.1) that had been sparged with $N_2$. The reaction was mixed at 23 ºC for 1 h before the residual DPDS was extracted with Hex (5 x 500 µL), and treated with hydrazine buffer (500 µL, 6 M GndHCl, 0.1 M $Na_2HPO_4$, 4% (v/v) hydrazine hydrate pH 7.4) that had been sparged with $N_2$. The reaction was stirred at 23 ºC for 15 min before it was treated with deselenization buffer (500 µL, 6 M GndHCl, 0.1 M $NaH_2PO_4$, 250 mM TCEP HCl, 25 mM DTT, pH 5.2). The reaction was stirred at 23 ºC for 5 min before it was purified over an Agilent C18 column (10 x 250 mm, 5 µm, 100 Å, 4.7 mL/min) 15%-35% MeCN/$H_2O$/0.1% TFA over 30 min. The fractions were lyophilized to afford **4a** (18.0 mg, 97% yield).

**Crude LCMS (Deselenization)**: Measured on an Agilent EC-C18 Poroshell column (4.6 x 100 mm, 4 µm, 100 Å, 40 °C, 1 mL/min) 5%-65% MeCN/$H_2O$/0.05% TFA over 60 min.

##### LC trace (214 nm) of unpurified peptide

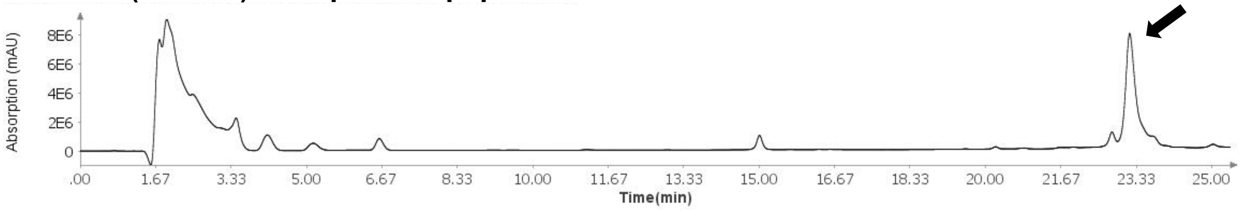

##### Total Ion Current (TIC) of unpurified peptide

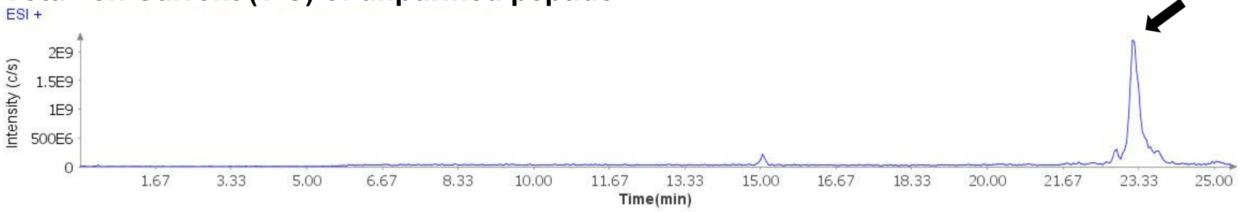

##### Extracted Ion Current (XIC) of unpurified peptide

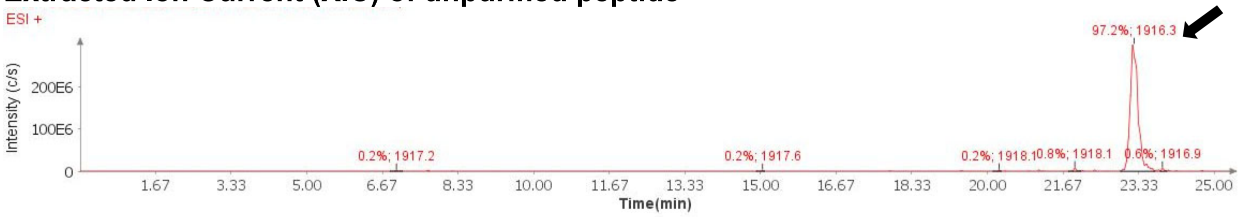

**Mass Spectrum (ESI) of the expected product (unpurified sample)**

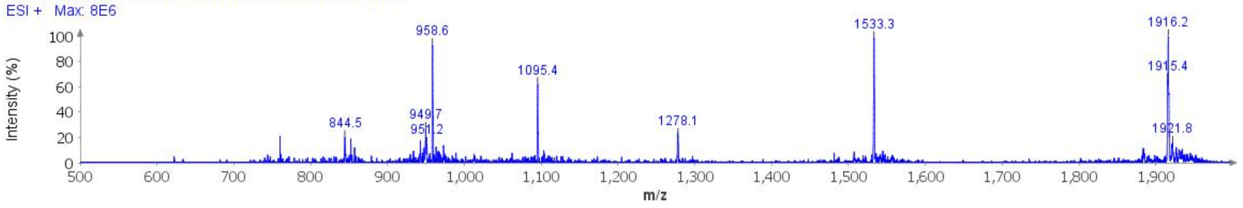

**Deconvoluted Mass Spectrum (ESI) of the expected product (unpurified sample)**

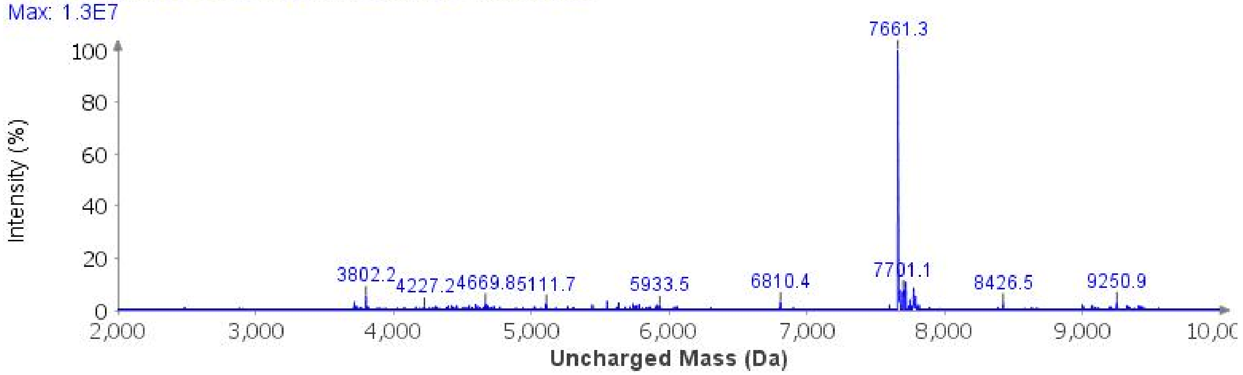

**Analytical LCMS**: Measured on an Agilent EC-C18 Poroshell column (4.6 x 100 mm, 4 µm, 100 Å, 40 °C, 1 mL/min) 5%-65% MeCN/$H_2O$/0.05% TFA over 60 min.

**Chemical Formula**: $C_{330}H_{537}N_{103}O_{105}S$
**Molecular Weight**: 7659.6020 g/mol

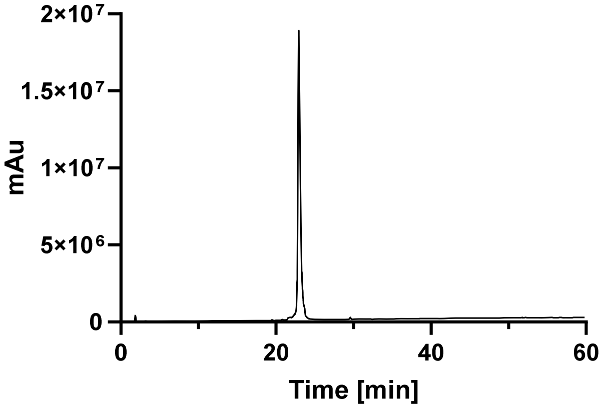

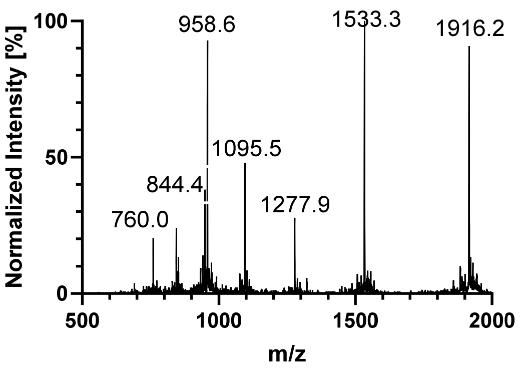
Normalized Intensity [%]
100
50
0
958.6
1533.3
1916.2
1095.5
844.4
1277.9
760.0
500
1000
1500
2000
m/z

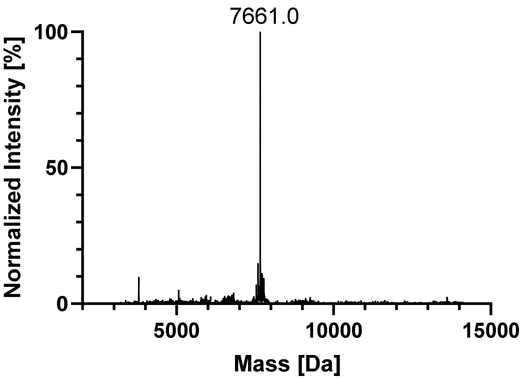
Normalized Intensity [%]
100
50
0
7661.0
5000
10000
15000
Mass [Da]

## pS356

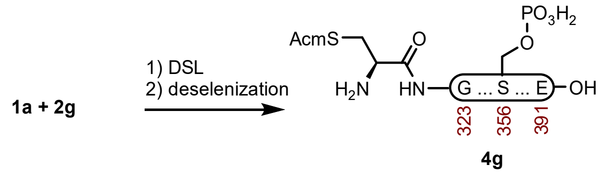

**Peptide 4g.** This one pot DLS-deselenization method was modified from the literature.[1] A 1-dram vial containing **1a** (20.7 mg, 2.73 μmol) and **2g** (32.0 mg, 7.73 μmol) was dissolved in fresh DSL buffer (400 μL, 6 M GndHCl, 0.1 M $Na_2HPO_4$, pH 6.1) that had been sparged with $N_2$. The reaction was mixed at 23 ºC for 1 h before the residual DPDS was extracted with Hex (5 x 400 μL), and treated with hydrazine buffer (500 μL, 6 M GndHCl, 0.1 M $Na_2HPO_4$, 4% (v/v) hydrazine hydrate pH 7.5) that had been sparged with $N_2$. The reaction was stirred at 23 ºC for 15 min before it was treated with deselenization buffer (400 μL, 6 M GndHCl, 0.1 M $NaH_2PO_4$, 250 mM TCEP HCl, 25 mM DTT, pH 5.2). The reaction was stirred at 23 ºC for 15 min before it was purified over a Luna PhenylHexyl column (10 x 250 mm, 5 μm, 100 Å, 4.7 mL/min) 15%-40% MeCN/$H_2O$/0.1% TFA over 30 min. The fractions were lyophilized to afford **4g** (16.3 mg, 39% yield).

**Prep HPLC trace of diselenide-selenoester-ligation-deselenization**

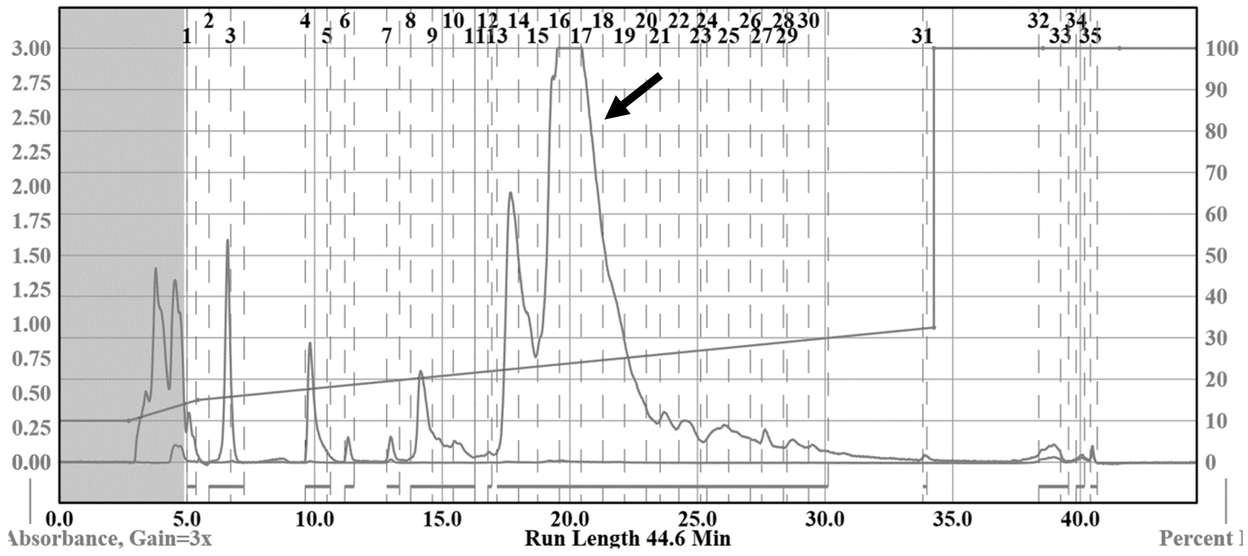

**Analytical LCMS**: Measured on an Agilent EC-C18 Poroshell column (4.6 x 100 mm, 4 μm, 100 Å, 40 °C, 1 mL/min) 5%-65% MeCN/$H_2O$/0.05% TFA over 60 min.

**Chemical Formula**: $C_{328}H_{536}N_{103}O_{107}PS$
**Molecular Weight**: 7697.5438 g/mol

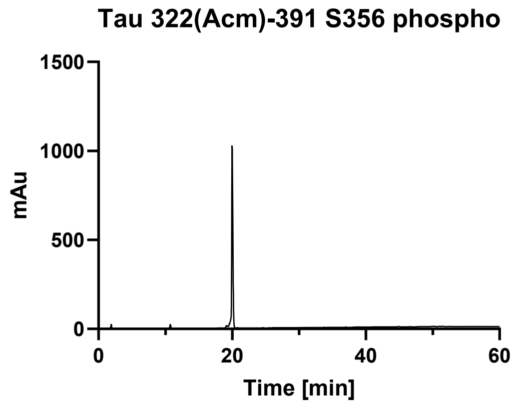
Tau 322(Acm)-391 S356 phospho
1500
1000
500
0
mAu
0
20
40
60
Time [min]

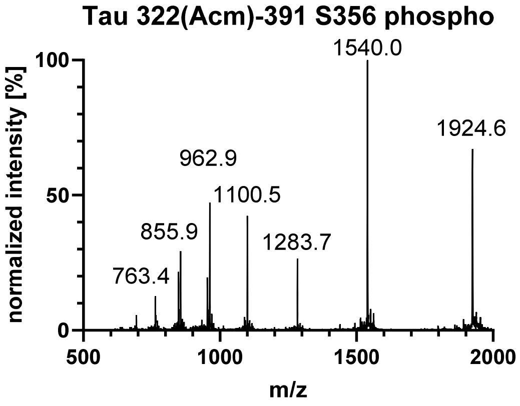
Tau 322(Acm)-391 S356 phospho
100
50
0
normalized intensity [%]
1540.0
1924.6
962.9
1100.5
855.9
1283.7
763.4
500
1000
1500
2000
m/z

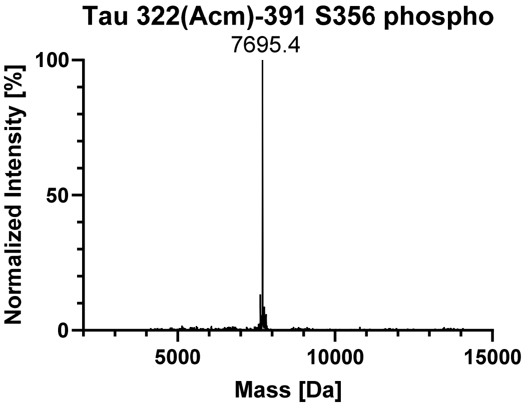
Tau 322(Acm)-391 S356 phospho
100
50
0
Normalized Intensity [%]
7695.4
5000
10000
15000
Mass [Da]

## pS352

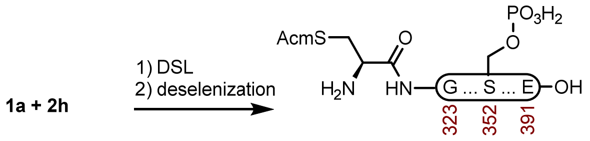

**4h**

**Peptide 4h.** This one pot DLS-deselenization method was modified from the literature.[1] A 1-dram vial containing **1a** (20.4 mg, 2.69 µmol) and **2h** (29.0 mg, 7.01 µmol) was dissolved in fresh DSL buffer (400 µL, 6 M GndHCl, 0.1 M $Na_2HPO_4$, pH 6.1) that had been sparged with $N_2$. The reaction was mixed at 23 ºC for 1 h before the residual DPDS was extracted with Hex (5 x 400 µL), and treated with hydrazine buffer (500 µL, 6 M GndHCl, 0.1 M $Na_2HPO_4$, 4% (v/v) hydrazine hydrate pH 7.5) that had been sparged with $N_2$. The reaction was stirred at 23 ºC for 15 min before it was treated with deselenization buffer (400 µL, 6 M GndHCl, 0.1 M $NaH_2PO_4$, 250 mM TCEP HCl, 25 mM DTT, pH 5.2). The reaction was stirred at 23 ºC for 15 min before it was purified over a Luna PhenylHexyl column (10 x 250 mm, 5 µm, 100 Å, 4.7 mL/min) 15%-40% MeCN/$H_2O$/0.1% TFA over 30 min. The fractions were lyophilized to afford **4h** (14.1 mg, 34% yield).

**Crude LCMS (Deselenization)**: Measured on an Agilent EC-C18 Poroshell column (4.6 x 100 mm, 4 µm, 100 Å, 40 °C, 1 mL/min) 5%-65% MeCN/$H_2O$/0.05% TFA over 60 min.

**LC trace (214 nm) of unpurified peptide**

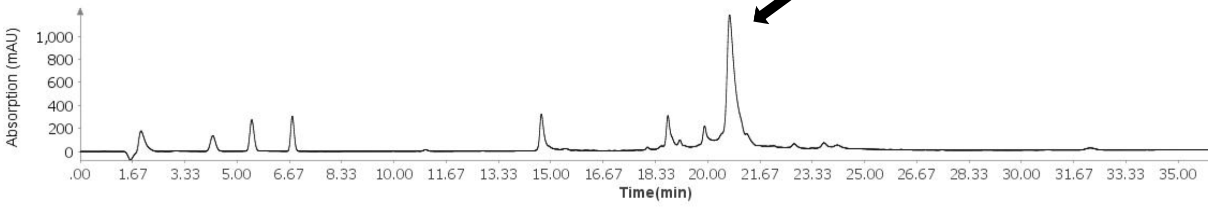

**Total Ion Current (TIC) of unpurified peptide**

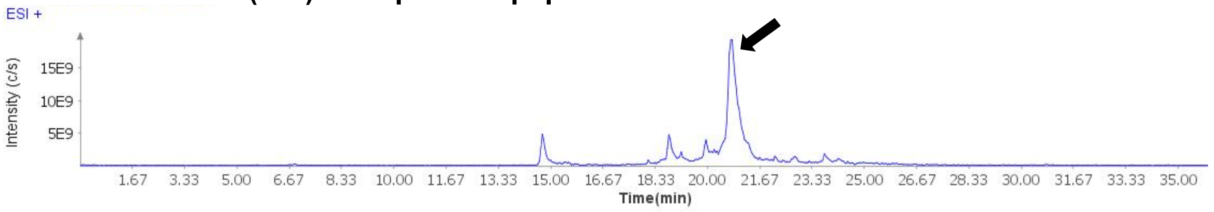

**Extracted Ion Current (XIC) of unpurified peptide**

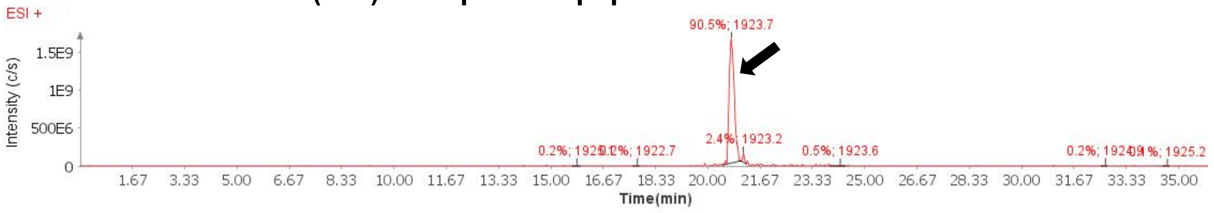

**Mass Spectrum (ESI) of the expected product (unpurified sample)**

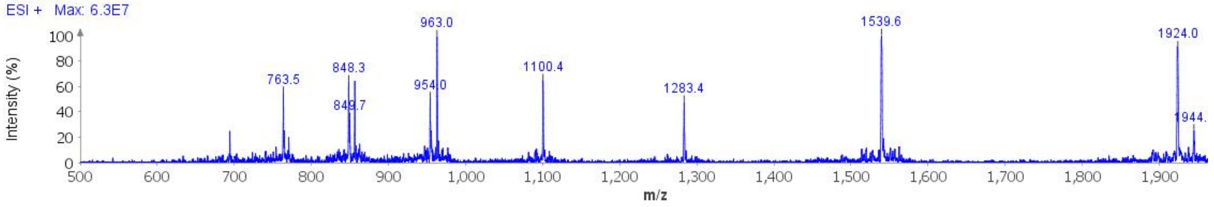

**Deconvoluted Mass Spectrum (ESI) of the expected product (unpurified sample)**

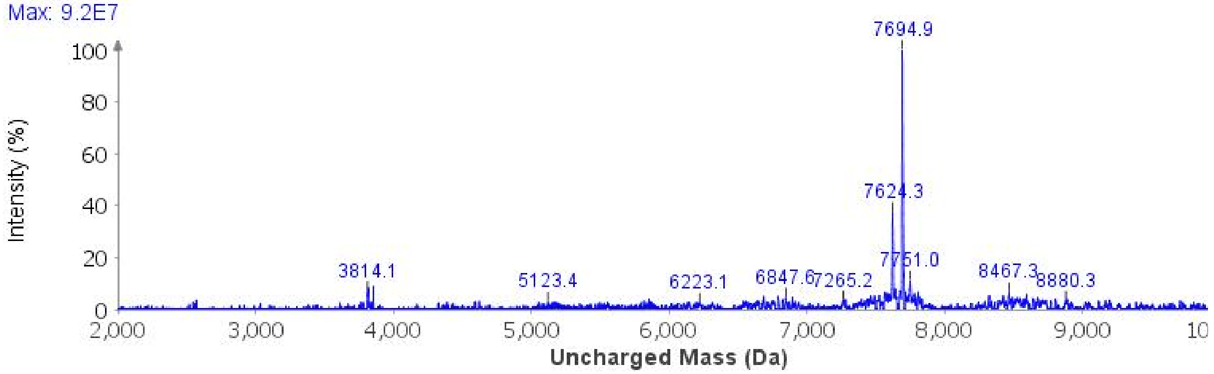

**Analytical LCMS**: Measured on an Agilent EC-C18 Poroshell column (4.6 x 100 mm, 4 µm, 100 Å, 40 °C, 1 mL/min) 5%-85% MeCN/$H_2O$/0.05% TFA over 30 min.

**Chemical Formula**: $C_{328}H_{536}N_{103}O_{107}PS$
**Molecular Weight**: 7697.5438 g/mol

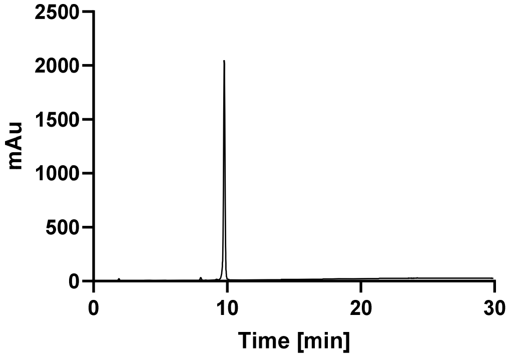

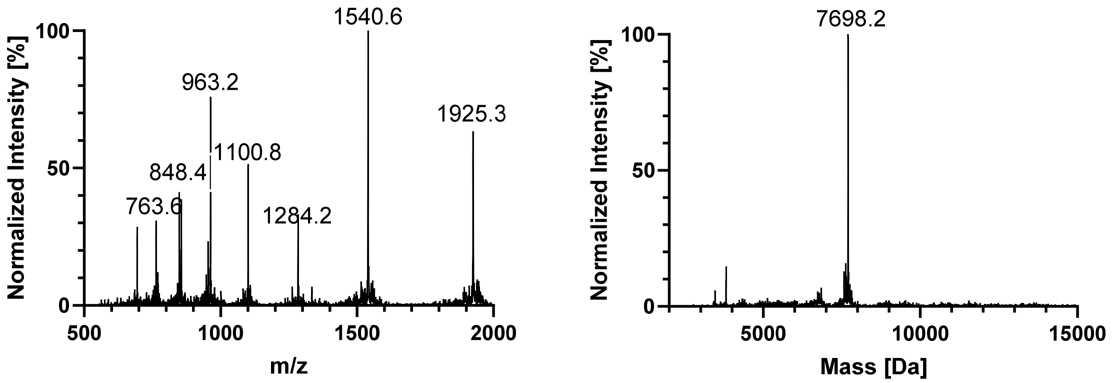

Normalized Intensity [%]
100
50
0
500
1000
1500
2000
m/z
1540.6
963.2
1925.3
1100.8
848.4
763.6
1284.2
Normalized Intensity [%]
100
50
0
5000
10000
15000
Mass [Da]
7698.2

#### GlcNAcN359

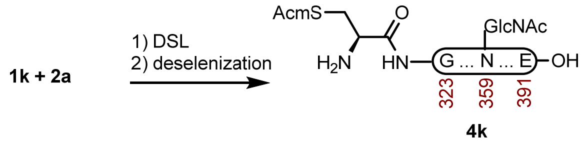

**Peptide 4k.** This one pot DLS-deselenization method was modified from the literature.[1] A 2-dram vial containing **1k** (12.5 mg, 1.56 µmol) and **2a** (15.5 mg, 3.82 µmol) was dissolved in fresh DSL buffer (500 µL, 6 M GndHCl, 0.1 M $Na_2HPO_4$, pH 6.2) that had been sparged with $N_2$. The reaction was mixed at 23 ºC for 1 h before the residual DPDS was extracted with Hex (5 x 500 µL), and treated with hydrazine buffer (500 µL, 6 M GndHCl, 0.1 M $Na_2HPO_4$, 4% (v/v) hydrazine hydrate pH 7.3) that had been sparged with $N_2$. The reaction was stirred at 23 ºC for 30 min before it was treated with deselenization buffer (500 µL, 6 M GndHCl, 0.1 M $NaH_2PO_4$, 250 mM TCEP HCl, 25 mM DTT, pH 5.2). The reaction was stirred at 23 ºC for 15 min before it was purified over a Luna C4 column (10 x 250 mm, 10 µm, 100 Å, 4.7 mL/min) 15%-35% MeCN/$H_2O$/0.1% TFA over 30 min. The fractions were lyophilized to afford **4k** (20.2 mg, 83% yield).

**Crude LCMS (Deselenization)**: Measured on an Agilent EC-C18 Poroshell column (4.6 x 100 mm, 4 µm, 100 Å, 40 °C, 1 mL/min) 5%-65% MeCN/$H_2O$/0.05% TFA over 60 min.

##### LC trace (214 nm) of unpurified peptide

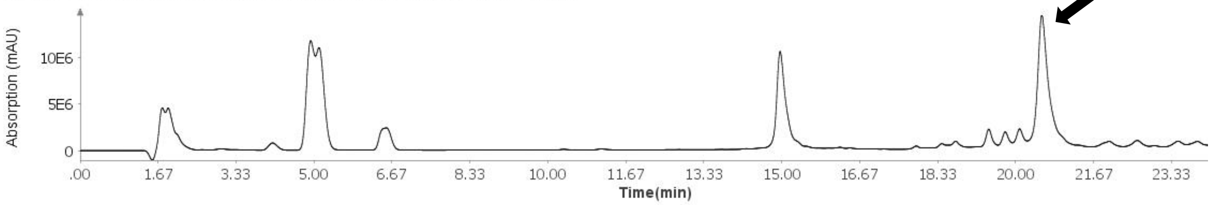

##### Total Ion Current (TIC) of unpurified peptide

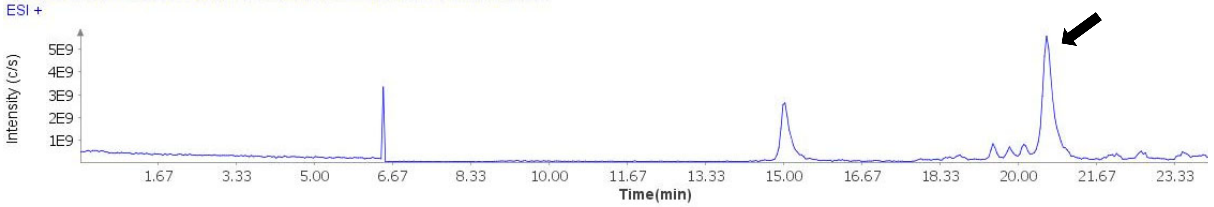

##### Extracted Ion Current (XIC) of unpurified peptide

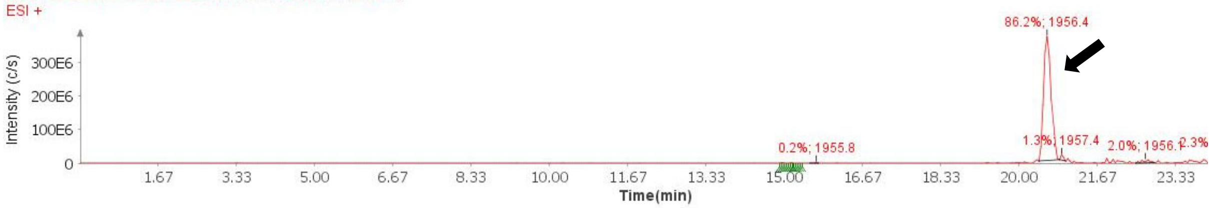

**Mass Spectrum (ESI) of the expected product (unpurified sample)**

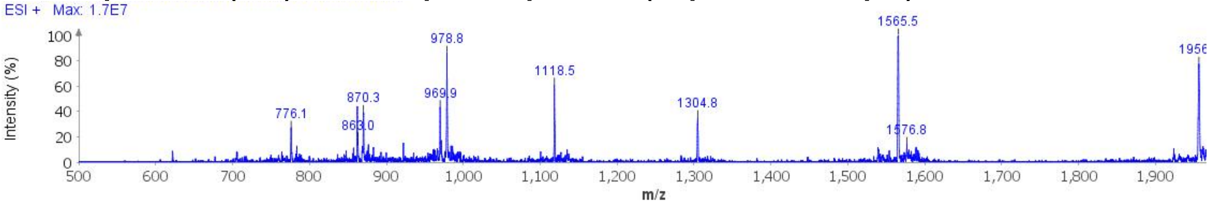

**Deconvoluted Mass Spectrum (ESI) of the expected product (unpurified sample)**

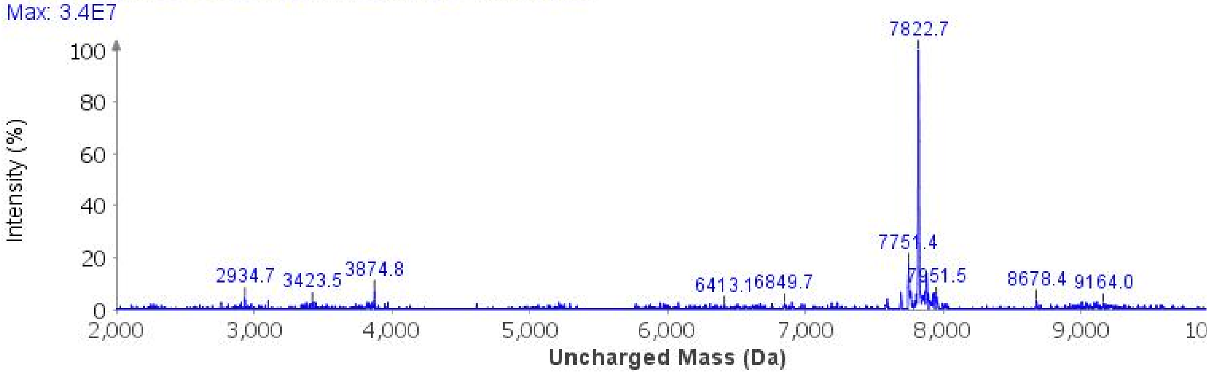

**Analytical LCMS**: Measured on an Agilent EC-C18 Poroshell column (4.6 x 100 mm, 4 µm, 100 Å, 40 °C, 1 mL/min) 5%-65% MeCN/$H_2O$/0.05% TFA over 60 min.

**Chemical Formula**: $C_{336}H_{548}N_{104}O_{109}S$
**Molecular Weight**: 7820.7590 g/mol

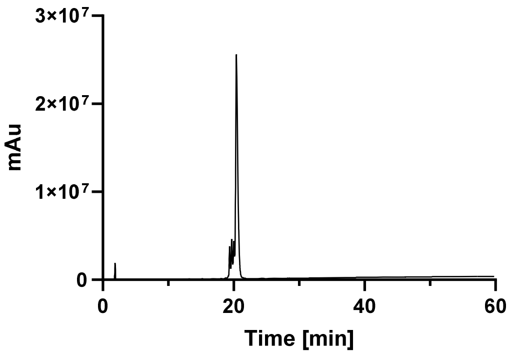

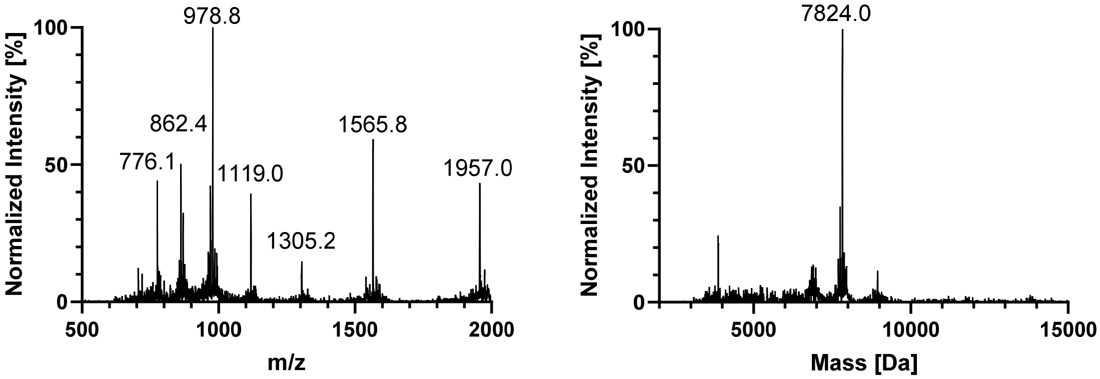
978.8
862.4
776.1
1119.0
1305.2
1565.8
1957.0
Normalized Intensity [%]
100
50
0
500
1000
1500
2000
m/z
7824.0
Normalized Intensity [%]
100
50
0
5000
10000
15000
Mass [Da]

#### GlcNAcN359 + AcK353

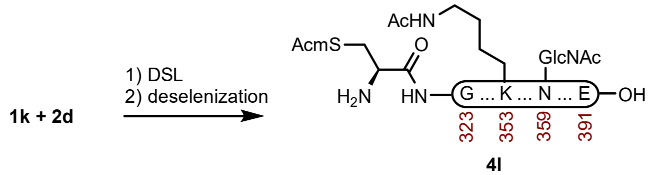

**Peptide 4l.** This one pot DLS-deselenization method was modified from the literature.[1] A 2-dram vial containing **1k** (6.3 mg,0.788 µmol) and **2d** (8.5 mg, 2.07 µmol) was dissolved in fresh DSL buffer (300 µL, 6 M GndHCl, 0.1 M $Na_2HPO_4$, pH 6.2) that had been sparged with $N_2$. The reaction was mixed at 23 ºC for 1 h before the residual DPDS was extracted with Hex (5 x 300 µL), and treated with hydrazine buffer (300 µL, 6 M GndHCl, 0.1 M $Na_2HPO_4$, 4% (v/v) hydrazine hydrate pH 7.3) that had been sparged with $N_2$. The reaction was stirred at 23 ºC for 30 min before it was treated with deselenization buffer (300 µL, 6 M GndHCl, 0.1 M $NaH_2PO_4$, 250 mM TCEP HCl, 25 mM DTT, pH 5.3). The reaction was stirred at 23 ºC for 15 min before it was purified over a Luna C4 column (10 x 250 mm, 10 µm, 100 Å, 4.7 mL/min) 15%-35% MeCN/$H_2O$/0.1% TFA over 30 min. The fractions were lyophilized to afford **4l** (10.0 mg, 81% yield).

**Prep HPLC trace of diselenide-selenoester-ligation-deselenization**

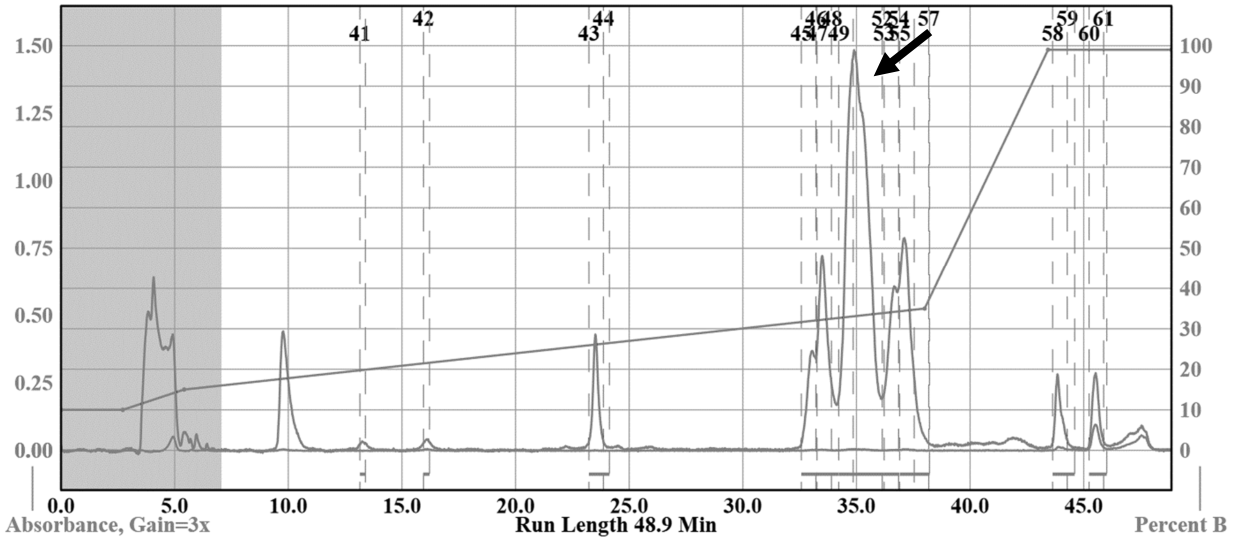

**Analytical LCMS**: Measured on an Agilent EC-C18 Poroshell column (4.6 x 100 mm, 4 µm, 100 Å, 40 °C, 1 mL/min) 5%-65% MeCN/$H_2O$/0.05% TFA over 60 min.

**Chemical Formula**: $C_{338}H_{550}N_{104}O_{110}S$
**Molecular Weight**: 7862.7960 g/mol

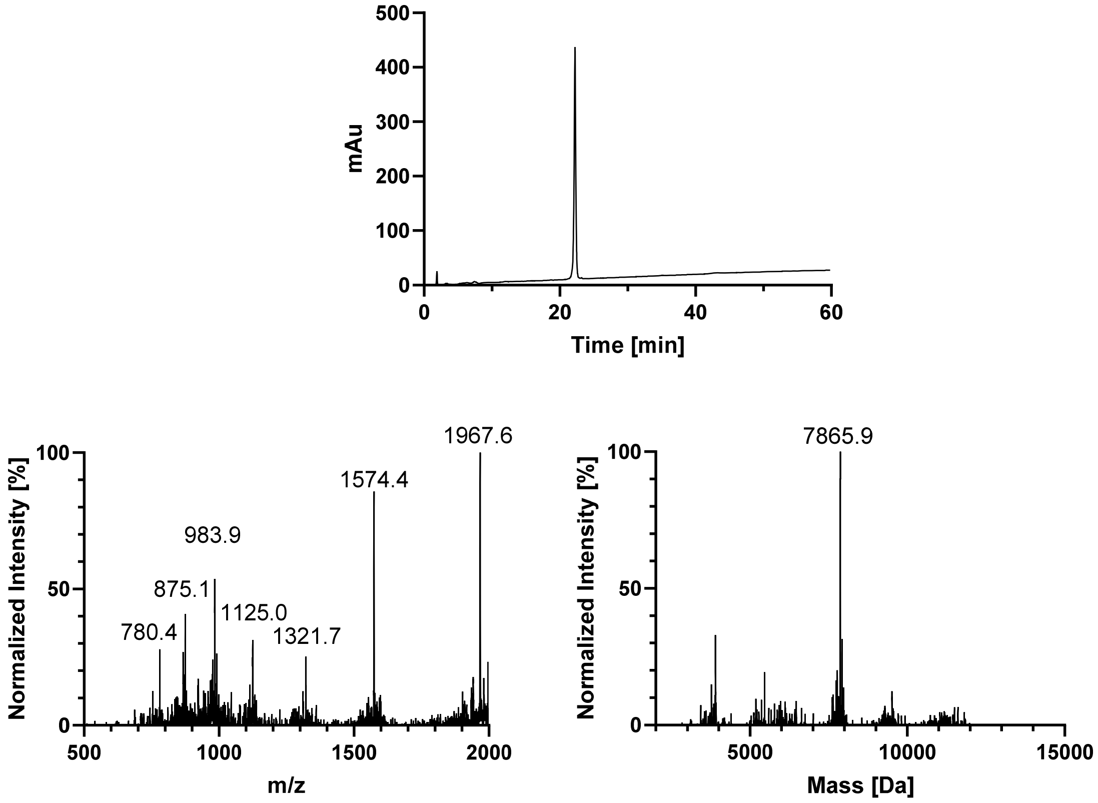

500
400
300
200
100
0
mAu
0
20
40
60
Time [min]
1967.6
1574.4
983.9
875.1
1125.0
780.4
1321.7
Normalized Intensity [%]
100
50
0
500
1000
1500
2000
m/z
7865.9
Normalized Intensity [%]
100
50
0
5000
10000
15000
Mass [Da]

#### 7. C322-E391 (Acm Removal)

### WT

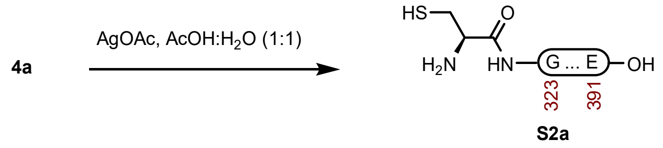

**Peptide S2a.** This Acm removal method was modified from the literature.[4] A 2 dram vial with **4a** (17.0 mg, 2.23 µmol) and AgOAc (7.4 mg, 47.3 µmol) was dissolved in AcOH:$H_2O$ buffer (1000 µL, 1:1) that had been sparged with $N_2$. The reaction was mixed at 23 ºC for 6 h before it was quenched with DTT. The resulting precipitate was centrifuged, and the supernatant was removed. The pellet was also washed twice with $H_2O$ +0.1% formic acid and the combined washes were purified over a Luna PhenylHexyl column (10 x 250 mm, 5 µm, 100 Å, 4.7 mL/min) 20%-40% MeCN/$H_2O$/0.1% TFA over 30 min. The fractions were lyophilized to afford **S2a** (15.0 mg, 89% yield).

**Prep HPLC trace of Acm removal**

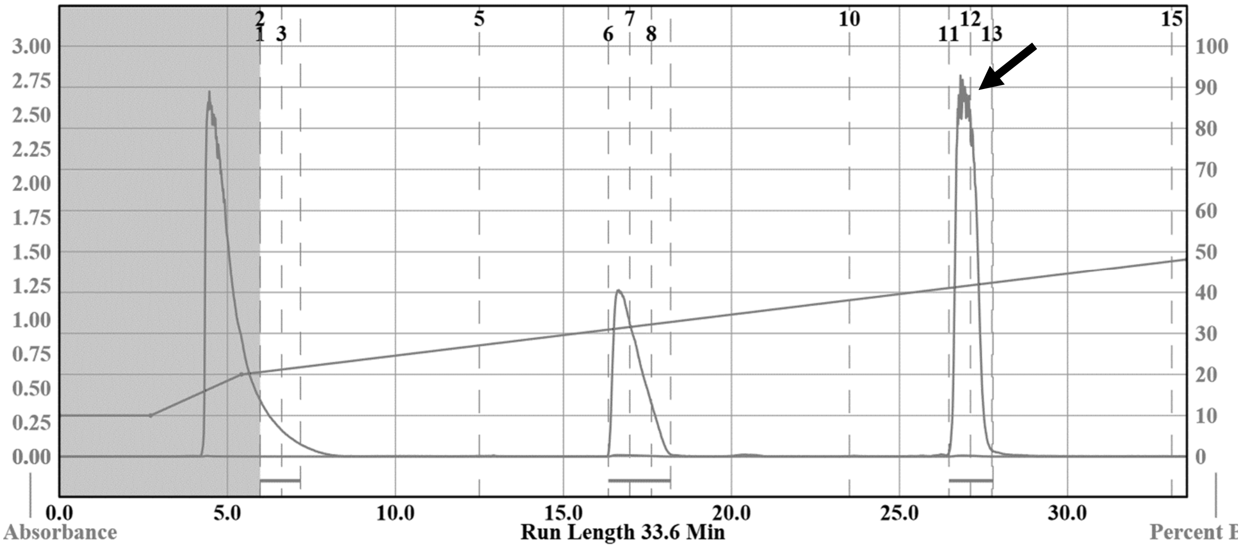

**Analytical LCMS**: Measured on an Agilent EC-C18 Poroshell column (4.6 x 100 mm, 4 µm, 100 Å, 40 °C, 1 mL/min) 5%-65% MeCN/$H_2O$/0.05% TFA over 60 min.

**Chemical Formula**: $C_{325}H_{530}N_{102}O_{103}S$
**Molecular Weight**: 7546.4860 g/mol

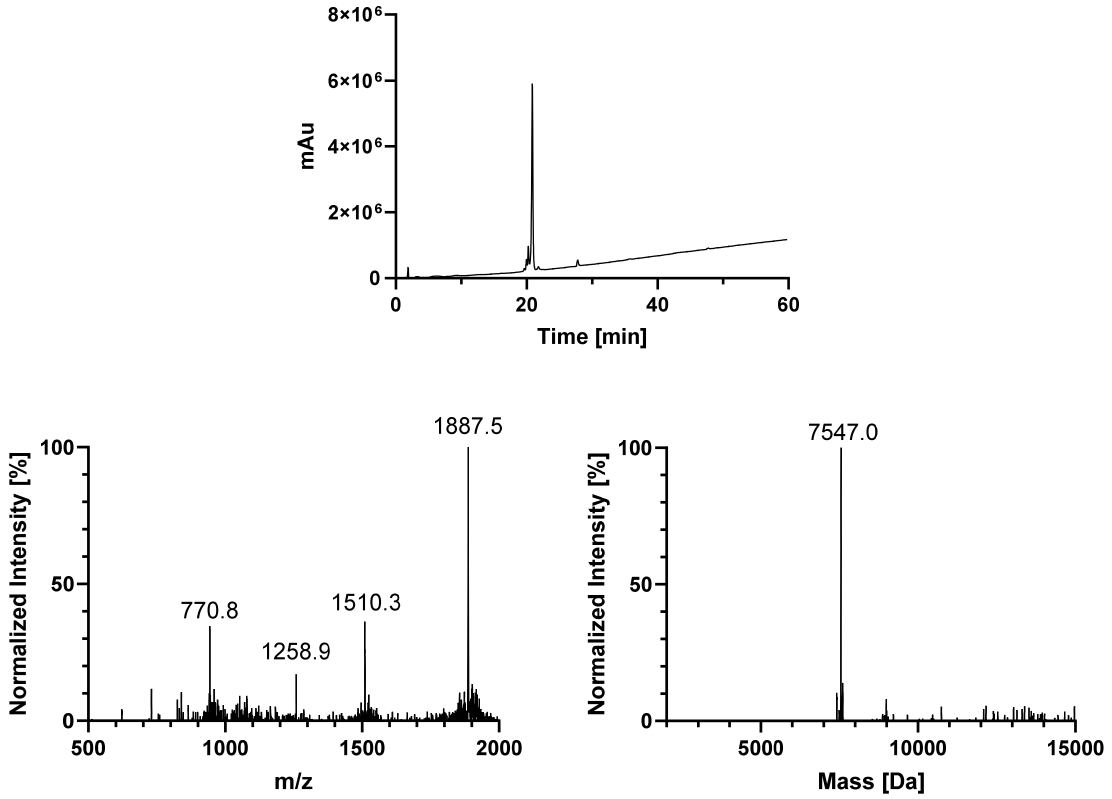
8×10⁶
6×10⁶
4×10⁶
2×10⁶
0
mAu
0
20
40
60
Time [min]
1887.5
770.8
1258.9
1510.3
100
50
0
Normalized Intensity [%]
500
1000
1500
2000
m/z
7547.0
100
50
0
Normalized Intensity [%]
5000
10000
15000
Mass [Da]

#### AcK369

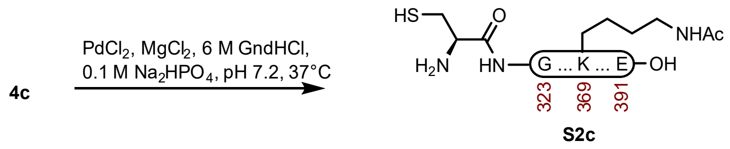

**Peptide S2c.** This Acm removal method was modified from the literature.[5] A 2 dram vial with **4c** (13.6 mg, 1.78 µmol) was dissolved in buffer (890 µL, 6 M GndHCl, 0.1 M $Na_2HPO_4$, pH 7.2) that had been sparged with $N_2$. The reaction was treated with $MgCl_2$ in buffer (25.4 µL, 0.1 mg/mL, 26.7 µmol) and $PdCl_2$ in buffer (15.8 µL, 0.1 mg/mL, 8.90 µmol). The reaction was mixed at 37 °C for 2 h before it was quenched with DTT. After 2 h a precipitate slowly formed. The resulting precipitate was centrifuged, and the supernatant was removed. The pellet was also washed twice with $H_2O$ +0.1% formic acid and the combined washes were purified over a Luna C4 column (10 x 250 mm, 10 µm, 100 Å, 4.7 mL/min) 20%-35% MeCN/$H_2O$/0.1% TFA over 30 min. The fractions were lyophilized to afford **S2c** (8.9 mg, 66% yield).

**Prep HPLC trace of Acm removal**

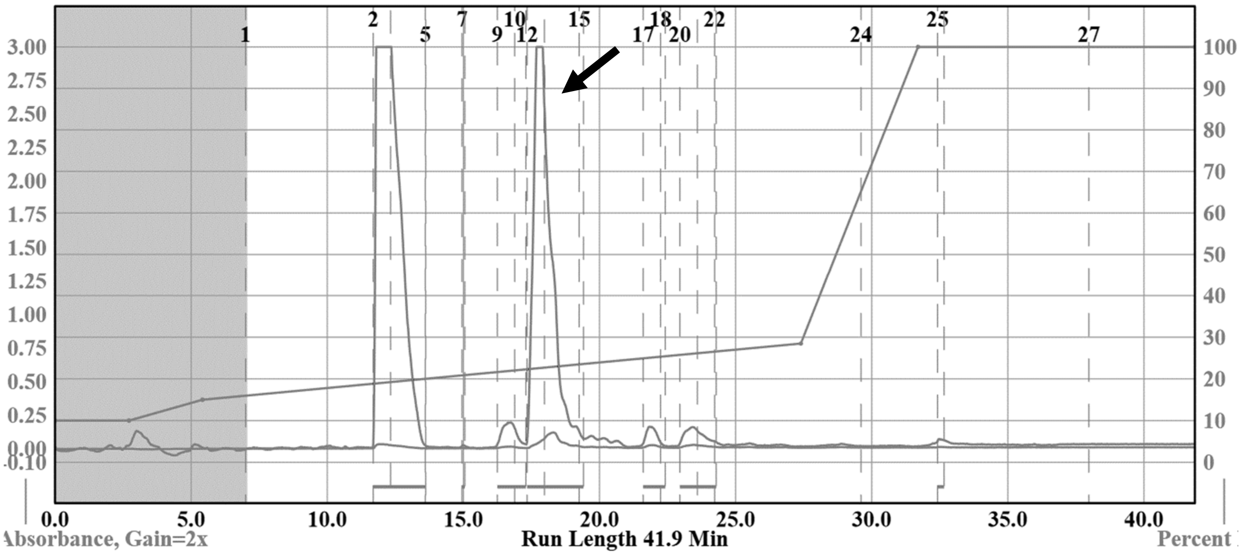

**Analytical LCMS**: Measured on an Agilent EC-C18 Poroshell column (4.6 x 100 mm, 4 µm, 100 Å, 40 °C, 1 mL/min) 5%-65% MeCN/$H_2O$/0.05% TFA over 60 min.

**Chemical Formula**: $C_{327}H_{532}N_{102}O_{104}S$
**Molecular Weight**: 7588.5230 g/mol

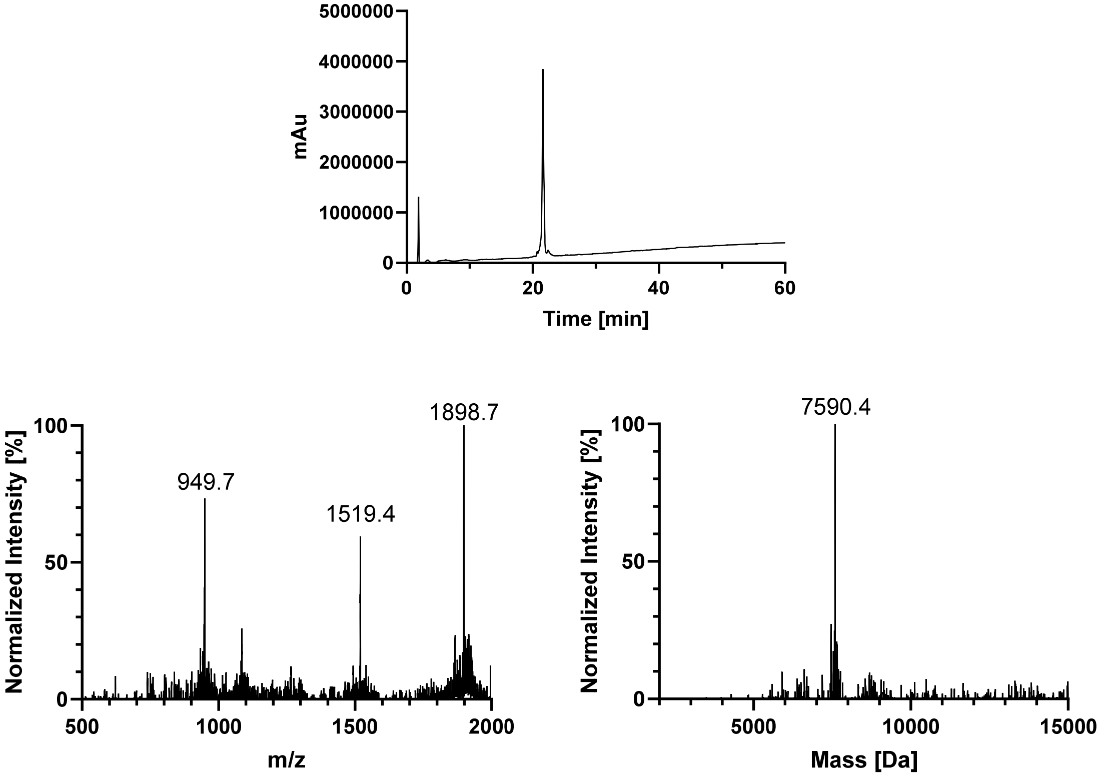
5000000
4000000
3000000
2000000
1000000
0
mAu
0
20
40
60
Time [min]
100
50
0
Normalized Intensity [%]
949.7
1519.4
1898.7
500
1000
1500
2000
m/z
100
50
0
Normalized Intensity [%]
7590.4
5000
10000
15000
Mass [Da]

#### AcK353

4d

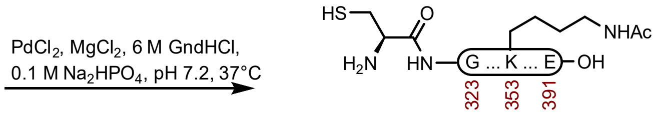

S2d

**Peptide S2d.** This Acm removal method was modified from the literature.[5] A 2 dram vial with **4d** (18.0 mg, 2.35 µmol) was dissolved in buffer (800 µL, 6 M GndHCl, 0.1 M $Na_2HPO_4$, pH 7.2) that had been sparged with $N_2$. The reaction was treated with $MgCl_2$ in buffer (33.6 µL, 0.1 mg/mL, 35.3 µmol) and $PdCl_2$ in buffer (20.8 µL, 0.1 mg/mL, 11.8 µmol). The reaction was mixed at 37 °C for 2 h before it was quenched with DTT. The resulting precipitate was centrifuged, and the supernatant was removed. The pellet was also washed twice with $H_2O$ +0.1% formic acid and the combined washes were purified over an Agilent C18 column (10 x 250 mm, 5 µm, 100 Å, 4.7 mL/min) 25%-35% MeCN/$H_2O$/0.1% TFA over 30 min. The fractions were lyophilized to afford **S2d** (7.0 mg, 39% yield).

**Prep HPLC trace of Acm removal**

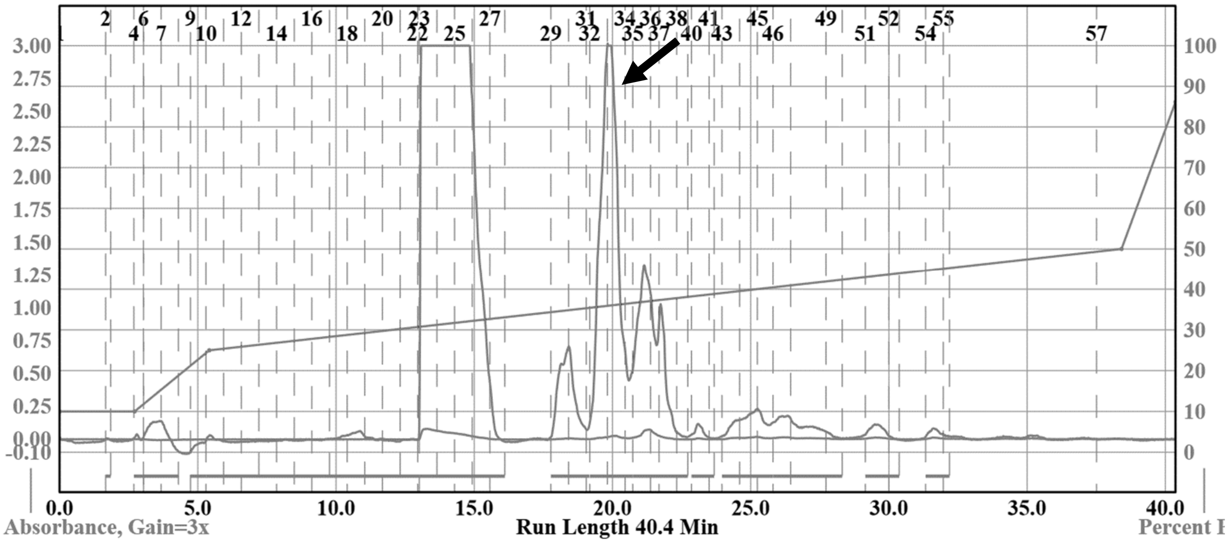

**Analytical LCMS**: Measured on an Agilent EC-C18 Poroshell column (4.6 x 100 mm, 4 µm, 100 Å, 40 °C, 1 mL/min) 5%-65% MeCN/$H_2O$/0.05% TFA over 60 min.

**Chemical Formula**: $C_{327}H_{532}N_{102}O_{104}S$
**Molecular Weight**: 7588.5230 g/mol

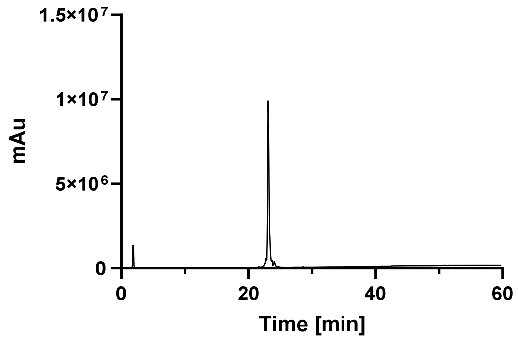
1.5×10⁷
1×10⁷
5×10⁶
0
mAu
0
20
40
60
Time [min]

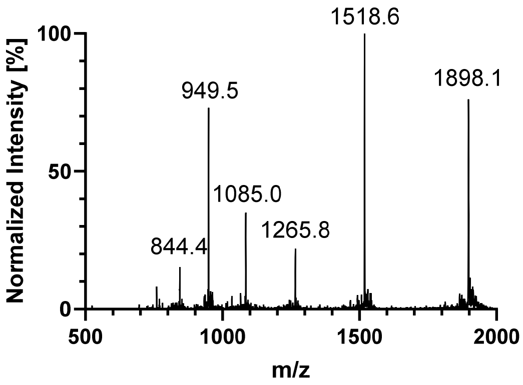
100
50
0
Normalized Intensity [%]
1518.6
949.5
1898.1
1085.0
1265.8
844.4
500
1000
1500
2000
m/z

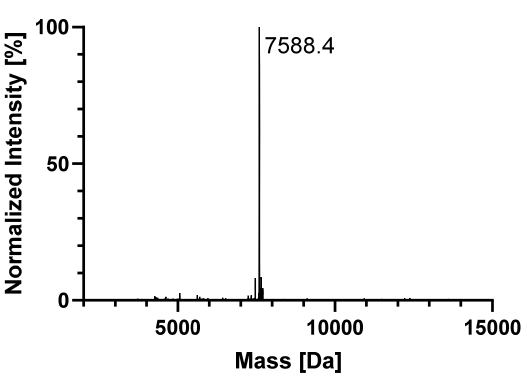
100
50
0
Normalized Intensity [%]
7588.4
5000
10000
15000
Mass [Da]

## pS356

4g

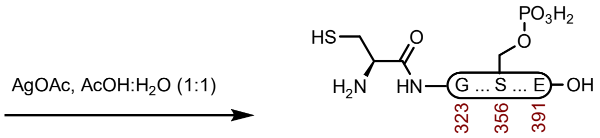

S2g

**Peptide S2g.** This Acm removal method was modified from the literature.[4] A 2 dram vial with **4g** (16.3 mg, 2.12 μmol) and AgOAc (8.2 mg, 49.1 μmol) was dissolved in AcOH:$H_2O$ buffer (530 μL, 1:1) that had been sparged with $N_2$. The reaction was mixed at 23 ºC for 6 h before it was quenched with DTT. The resulting precipitate was centrifuged, and the supernatant was removed. The pellet was also washed twice with $H_2O$+0.1% formic acid and the combined washes were purified over a Luna PhenylHexyl column (10 x 250 mm, 5 μm, 100 Å, 4.7 mL/min) 18%-30% MeCN/$H_2O$/0.1% TFA over 30 min. The fractions were lyophilized to afford **S2g** (15.4 mg, 95% yield).

**Prep HPLC trace of Acm removal**

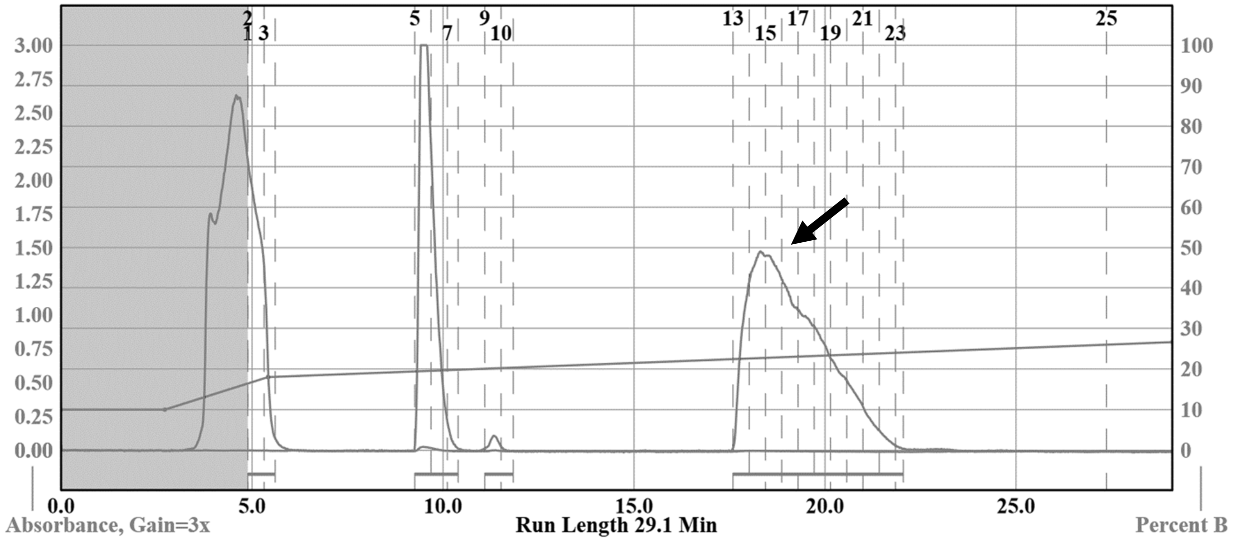

**Analytical LCMS**: Measured on an Agilent EC-C18 Poroshell column (4.6 x 100 mm, 4 μm, 100 Å, 40 °C, 1 mL/min) 5%-65% MeCN/$H_2O$/0.05% TFA over 60 min.

**Chemical Formula**: $C_{325}H_{531}N_{102}O_{106}PS$
**Molecular Weight**: 7626.4648 g/mol

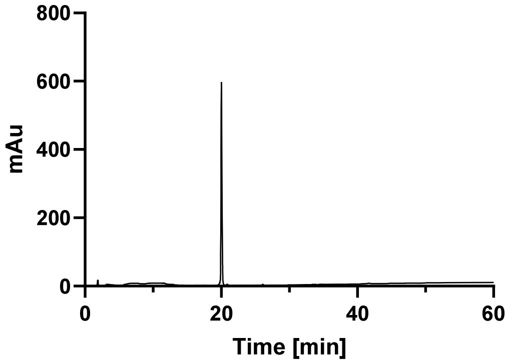
800
600
400
200
0
mAu
0
20
40
60
Time [min]

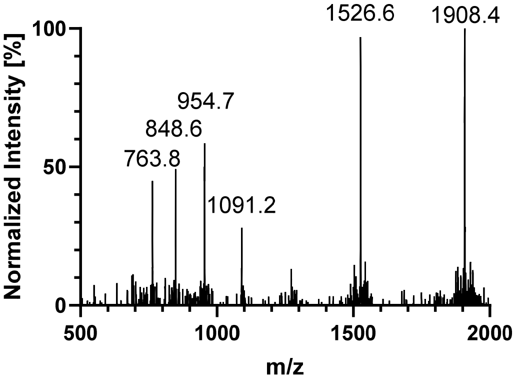
Normalized Intensity [%]
100
50
0
500
1000
1500
2000
m/z
1526.6
1908.4
954.7
848.6
763.8
1091.2

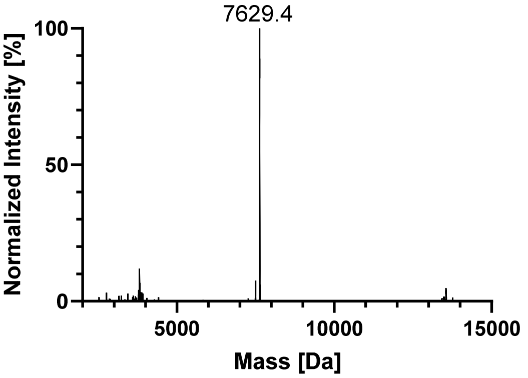
Normalized Intensity [%]
100
50
0
5000
10000
15000
Mass [Da]
7629.4

## pS352

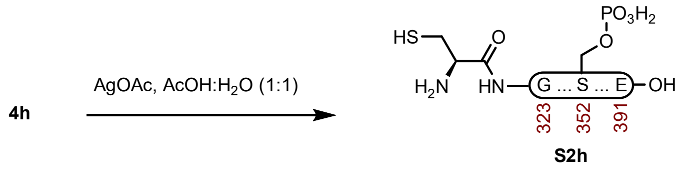

**Peptide S2h.** This Acm removal method was modified from the literature.[4] A 2 dram vial with **4h** (14.1 mg, 1.83 μmol) and AgOAc (8.2 mg, 49.1 μmol) was dissolved in AcOH:$H_2O$ buffer (450 μL, 1:1) that had been sparged with $N_2$. The reaction was mixed at 23 ºC for 5 h before it was quenched with DTT. The resulting precipitate was centrifuged, and the supernatant was removed. The pellet was also washed twice with $H_2O$ +0.1% formic acid and the combined washes were purified over a Luna PhenylHexyl column (10 x 250 mm, 5 μm, 100 Å, 4.7 mL/min) 18%-30% MeCN/$H_2O$/0.1% TFA over 30 min. The fractions were lyophilized to afford **S2h** (13.1 mg, 94% yield).

**Prep HPLC trace of Acm removal**

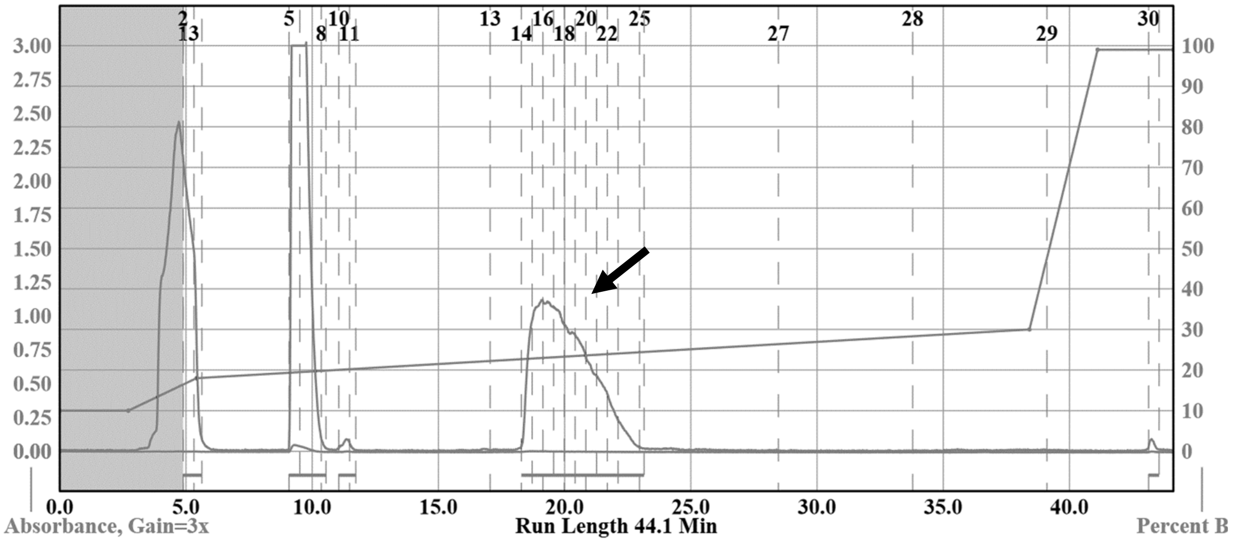

**Analytical LCMS**: Measured on an Agilent EC-C18 Poroshell column (4.6 x 100 mm, 4 μm, 100 Å, 40 °C, 1 mL/min) 5%-65% MeCN/$H_2O$/0.05% TFA over 60 min.

**Chemical Formula**: $C_{325}H_{531}N_{102}O_{106}PS$
**Molecular Weight**: 7626.4648 g/mol

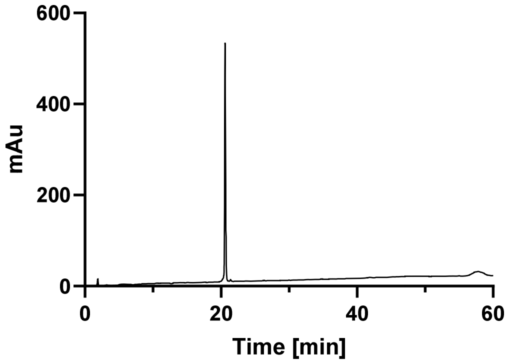
600
400
200
0
mAu
0
20
40
60
Time [min]

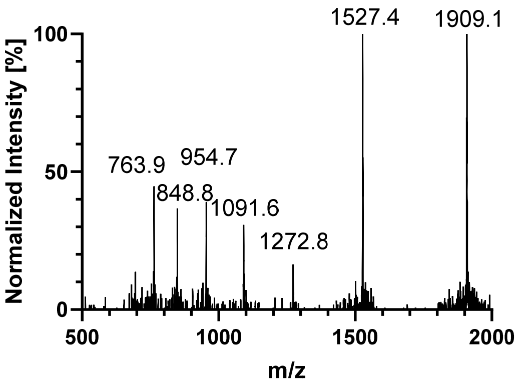
1527.4
1909.1
763.9
954.7
848.8
1091.6
1272.8
100
50
0
Normalized Intensity [%]
500
1000
1500
2000
m/z

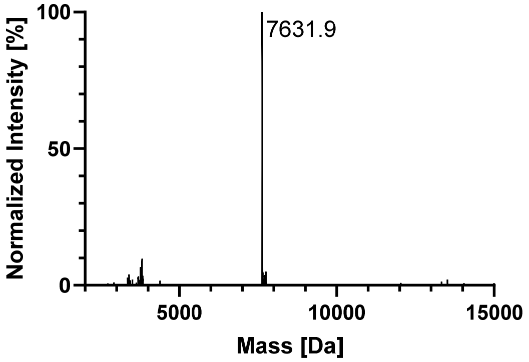
100
7631.9
50
0
Normalized Intensity [%]
5000
10000
15000
Mass [Da]

#### GlcNAcN359

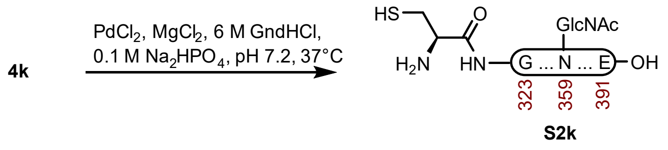

**Peptide S2k.** This Acm removal method was modified from the literature.[5] A 2 dram vial with **4k** (20.2 mg, 2.58 μmol) was dissolved in buffer (1300 μL, 6 M GndHCl, 0.1 M $Na_2HPO_4$, pH 7.2) that had been sparged with $N_2$. The reaction was treated with $MgCl_2$ in buffer (36.8 μL, 0.1 mg/mL, 38.7 μmol) and $PdCl_2$ in buffer (22.8 μL, 0.1 mg/mL, 12.9 μmol). The reaction was mixed at 37 °C for 2 h before it was quenched with DTT. The resulting precipitate was centrifuged, and the supernatant was removed. The pellet was also washed twice with $H_2O$ +0.1% formic acid and the combined washes were purified over a Luna C4 column (10 x 250 mm, 10 μm, 100 Å, 4.7 mL/min) 20%-55% MeCN/$H_2O$/0.1% TFA over 30 min. The fractions were lyophilized to afford **S2k** (17.5 mg, 88% yield).

**Prep HPLC trace of Acm removal**

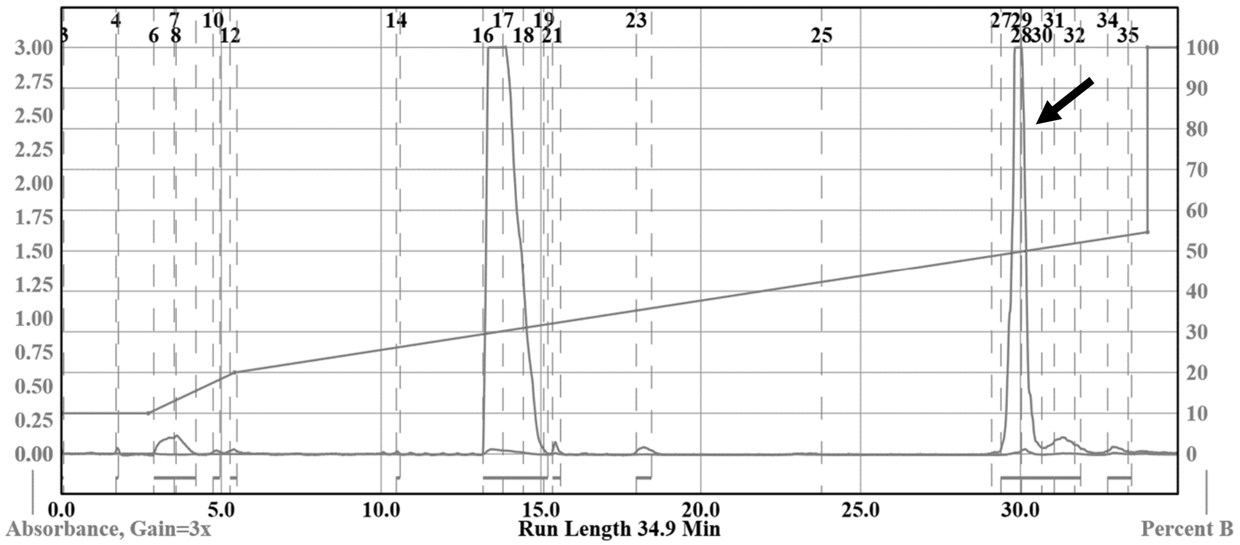

**Analytical LCMS**: Measured on an Agilent EC-C18 Poroshell column (4.6 x 100 mm, 4 μm, 100 Å, 40 °C, 1 mL/min) 5%-65% MeCN/$H_2O$/0.05% TFA over 60 min.

**Chemical Formula**: $C_{333}H_{543}N_{103}O_{108}S$
**Molecular Weight**: 7749.6800 g/mol

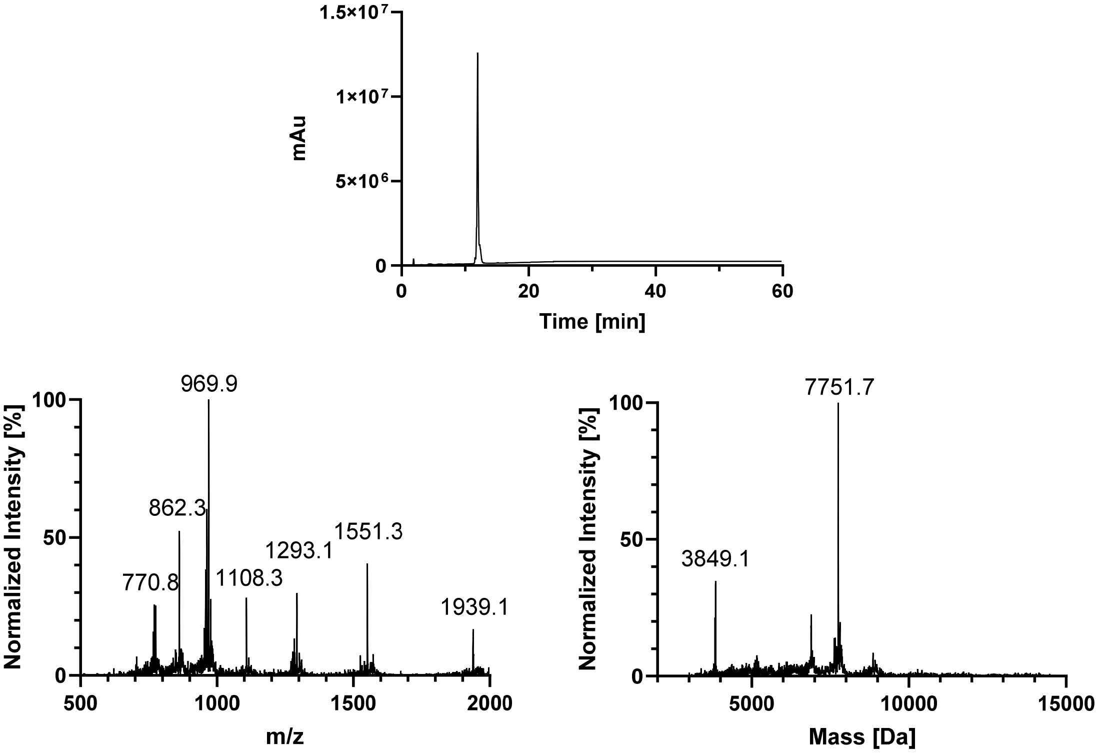
1.5×10⁷
1×10⁷
5×10⁶
0
mAu
0
20
40
60
Time [min]
Normalized Intensity [%]
100
50
0
969.9
862.3
770.8
1108.3
1293.1
1551.3
1939.1
500
1000
1500
2000
m/z
Normalized Intensity [%]
100
50
0
7751.7
3849.1
5000
10000
15000
Mass [Da]

#### GlcNAcN359 + AcK353

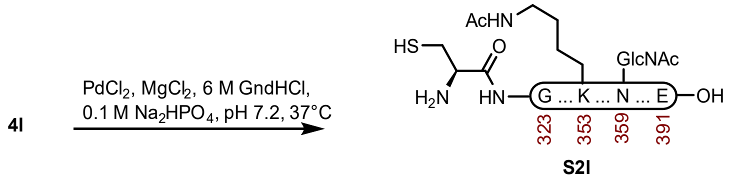

**Peptide S2l.** This Acm removal method was modified from the literature.[5] A 2 dram vial with **4l** (10.0 mg, 1.27 μmol) was dissolved in buffer (635 μL, 6 M GndHCl, 0.1 M $Na_2HPO_4$, pH 7.2) that had been sparged with $N_2$. The reaction was treated with $MgCl_2$ in buffer (18.1 μL, 0.1 mg/mL, 19.1 μmol) and $PdCl_2$ in buffer (11.3 μL, 0.1 mg/mL, 6.35 μmol). The reaction was mixed at 37 °C for 2 h before it was quenched with DTT (no metal sulfide precipitate formed) and purified over a Luna C4 column (10 x 250 mm, 10 μm, 100 Å, 4.7 mL/min) 20%-35% MeCN/$H_2O$/0.1% TFA over 30 min. The fractions were lyophilized to afford **S2l** (8.9 mg, 90% yield).

##### Prep HPLC trace of Acm removal

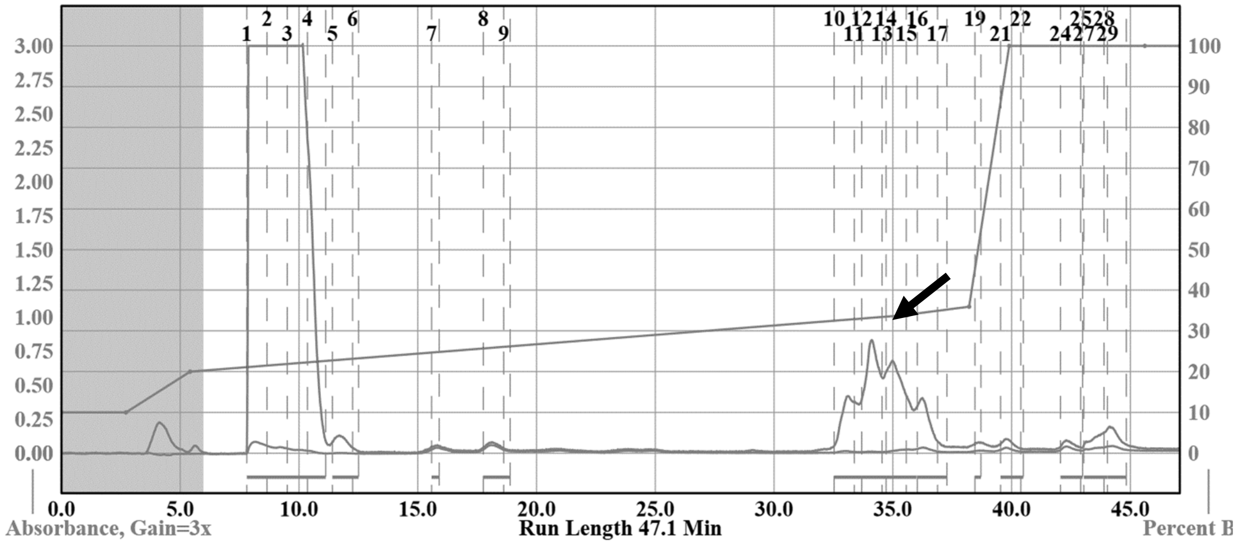

**Analytical LCMS**: Measured on an Agilent EC-C18 Poroshell column (4.6 x 100 mm, 4 μm, 100 Å, 40 °C, 1 mL/min) 5%-65% MeCN/$H_2O$/0.05% TFA over 60 min.

**Chemical Formula**: $C_{335}H_{545}N_{103}O_{109}S$
**Molecular Weight**: 7791.7170 g/mol

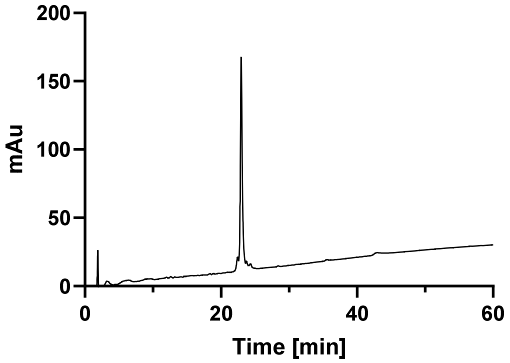
mAu
200
150
100
50
0
0
20
40
60
Time [min]

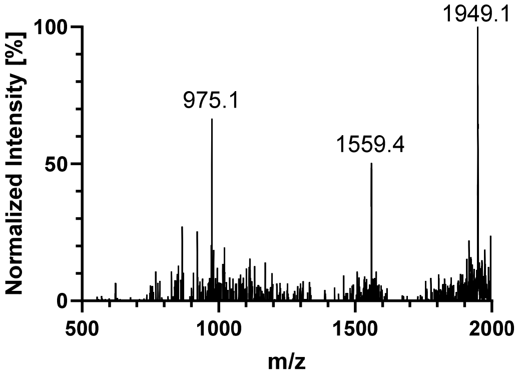
Normalized Intensity [%]
100
50
0
975.1
1559.4
1949.1
500
1000
1500
2000
m/z

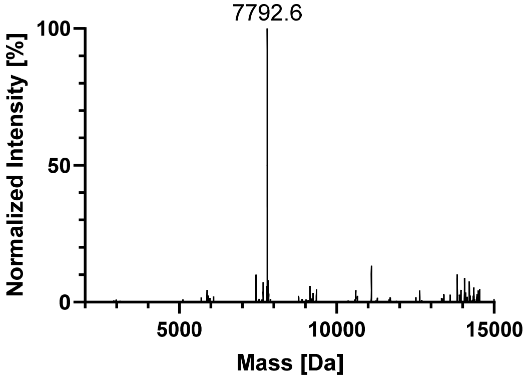
Normalized Intensity [%]
100
50
0
7792.6
5000
10000
15000
Mass [Da]

#### 8. C291-E391 (NCL-Thz Removal)

### WT

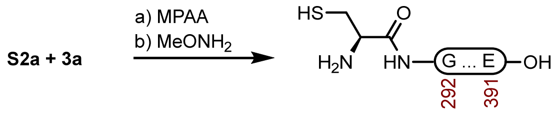

**5a**

**Peptide 5a.** A 1 dram vial with **S2a** (15.8 mg, 2.09 µmol) and **3a** (9.7 mg, 2.94 µmol) was dissolved in fresh NCL buffer (1000 µL, 200 mM MPAA, 6 M GndHCl, 0.2 M $Na_2HPO_4$, 20 mM TCEP HCl, pH 7.2) that had been sparged with $N_2$. The reaction was mixed at 23 ºC for 3 h before adding Thz removal buffer (1000 µL, 400 mM $MeONH_2$ HCl, 6 M GndHCl, 0.2 M $NaH_2PO_4$, 20 mM TCEP HCl, pH 2.5), and adjusted to pH 4.1. The reaction was mixed at 23 ºC for 3 h before it was purified over a Luna PhenylHexyl column (10 x 250 mm, 5 µm, 100 Å, 4.7 mL/min) 20%-40% MeCN/$H_2O$/0.1% TFA over 30 min. The fractions were lyophilized to afford **5a** (13.5 mg, 60% yield).

**Crude LCMS (NCL)**: Measured on an Agilent EC-C18 Poroshell column (4.6 x 100 mm, 4 µm, 100 Å, 40 °C, 1 mL/min) 5%-65% MeCN/$H_2O$/0.05% TFA over 60 min.

**LC trace (214 nm) of unpurified peptide**

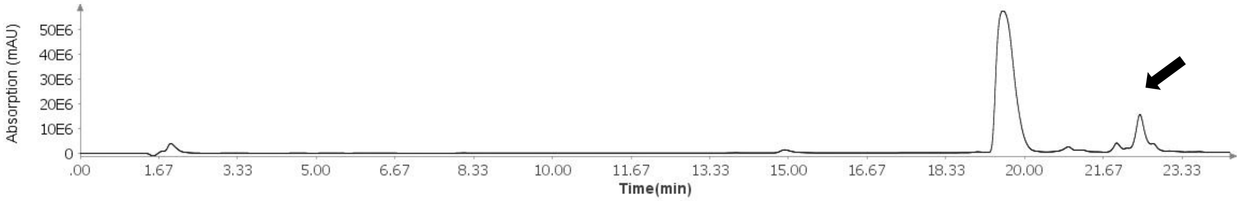

**Total Ion Current (TIC) of unpurified peptide**

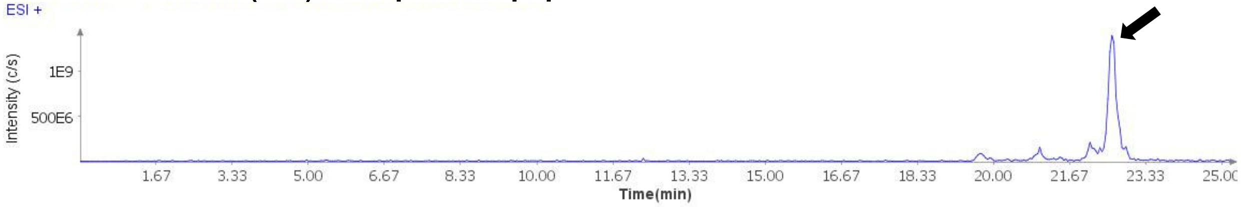

**Extracted Ion Current (XIC) of unpurified peptide**

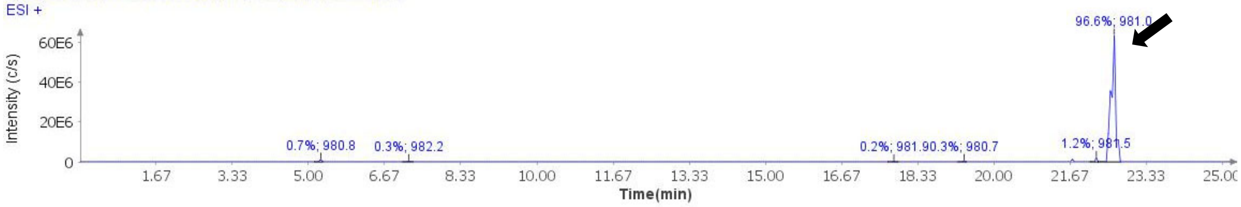

**Mass Spectrum (ESI) of the expected product (unpurified sample)**

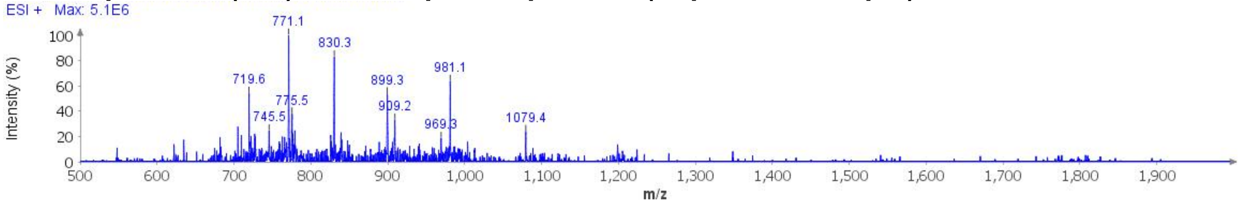

**Deconvoluted Mass Spectrum (ESI) of the expected product (unpurified sample)**

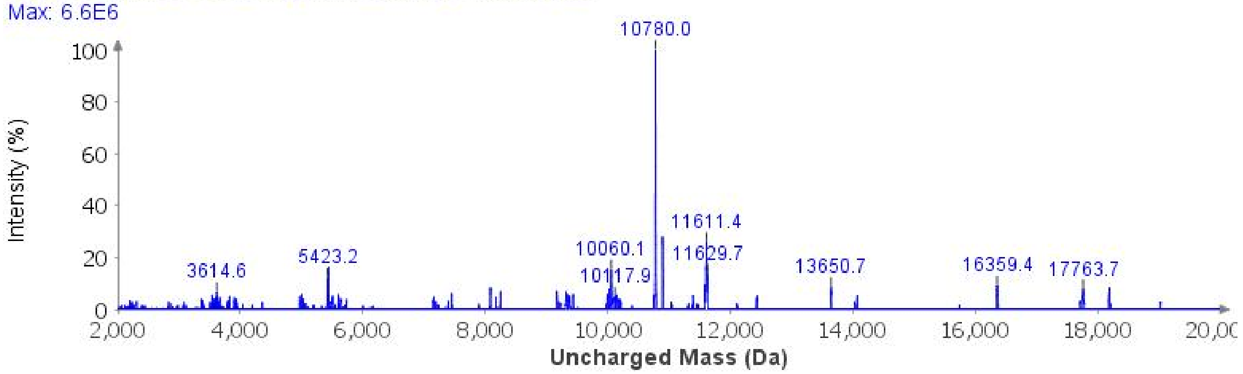

**Sample Displacement Mode Chromatography**:[6] The material was further purified by sample displacement mode over a Zorbax C18 column (4.6 x 250 mm, 5 µm, 300 Å, 4.7 mL/min, 40 °C) 5%-45% MeCN/$H_2O$/0.05%TFA over 60 min.

**Sample Displacement Mode trace (214 nm)**

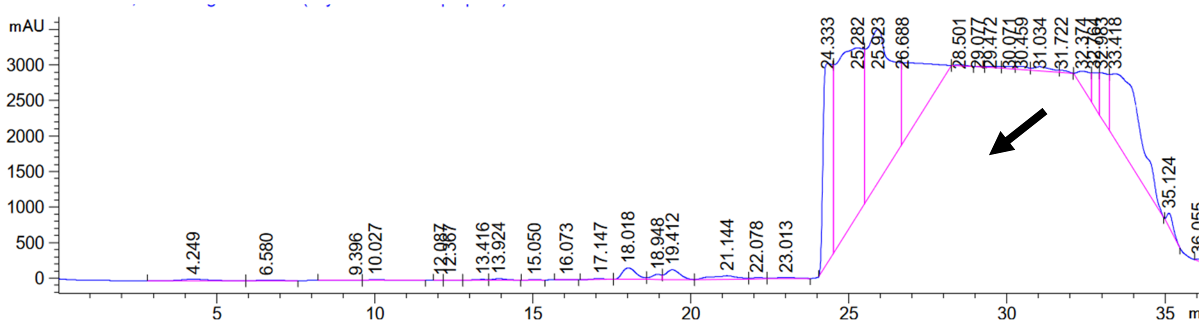

**Analytical LCMS**: Measured on an Agilent EC-C18 Poroshell column (4.6 x 100 mm, 4 µm, 100 Å, 40 °C, 1 mL/min) 5%-65% MeCN/$H_2O$/0.05% TFA over 60 min.

**Chemical Formula**: $C_{467}H_{766}N_{142}O_{146}S_2$
**Molecular Weight**: 10770.2330 g/mol

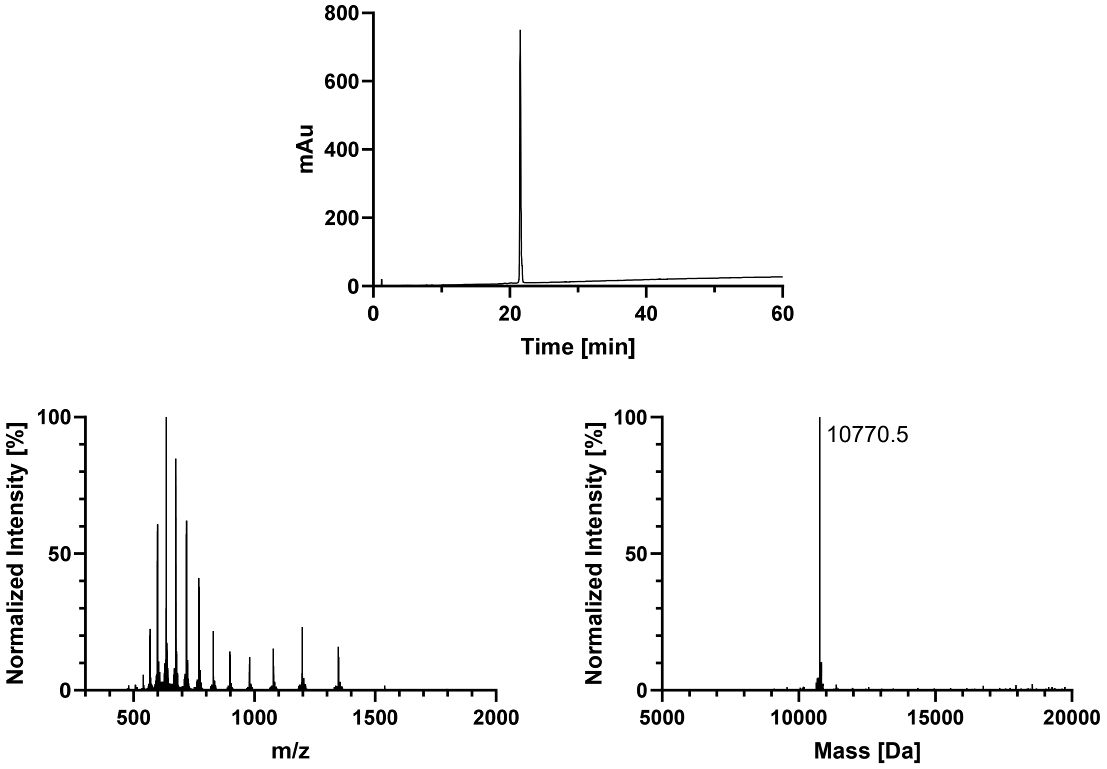

mAu
Time [min]
Normalized Intensity [%]
m/z
Normalized Intensity [%]
10770.5
Mass [Da]

## P301S

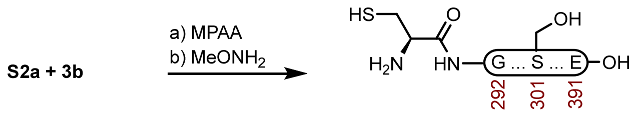

**5b**

**Peptide 5b.** A 1 dram vial with **S2a** (9.5 mg, 1.26 µmol) and **3b** (5.8 mg, 1.69 µmol) was dissolved in fresh NCL buffer (1000 µL, 200 mM MPAA, 6 M GndHCl, 0.2 M $Na_2HPO_4$, 20 mM TCEP HCl, pH 7.0) that had been sparged with $N_2$. The reaction was mixed at 37 °C for 5 h before adding Thz removal buffer (1000 µL, 400 mM $MeONH_2$ HCl, 6 M GndHCl, 0.2 M $NaH_2PO_4$, 20 mM TCEP HCl, pH 2.4), and adjusted to pH 4.0. The reaction was mixed at 23 ºC for 2 h before it was purified over a Luna PhenylHexyl column (10 x 250 mm, 5 µm, 100 Å, 4.7 mL/min) 25%-35% MeCN/$H_2O$/0.1% TFA over 30 min. The fractions were lyophilized to afford **5b** (5.8 mg, 43% yield).

**Crude LCMS (NCL)**: Measured on an Agilent EC-C18 Poroshell column (4.6 x 100 mm, 4 µm, 100 Å, 40 °C, 1 mL/min) 5%-65% MeCN/$H_2O$/0.05% TFA over 60 min.

##### LC trace (214 nm) of unpurified peptide

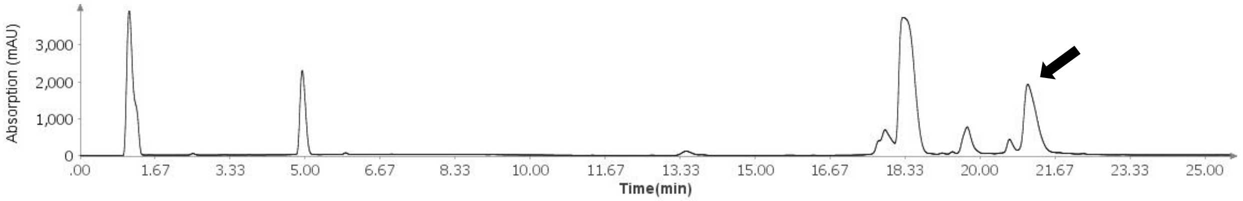

##### Total Ion Current (TIC) of unpurified peptide

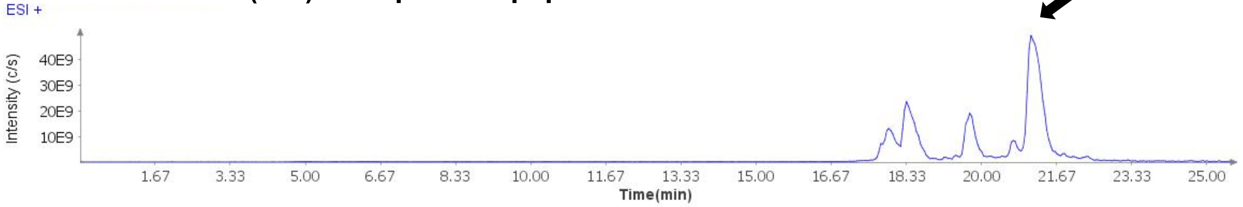

##### Extracted Ion Current (XIC) of unpurified peptide

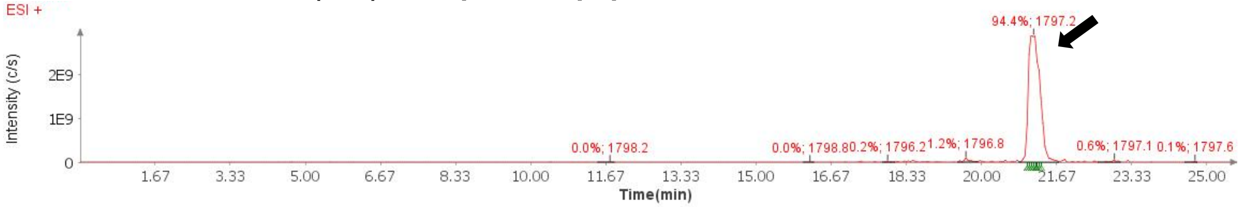

##### Mass Spectrum (ESI) of the expected product (unpurified sample)

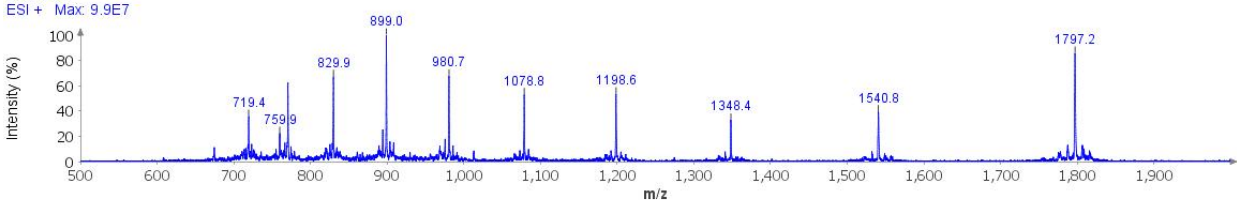

**Deconvoluted Mass Spectrum (ESI) of the expected product (unpurified sample)**

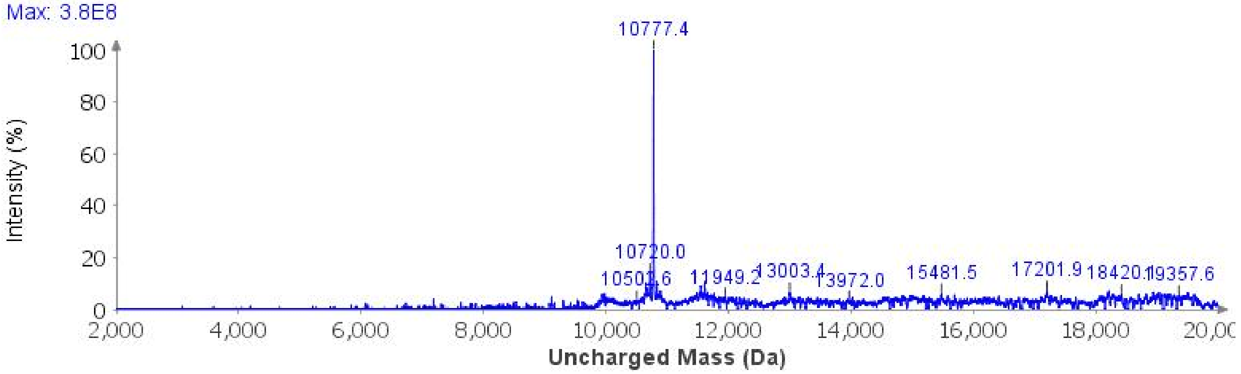

**Sample Displacement Mode Chromatography**:[6] The material was further purified by sample displacement mode over a Zorbax C18 column (4.6 x 250 mm, 5 µm, 300 Å, 4.7 mL/min, 40 °C) 5%-45% MeCN/$H_2O$/0.05%TFA over 60 min.

**Sample Displacement Mode trace (214 nm)**

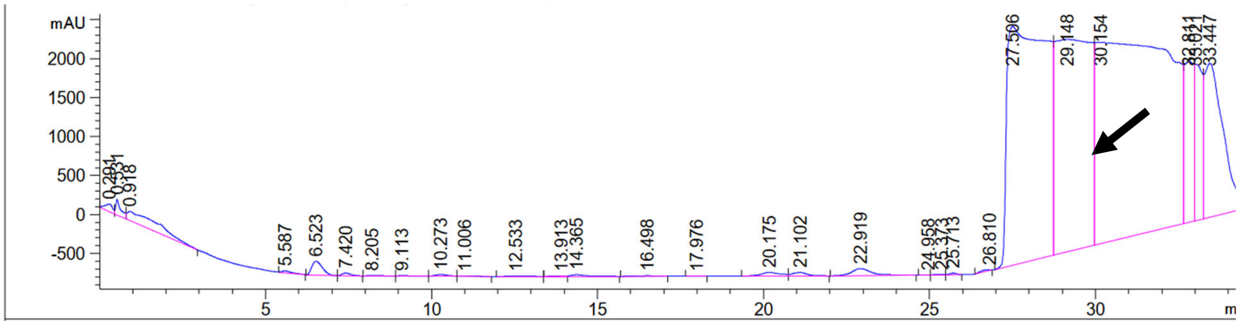

**Analytical LCMS**: Measured on an Agilent EC-C18 Poroshell column (4.6 x 100 mm, 4 µm, 100 Å, 40 °C, 1 mL/min) 5%-65% MeCN/$H_2O$/0.05% TFA over 60 min.

**Chemical Formula**: $C_{465}H_{764}N_{142}O_{147}S_2$
**Molecular Weight**: 10760.1940 g/mol

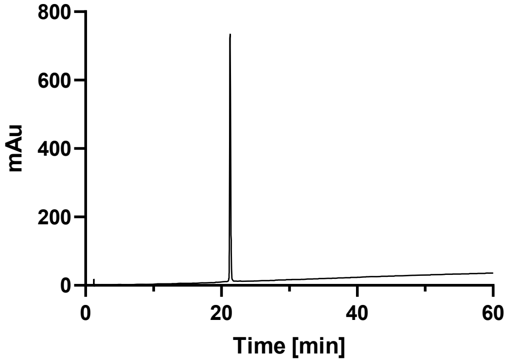

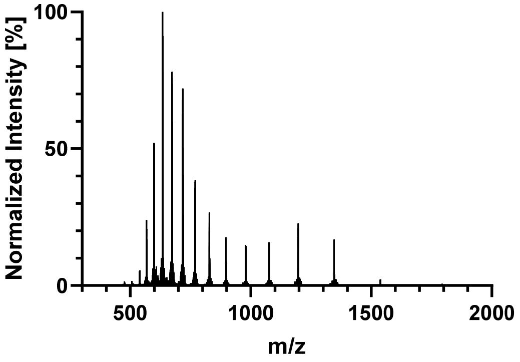
Normalized Intensity [%]
100
50
0
500
1000
1500
2000
m/z

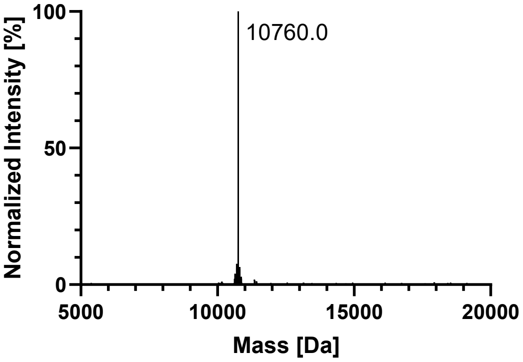
Normalized Intensity [%]
100
50
0
10760.0
5000
10000
15000
20000
Mass [Da]

#### AcK369

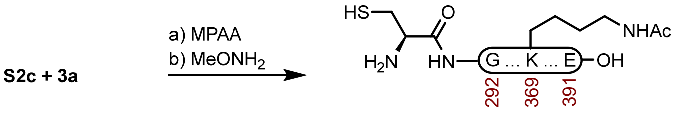

**5c**

**Peptide 5c.** A 2 dram vial with **S2c** (8.9 mg, 1.17 μmol) and **3a** (5.3 mg, 1.61 μmol) was dissolved in fresh NCL buffer (585 μL, 200 mM MPAA, 6 M GndHCl, 0.2 M $Na_2HPO_4$, 20 mM TCEP HCl, pH 7.2) that had been sparged with $N_2$. The reaction was mixed at 23 ºC for 3 h before adding Thz removal buffer (585 μL, 400 mM $MeONH_2$ HCl, 6 M GndHCl, 0.2 M $NaH_2PO_4$, 20 mM TCEP HCl, pH 2.5), and adjusted to pH 4.0. The reaction was mixed at 23 ºC for 5 h before it was purified over a Luna PhenylHexyl column (10 x 250 mm, 5 μm, 100 Å, 4.7 mL/min) 15%-40% MeCN/$H_2O$/0.1% TFA over 30 min. The fractions were lyophilized to afford **5c** (11.9 mg, 94% yield).

**Crude LCMS (NCL)**: Measured on an Agilent EC-C18 Poroshell column (4.6 x 100 mm, 4 μm, 100 Å, 40 °C, 1 mL/min) 5%-65% MeCN/$H_2O$/0.05% TFA over 60 min.

##### LC trace (214 nm) of unpurified peptide

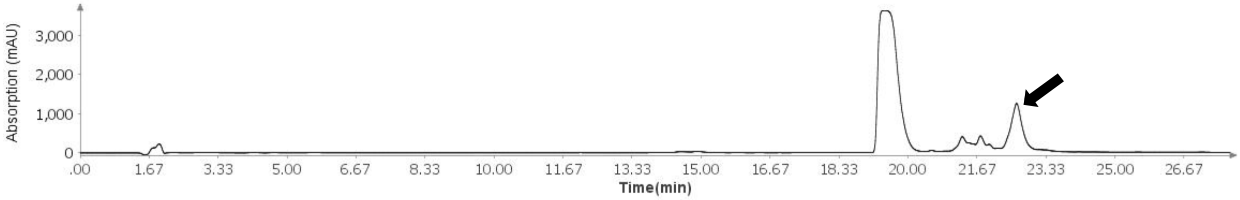

##### Total Ion Current (TIC) of unpurified peptide

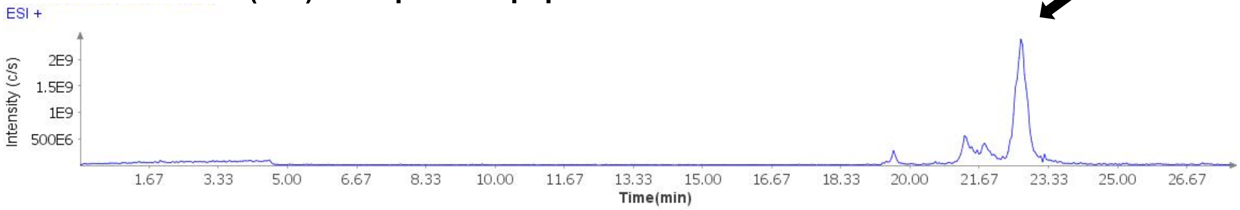

##### Extracted Ion Current (XIC) of unpurified peptide

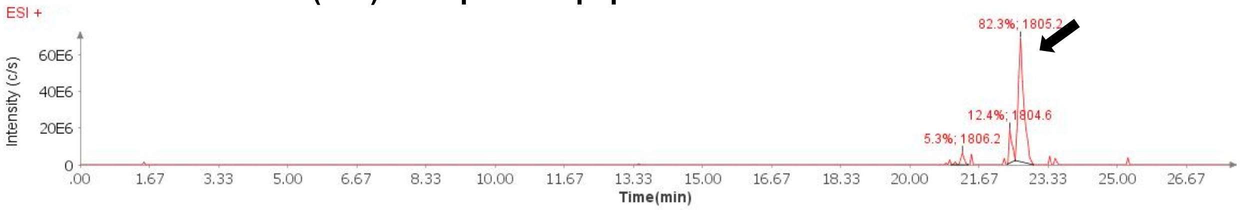

##### Mass Spectrum (ESI) of the expected product (unpurified sample)

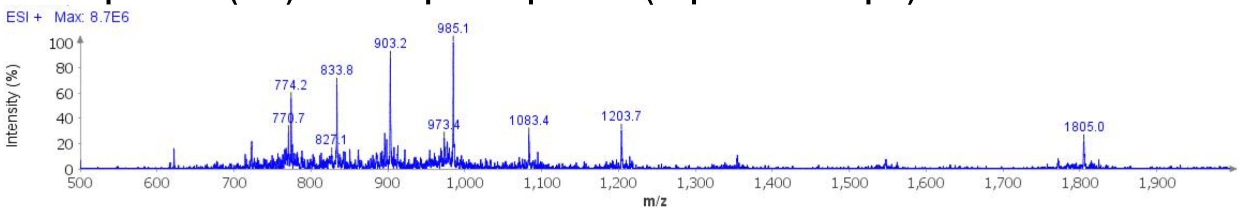

**Deconvoluted Mass Spectrum (ESI) of the expected product (unpurified sample)**

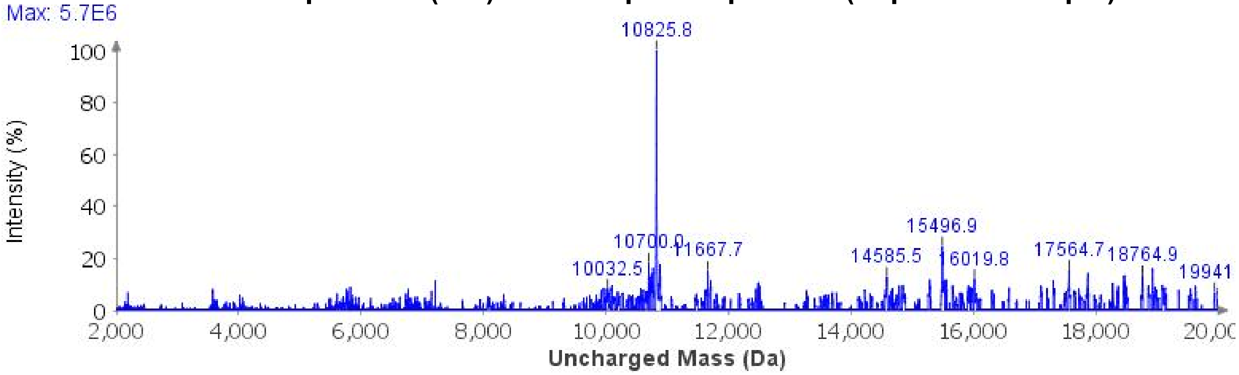

**Sample Displacement Mode Chromatography**:[6] The material was further purified by sample displacement mode over a Zorbax C18 column (4.6 x 250 mm, 5 µm, 300 Å, 4.7 mL/min, 40 °C) 5%-45% MeCN/$H_2O$/0.05%TFA over 60 min.

**Sample Displacement Mode trace (214 nm)**

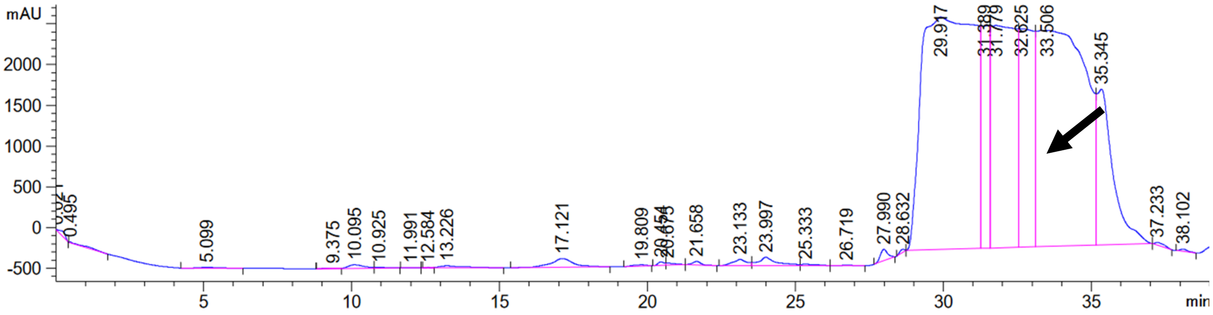

**Analytical LCMS**: Measured on an Agilent EC-C18 Poroshell column (4.6 x 100 mm, 4 µm, 100 Å, 40 °C, 1 mL/min) 5%-65% MeCN/$H_2O$/0.05% TFA over 60 min.

**Chemical Formula**: $C_{469}H_{768}N_{142}O_{147}S_2$
**Molecular Weight**: 10812.2700 g/mol

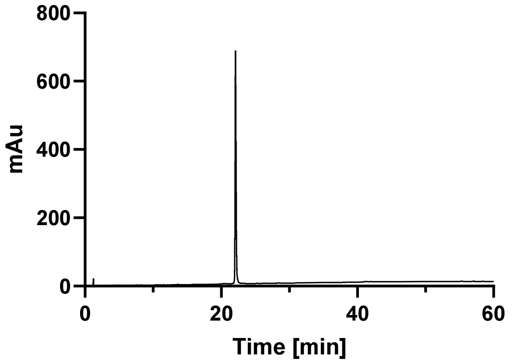
800
600
400
200
0
mAu
0
20
40
60
Time [min]

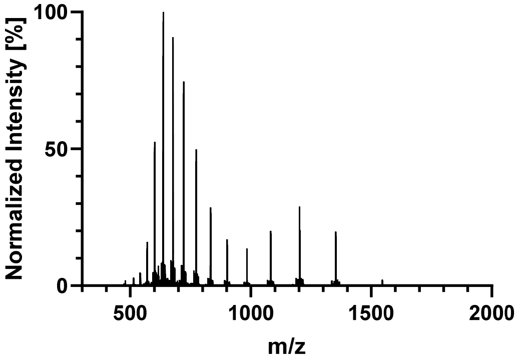
100
50
0
Normalized Intensity [%]
500
1000
1500
2000
m/z

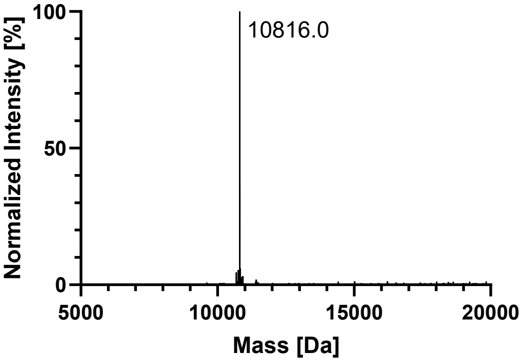
100
50
0
Normalized Intensity [%]
10816.0
5000
10000
15000
20000
Mass [Da]

##### AcK353

S2d + 3a

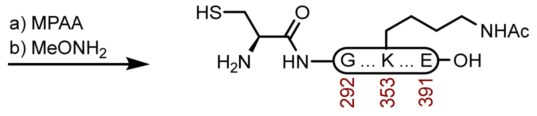

**5d**

**Peptide 5d.** A 2 dram vial with **S2d** (7.0 mg, 0.922 µmol) and **3a** (4.4 mg, 1.33 µmol) was dissolved in fresh NCL buffer (400 µL, 200 mM MPAA, 6 M GndHCl, 0.2 M $Na_2HPO_4$, 20 mM TCEP HCl, pH 7.1) that had been sparged with $N_2$. The reaction was mixed at 23 ºC for 4 h before adding Thz removal buffer (400 µL, 400 mM $MeONH_2$ HCl, 6 M GndHCl, 0.2 M $NaH_2PO_4$, 20 mM TCEP HCl, pH 2.4), and adjusted to pH 4.0. The reaction was mixed at 23 ºC for 5 h before it was purified over a Luna PhenylHexyl column (10 x 250 mm, 5 µm, 100 Å, 4.7 mL/min) 20%-60% $MeCN/H_2O$/0.1% TFA over 30 min. The fractions were lyophilized to afford **5d** (8.4 mg, 84% yield).

**Crude LCMS (NCL)**: Measured on an Agilent EC-C18 Poroshell column (4.6 x 100 mm, 4 µm, 100 Å, 40 °C, 1 mL/min) 5%-65% $MeCN/H_2O$/0.05% TFA over 60 min.

**LC trace (214 nm) of unpurified peptide**

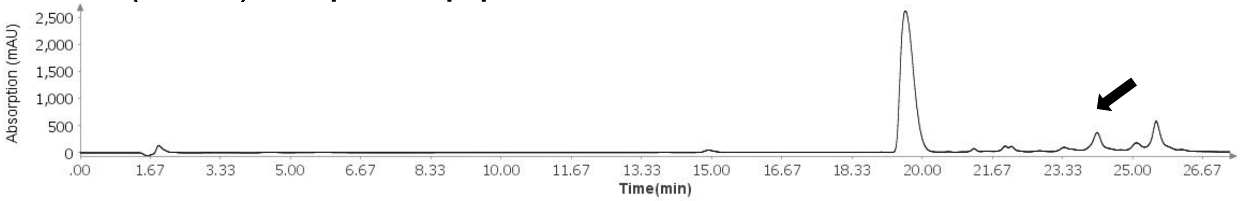

**Total Ion Current (TIC) of unpurified peptide**

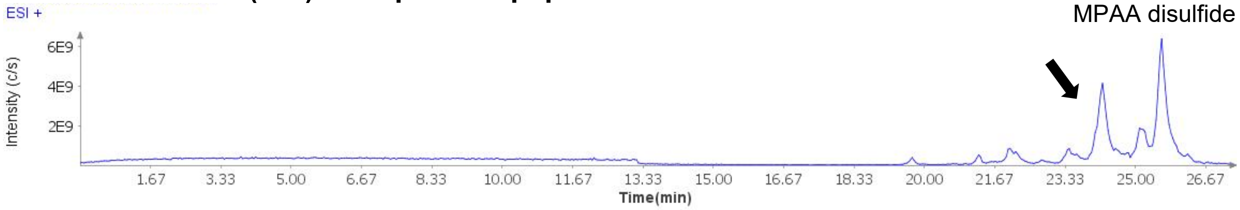

**Extracted Ion Current (XIC) of unpurified peptide**

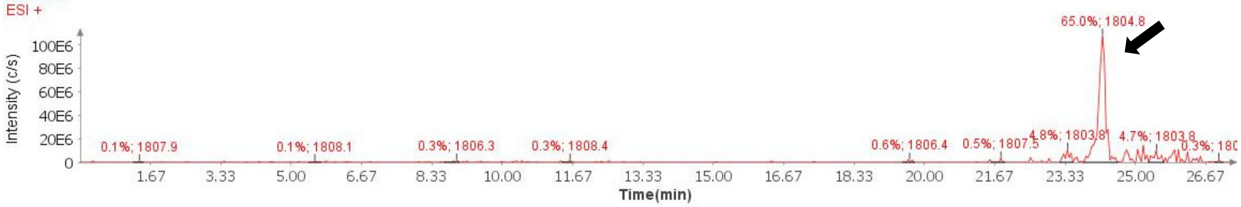

**Mass Spectrum (ESI) of the expected product (unpurified sample)**

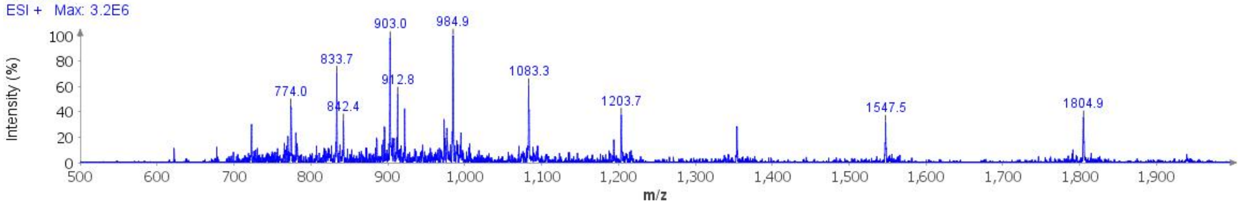

**Deconvoluted Mass Spectrum (ESI) of the expected product (unpurified sample)**

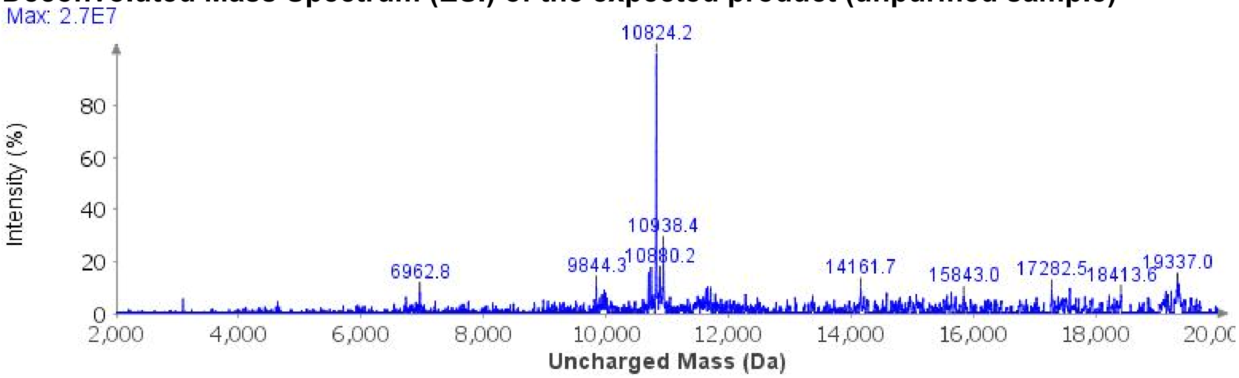

**Sample Displacement Mode Chromatography**:[6] The material was further purified by sample displacement mode over a Zorbax C18 column (4.6 x 250 mm, 5 µm, 300 Å, 4.7 mL/min, 40 °C) 5%-45% MeCN/$H_2O$/0.05%TFA over 60 min.

**Sample Displacement Mode trace (214 nm)**

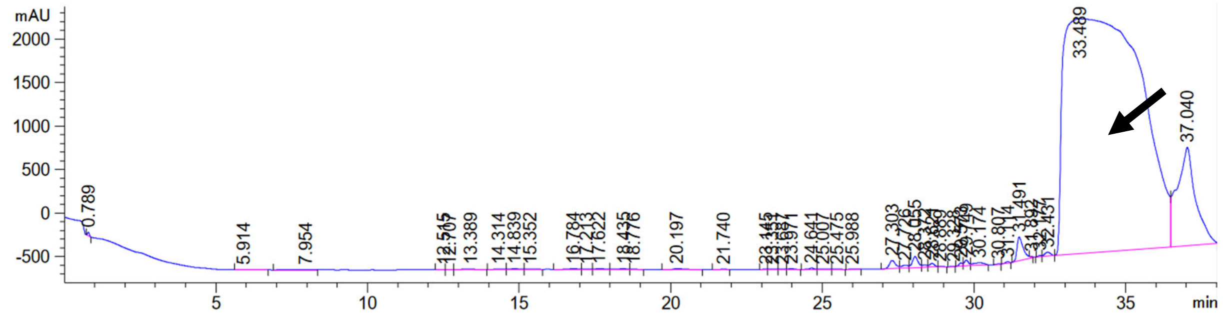

**Analytical LCMS**: Measured on an Agilent EC-C18 Poroshell column (4.6 x 100 mm, 4 µm, 100 Å, 40 °C, 1 mL/min) 5%-65% MeCN/$H_2O$/0.05% TFA over 60 min.

**Chemical Formula**: $C_{469}H_{768}N_{142}O_{147}S_2$
**Molecular Weight**: 10812.2700 g/mol

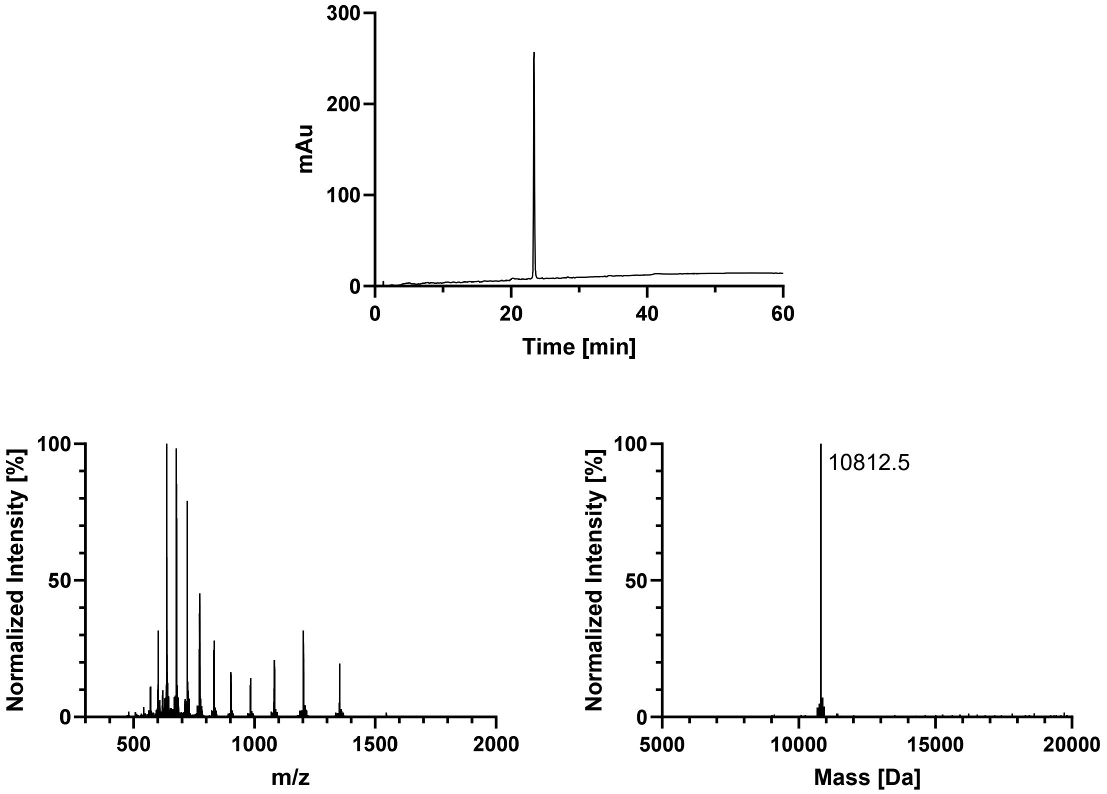
300
200
100
0
mAu
0
20
40
60
Time [min]
Normalized Intensity [%]
100
50
0
500
1000
1500
2000
m/z
Normalized Intensity [%]
100
50
0
10812.5
5000
10000
15000
20000
Mass [Da]

#### AcK317

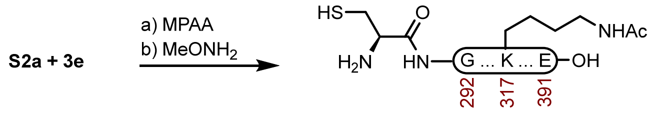

**5e**

**Peptide 5e.** A 1-dram vial with **S2a** (5.0 mg, 0.662 μmol) and **3e** (3.7 mg, 1.07 μmol) was dissolved in fresh NCL buffer (331 μL, 200 mM MPAA, 6 M GndHCl, 0.2 M $Na_2HPO_4$, 20 mM TCEP HCl, pH 7.0) that had been sparged with $N_2$. The reaction was mixed at 23 ºC for 5 h before adding Thz removal buffer (331 μL, 400 mM $MeONH_2$ HCl, 6 M GndHCl, 0.2 M $NaH_2PO_4$, 20 mM TCEP HCl, pH 2.4), and adjusted to pH 4.0. The reaction was mixed at 23 ºC for 3 h before it was purified over a Luna PhenylHexyl column (10 x 250 mm, 5 μm, 100 Å, 4.7 mL/min) 15%-40% MeCN/$H_2O$/0.1% TFA over 30 min. The fractions were lyophilized to afford **5e** (4.8 mg, 67% yield).

**Crude LCMS (NCL)**: Measured on an Agilent EC-C18 Poroshell column (4.6 x 100 mm, 4 μm, 100 Å, 40 °C, 1 mL/min) 5%-65% MeCN/$H_2O$/0.05% TFA over 60 min.

##### LC trace (214 nm) of unpurified peptide

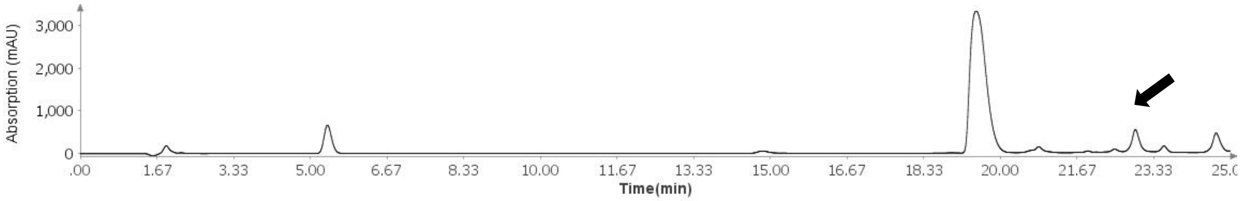

##### Total Ion Current (TIC) of unpurified peptide

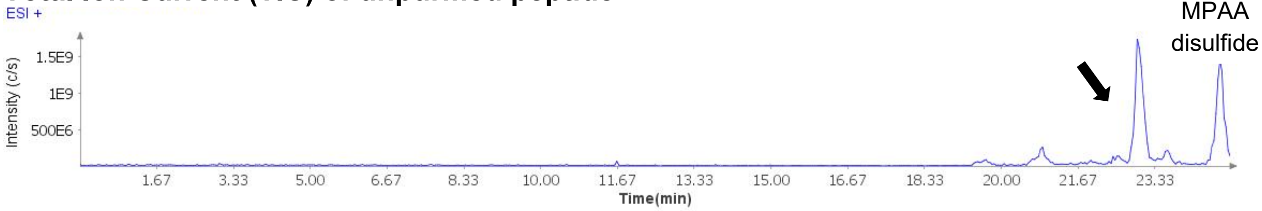

##### Extracted Ion Current (XIC) of unpurified peptide

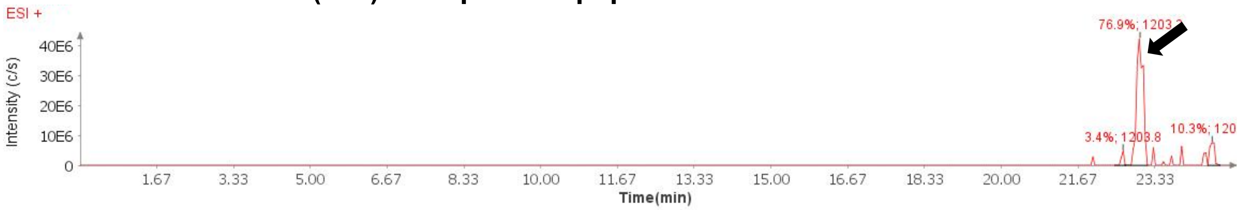

##### Mass Spectrum (ESI) of the expected product (unpurified sample)

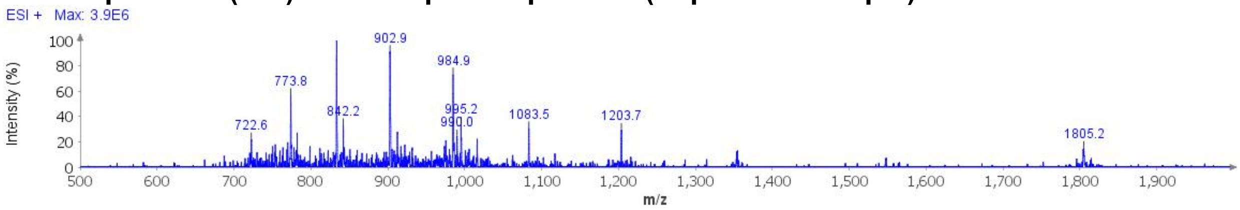

**Deconvoluted Mass Spectrum (ESI) of the expected product (unpurified sample)**

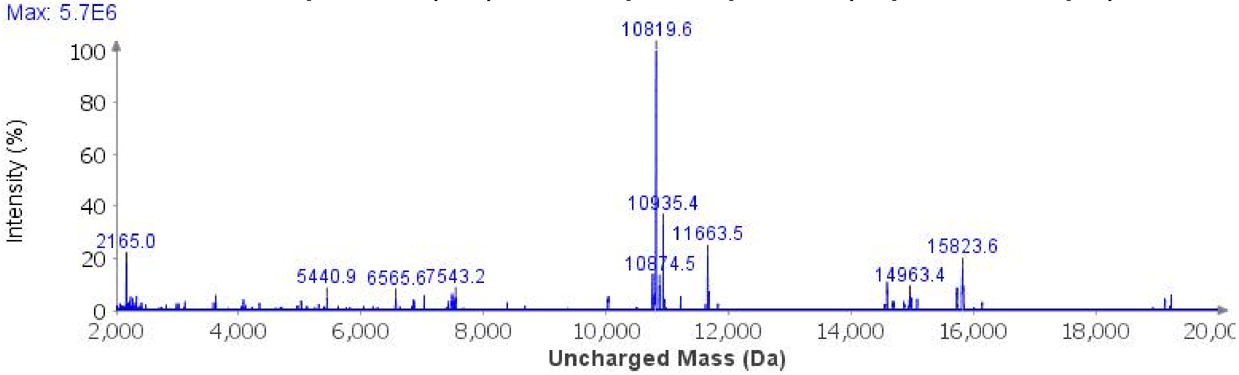

**Sample Displacement Mode Chromatography**:[6] The material was further purified by sample displacement mode over a Zorbax C18 column (4.6 x 250 mm, 5 μm, 300 Å, 4.7 mL/min, 40 °C) 5%-45% MeCN/$H_2O$/0.05%TFA over 60 min.

**Sample Displacement Mode trace (214 nm)**

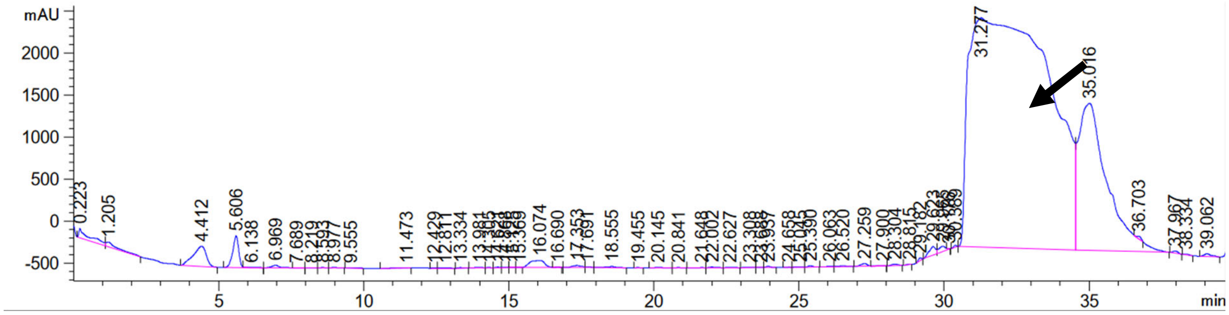

**Analytical LCMS**: Measured on an Agilent EC-C18 Poroshell column (4.6 x 100 mm, 4 μm, 100 Å, 40 °C, 1 mL/min) 5%-65% MeCN/$H_2O$/0.05% TFA over 60 min.

**Chemical Formula**: $C_{469}H_{768}N_{142}O_{147}S_2$
**Molecular Weight**: 10812.2700 g/mol

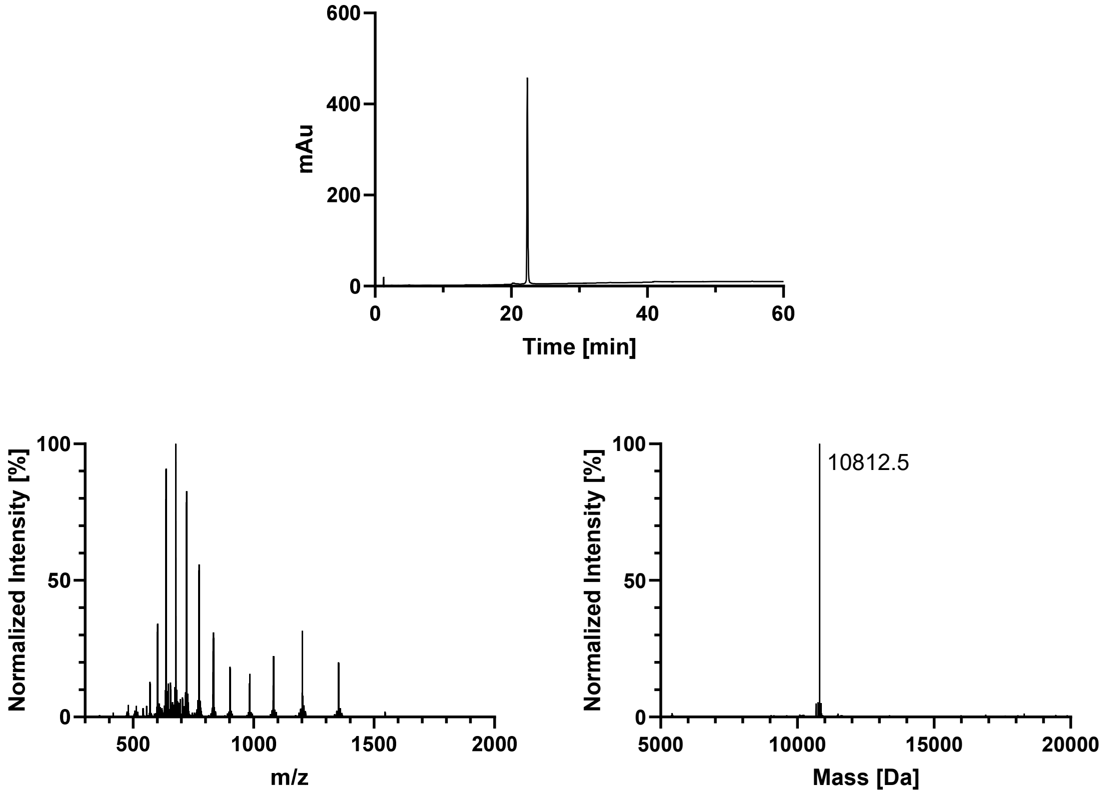
600
400
200
0
mAu
0
20
40
60
Time [min]
Normalized Intensity [%]
100
50
0
500
1000
1500
2000
m/z
Normalized Intensity [%]
100
50
0
10812.5
5000
10000
15000
20000
Mass [Da]

#### AcK311

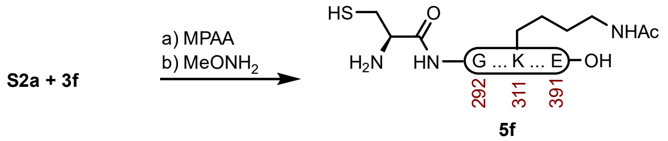

**Peptide 5f.** A 1-dram vial with **S2a** (5.6 mg, 0.741 μmol) and **3f** (4.0 mg, 1.15 μmol) was dissolved in fresh NCL buffer (370 μL, 200 mM MPAA, 6 M GndHCl, 0.2 M $Na_2HPO_4$, 20 mM TCEP HCl, pH 7.0) that had been sparged with $N_2$. The reaction was mixed at 23 ºC for 5 h before adding Thz removal buffer (370 μL, 400 mM $MeONH_2$ HCl, 6 M GndHCl, 0.2 M $NaH_2PO_4$, 20 mM TCEP HCl, pH 2.4), and adjusted to pH 4.0. The reaction was mixed at 23 ºC for 3 h before it was purified over a Luna PhenylHexyl column (10 x 250 mm, 5 μm, 100 Å, 4.7 mL/min) 15%-40% MeCN/$H_2O$/0.1% TFA over 30 min. The fractions were lyophilized to afford **5f** (5.0 mg, 62% yield).

##### Prep HPLC trace of NCL-Thz removal

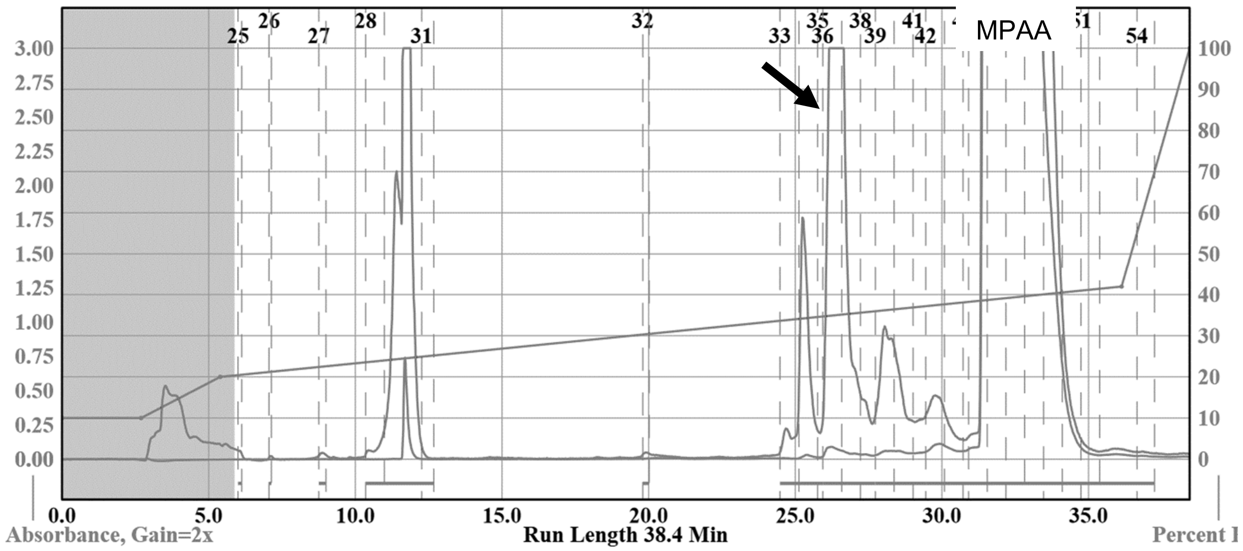

**Sample Displacement Mode Chromatography**:[6] The material was further purified by sample displacement mode over a Zorbax C18 column (4.6 x 250 mm, 5 μm, 300 Å, 4.7 mL/min, 40 °C) 5%-45% MeCN/$H_2O$/0.05%TFA over 60 min.

##### Sample Displacement Mode trace (214 nm)

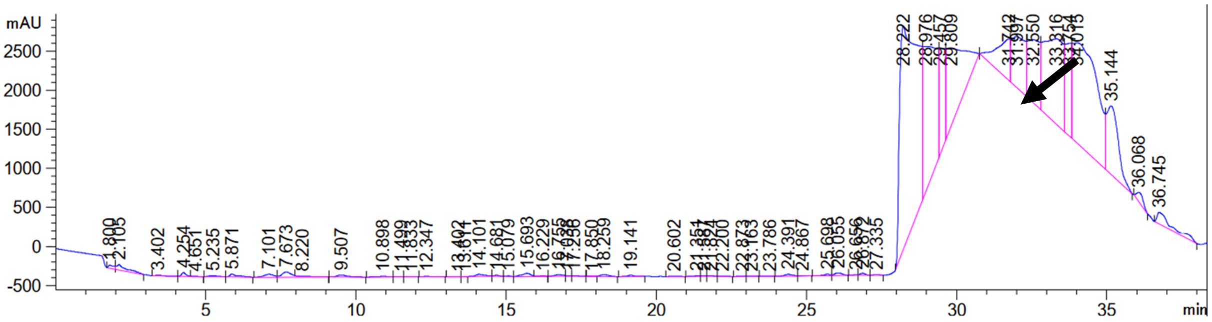

**Analytical LCMS**: Measured on an Agilent EC-C18 Poroshell column (4.6 x 100 mm, 4 µm, 100 Å, 40 °C, 1 mL/min) 5%-65% MeCN/$H_2O$/0.05% TFA over 60 min.

**Chemical Formula**: $C_{469}H_{768}N_{142}O_{147}S_2$
**Molecular Weight**: 10812.2700 g/mol

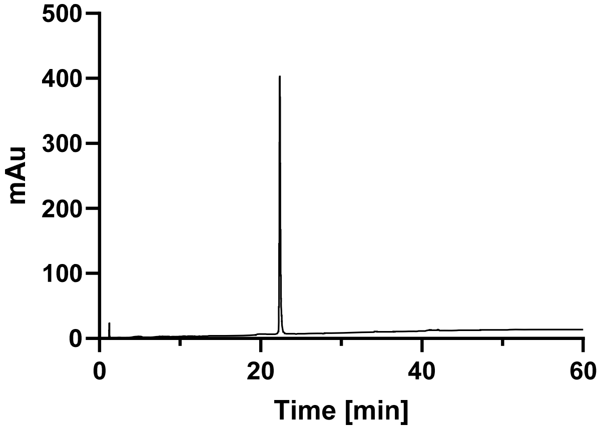

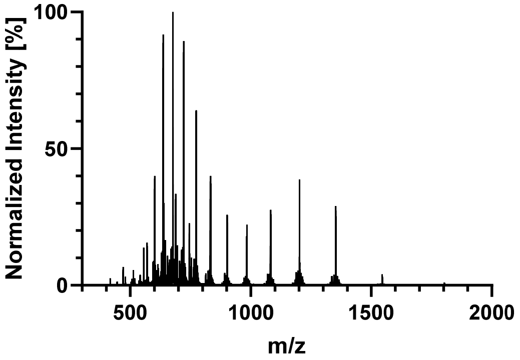

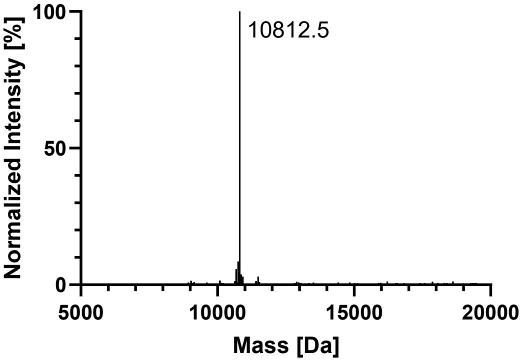

## pS356

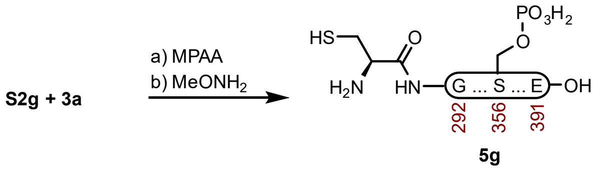

**Peptide 5g.** A 2-dram vial with **S2g** (16.4 mg, 2.15 µmol) and **3a** (12.2 mg, 3.70 µmol) was dissolved in fresh NCL buffer (717 µL, 200 mM MPAA, 6 M GndHCl, 0.2 M $Na_2HPO_4$, 20 mM TCEP HCl, pH 7.0) that had been sparged with $N_2$. The reaction was mixed at 23 ºC for 4 h before adding Thz removal buffer (717 µL, 400 mM $MeONH_2$ HCl, 6 M GndHCl, 0.2 M $NaH_2PO_4$, 20 mM TCEP HCl, pH 2.3), and adjusted to pH 4.0. The reaction was mixed at 23 ºC for 4 h before it was purified over a Luna PhenylHexyl column (10 x 250 mm, 5 µm, 100 Å, 4.7 mL/min) 20%-40% MeCN/$H_2O$/0.1% TFA over 30 min. The fractions were lyophilized to afford **5g** (9.7 mg, 42% yield).

##### Prep HPLC trace of NCL-Thz removal

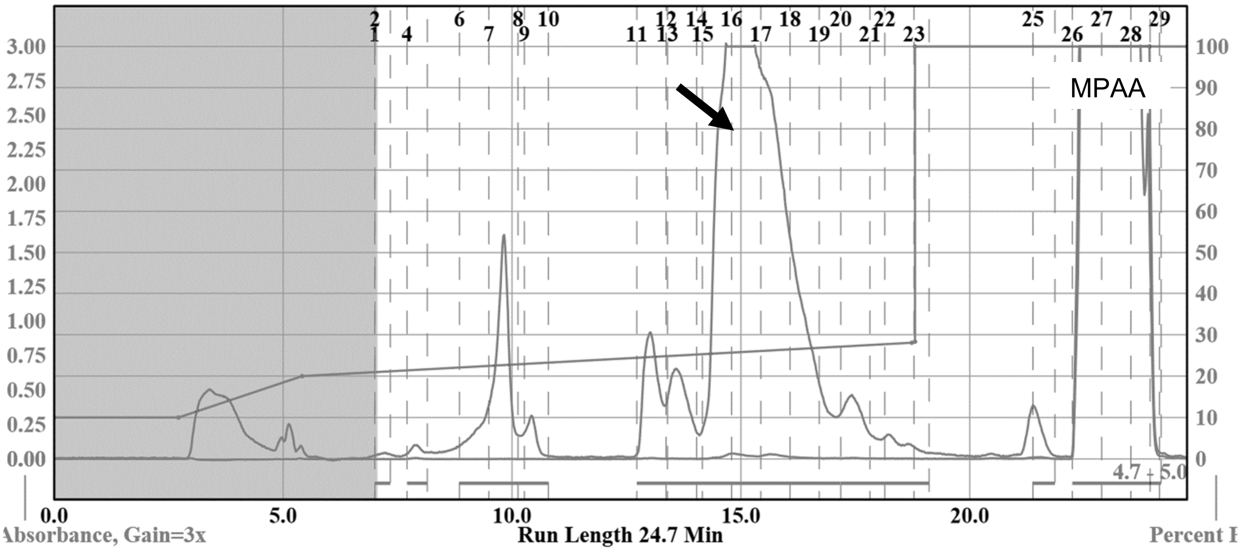

**Sample Displacement Mode Chromatography**:[6] The material was further purified by sample displacement mode over a Zorbax C18 column (4.6 x 250 mm, 5 µm, 300 Å, 4.7 mL/min, 40 °C) 5%-45% MeCN/$H_2O$/0.05%TFA over 60 min.

##### Sample Displacement Mode trace (214 nm)

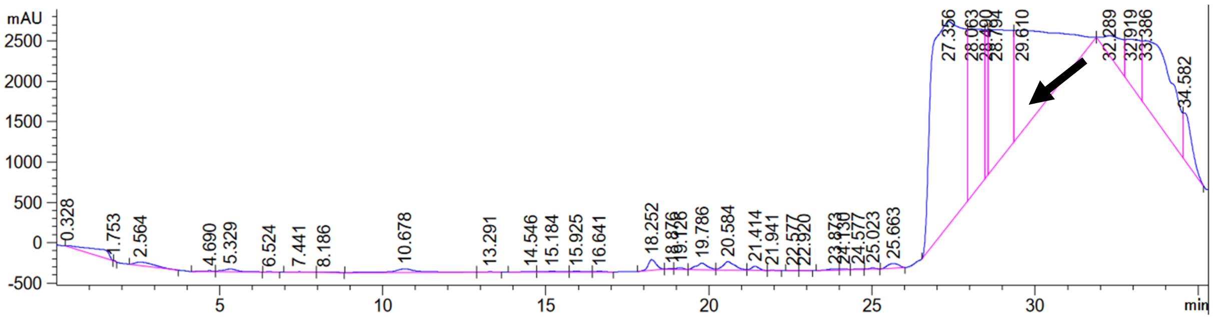

**Analytical LCMS**: Measured on an Agilent EC-C18 Poroshell column (4.6 x 100 mm, 4 µm, 100 Å, 40 °C, 1 mL/min) 5%-65% MeCN/$H_2O$/0.05% TFA over 60 min.

**Chemical Formula**: $C_{467}H_{767}N_{142}O_{149}PS_2$
**Molecular Weight**: 10850.2118 g/mol

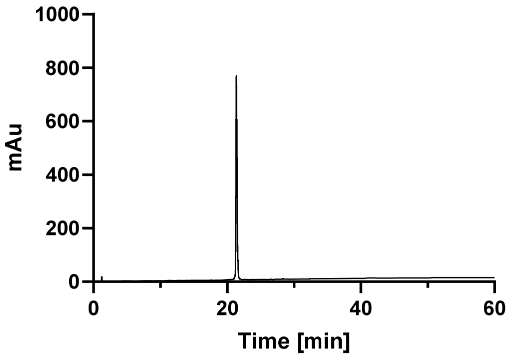

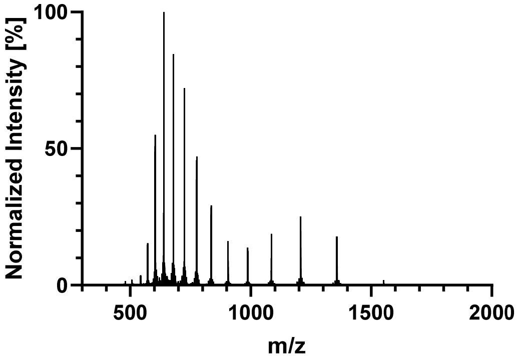

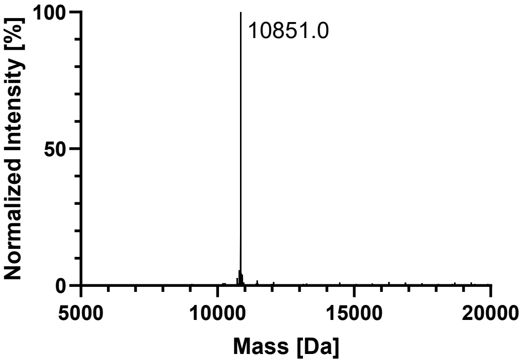

*pS352*

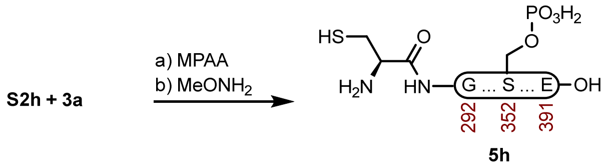

**<u>Peptide 5h.</u>** A 2-dram vial with **S2h** (14.1 mg, 1.85 μmol) and **3a** (10.1 mg, 3.06 μmol) was dissolved in fresh NCL buffer (617 μL, 200 mM MPAA, 6 M GndHCl, 0.2 M $Na_2HPO_4$, 20 mM TCEP HCl, pH 7.0) that had been sparged with $N_2$. The reaction was mixed at 23 ºC for 4 h before adding Thz removal buffer (617 μL, 400 mM $MeONH_2$ HCl, 6 M GndHCl, 0.2 M $NaH_2PO_4$, 20 mM TCEP HCl, pH 2.3), and adjusted to pH 4.0. The reaction was mixed at 23 ºC for 4 h before it was purified over a Luna PhenylHexyl column (10 x 250 mm, 5 μm, 100 Å, 4.7 mL/min) 15%-35% $MeCN/H_2O$/0.1% TFA over 30 min. The fractions were lyophilized to afford **5h** (7.5 mg, 37% yield).

**Crude LCMS (NCL)**: Measured on an Agilent EC-C18 Poroshell column (4.6 x 100 mm, 4 μm, 100 Å, 40 °C, 1 mL/min) 5%-65% $MeCN/H_2O$/0.05% TFA over 60 min.

**LC trace (214 nm) of unpurified peptide**

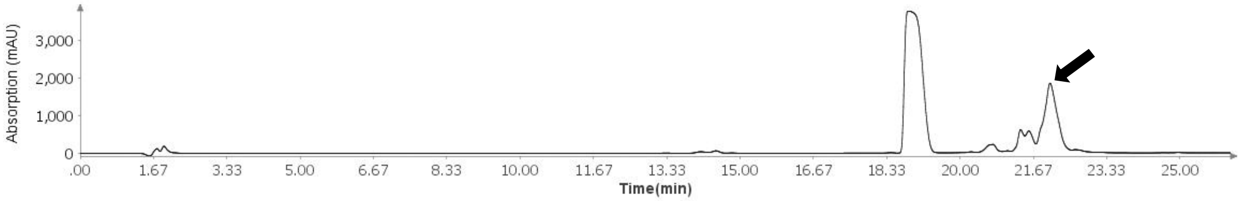

**Total Ion Current (TIC) of unpurified peptide**

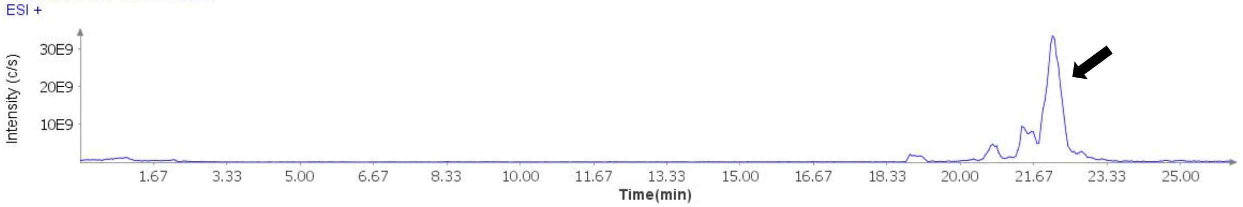

**Extracted Ion Current (XIC) of unpurified peptide**

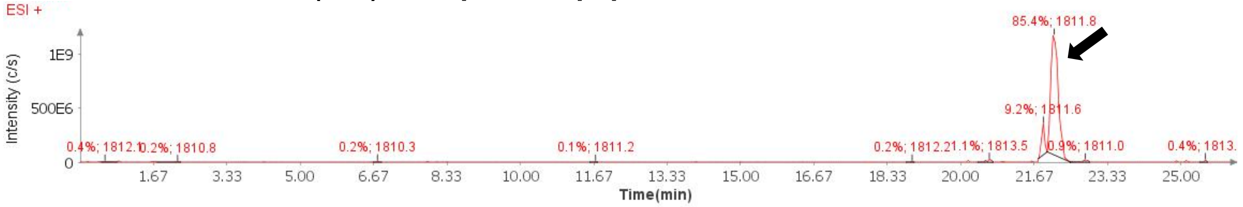

**Mass Spectrum (ESI) of the expected product (unpurified sample)**

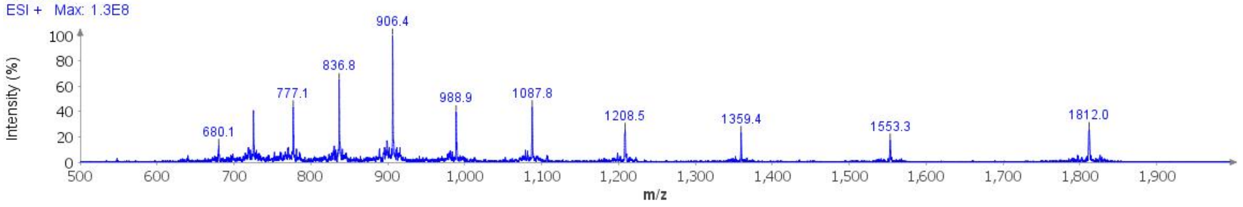

**Deconvoluted Mass Spectrum (ESI) of the expected product (unpurified sample)**

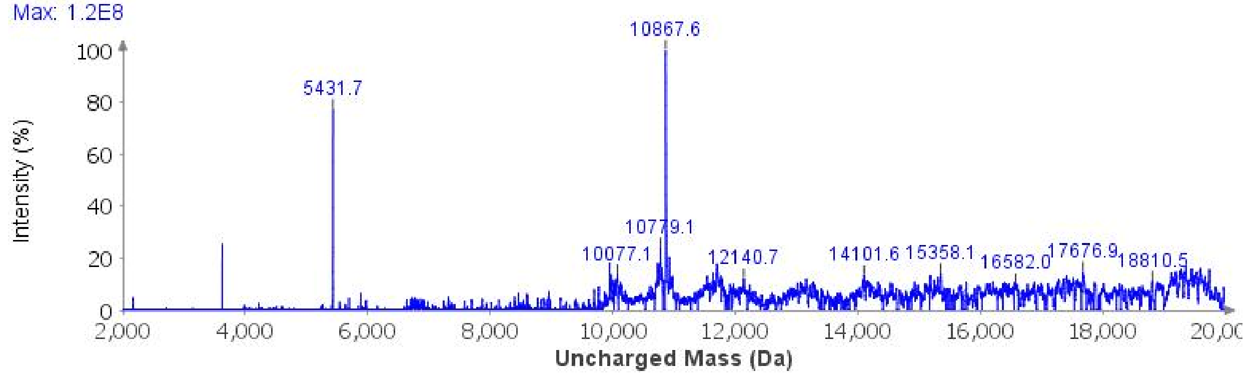

**Sample Displacement Mode Chromatography**:[6] The material was further purified by sample displacement mode over a Zorbax C18 column (4.6 x 250 mm, 5 µm, 300 Å, 4.7 mL/min, 40 °C) 5%-45% MeCN/$H_2O$/0.05%TFA over 60 min.

**Sample Displacement Mode trace (214 nm)**

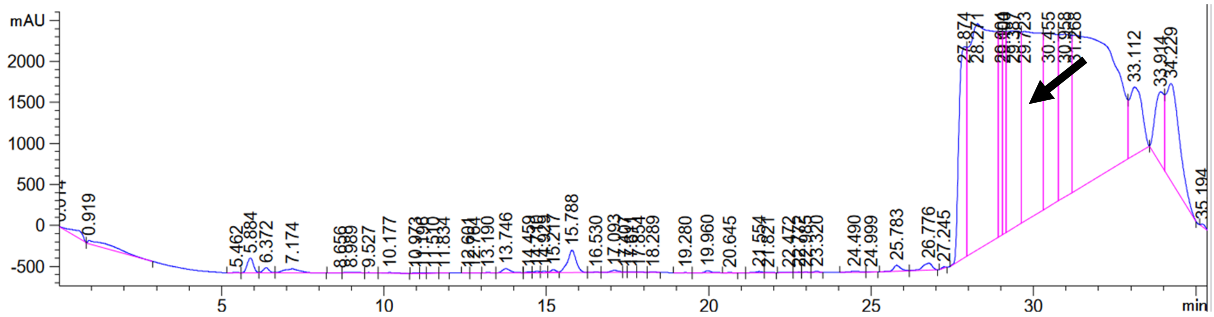

**Analytical LCMS**: Measured on an Agilent EC-C18 Poroshell column (4.6 x 100 mm, 4 µm, 100 Å, 40 °C, 1 mL/min) 5%-65% MeCN/$H_2O$/0.05% TFA over 60 min.

**Chemical Formula**: $C_{467}H_{767}N_{142}O_{149}PS_2$
**Molecular Weight**: 10850.2118 g/mol

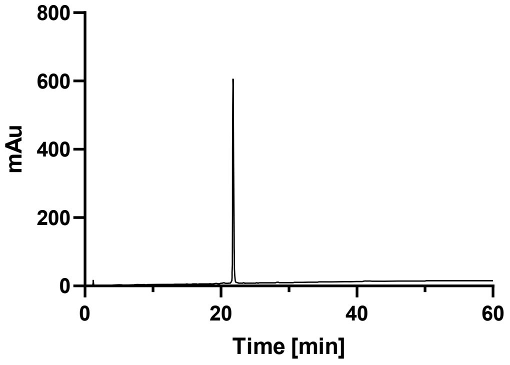

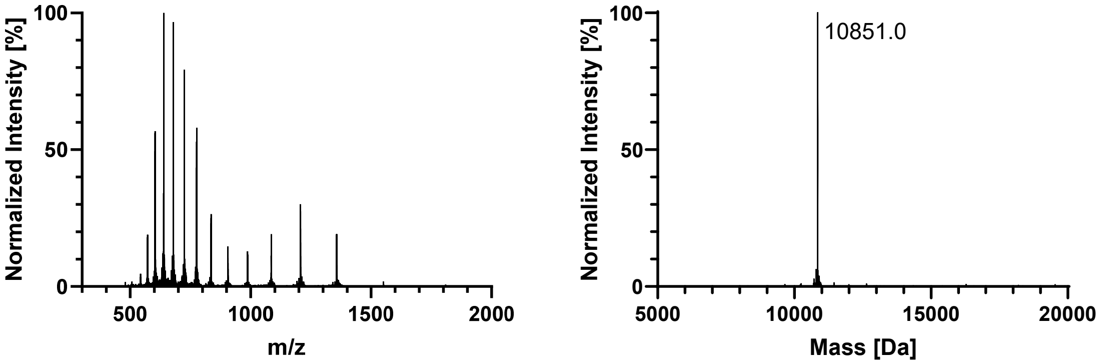
Normalized Intensity [%]
100
50
0
500
1000
1500
2000
m/z
Normalized Intensity [%]
100
50
0
10851.0
5000
10000
15000
20000
Mass [Da]

## pY310

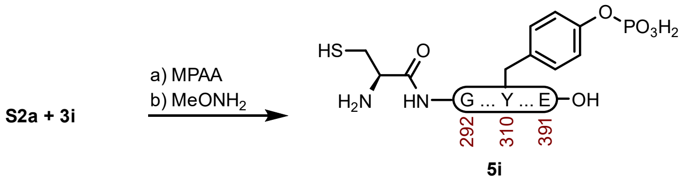

**Peptide 5i.** A 2-dram vial with **S2a** (8.4 mg, 1.11 μmol) and **3i** (3.8 mg, 1.08 μmol) was dissolved in fresh NCL buffer (500 μL, 200 mM MPAA, 6 M GndHCl, 0.2 M $Na_2HPO_4$, 20 mM TCEP HCl, pH 7.1) that had been sparged with $N_2$. The reaction was mixed at 23 ºC for 3 h before adding Thz removal buffer (500 μL, 400 mM $MeONH_2$ HCl, 6 M GndHCl, 0.2 M $NaH_2PO_4$, 20 mM TCEP HCl, pH 2.5), and adjusted to pH 3.9. The reaction was mixed at 23 ºC for 2 h before it was purified over a Luna PhenylHexyl column (10 x 250 mm, 5 μm, 100 Å, 4.7 mL/min) 20%-40% MeCN/$H_2O$/0.1% TFA over 30 min. The fractions were lyophilized to afford **5i** (5.1 mg, 42% yield).

**Prep HPLC trace of NCL-Thz removal**

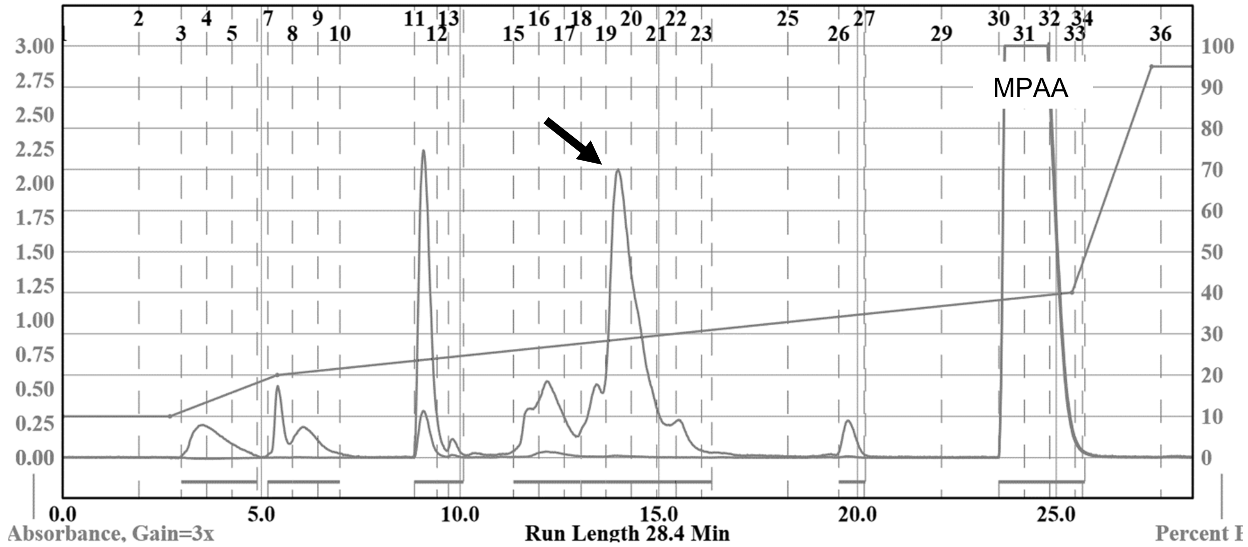

**Sample Displacement Mode Chromatography**:[6] The material was further purified by sample displacement mode over a Zorbax C18 column (4.6 x 250 mm, 5 μm, 300 Å, 4.7 mL/min, 40 °C) 5%-45% MeCN/$H_2O$/0.05%TFA over 60 min.

**Sample Displacement Mode trace (214 nm)**

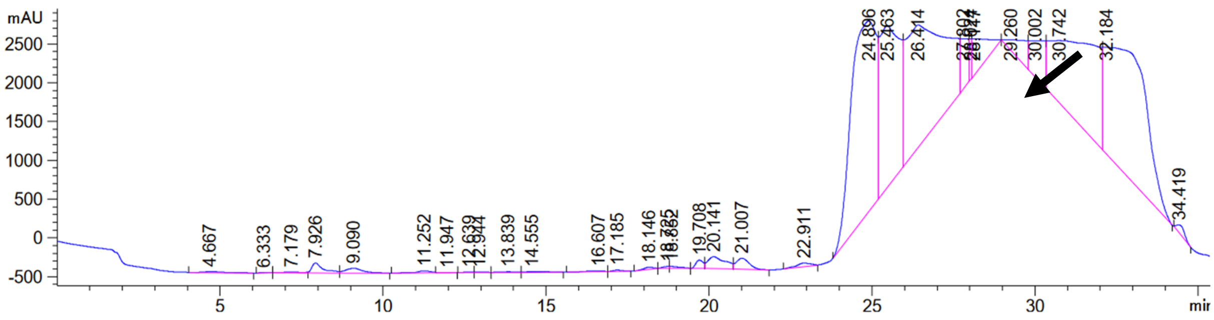

**Analytical LCMS**: Measured on an Agilent EC-C18 Poroshell column (4.6 x 100 mm, 4 µm, 100 Å, 40 °C, 1 mL/min) 5%-65% MeCN/$H_2O$/0.05% TFA over 60 min.

**Chemical Formula**: $C_{467}H_{767}N_{142}O_{149}PS_2$
**Molecular Weight**: 10850.2118 g/mol

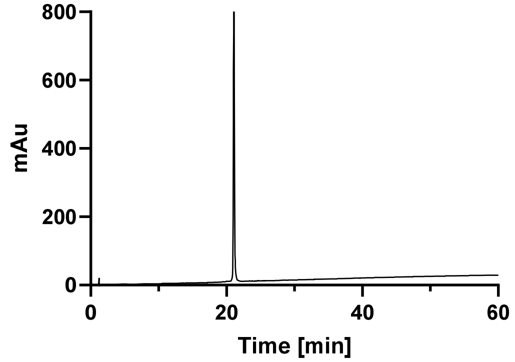

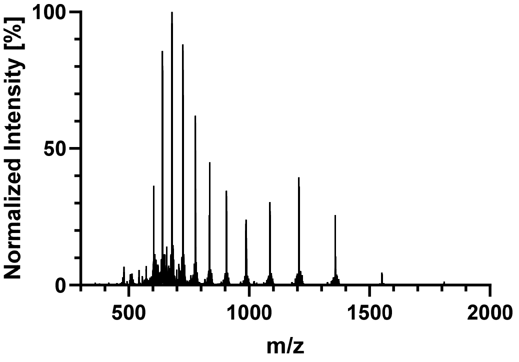

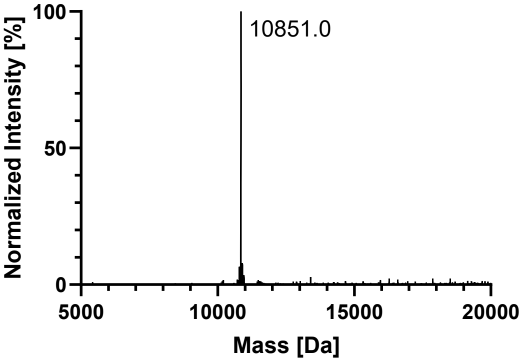

## pS305

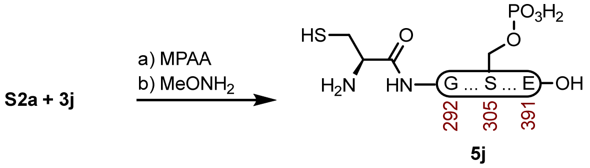

**Peptide 5j.** A 1-dram vial with **S2a** (5.0 mg, 0.662 μmol) and **3j** (3.2 mg, 0.913 μmol) was dissolved in fresh NCL buffer (331 μL, 200 mM MPAA, 6 M GndHCl, 0.2 M $Na_2HPO_4$, 20 mM TCEP HCl, pH 7.2) that had been sparged with $N_2$. The reaction was mixed at 23 ºC for 3 h before adding Thz removal buffer (331 μL, 400 mM $MeONH_2$ HCl, 6 M GndHCl, 0.2 M $NaH_2PO_4$, 20 mM TCEP HCl, pH 2.4), and adjusted to pH 4.0. The reaction was mixed at 23 ºC for 6 h before it was purified over a Luna PhenylHexyl column (10 x 250 mm, 5 μm, 100 Å, 4.7 mL/min) 15%-40% MeCN/$H_2O$/0.1% TFA over 30 min. The fractions were lyophilized to afford **5j** (4.1 mg, 57% yield).

**Crude LCMS (NCL)**: Measured on an Agilent EC-C18 Poroshell column (4.6 x 100 mm, 4 μm, 100 Å, 40 °C, 1 mL/min) 5%-65% MeCN/$H_2O$/0.05% TFA over 60 min.

##### LC trace (214 nm) of unpurified peptide

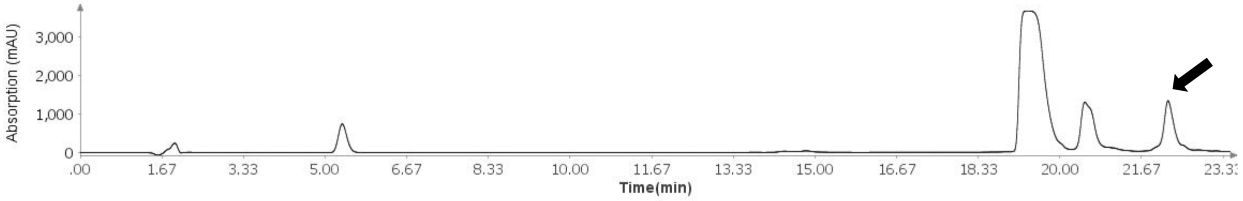

##### Total Ion Current (TIC) of unpurified peptide

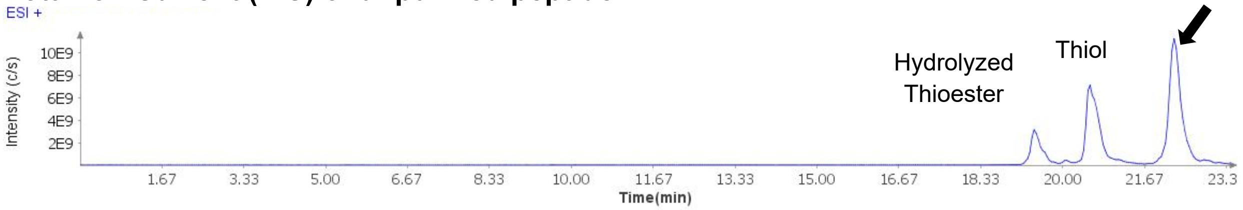

##### Extracted Ion Current (XIC) of unpurified peptide

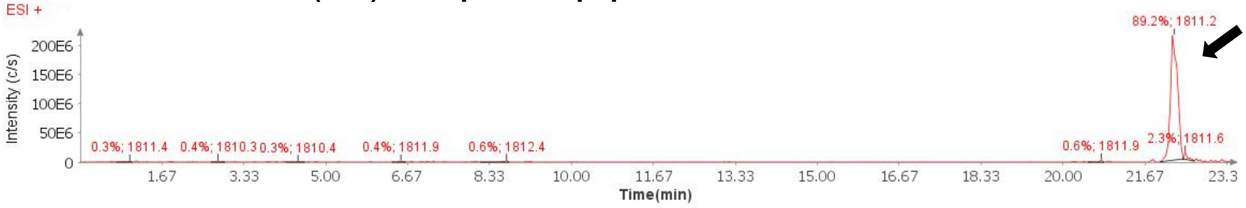

##### Mass Spectrum (ESI) of the expected product (unpurified sample)

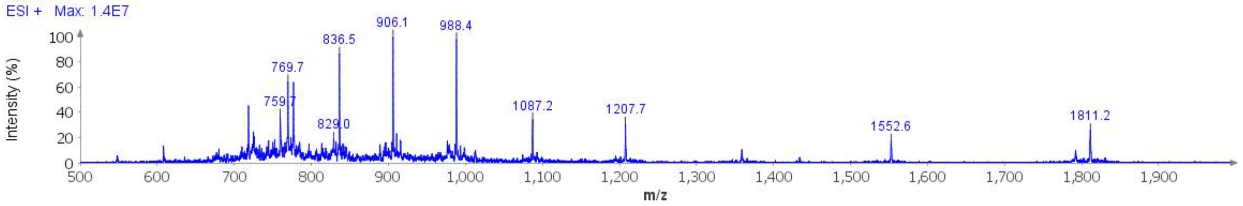

**Deconvoluted Mass Spectrum (ESI) of the expected product (unpurified sample)**

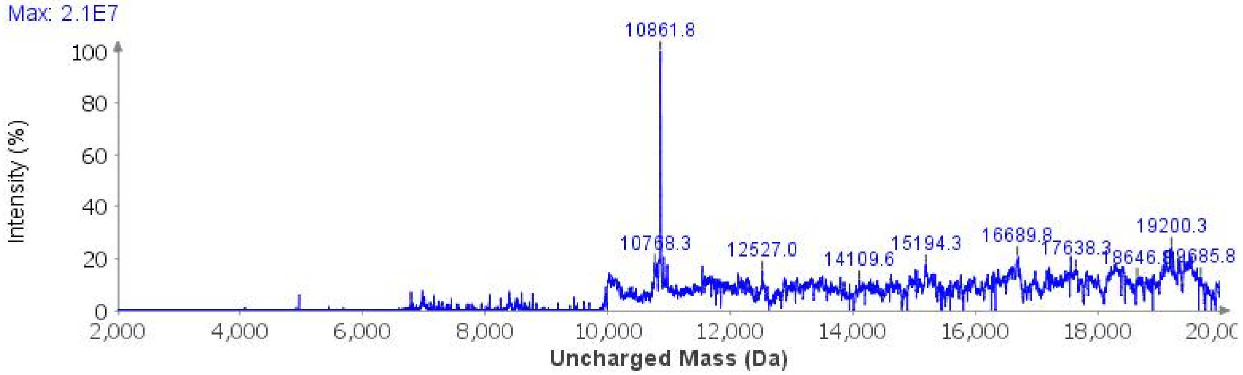

**Sample Displacement Mode Chromatography**:[6] The material was further purified by sample displacement mode over a Zorbax C18 column (4.6 x 250 mm, 5 µm, 300 Å, 4.7 mL/min, 40 °C) 5%-45% MeCN/$H_2O$/0.05%TFA over 60 min.

**Sample Displacement Mode trace (214 nm)**

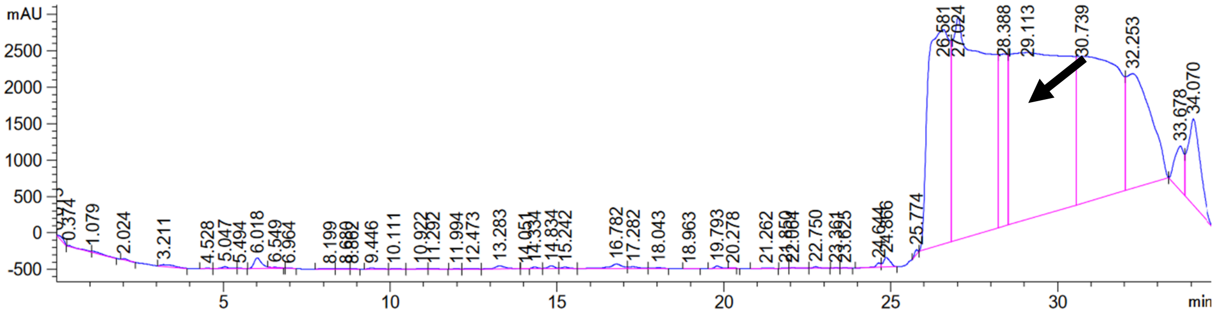

**Analytical LCMS**: Measured on an Agilent EC-C18 Poroshell column (4.6 x 100 mm, 4 µm, 100 Å, 40 °C, 1 mL/min) 5%-65% MeCN/$H_2O$/0.05% TFA over 60 min.

**Chemical Formula**: $C_{467}H_{767}N_{142}O_{149}PS_2$
**Molecular Weight**: 10850.2118 g/mol

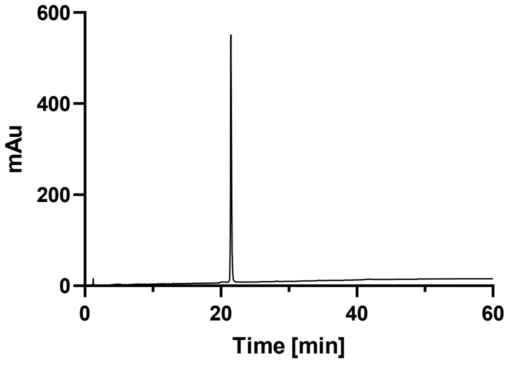

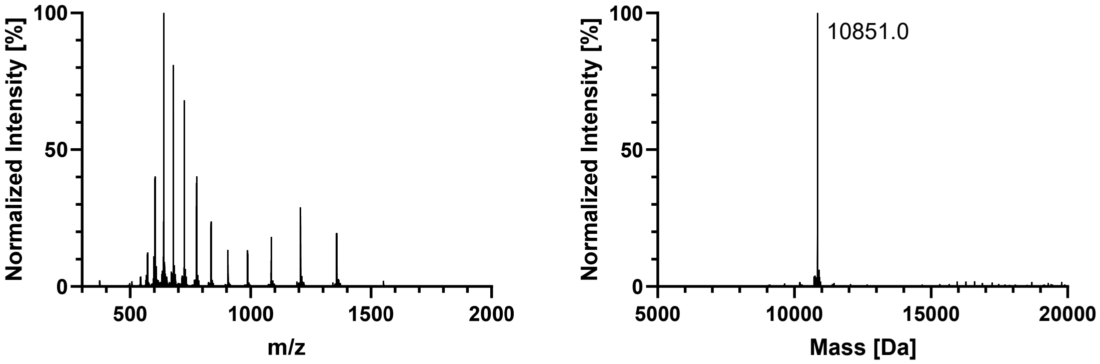
Normalized Intensity [%]
100
50
0
500
1000
1500
2000
m/z
Normalized Intensity [%]
100
50
0
10851.0
5000
10000
15000
20000
Mass [Da]

#### GlcNAcN359

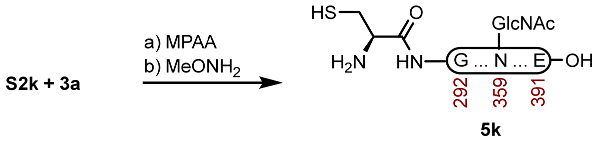

**Peptide 5k.** A 1-dram vial with **S2k** (9.8 mg, 1.26 µmol) and **3a** (6.8 mg, 2.06 µmol) was dissolved in fresh NCL buffer (630 µL, 200 mM MPAA, 6 M GndHCl, 0.2 M $Na_2HPO_4$, 20 mM TCEP HCl, pH 7.2) that had been sparged with $N_2$. The reaction was mixed at 23 ºC for 3 h before adding Thz removal buffer (630 µL, 400 mM $MeONH_2$ HCl, 6 M GndHCl, 0.2 M $NaH_2PO_4$, 20 mM TCEP HCl, pH 2.5), and adjusted to pH 4.0. The reaction was mixed at 23 ºC for 3 h before it was purified over a Luna PhenylHexyl column (10 x 250 mm, 5 µm, 100 Å, 4.7 mL/min) 20%-40% MeCN/$H_2O$/0.1% TFA over 30 min. The fractions were lyophilized to afford **5k** (10.4 mg, 75% yield).

**Crude LCMS (NCL)**: Measured on an Agilent EC-C18 Poroshell column (4.6 x 100 mm, 4 µm, 100 Å, 40 °C, 1 mL/min) 5%-65% MeCN/$H_2O$/0.05% TFA over 60 min.

##### LC trace (214 nm) of unpurified peptide

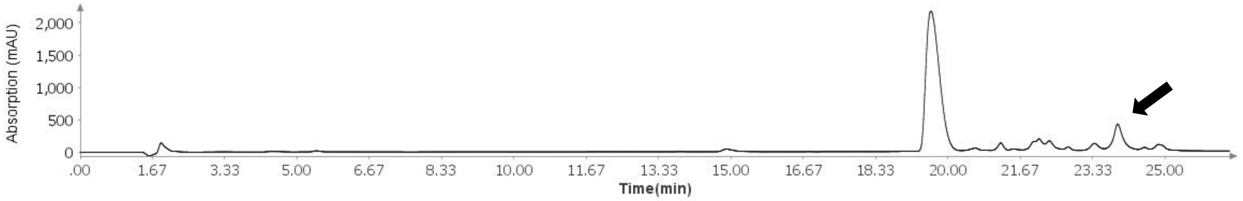

##### Total Ion Current (TIC) of unpurified peptide

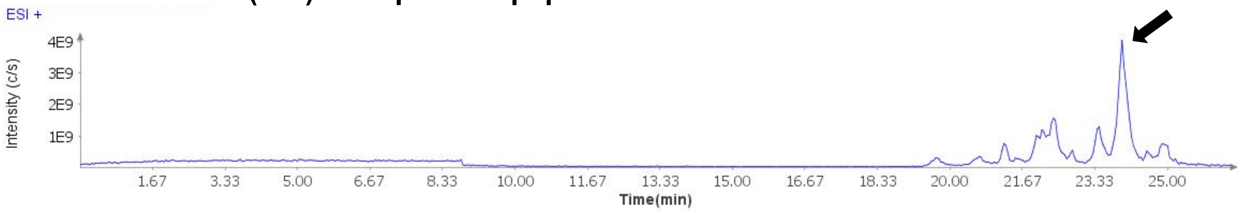

##### Extracted Ion Current (XIC) of unpurified peptide

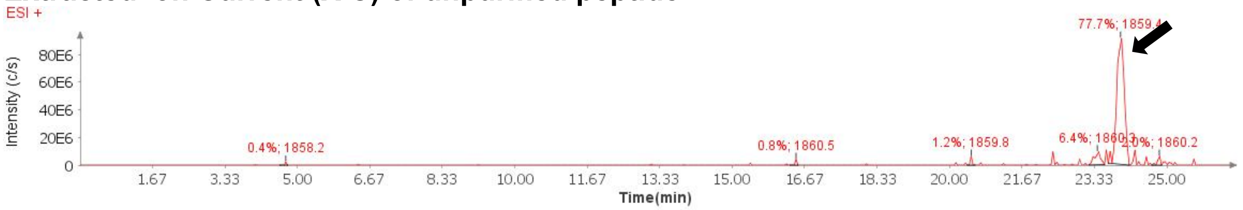

##### Mass Spectrum (ESI) of the expected product (unpurified sample)

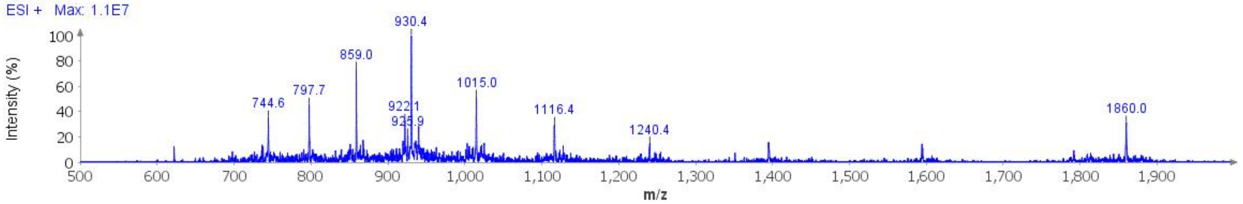

**Deconvoluted Mass Spectrum (ESI) of the expected product (unpurified sample)**

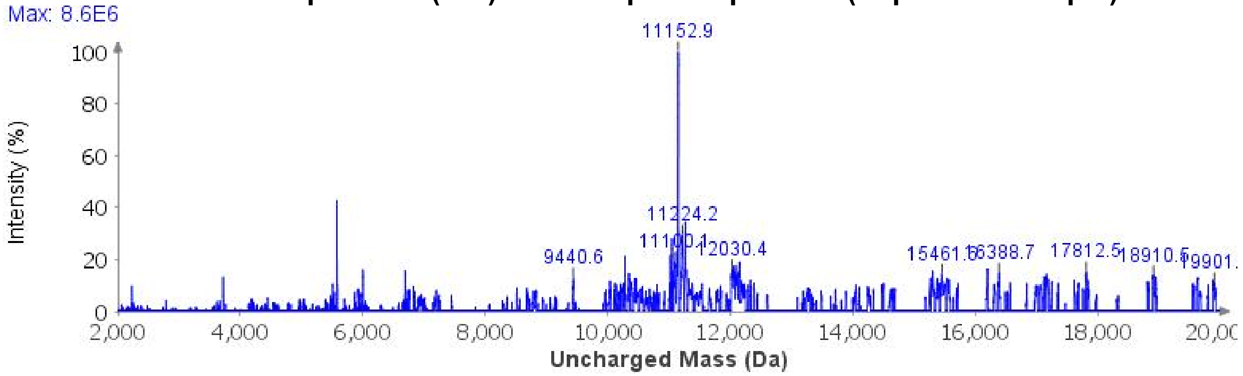

**Sample Displacement Mode Chromatography**:[6] The material was further purified by sample displacement mode over a Zorbax C18 column (4.6 x 250 mm, 5 µm, 300 Å, 4.7 mL/min, 40 °C) 5%-45% MeCN/$H_2O$/0.05%TFA over 60 min.

**Sample Displacement Mode trace (214 nm)**

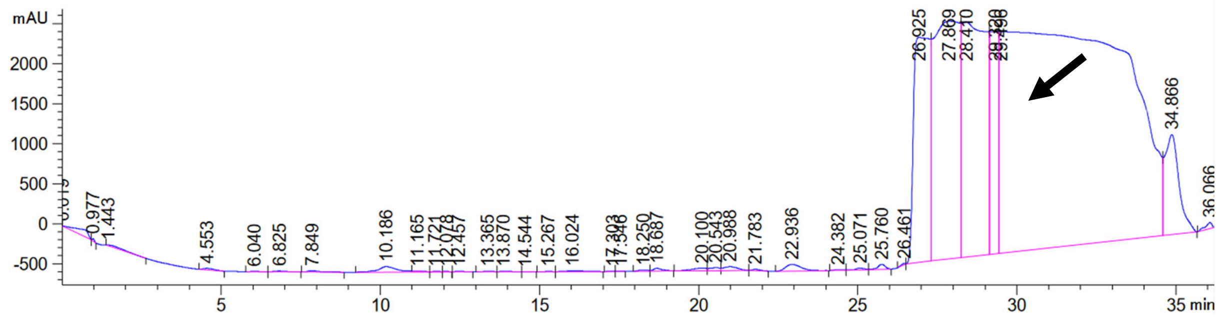

**Analytical LCMS**: Measured on an Agilent EC-C18 Poroshell column (4.6 x 100 mm, 4 µm, 100 Å, 40 °C, 1 mL/min) 5%-65% MeCN/$H_2O$/0.05% TFA over 60 min.

**Chemical Formula**: $C_{475}H_{779}N_{143}O_{151}S_2$
**Molecular Weight**: 10973.4270 g/mol

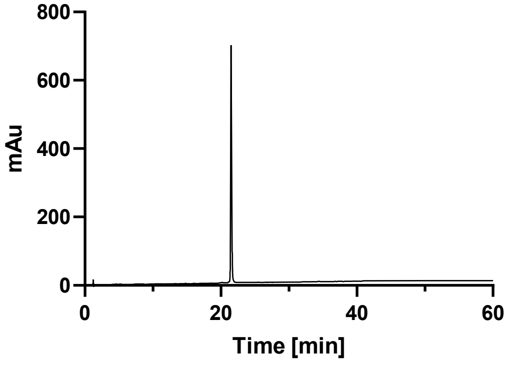

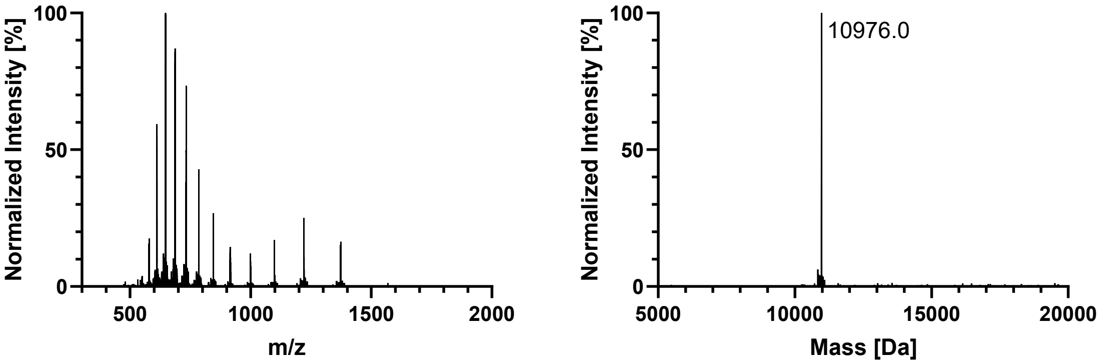
Normalized Intensity [%]
100
50
0
500
1000
1500
2000
m/z
Normalized Intensity [%]
100
50
0
10976.0
5000
10000
15000
20000
Mass [Da]

#### GlcNAcN359 + AcK353

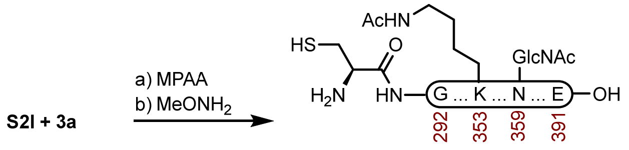

**5l**

**Peptide 5l.** A 2-dram vial with **S2l** (8.9 mg, 1.14 µmol) and **3a** (5.5 mg, 1.67 µmol) was dissolved in fresh NCL buffer (570 µL, 200 mM MPAA, 6 M GndHCl, 0.2 M $Na_2HPO_4$, 20 mM TCEP HCl, pH 7.0) that had been sparged with $N_2$. The reaction was mixed at 23 ºC for 6 h before adding Thz removal buffer (570 µL, 400 mM $MeONH_2$ HCl, 6 M GndHCl, 0.2 M $NaH_2PO_4$, 20 mM TCEP HCl, pH 2.7), and adjusted to pH 4.0. The reaction was mixed at 23 ºC for 4 h before it was purified over a Luna PhenylHexyl column (10 x 250 mm, 5 µm, 100 Å, 4.7 mL/min) 15%-40% MeCN/$H_2O$/0.1% TFA over 30 min. The fractions were lyophilized to afford **5l** (6.9 mg, 55% yield).

**Crude LCMS (NCL)**: Measured on an Agilent EC-C18 Poroshell column (4.6 x 100 mm, 4 µm, 100 Å, 40 °C, 1 mL/min) 5%-65% MeCN/$H_2O$/0.05% TFA over 60 min.

##### LC trace (214 nm) of unpurified peptide

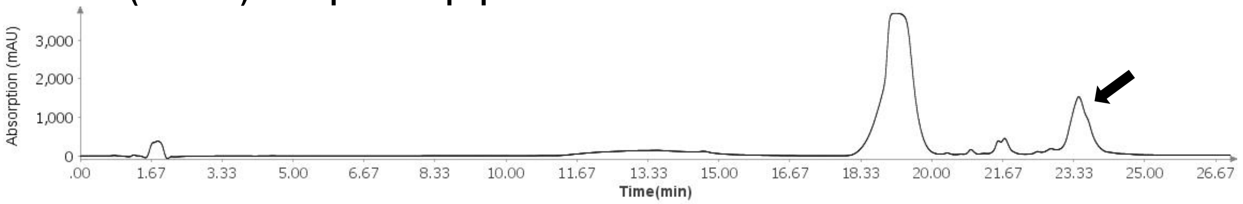

##### Total Ion Current (TIC) of unpurified peptide

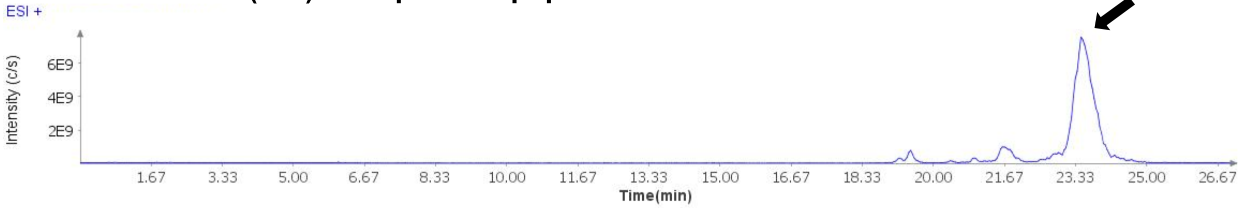

##### Extracted Ion Current (XIC) of unpurified peptide

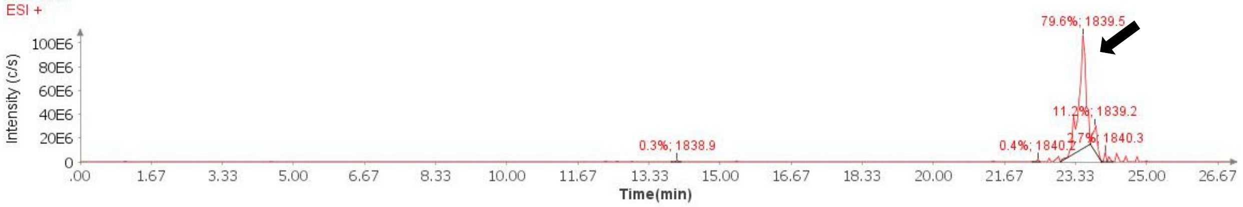

##### Mass Spectrum (ESI) of the expected product (unpurified sample)

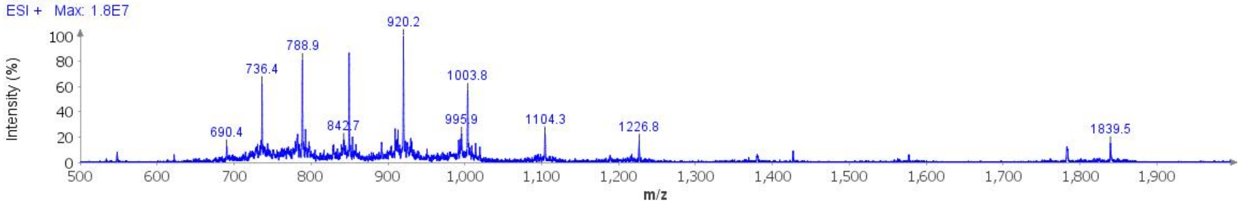

**Deconvoluted Mass Spectrum (ESI) of the expected product (unpurified sample)**

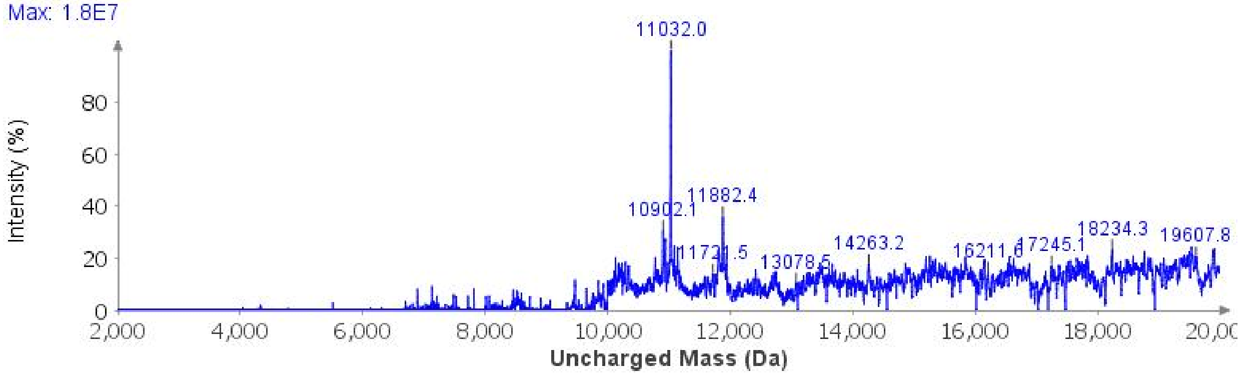

**Sample Displacement Mode Chromatography**:[6] The material was further purified by sample displacement mode over a Zorbax C18 column (4.6 x 250 mm, 5 µm, 300 Å, 4.7 mL/min, 40 °C) 5%-45% MeCN/$H_2O$/0.05%TFA over 60 min.

**Sample Displacement Mode trace (214 nm)**

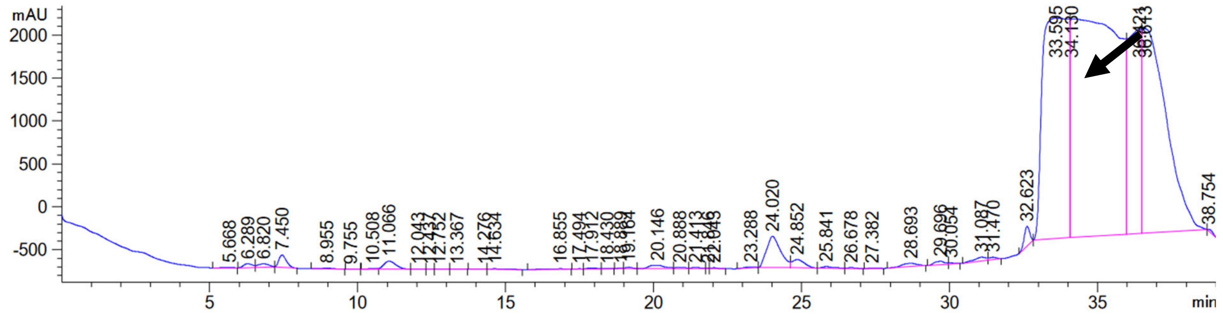

**Analytical LCMS**: Measured on an Agilent EC-C18 Poroshell column (4.6 x 100 mm, 4 µm, 100 Å, 40 °C, 1 mL/min) 5%-65% MeCN/$H_2O$/0.05% TFA over 60 min.

**Chemical Formula**: $C_{477}H_{781}N_{143}O_{152}S_2$
**Molecular Weight**: 11015.4640 g/mol

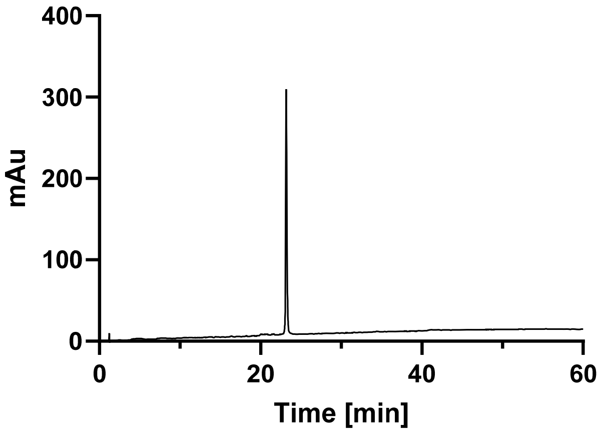
400
300
200
100
0
mAu
0
20
40
60
Time [min]

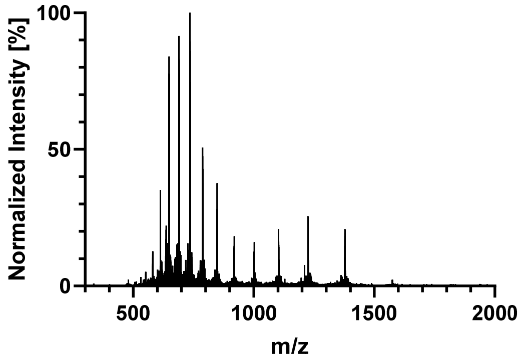
100
50
0
Normalized Intensity [%]
500
1000
1500
2000
m/z

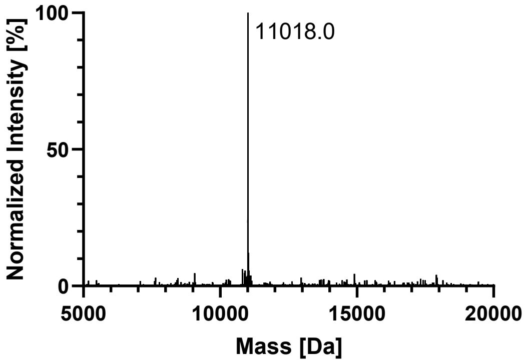
100
50
0
Normalized Intensity [%]
11018.0
5000
10000
15000
20000
Mass [Da]

#### GlcNAcN359 + pS305

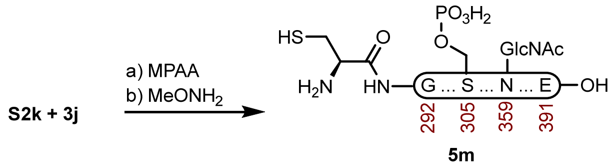

**Peptide 5m.** A 1-dram vial with **S2k** (9.1 mg, 1.22 µmol) and **3j** (6.5 mg, 1.85 µmol) was dissolved in fresh NCL buffer (750 µL, 200 mM MPAA, 6 M GndHCl, 0.2 M $Na_2HPO_4$, 20 mM TCEP HCl, pH 7.2) that had been sparged with $N_2$. The reaction was mixed at 23 ºC for 4 h before adding Thz removal buffer (750 µL, 400 mM $MeONH_2$ HCl, 6 M GndHCl, 0.2 M $NaH_2PO_4$, 20 mM TCEP HCl, pH 2.4), and adjusted to pH 3.9. The reaction was mixed at 23 ºC for 4 h before it was purified over a Luna PhenylHexyl column (10 x 250 mm, 5 µm, 100 Å, 4.7 mL/min) 15%-40% MeCN/$H_2O$/0.1% TFA over 30 min. The fractions were lyophilized to afford **5m** (5.0 mg, 37% yield).

**Crude LCMS (NCL)**: Measured on an Agilent EC-C18 Poroshell column (4.6 x 100 mm, 4 µm, 100 Å, 40 °C, 1 mL/min) 5%-65% MeCN/$H_2O$/0.05% TFA over 60 min.

##### LC trace (214 nm) of unpurified peptide

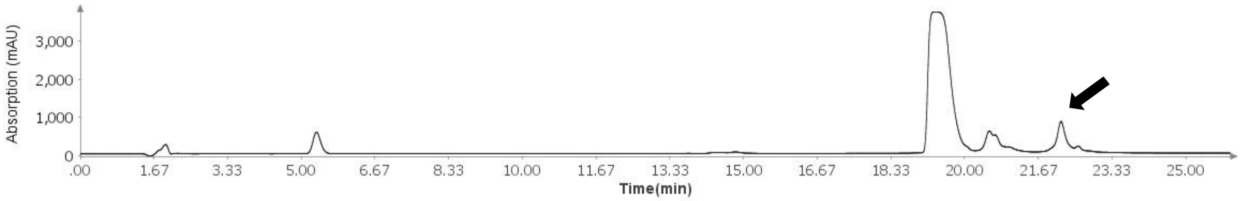

##### Total Ion Current (TIC) of unpurified peptide

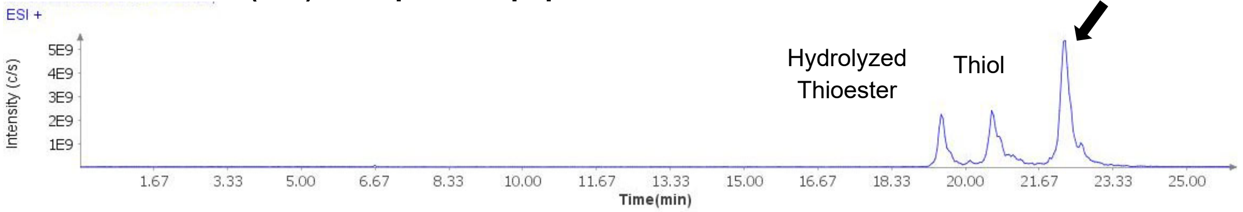

##### Extracted Ion Current (XIC) of unpurified peptide

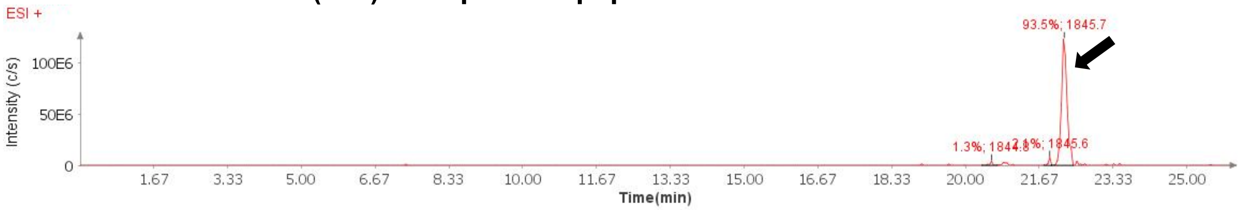

##### Mass Spectrum (ESI) of the expected product (unpurified sample)

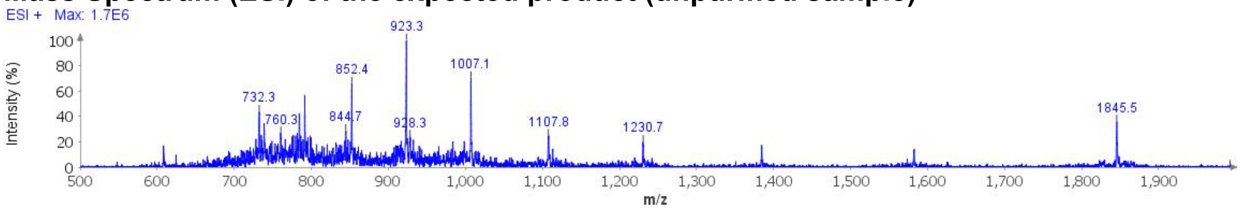

**Deconvoluted Mass Spectrum (ESI) of the expected product (unpurified sample)**

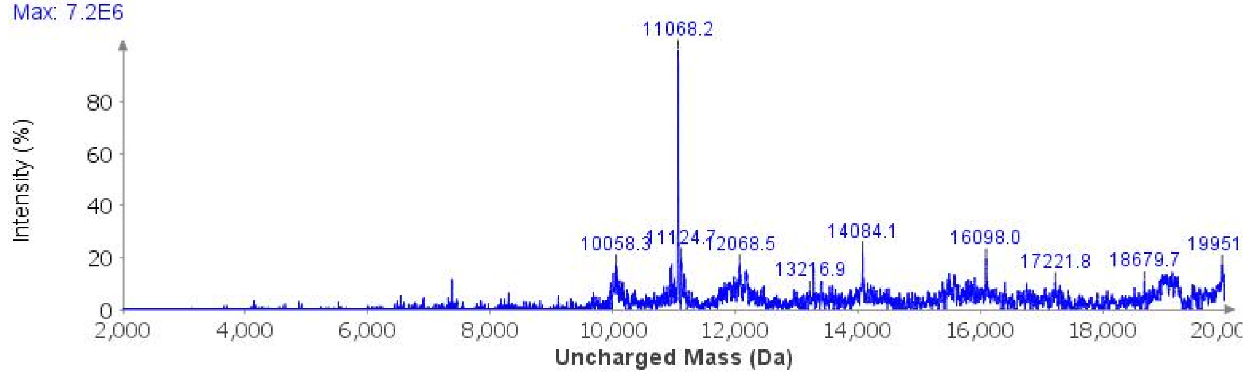

**Sample Displacement Mode Chromatography**:[6] The material was further purified by sample displacement mode over a Zorbax C18 column (4.6 x 250 mm, 5 μm, 300 Å, 4.7 mL/min, 40 °C) 5%-45% MeCN/$H_2O$/0.05%TFA over 60 min.

**Sample Displacement Mode trace (214 nm)**

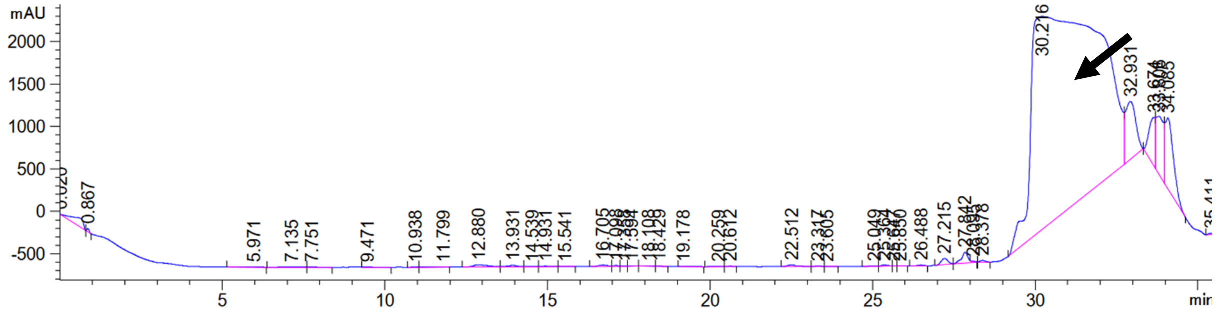

**Analytical LCMS**: Measured on an Agilent EC-C18 Poroshell column (4.6 x 100 mm, 4 μm, 100 Å, 40 °C, 1 mL/min) 5%-65% MeCN/$H_2O$/0.05% TFA over 60 min.

**Chemical Formula**: $C_{475}H_{780}N_{143}O_{154}PS_2$
**Molecular Weight**: 11053.4058 g/mol

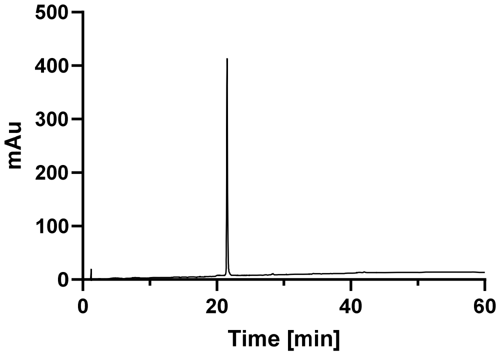

500
400
300
200
100
0
mAu
0
20
40
60
Time [min]

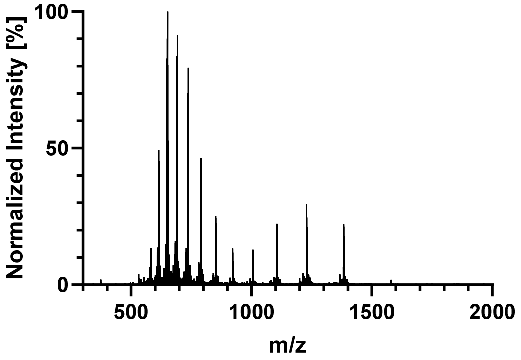

100
50
0
Normalized Intensity [%]
500
1000
1500
2000
m/z

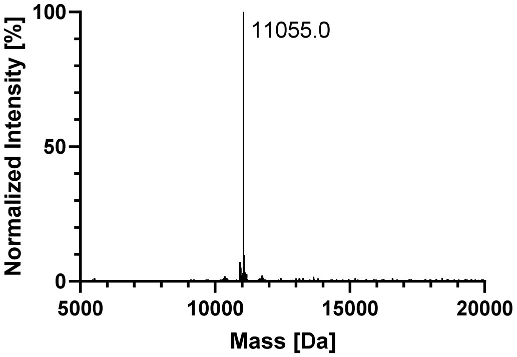

100
50
0
Normalized Intensity [%]
11055.0
5000
10000
15000
20000
Mass [Da]

#### 9. PROTEIN CONCENTRATION

Lyophilized Tau(291-391) was dissolved in the buffer, and the protein concentration was estimated at A280 on a Nanodrop with a calculated extinction coefficient of 1490 $M^{-1}cm^{-1}$. The tau protein stock solutions were typically 150-300 µM for 25 mM HEPES or ~1 mM for 10 mM sodium phosphate.

#### 10. GEL ELECTROPHORESIS

A 0.65 mL Eppendorf tube was charged with 3 µg of the sample, 4 µL NuPAGE LDS Sample Buffer (Invitrogen, Cat. No. NP0007), and 1 µL of NuPAGE Sample Reducing Agent (Invitrogen, Cat. No. NP0004). The samples were incubated at 70 ºC for 5 min and then loaded into a Novex 16% Tricine Gel (Invitrogen, Cat. No. EC66955BOX). The gel was run at 125 V for 90 min using Tris-Glycine SDS running buffer (Invitrogen, Cat. No. LC2675-4). The gel was destained with 200 mL of $H_2O$ (3 x 5 min) and then stained with 0.04% (w/v) colloidal Coomassie blue (G250) + 3.5% (w/v) perchloric acid for 2 h. The gel was destained with $H_2O$ and imaged.

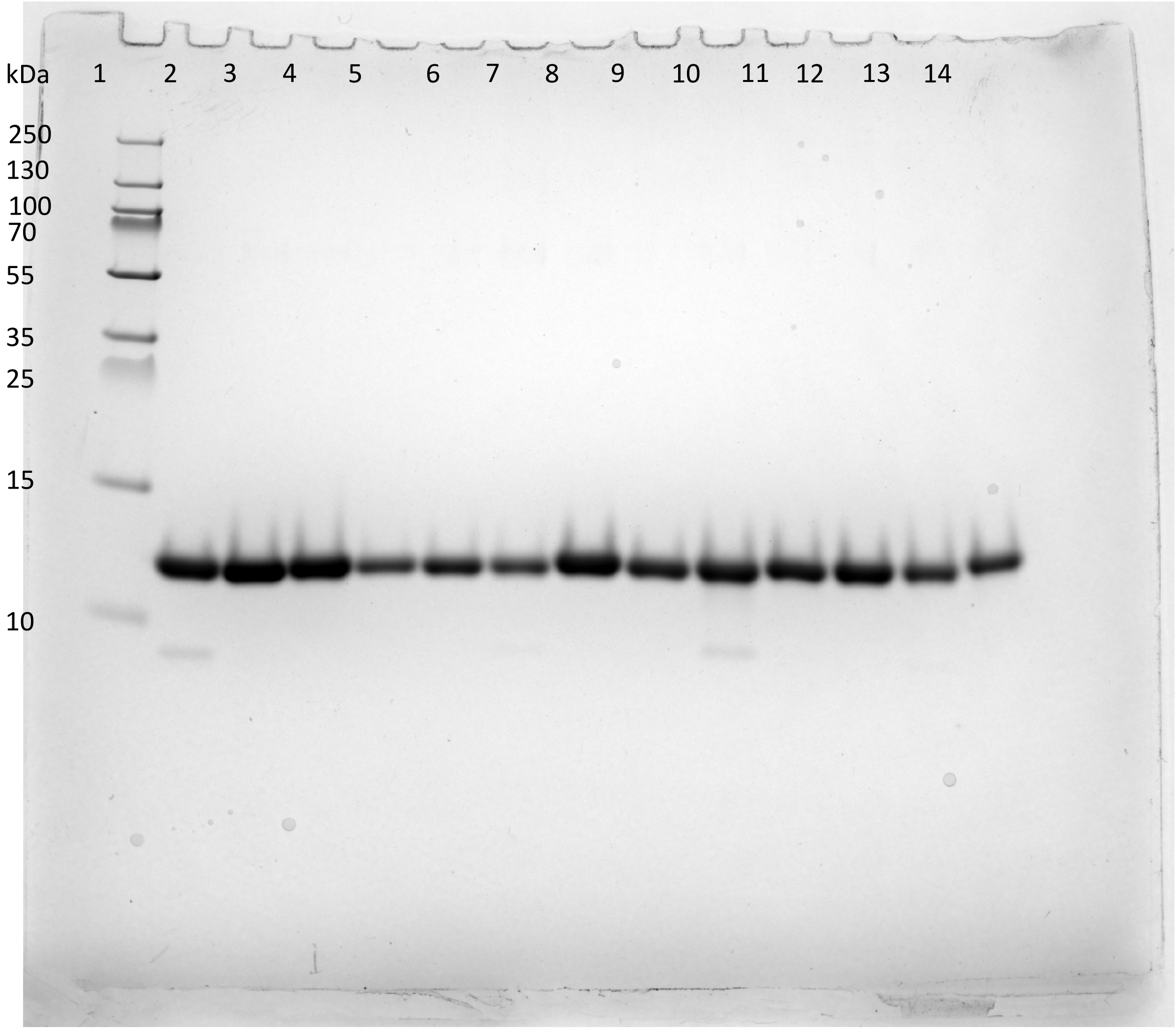

**1:** PageRuler Plus Prestained Protein Ladder (Thermo Scientific, Cat. No. 26619). **2:** Unmodified. **3:** P301S. **4:** AcK369. **5:** AcK353. **6:** AcK317. **7:** AcK311. **8:** pS356. **9:** pS352. **10:** pY310. **11:** pS305. **12:** GlcNAcN359. **13:** GlcNAcN359 + AcK353. **14:** GlcNAcN359 + pS305

#### 11. CD SPECTROSCOPY

Lyophilized Tau(291-391) was dissolved in 20 mM sodium phosphate buffer (pH 7.4), and the residual TFA salts were removed using a 7,000 MWCO Slide-A-Lyzer MINI Dialysis Unit (Thermo Scientific, Cat. No. 69562). The proteins were adjusted to 0.4 mg/mL in 20 mM sodium phosphate buffer (pH 7.4) and analyzed in triplicate by CD spectroscopy. The triplicate CD data were averaged and reported as molar eplicity.

**CD Spectrometer Settings:**
Pathlength: 0.05 cm
Wavelength: 180 nm – 260 nm
Step Size: 0.5 nm
Bandwidth: 1 nm
Repeats: 5 repeats for each sample, one repeat for each baseline

**Temperature Dependent CD Spectra of Tau(291-391):**

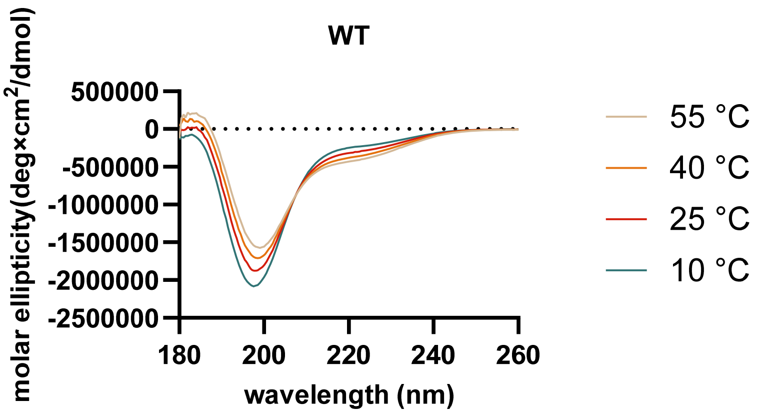

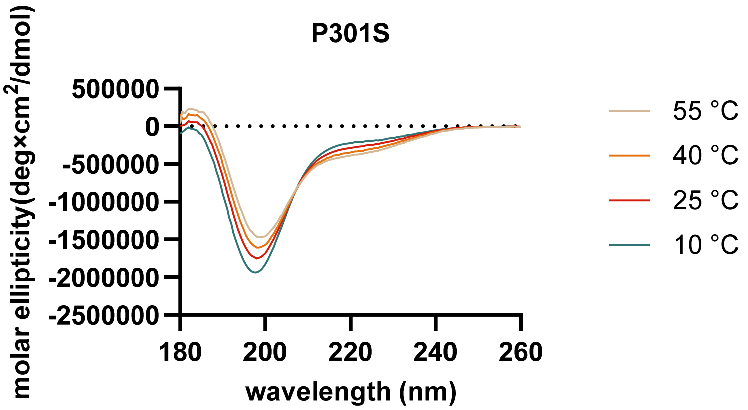

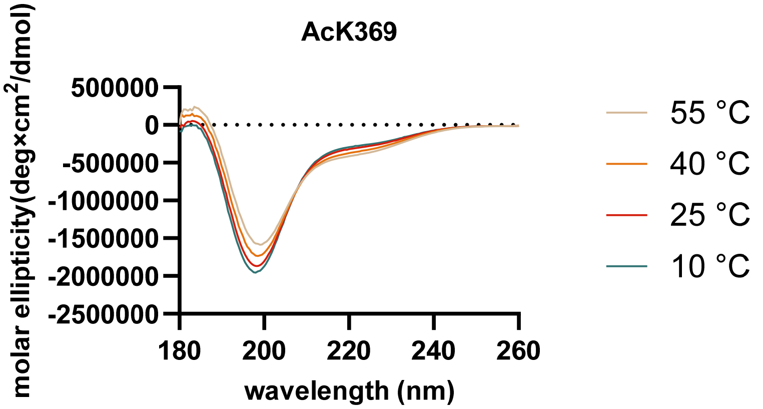
AcK369
500000
0
-500000
-1000000
-1500000
-2000000
-2500000
180
200
220
240
260
wavelength (nm)
molar ellipticity(deg×cm2/dmol)
55 °C
40 °C
25 °C
10 °C

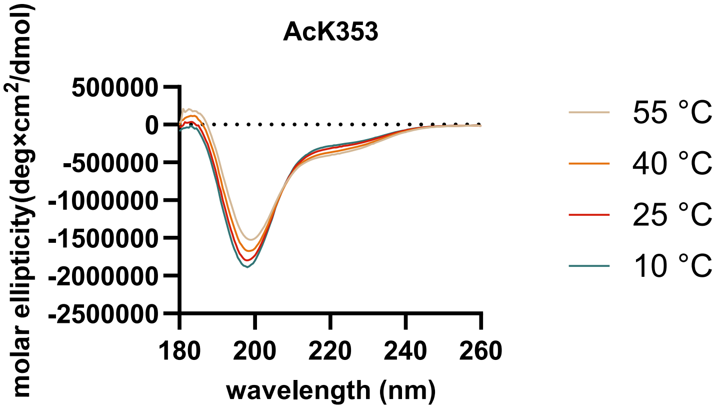
AcK353
500000
0
-500000
-1000000
-1500000
-2000000
-2500000
180
200
220
240
260
wavelength (nm)
molar ellipticity(deg×cm2/dmol)
55 °C
40 °C
25 °C
10 °C

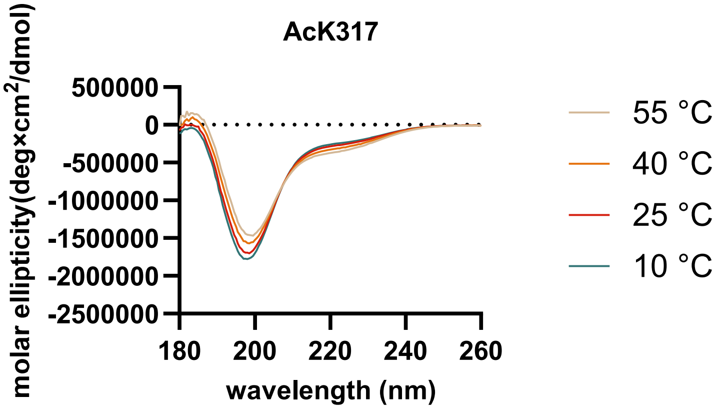
AcK317
500000
0
-500000
-1000000
-1500000
-2000000
-2500000
180
200
220
240
260
wavelength (nm)
molar ellipticity(deg×cm2/dmol)
55 °C
40 °C
25 °C
10 °C

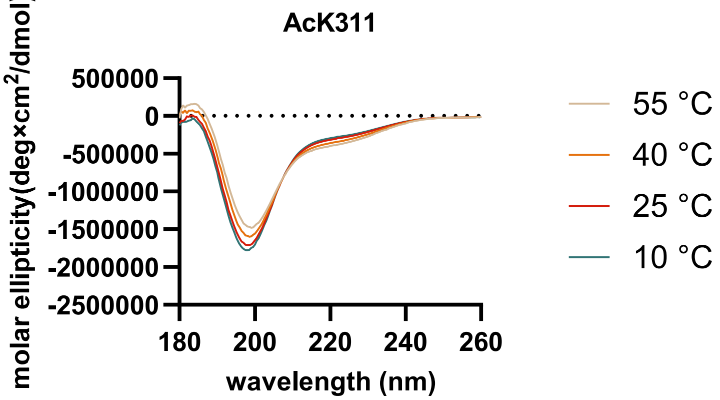
AcK311
molar ellipticity(deg×cm2/dmol)
500000
0
-500000
-1000000
-1500000
-2000000
-2500000
180
200
220
240
260
wavelength (nm)
55 °C
40 °C
25 °C
10 °C

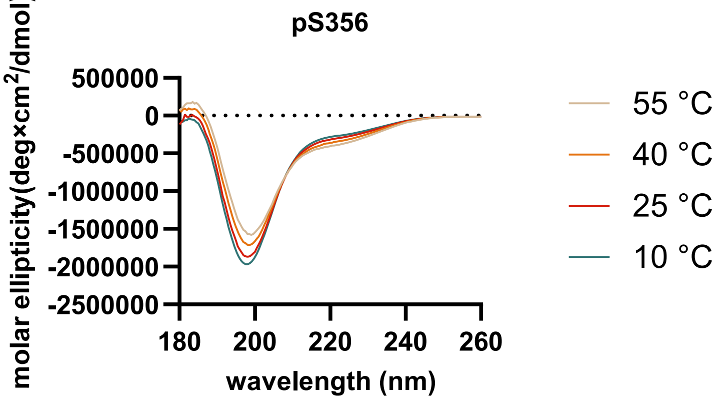
pS356
molar ellipticity(deg×cm2/dmol)
500000
0
-500000
-1000000
-1500000
-2000000
-2500000
180
200
220
240
260
wavelength (nm)
55 °C
40 °C
25 °C
10 °C

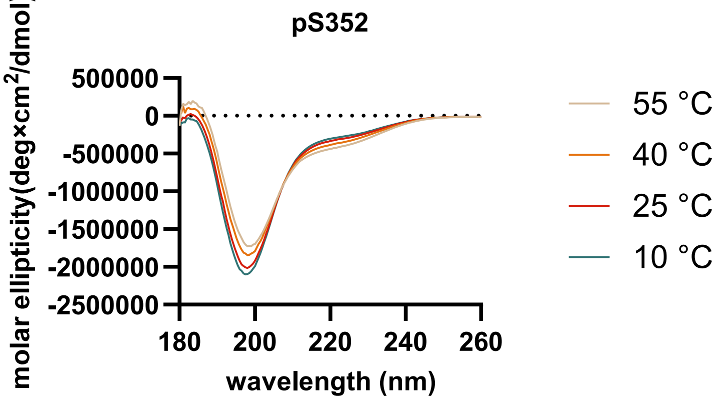
pS352
molar ellipticity(deg×cm2/dmol)
500000
0
-500000
-1000000
-1500000
-2000000
-2500000
180
200
220
240
260
wavelength (nm)
55 °C
40 °C
25 °C
10 °C

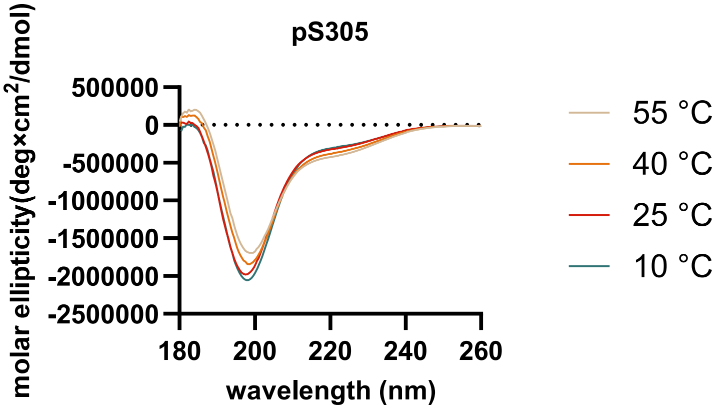
pS305
molar ellipticity(deg×cm2/dmol)
500000
0
-500000
-1000000
-1500000
-2000000
-2500000
180
200
220
240
260
wavelength (nm)
55 °C
40 °C
25 °C
10 °C

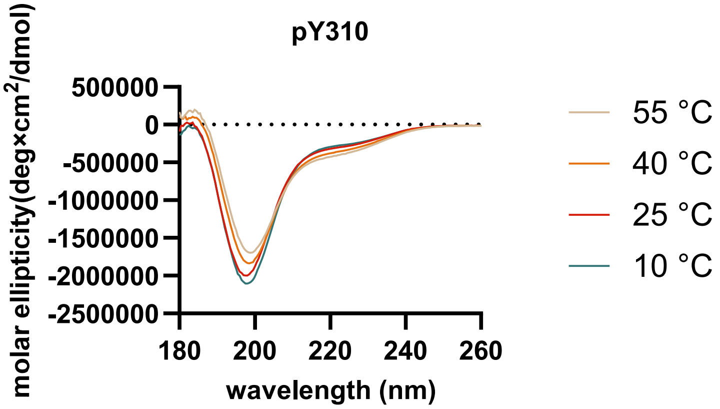
pY310
molar ellipticity(deg×cm2/dmol)
500000
0
-500000
-1000000
-1500000
-2000000
-2500000
180
200
220
240
260
wavelength (nm)
55 °C
40 °C
25 °C
10 °C

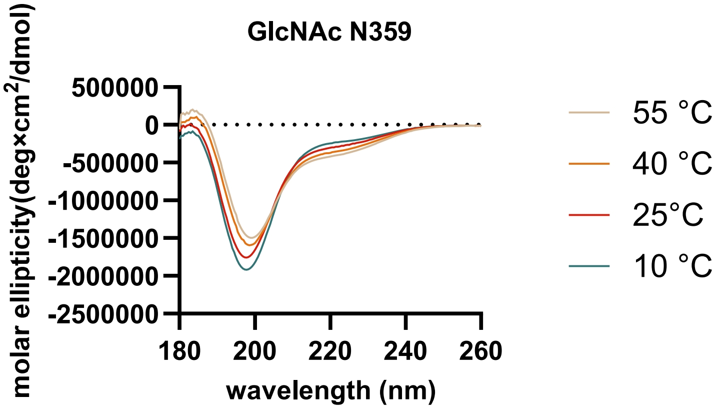
GlcNAc N359
molar ellipticity(deg×cm2/dmol)
500000
0
-500000
-1000000
-1500000
-2000000
-2500000
180
200
220
240
260
wavelength (nm)
55 °C
40 °C
25°C
10 °C

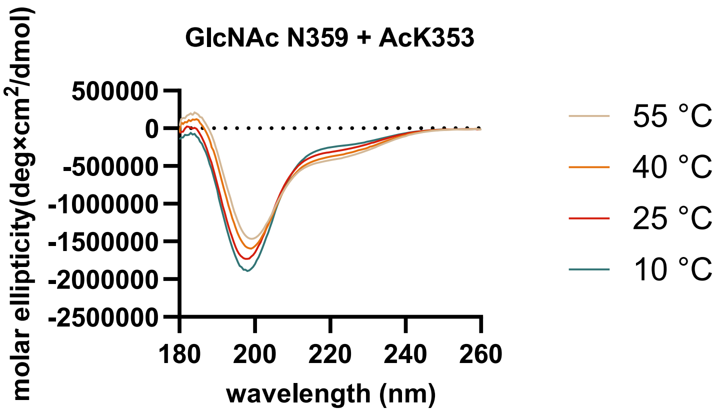
GlcNAc N359 + AcK353
molar ellipticity(deg×cm2/dmol)
500000
0
-500000
-1000000
-1500000
-2000000
-2500000
180
200
220
240
260
wavelength (nm)
55 °C
40 °C
25 °C
10 °C

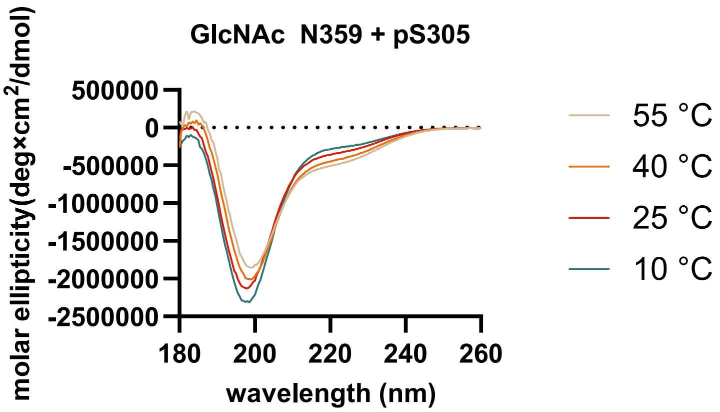
GlcNAc N359 + pS305
molar ellipticity(deg×cm2/dmol)
500000
0
-500000
-1000000
-1500000
-2000000
-2500000
180
200
220
240
260
wavelength (nm)
55 °C
40 °C
25 °C
10 °C

#### 12. NEGATIVE STAIN TEM

The fibril samples were directly imaged by transmission electron microscopy (TEM) after the aggregation. Filament preparations (5 µL) were applied to glow-discharged, carbon film 200 Mesh Cu grids (Electron Microscopy Sciences, CF200-CU) for 60s. The sample was blotted off and washed twice with $H_2O$. The sample was negatively stained twice with 3% uranyl acetate (5 µL) and imaged on an FEI Tecnai T12 Spirit at 120 kV with a LaB6 filament.

#### 13. COFACTOR-FREE SELF ASSEMBLY

This protocol was adapted from the literature.[7] Lyophilized Tau(291-391) was dissolved in 10 mM sodium phosphate, 10 mM DTT (pH 7.4) and purified by SEC over a Superdex 75 10/300 GL into 10 mM sodium phosphate, 10 mM DTT (pH 7.4). The fractions containing protein were concentrated to 8 mg/mL using a Pierce Concentrator (PES, 3K MWCO, 0.5 mL, 88512).

The cofactor free self-assembly was run in a 96 well plate (Greiner BioOne, 96 well, PS, F Bottom, Chimney well, black, medium binding, 655096). The wells were filled with 100 µL of the sample with 4 mg/mL Tau(291-391), 200 mM $MgCl_2$, 10 mM DTT, 10 mM sodium phosphate (pH 7.4). The plate was sealed with an adhesive film (VWR, polyester foil, Cat. No. 89134-430) and mixed at 37 ºC at 300 RPM on a thermomixer (Ika Matrix Orbital). After 48 h, the aggregation mixtures were directly assayed for fibrils by negative stain TEM.

**Stock Solutions.** Filtered through a 0.22 µM filter prior to use.

- 10 mM sodium phosphate (pH 7.4)
- 1 M $MgCl_2$ in 10 mM sodium phosphate (pH 7.4). Readjusted to pH 7.4 after dissolving $MgCl_2$.
- 100 mM DTT in 10 mM sodium phosphate (pH 7.4). Readjusted to pH 7.4 after dissolving DTT.
- 8 mg/mL Tau(291-391) and 10 mM DTT in 10 mM sodium phosphate (pH 7.4).

**Final Concentration.** 4 mg/mL Tau(291-391), 200 mM $MgCl_2$, 10 mM DTT, 10 mM sodium phosphate (pH 7.4), 37 ºC, 300 RPM.

#### 14. ELISA OF TAU-TAU BINDING

This protocol was taken from literature with minor modifications.[8,9] The difference with our protocol is that we used 2N4R Tau(2-441) in the solution phase instead of Tau(297-391), and we used anti-Tau 15-25 mouse antibody (binds residues 15-25 on 2N4R tau, BioLegend, 835201) as the primary antibody instead of mAb 423, and anti-mouse IgG (H+L), HRP conjugate (Promega, W4021) as the secondary antibody.

The dilutions were an 8-point curve with 3 wells each. This makes for a total of 24 wells per protein, per run. The experiment was repeated three times, and the data from each experiment was averaged, normalized, and reported graphically. The error bars on the graphs represent the standard deviation.

Detailed protocol:

1. Pipette 50 µL of 1 µM Tau(291-391) into 24 wells of a PVC microplate (costar Serocluster, 96 well U Bottom Plate, Non-Treated, Vinyl, Ref. 2797). Incubate the plate for 1 h at 37 ºC. Wash plate with wash buffer (200 µL x 3).
2. Pipette 200 µL of block buffer per well. Incubate plate for 1 hour at 37 ºC. Wash plate with wash buffer (200 µL x 3).
3. Add 50 µL of each Tau(2-441) dilution in binding buffer into three wells (21 wells total). Pipette 50 µL of binding buffer into the last three wells. Incubate for 1 hour at 37 ºC. Wash plate with wash buffer (200 µL x 3).
4. Pipette 50 µL of primary antibody in blocking buffer to each well. Incubate plate for 1 h at 37 ºC. Wash plate with wash buffer (200 µL x 3).
5. Pipette 50 µL of secondary antibody solution into each well. Incubate plate for 1 h at 37 ºC. Incubate plate for 1 hour at 37 ºC. Wash plate with wash buffer (200 µL x 3). Pipette 50 µL of TMB solution to each well, and immediately transfer to the SpectraMax iD5 plate reader and run the kinetic assay protocol.
6. ELISA-TMB Kinetic Assay: Monitor absorbance at 650 nm. Read time: 3 minutes, every 22 s. Shaking before the first read for 5 s, and then no shaking.
7. The data was baseline corrected to the wells without Tau(2-441). Using Prism/Graphpad, the data from three experiments is normalized, averaged, and presented as the standard deviation from three experiments.
8. The data is fit to a nonlinear regression.
9. The curve fit is "One site- Total and nonspecific binding" with a constraint of background binding equal to zero.
10. This assay is repeated for each modified Tau(291-391) construct, to obtain the $K_d$ of tau-tau binding.

**Stock Solutions**:

- Carbonate Buffer: 50 mM sodium carbonate-sodium bicarbonate (pH 9.6).
- Blocking Buffer: 2 % (w/v) dried milk powder in PBS (pH 7.4).
- Wash Buffer: 0.05% Tween 20 in $H_2O$.
- PBST: 0.05% Tween 20 in PBS (pH 7.4)
- Binding Buffer: 50 mM MES, 25 mM NaCl, 10 mM PIPES, 0.05 % Tween® 20, 1 % (w/v) fish skin gelatine (pH 6.0).
- Primary detection antibody: 1:1000 dilution of Anti-Tau, 15-25 mouse antibody in block buffer.
- Secondary detection antibody: 1:1000 dilution of anti-mouse IgG (H+L), HRP conjugate in PBST.

- Tau(291-391): 1 µM Tau(291-391) in binding buffer.
- Tau(2-441): Prepare a 1:1 serial dilution series of Tau(2-441) (Acro Biosystems, Cat. No. H5117) in binding buffer. The concentration of Tau(2-441) in the dilution series is: 1 µM, 0.5 µM, 0.25 µM, 0.0125 µM, 0.0625 µM, 0.0313 µM, and 0.0156 µM.
- TMB solution: 0.1 mg/mL 3,3,5',5'-tetramethylbenzidine (Sigma Aldrich), 0.003% hydrogen peroxide, 50 mM sodium acetate (pH 5.0). Prepared fresh before use.

**Microplate Settings:**

- Wavelength: Absorbance at 650 nm.
- Timing: Read every 22 s over 3 min.
- Shaking: Shake for 30 s before the first read, and then no shaking.

**One site total and nonspecific binding equation:**

$$y = \frac{B_{max} \cdot x}{x + K_d} + NS \cdot x + BG$$

- BG: Background, BG = 0 from normalization.
- $B_{max}$: Maximum specific binding, in same units as y.
- $K_d$: Equilibrium dissociation constant, in same units as x.
- NS: The slope of non-specific binding, in units of y/x.

**Table S1: Calculated Binding Values**

| Tau | $K_d$ (nM) | SD of $K_d$ (nM) | Replicates |
|---|---|---|---|
| WT | 6.017 | 1.324 | 3 |
| P301S | 8.088 | 2.1885 | 3 |
| AcK369 | 3.716 | 1.58 | 3 |
| AcK353 | 7.263 | 2.468 | 3 |
| AcK317 | 8.631 | 3.395 | 3 |
| AcK311 | 10.5 | 2.33 | 3 |
| pS356 | 15.55 | 4.025 | 3 |
| pS352 | 13.08 | 3.535 | 3 |
| pY310 | 6.466 | 1.487 | 3 |
| pS305 | 23.34 | 5.035 | 3 |
| GlcNAc N359 | 19.94 | 5.65 | 3 |
| GlcNAc N359 + AcK353 | 17.78 | 2.3 | 3 |
| GlcNAc N359 + pS305 | 7.114 | 1.267 | 3 |

**Langmuir curves for binding 2N4R Tau(2-441) to Tau(291-391):**

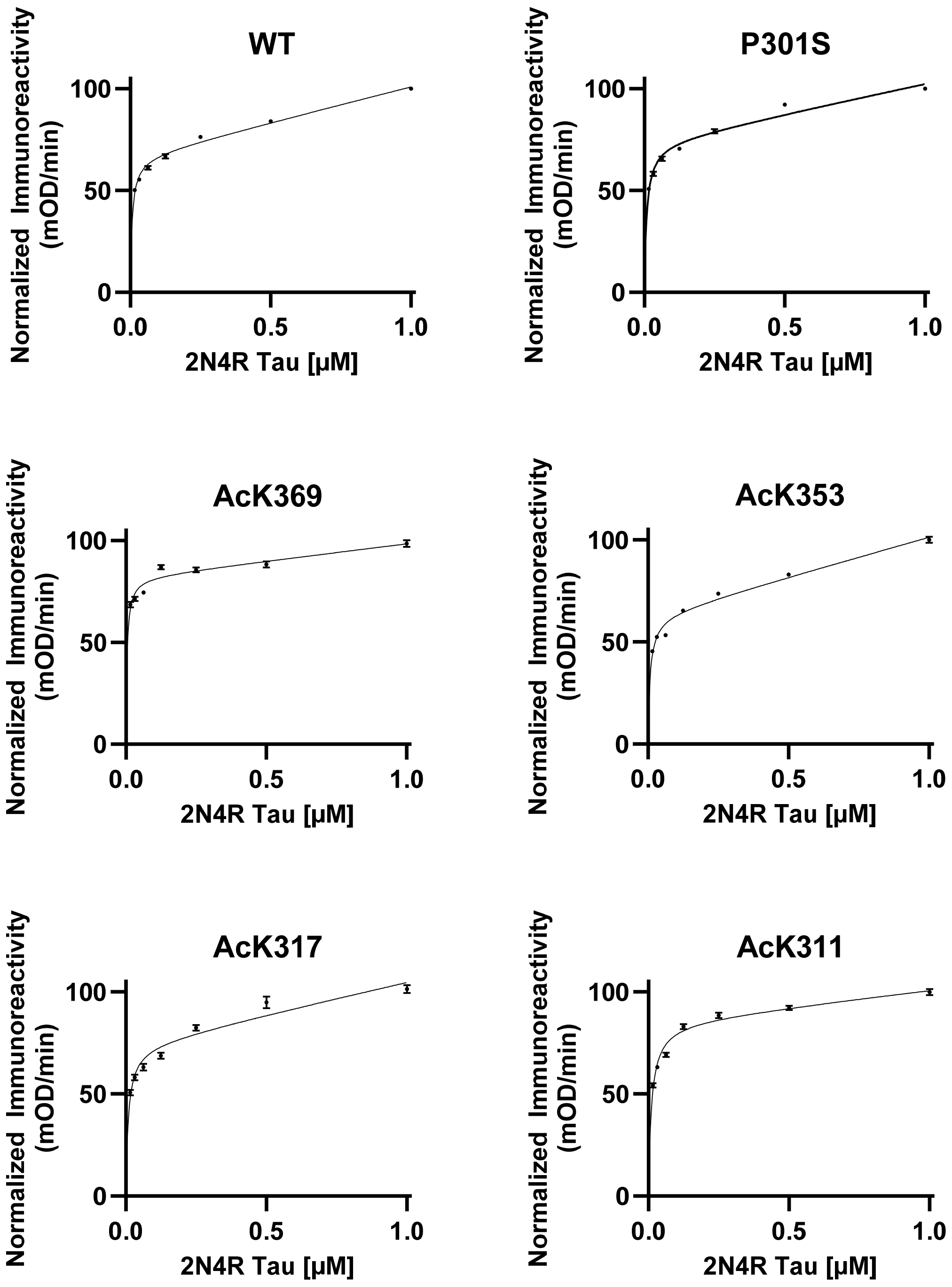

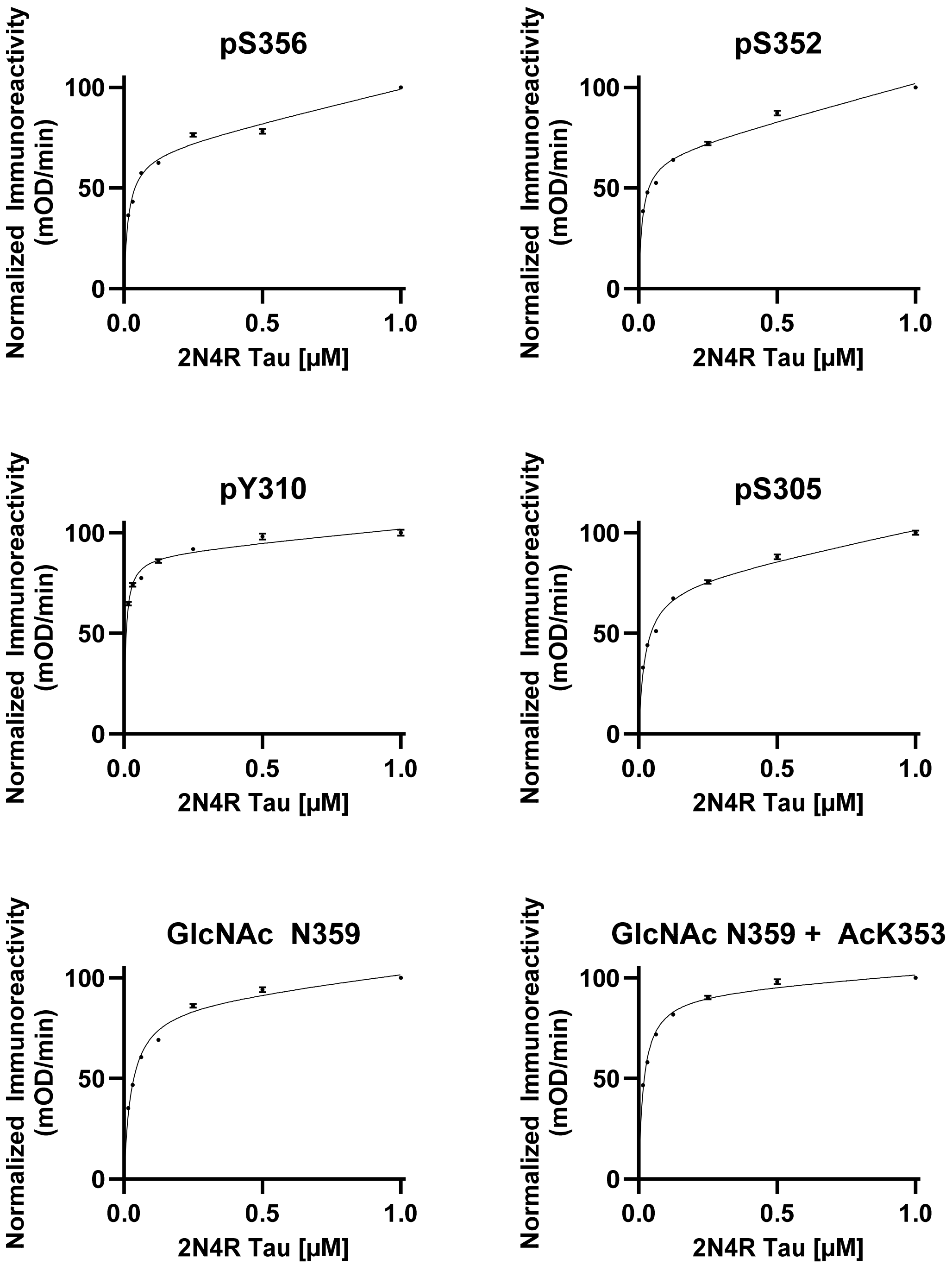
pS356
pS352
pY310
pS305
GlcNAc N359
GlcNAc N359 + AcK353
Normalized Immunoreactivity (mOD/min)
2N4R Tau [μM]
0
50
100
0.0
0.5
1.0

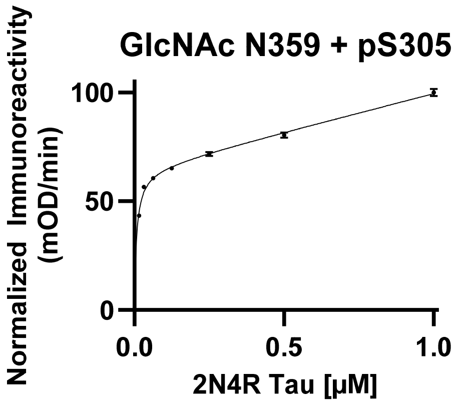
GlcNAc N359 + pS305
Normalized Immunoreactivity
(mOD/min)
100
50
0
0.0
0.5
1.0
2N4R Tau [µM]

#### 15. HEPARIN INDUCED AGGREGATION

A 0.65 µL Eppendorf tube was charged with 50 µL of aggregation assay mixture at the final concentration listed. The buffers and salts were added first, followed by protein, and heparin was added last. Three wells on a black 384-well non-binding microplate (Greiner BioOne, PS, F Bottom, small volume, HiBase, REF 784900) were filled with 15 µL of the aggregation assay mixture, the plate was sealed with polyester adhesive film (VWR, Cat. No. 89134-430), and the plate was incubated at 37 ºC in the microplate reader. The data from the three wells were averaged, and the experiment was repeated for a total of three times. The data was averaged then normalized, and the error bars on the graphs were reported as the standard error of the mean.

**Stock Solutions:**

- 25 mM HEPES (pH 7.4).
- 200 mM DTT in 25 mM HEPES (pH 7.4).
- 440 mM Heparin sodium salt from porcine intestinal mucosa (Sigma H4784, 18 kDa average molar weight, 1 mg/mL = 55 µM) in 25 mM HEPES (pH 7.4).
- 150-300 µM Tau(291-391) in 25 mM HEPES (pH 7.4).
- 500 µM Thioflavin t in 25 mM HEPES (pH 7.4).

**Final Concentration:**

- 100 µM Tau(291-391), 40 µM heparin, 20 µM thioflavin T, 10 mM DTT, 50 mM NaCl, 25 mM HEPES (pH 7.4), 37 ºC.

**Microplate Settings:**

- Wavelength: EX 440 nm, EM 480 nm.
- PMT gain: 500 V, read 9 mm from the top of the plate.
- Integration time: 400 ms.
- Timing: Read every 10 minutes over 72 h.
- Shaking: shake for 30 s before the first read, and then no shaking.

**Thioflavin T Plots of Tau(291-391) Aggregation:**

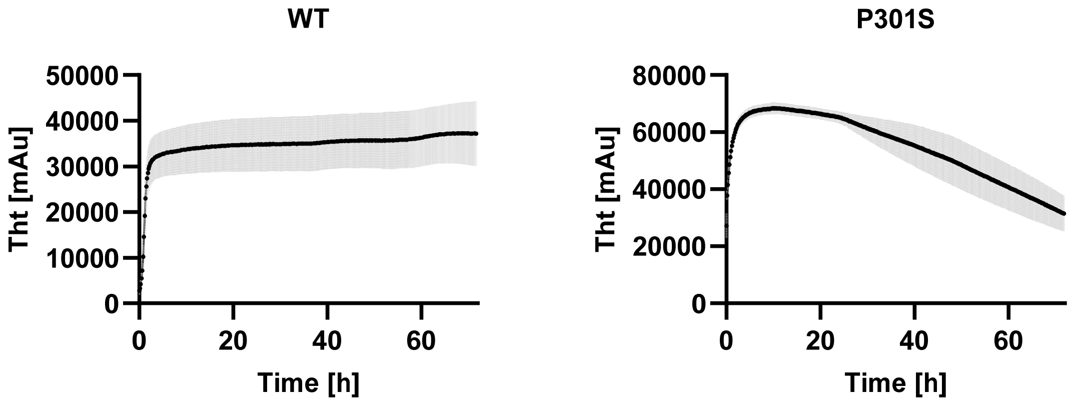

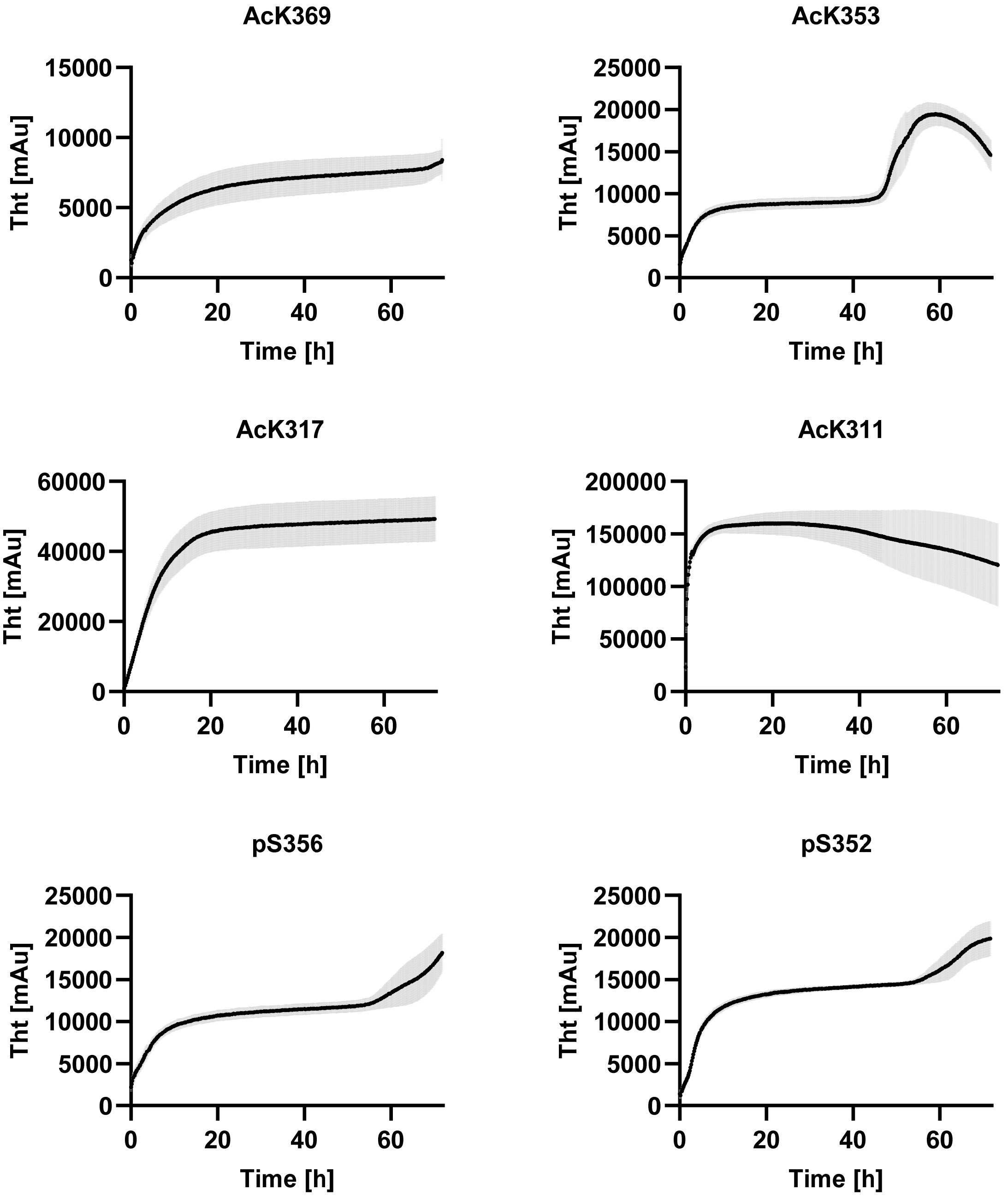
AcK369
Tht [mAu]
15000
10000
5000
0
0
20
40
60
Time [h]
AcK353
Tht [mAu]
25000
20000
15000
10000
5000
0
0
20
40
60
Time [h]
AcK317
Tht [mAu]
60000
40000
20000
0
0
20
40
60
Time [h]
AcK311
Tht [mAu]
200000
150000
100000
50000
0
0
20
40
60
Time [h]
pS356
Tht [mAu]
25000
20000
15000
10000
5000
0
0
20
40
60
Time [h]
pS352
Tht [mAu]
25000
20000
15000
10000
5000
0
0
20
40
60
Time [h]

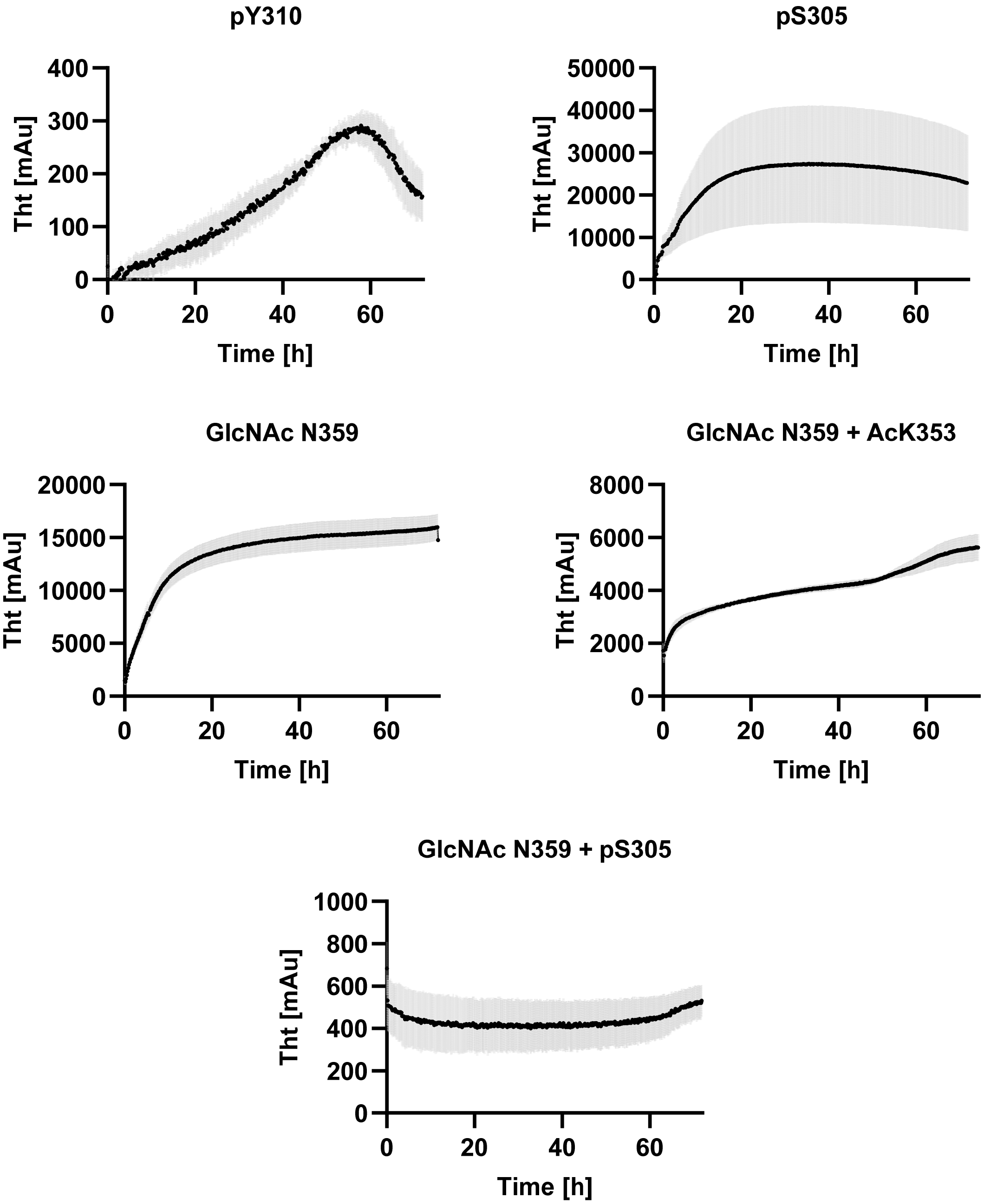

pY310
Tht [mAu]
400
300
200
100
0
0
20
40
60
Time [h]
pS305
Tht [mAu]
50000
40000
30000
20000
10000
0
0
20
40
60
Time [h]
GlcNAc N359
Tht [mAu]
20000
15000
10000
5000
0
0
20
40
60
Time [h]
GlcNAc N359 + AcK353
Tht [mAu]
8000
6000
4000
2000
0
0
20
40
60
Time [h]
GlcNAc N359 + pS305
Tht [mAu]
1000
800
600
400
200
0
0
20
40
60
Time [h]

#### 16. FIBRIL J ANALYSIS

The heparin induced aggregated samples were diluted (1:10) prior to staining. For the analysis, 3 replicates of each Tau(291-391) construct were imaged and 10 random locations. The 10 images for each replicate were taken at 30,000x (2.119 nm/pixel). Data from these replicates were pooled for subsequent analysis by FibrilJ.

Tau(291-391) fibrils were analyzed with the ImageJ fibril analysis plugin FibrilJ.[10] A modified FibrilJ that included a more detailed skeleton pruning algorithm was used to reduce the fibrils to lines and minimize branches mistakenly created during the skeletonization. Prior to analysis, images were binarized using custom segmentation algorithms. Out of the three diameter algorithms available in FibrilJ, results from the D2 (human-like) algorithm were presented in the results section, as D1 (area/length) and D3 (distance map) were not consistently calculated.

FibrilJ is limited in accurately distinguishing fibril lengths due to the innate limitations of the ImageJ 2D/3D Skeleton program and the AnalyzeSkeletons-based pruning plugin. When pruning the skeletons of the fibrils, FibrilJ trims off the shortest branches, which occasionally bisects the fibrils at random parts throughout their length. This frequently occurs when fibrils intersect or overlap, or when the positive stain is uneven, affecting the binary image. In these cases, FibrilJ often skeletonizes one fibril as multiple, providing the lengths of multiple sections of the fibril rather than the fibril in its entirety. The properties of the fibrils in the image were exported as an Excel sheet, with IDs for each fibril that FibrilJ detected and measured. Thus, the length data for the fibrils was visually examined and false fibrils detected on the images, as well as combining different fibril IDs that align to form one fibril. This increased the accuracy of length data compared to the program's original protocol. Obtaining accurate fibril length was further optimized by using lower density images with minimal fibril overlap, and regions of good negative (or positive) stain, permitting a more even image binarization and preventing fibril bisection caused by stain intensity differences on the image.

Data from the 3 fibril replicates in each group was compared and collated for inter-group comparison. The Kolmogorov-Smirnoff test was utilized to compare the length and diameter distributions of different fibril types.

The fibril diameters and lengths were analyzed with Fibril J. The diameters of the wild type (WT) Tau(291-391) fibrils, compared to the fibrils modified by acetylation, glycosylation, P301S mutation, and phosphorylation. There were clear differences in diameter between fibril types. Acetylations at all positions tested decreased the average diameter by 1-3 nanometers. Glycosylation had less of an effect on decreasing fibril diameter, the glycosylated K353Ac sample had a 1 nm narrower mean diameter than non-glycosylated K353Ac, while a glycosylation at N359 had a mean diameter only 0.2 nm smaller than WT. Phosphorylation of serine 352 and 356 had a similarly low effect on fibril diameter compared to glycosylation of other positions; they have effectively the same mean diameter, both 1 nm narrower than WT.

However, it is noteworthy that the pS356 sample appeared to have a bimodal distribution with a smaller peak at around 8 nm along with the larger peak at 14 nm. Mutated fibril P301S had a much broader distribution of diameters than all other modifications, with a mean diameter approximately 1.5 nm smaller than the wild type. Kolmogorov-Smirnoff tests were conducted to compare the diameter distributions of WT fibrils to those of modified fibrils, and the differences were very statistically significant ($p < 0.0001$) for all samples.

Ranges of fibril lengths across samples were shown in the insert (Figure 5e). All modified groups, excluding AcK369, had no fibrils longer than present in the WT sample; however, a few fibrils of K369Ac were much longer than those in other samples. P301S fibrils were the shortest with no fibrils reaching 1000 nm in length, and the vast majority being between 100-200 nm long. Kolmogorov-Smirnoff tests comparing the fibril groups concluded that these differences in length were statistically significant (all comparisons had p-values <0.0001).

**Table S2: Average Fibril Diameter**

| Sample | Mean Diameter (nm) (SD) |
|---|---|
| WT | 13.63 (2.099) |
| P301S | 11.79 (2.505) |
| S352Phos | 12.79 (1.580) |
| S356Phos | 12.39 (2.494) |
| K311Ac | 12.07 (2.421) |
| K317Ac | 11.39 (2.326) |
| K353Ac | 10.70 (3.064) |
| GlcNAcK353Ac | 9.649 (1.612) |
| N359GlcNAc | 13.41 (2.099) |
| K369Ac | 12.51 (2.912) |

**Table S3: Average Fibril Length**

| Sample | Mean Length (nm) (SD) |
|---|---|
| WT | 231.3 (256.5) |
| P301S | 214.9 (133.5) |
| S352Phos | 228.2 (234.9) |
| S356Phos | 150.1 (134.2) |
| K311Ac | 247.3 (230.8) |
| K317Ac | 256.7 (281.9) |
| K353Ac | 288.6 (282.8) |
| GlcNAcK353Ac | 185.0 (199.1) |
| N359GlcNAc | 237.3 (207.9) |
| K369Ac | 236.5 (461.1) |

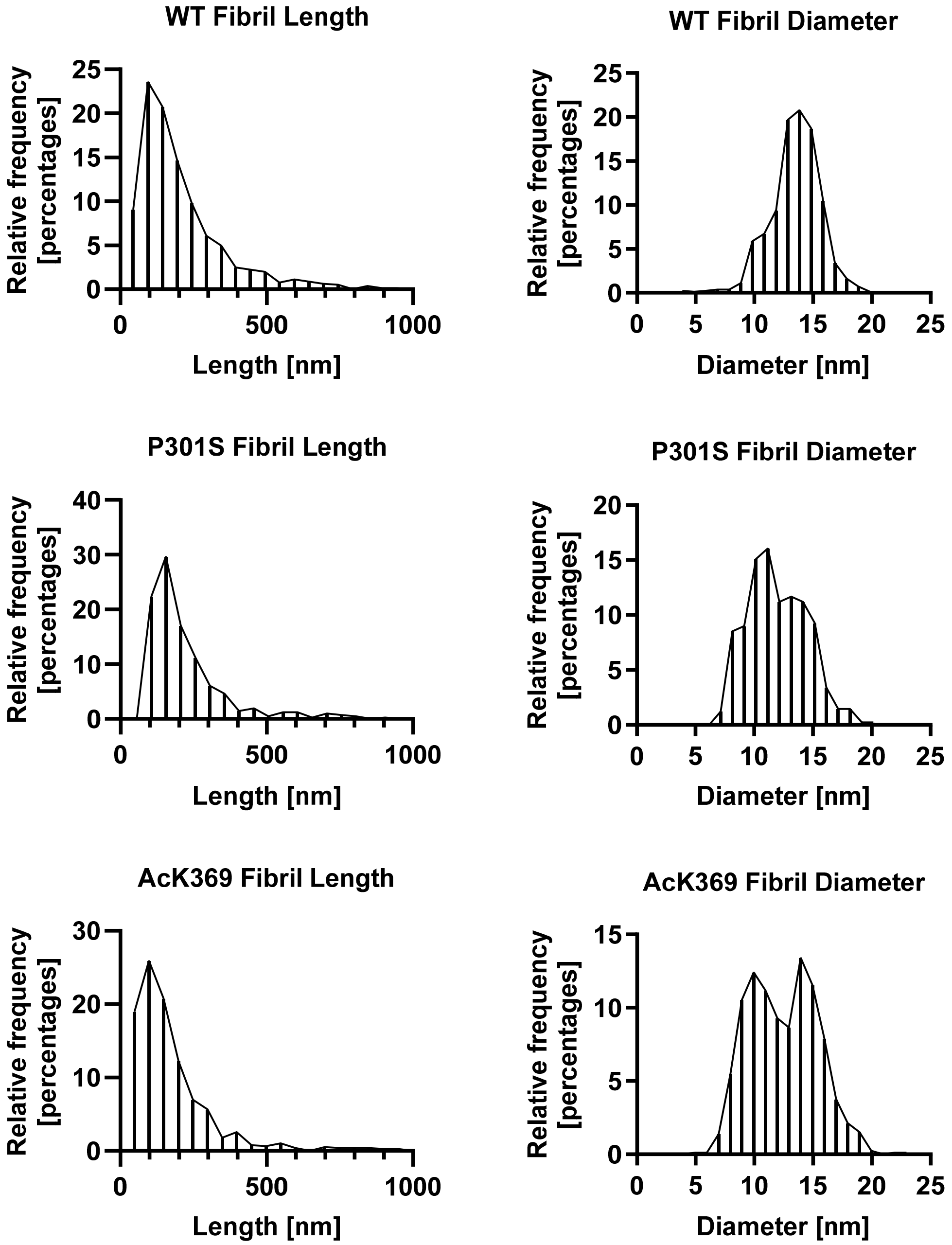
WT Fibril Length
Relative frequency [percentages]
Length [nm]
WT Fibril Diameter
Relative frequency [percentages]
Diameter [nm]
P301S Fibril Length
Relative frequency [percentages]
Length [nm]
P301S Fibril Diameter
Relative frequency [percentages]
Diameter [nm]
AcK369 Fibril Length
Relative frequency [percentages]
Length [nm]
AcK369 Fibril Diameter
Relative frequency [percentages]
Diameter [nm]

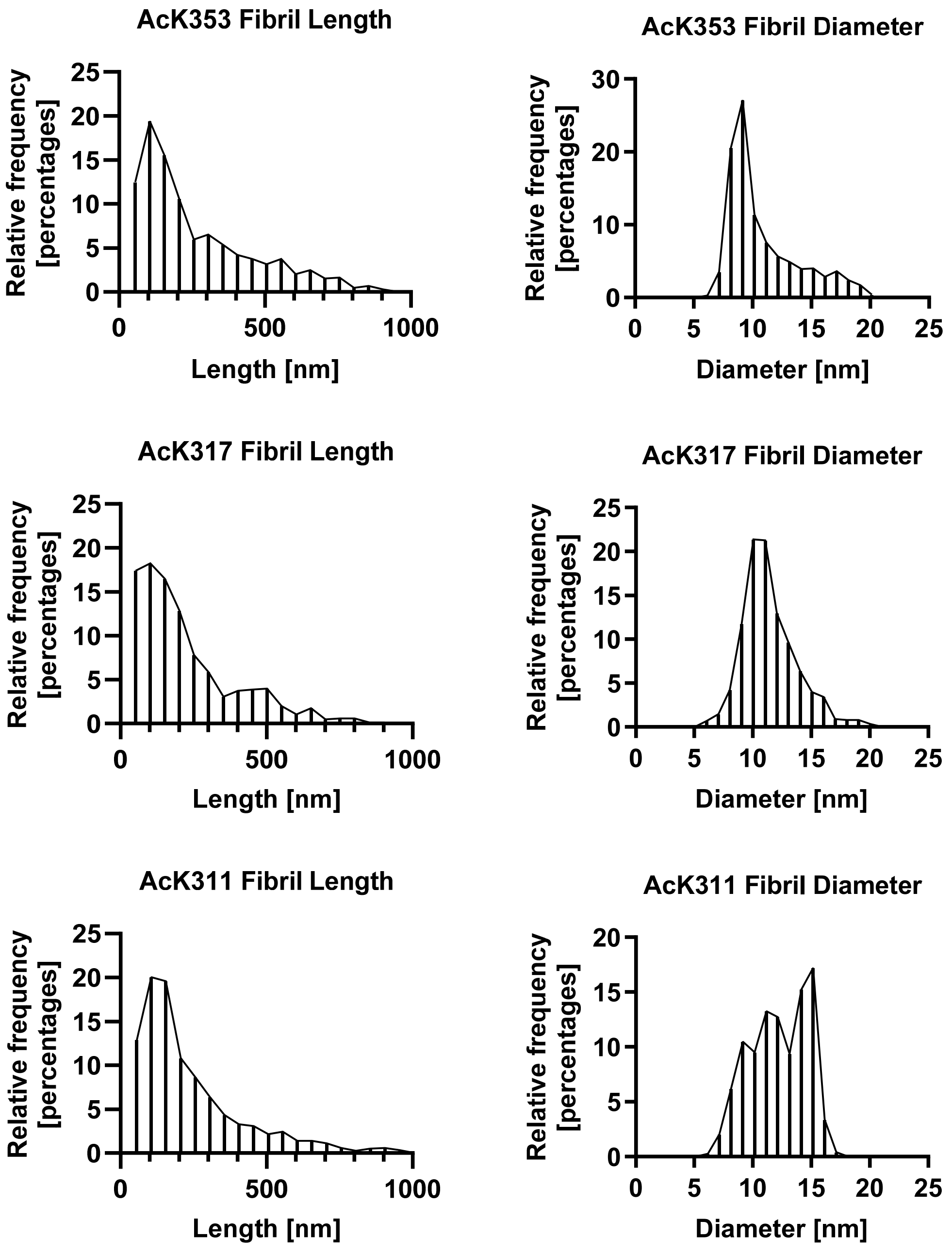
AcK353 Fibril Length
Relative frequency [percentages]
Length [nm]
AcK353 Fibril Diameter
Relative frequency [percentages]
Diameter [nm]
AcK317 Fibril Length
Relative frequency [percentages]
Length [nm]
AcK317 Fibril Diameter
Relative frequency [percentages]
Diameter [nm]
AcK311 Fibril Length
Relative frequency [percentages]
Length [nm]
AcK311 Fibril Diameter
Relative frequency [percentages]
Diameter [nm]

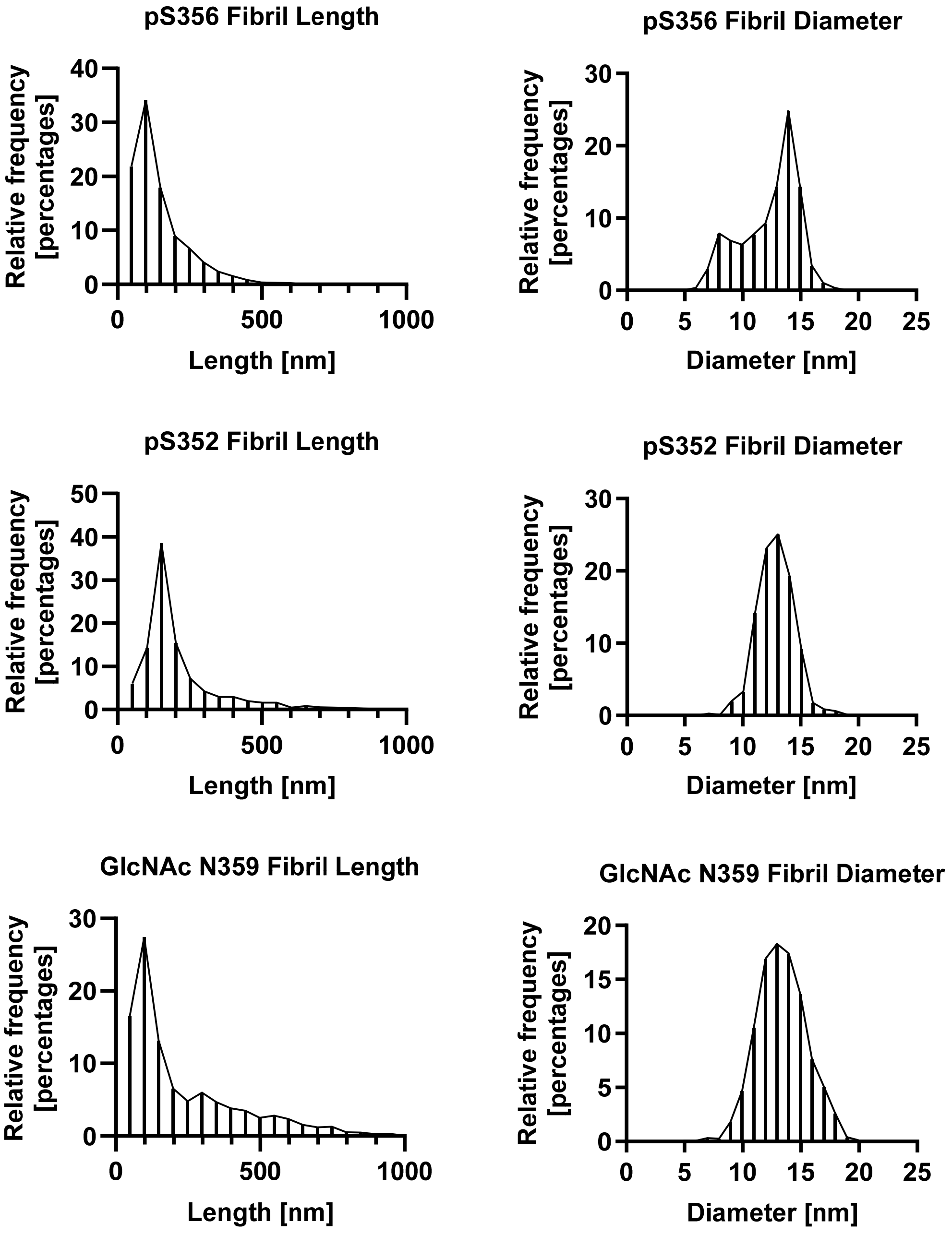
pS356 Fibril Length
Relative frequency [percentages]
Length [nm]
pS356 Fibril Diameter
Relative frequency [percentages]
Diameter [nm]
pS352 Fibril Length
Relative frequency [percentages]
Length [nm]
pS352 Fibril Diameter
Relative frequency [percentages]
Diameter [nm]
GlcNAc N359 Fibril Length
Relative frequency [percentages]
Length [nm]
GlcNAc N359 Fibril Diameter
Relative frequency [percentages]
Diameter [nm]

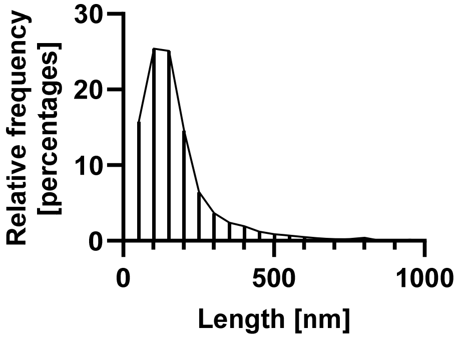
Relative frequency [percentages]
30
20
10
0
0
500
1000
Length [nm]

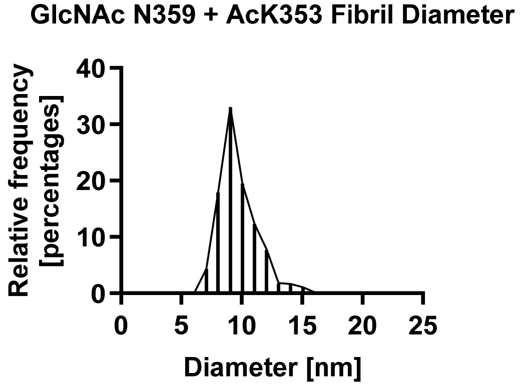
GlcNAc N359 + AcK353 Fibril Diameter
Relative frequency [percentages]
40
30
20
10
0
0
5
10
15
20
25
Diameter [nm]

#### 17. FIBRIL SORTING

The heparin induced aggregated samples were diluted (1:10) prior to staining. For the analysis, 3 replicates of each Tau(291-391) fibril were negatively stained and 3 random images at 30,000 x (2.119 nm/pixel) were obtained per replicate (9 images total). All the fibrils in each image were sorted into groups of fibrils that appear straight, jagged, hosed, or helical. The fibrils were then totaled, and the data were pooled for subsequent analysis. Clumped fibrils and overlapping fibrils were not counted in the analysis. Data from the replicates were averaged and displayed graphically with error bars that represent the standard deviation.

**Sorted Fibril Morphology of Tau(291-391)-Heparin Fibrils**

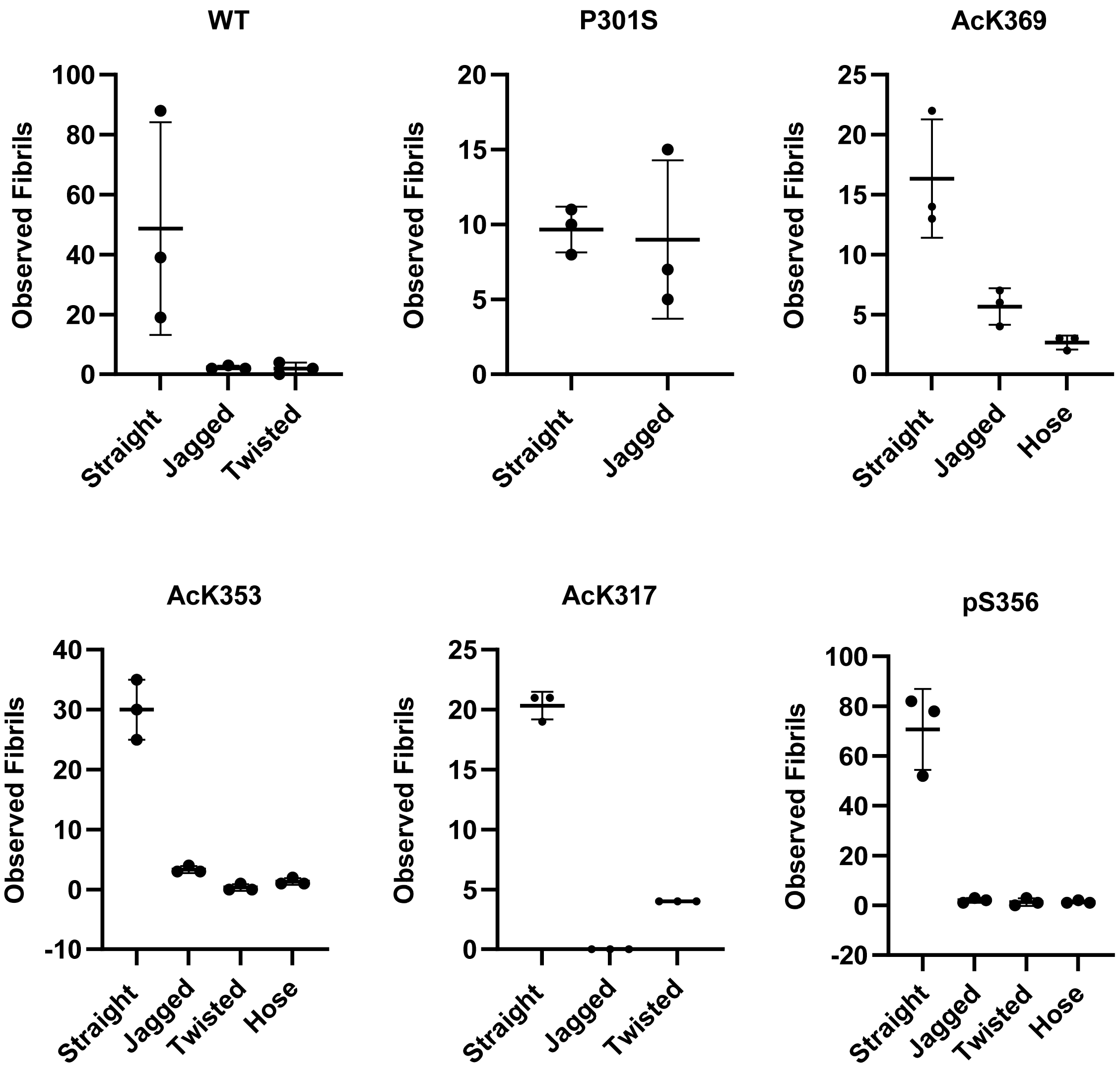

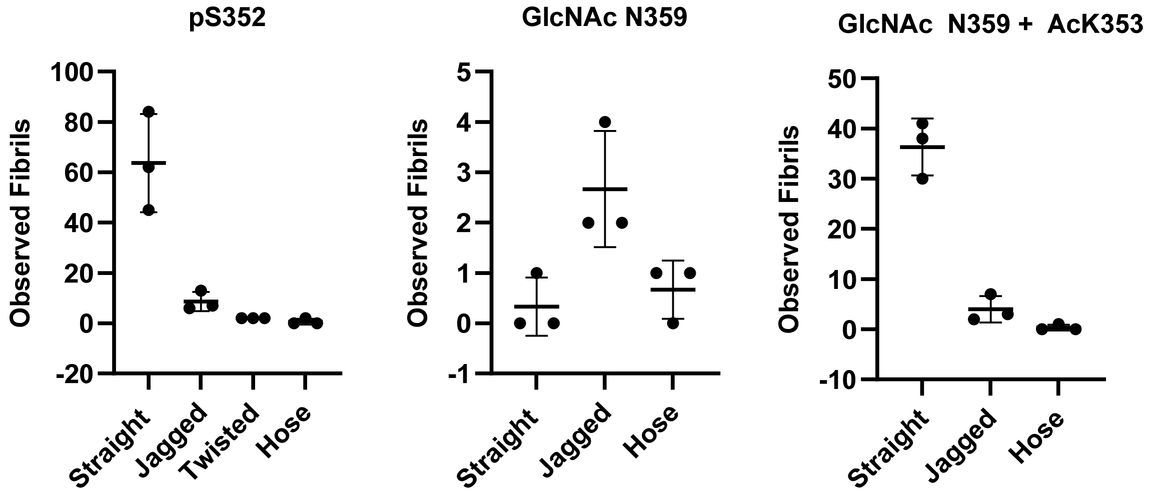
pS352
GlcNAc N359
GlcNAc N359 + AcK353
Observed Fibrils
100
80
60
40
20
0
-20
5
4
3
2
1
0
-1
50
40
30
20
10
0
-10
Straight
Jagged
Twisted
Hose

#### 18. CROSS OVER DISTANCE ANALYSIS

The crossover distance was measured at dark spots or thin regions, which were visible by negative stain TEM. The same images used for the fibril sorting were used, and the crossover distances were measured using ImageJ. One image from each replicate was selected, two to six fibrils were chosen of each morphology, and the crossover distance was measured at 5-20 locations along the entire fibril. The values were averaged, and the error bars were reported graphically the standard deviation.

**Cross Over Distance Measurements of Tau(291-391)-Heparin Fibrils**

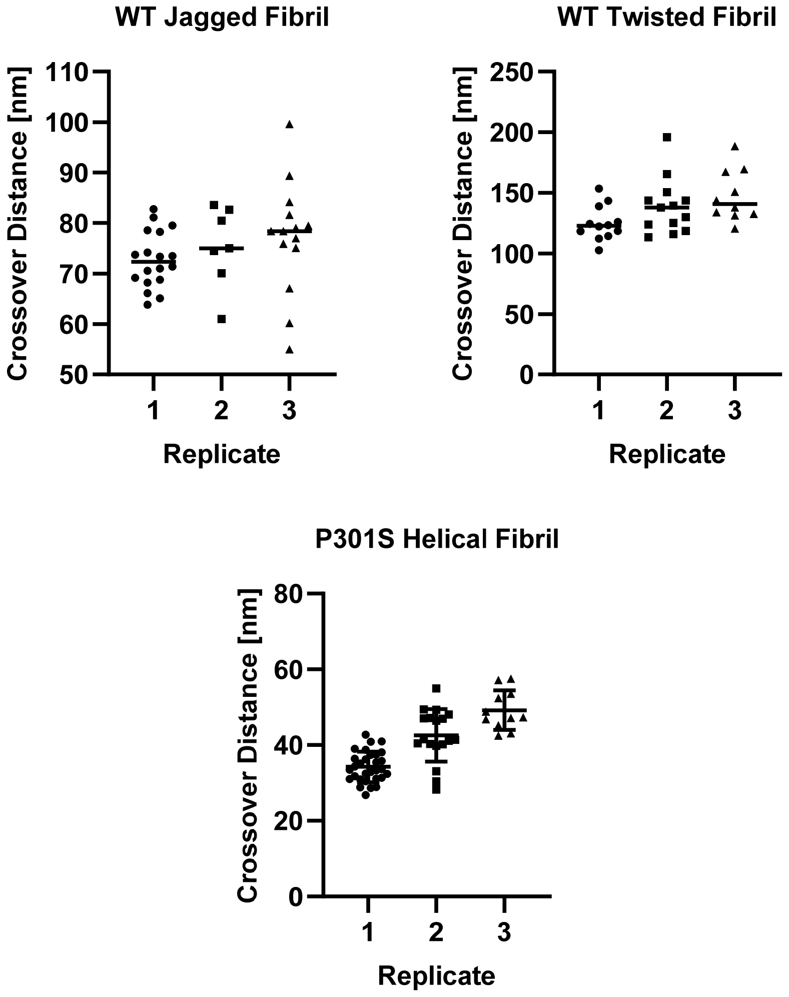

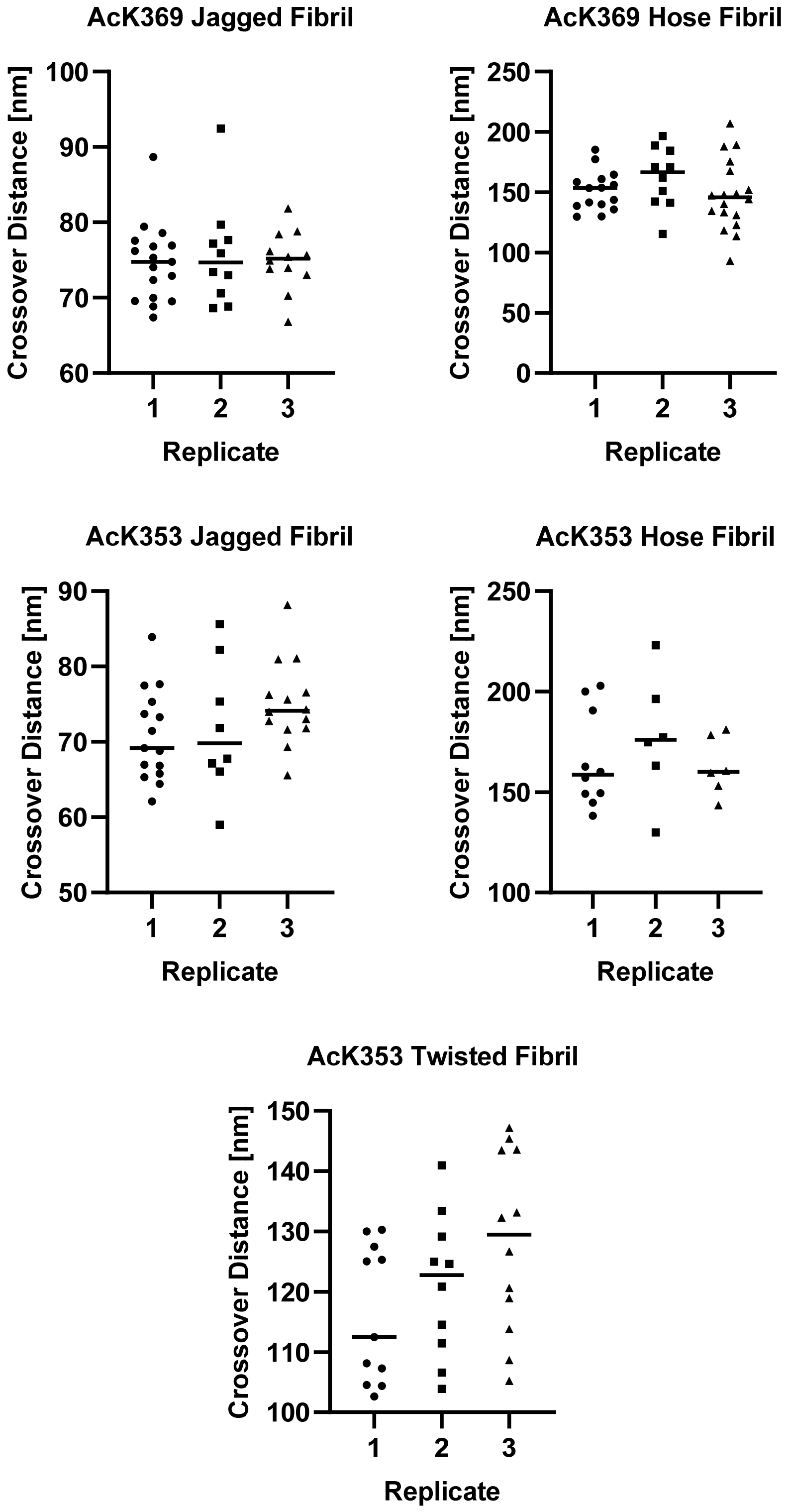
AcK369 Jagged Fibril
Crossover Distance [nm]
100
90
80
70
60
1
2
3
Replicate
AcK369 Hose Fibril
Crossover Distance [nm]
250
200
150
100
50
0
1
2
3
Replicate
AcK353 Jagged Fibril
Crossover Distance [nm]
90
80
70
60
50
1
2
3
Replicate
AcK353 Hose Fibril
Crossover Distance [nm]
250
200
150
100
1
2
3
Replicate
AcK353 Twisted Fibril
Crossover Distance [nm]
150
140
130
120
110
100
1
2
3
Replicate

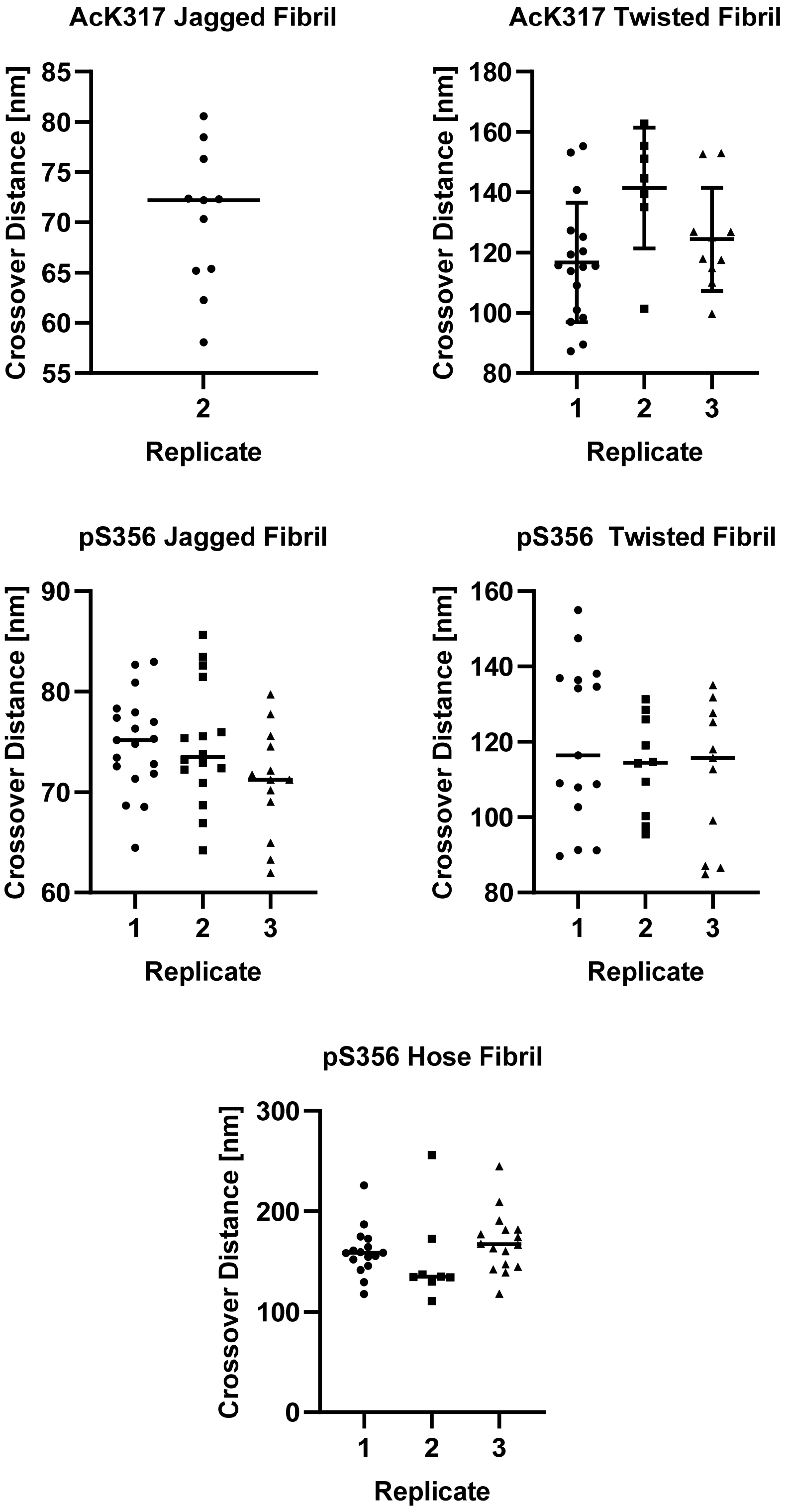
AcK317 Jagged Fibril
Crossover Distance [nm]
85
80
75
70
65
60
55
2
Replicate
AcK317 Twisted Fibril
Crossover Distance [nm]
180
160
140
120
100
80
1
2
3
Replicate
pS356 Jagged Fibril
Crossover Distance [nm]
90
80
70
60
1
2
3
Replicate
pS356 Twisted Fibril
Crossover Distance [nm]
160
140
120
100
80
1
2
3
Replicate
pS356 Hose Fibril
Crossover Distance [nm]
300
200
100
0
1
2
3
Replicate

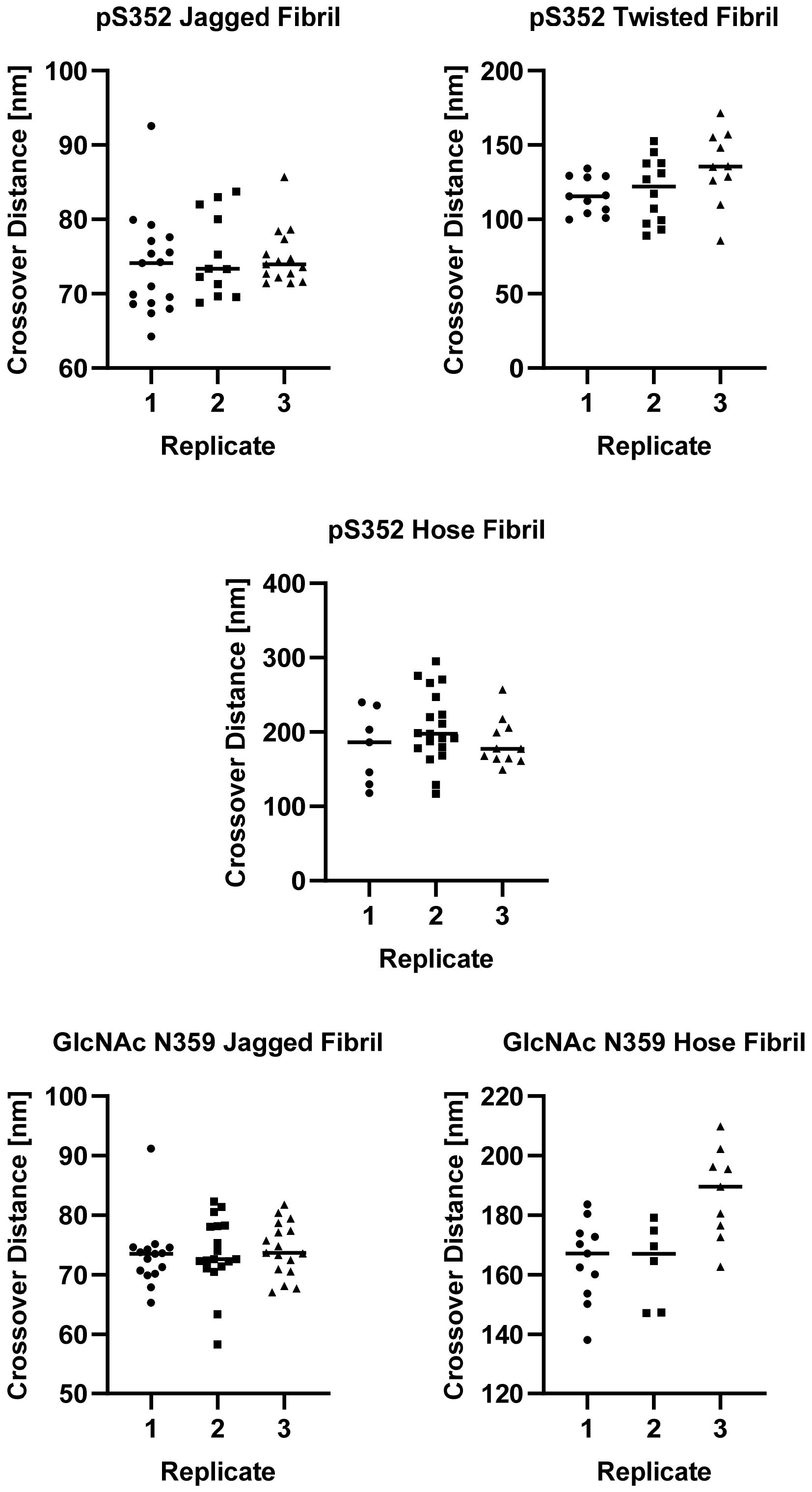
pS352 Jagged Fibril
Crossover Distance [nm]
100
90
80
70
60
1
2
3
Replicate
pS352 Twisted Fibril
Crossover Distance [nm]
200
150
100
50
0
1
2
3
Replicate
pS352 Hose Fibril
Crossover Distance [nm]
400
300
200
100
0
1
2
3
Replicate
GlcNAc N359 Jagged Fibril
Crossover Distance [nm]
100
90
80
70
60
50
1
2
3
Replicate
GlcNAc N359 Hose Fibril
Crossover Distance [nm]
220
200
180
160
140
120
1
2
3
Replicate

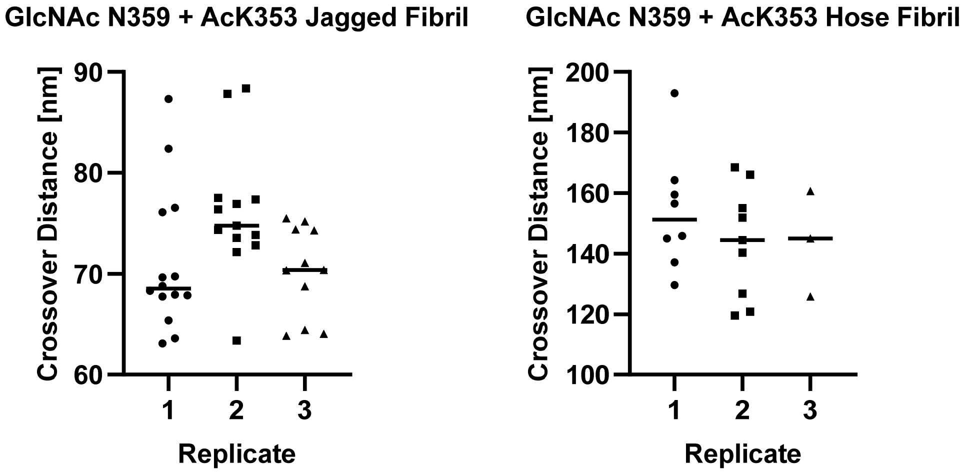
GlcNAc N359 + AcK353 Jagged Fibril
Crossover Distance [nm]
90
80
70
60
1
2
3
Replicate
GlcNAc N359 + AcK353 Hose Fibril
Crossover Distance [nm]
200
180
160
140
120
100
1
2
3
Replicate

#### 19. TURBIDITY DIAGRAMS

Turbidity phase diagrams were measured on a Nanodrop instrument. The absorbance was measured at 350 nm with a path length of 0.1 cm. A 0.65 µL Eppendorf tube was charged with 3 µL of 2x protein stock. Next, 3 µL of 2x poly(U) RNA stock solution was added, and the mixture was pipetted up and down several times. A minute after mixing, 2 µL of the sample was transferred to Nanodrop, and the absorbance was measured three times. Using the same LLPS mixture, this was repeated twice more. This experiment was performed in triplicate, and the data from the three experiments was averaged and reported as the standard error of the mean.

**Stock Solutions.** Filtered through a 0.22 µM filter prior to use.

- 2x protein stock: 60 µM Tau(291-391), 10 mM DTT, and 25 mM HEPES (pH 7.4).
- 2x poly(U) RNA stock: 0, 20, 40, 60, 80, 100, 120, 140, 160, 180, 200, 220 µg/mL poly(U) RNA in 25 mM HEPES (pH 7.4).

**Final Concentration.** 30 µM Tau(291-391), x µg/mL poly(U) RNA, 5 mM DTT, 25 mM HEPES (pH 7.4), 23 ºC.

**Turbidity Diagrams of Tau(291-391) with Poly(U) RNA**

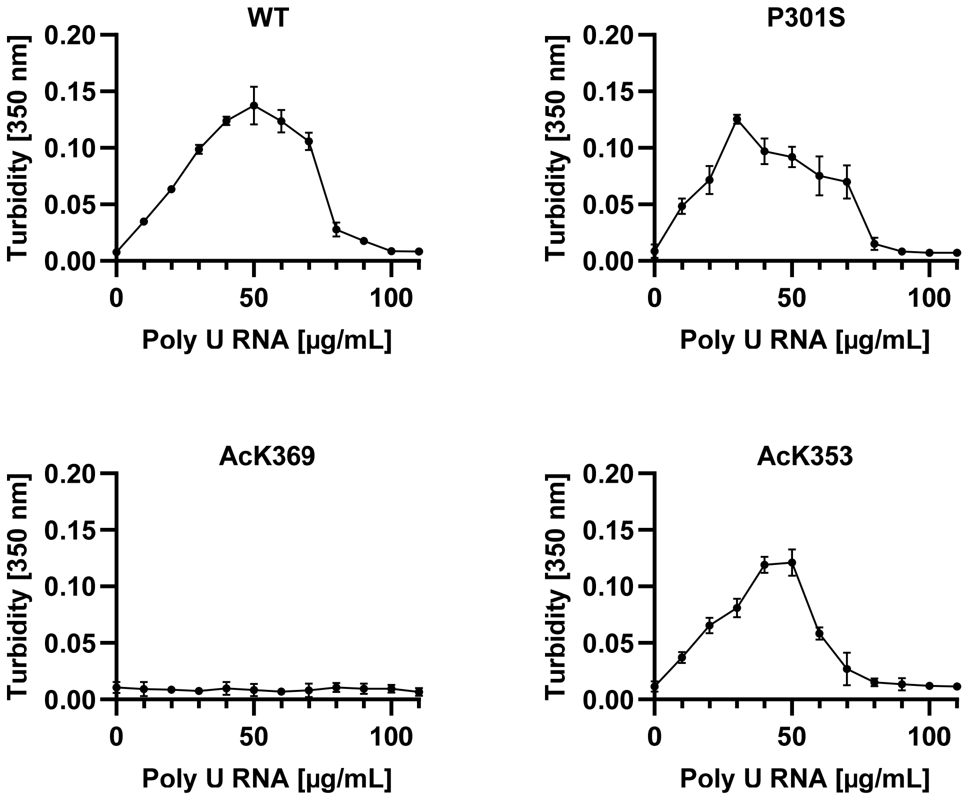

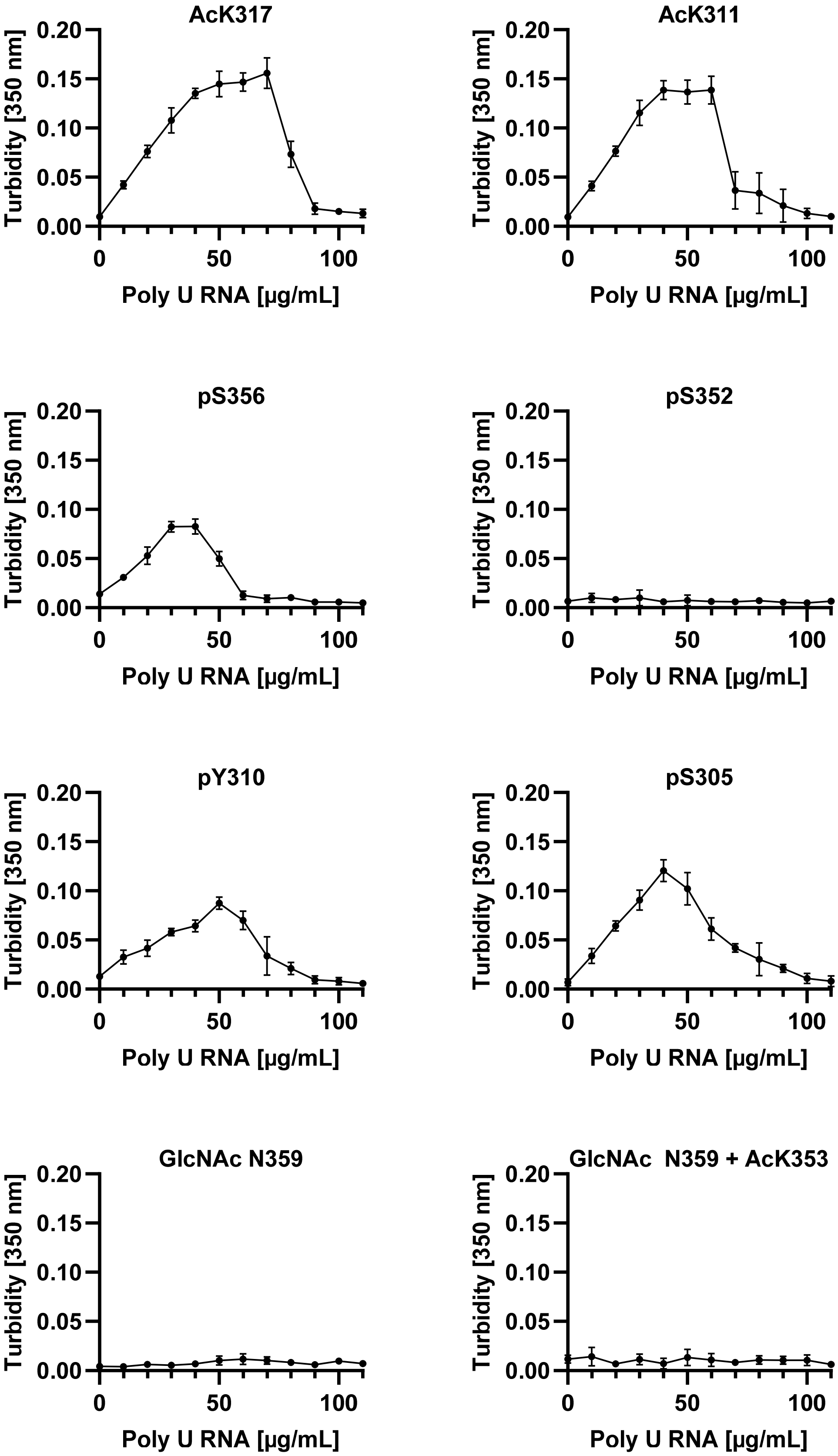
AcK317
AcK311
pS356
pS352
pY310
pS305
GlcNAc N359
GlcNAc N359 + AcK353
Turbidity [350 nm]
0.00
0.05
0.10
0.15
0.20
Poly U RNA [µg/mL]
0
50
100

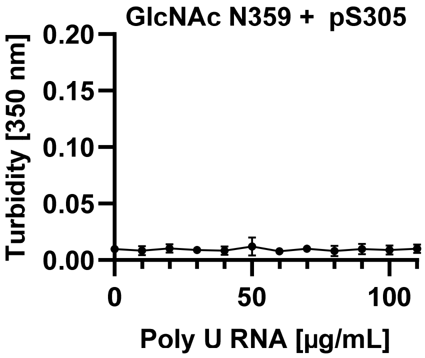
GlcNAc N359 + pS305
Turbidity [350 nm]
0.20
0.15
0.10
0.05
0.00
0
50
100
Poly U RNA [µg/mL]

### 20. SALT RESISTANCE

The salt resistance assay was measured by turbidity on Nanodrop. The absorbance was measured at 350 nm with a path length of 0.1 cm. A 0.65 µL Eppendorf tube was charged with 3 µL of 3x protein stock. The mixture was treated with 1.5 µL of 6x NaCl stock. Next, 4.5 µL of 2x RNA stock was added and pipetted up and down several times. A minute after mixing, 2 µL of the droplet solution was transferred to Nanodrop, and the absorbance was measured three times. Using the same LLPS mixture, this was repeated twice more. This experiment was performed in triplicate, and the data from the three experiments was averaged and reported as the standard error of the mean.

**Stock Solutions:**

- 3x protein stock: 90 µM Tau(291-391), 15 mM DTT, and 25 mM HEPES (pH 7.4).
- 6x NaCl stock: 30, 60, 90, 120 mM NaCl in 25 mM HEPES (pH 7.4).
- 2x poly U RNA stock: 80 µg/mL poly(U) RNA in 25 mM HEPES (pH 7.4).

**Final Concentration:**

- 30 µM Tau(291-391), 40 µg/mL poly(U) RNA, x mM NaCl, 5 mM DTT, 25 mM HEPES (pH 7.4), 23 ºC.

**Salt Resistance Turbidity Diagrams of Tau(291-391) with Poly(U) RNA**

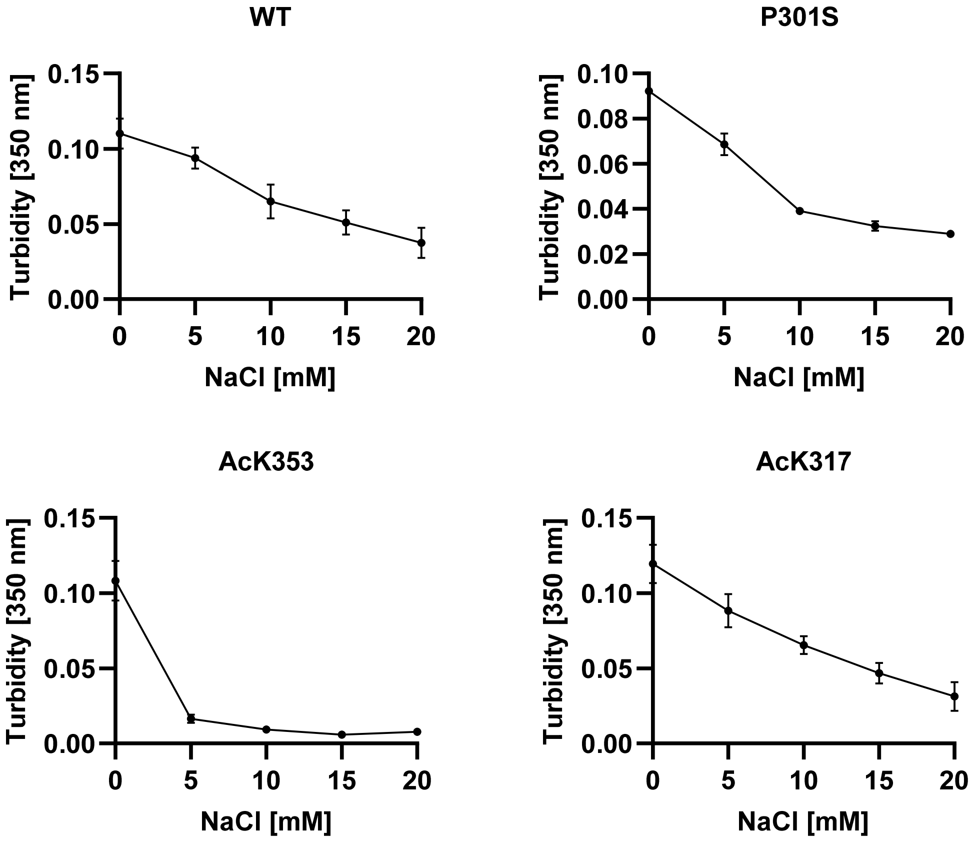

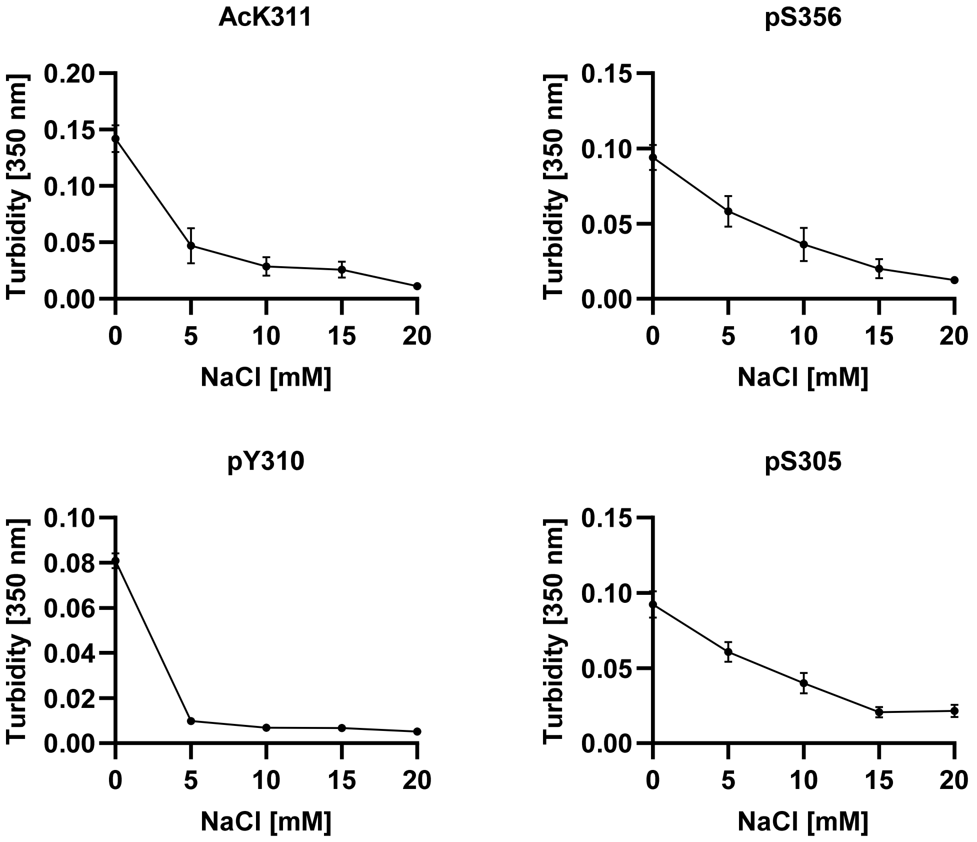
AcK311
pS356
pY310
pS305
Turbidity [350 nm]
NaCl [mM]
0.20
0.15
0.10
0.05
0.00
0.08
0.06
0.04
0.02
0
5
10
15
20

#### 21. LABELING OF TAU(291-391)

The labeling protocol is adapted from the AlexaFluor™ 488 5-SDP Ester (Thermo Scientific, Ref A30052) manufactures protocol. Tau(291-391) (8 mg/mL, 25 µL, 200 µg) in $H_2O$ was treated with $NaHCO_3$ (1 M, 3 µL) and Alexafluor488 SDP ester in DMSO (10 mg/mL, 3 µL). The reaction was incubated at 23 ºC for 2 h before it was quenched with hydroxylamine (50% (w/v), 10 µL). The excess label was removed by desalting over Sephadex G25 medium into $H_2O$, and the fractions were assayed for protein with gel electrophoresis. The fractions with protein were combined and lyophilized. The protein was redissolved in 25 mM HEPES (25 µL) and further purified by dialysis into 25 mM HEPES (pH 7.4) over a 7,000 MWCO Slide-A-Lyzer MINI Dialysis Unit (Thermo Scientific, Ref 69560). After 48 h, the protein concentration was determined by the Pierce BCA protein assay kit (Thermo Scientific), and the dye concentration was determined on Nanodrop by absorbance at 495 nm, with an extinction coefficient of 71,000 $M^{-1}cm^{-1}$. The protein-dye ratio was around 1:1 for all of the different Tau(291-391) constructs, and it is reported in the table below.

**Table S4: Tau(291-391) and AF488 Concentrations After Labeling**

| Tau | MW [g/mol] | Tau (BCA) [µM] | AF488 (A495) [µM] |
|---|---|---|---|
| WT | 10770.233 | 101 | 166 |
| P301S | 10760.194 | 72.5 | 76.2 |
| AcK369 | 10812.279 | 135 | 137 |
| AcK353 | 10812.279 | 174 | 161 |
| AcK317 | 10812.279 | 27.1 | 26.7 |
| AcK311 | 10812.279 | 131 | 107 |
| pS356 | 10849.204 | 168 | 223 |
| pS352 | 10849.204 | 121 | 166 |
| pY310 | 10849.204 | 108 | 159 |
| pS305 | 10849.204 | 158 | 125 |
| GlcNAcN359 | 10973.427 | 146 | 152 |
| GlcNAcN359 + AcK353 | 11015.464 | 306 | 305 |
| GlcNAcN359 + pS305 | 11064.488 | 285 | 233 |

#### 22. DROPLET SIZE MEASUREMENTS BY FLUORESCENCE MICROSCOPY

A 0.65 µL Eppendorf tube was charged with 3 µL of 2x protein. Next, 3 µL of 2x poly(U) RNA stock is added, and the mixture was pipetted up and down several times. The sample was transferred to the center of the micro-well in a 35 mm Glass bottom dish with 14 mm micro-well. The edge of the microwell was lined with 25 mM HEPES (~ 12 µL) to provide an evaporation shield, the microwell was covered with a glass coverslip, and the chamber was sealed with nail polish. For each droplet preparation, five random locations were imaged in 10-minute intervals. The experiment was repeated for a total of 3 or 4 independent experiments.

**Stock Solutions:**

- 2x protein stock: 60 µM Tau(291-391), 0.6 µM AF488, 10 mM DTT, and 25 mM HEPES (pH 7.4).
- 2x poly U RNA stock: 80 µg/mL poly U RNA in 25 mM HEPES (pH 7.4).

**Final Concentration:**

- 30 µM Tau(291-391), 0.3 µM AF488 (labeled Tau(291-391)), 40 µg/mL poly U RNA, 5 mM DTT, 25 mM HEPES (pH 7.4), 23 ºC.

**Microscope Settings:**

A Nikon Eclipse Ti Laser Scanning Confocal microscope and 60x oil immersion objective were used for this experiment. Droplets were imaged in the EGFP channel with 3 laser power and 5 detector gain. Pinhole size is 1.2. All imaging fields use Nyquist 0.09 µm and a 512x512 field. Droplets were imaged at five locations at the bottom of the dish using perfect focus (PFS) and autoscaled to avoid saturation. Movies were analyzed using FIJI/ImageJ using the following workflow. Open Movie -> Image -> 8 bit -> Thresholding -> Huang Threshold -> Analyze Particles -> size = 0.1-Infinity; circularity = 0-1; show: show overlay
masks; summarize, clear results, don't exclude edges. The experiment is repeated three independent times.

Replicates were processed separately and averaged. The data from the three triplicate experiments was averaged and displayed graphically with error bars representing standard error of the mean. Area over time represents the average area of the image covered by fluorescent droplets. Size over time represents the average area of individual circular droplets.

#### Percent Area and Droplet Area Kinetic Plots of Tau(291-391)-Poly(U) RNA Droplets

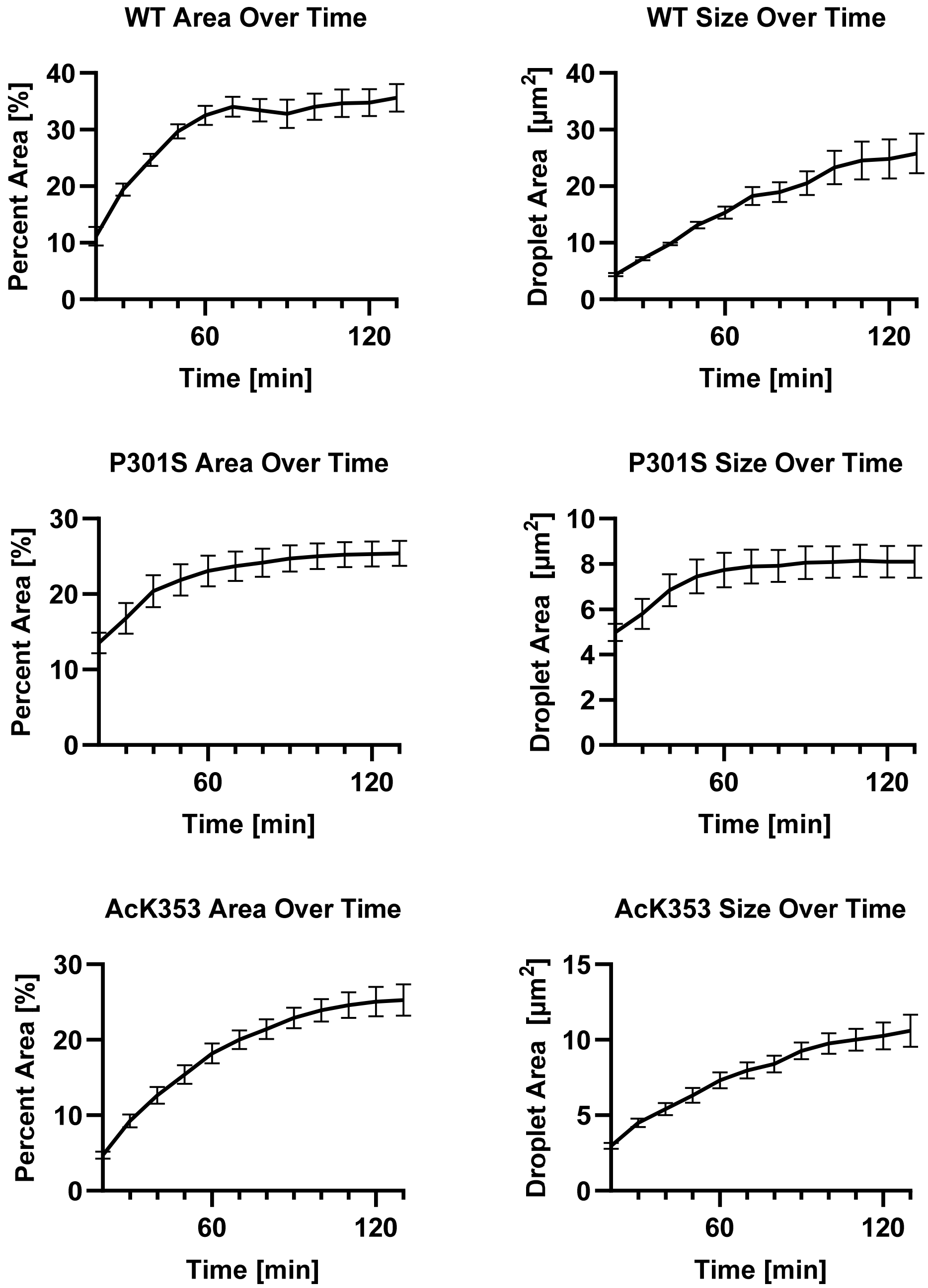

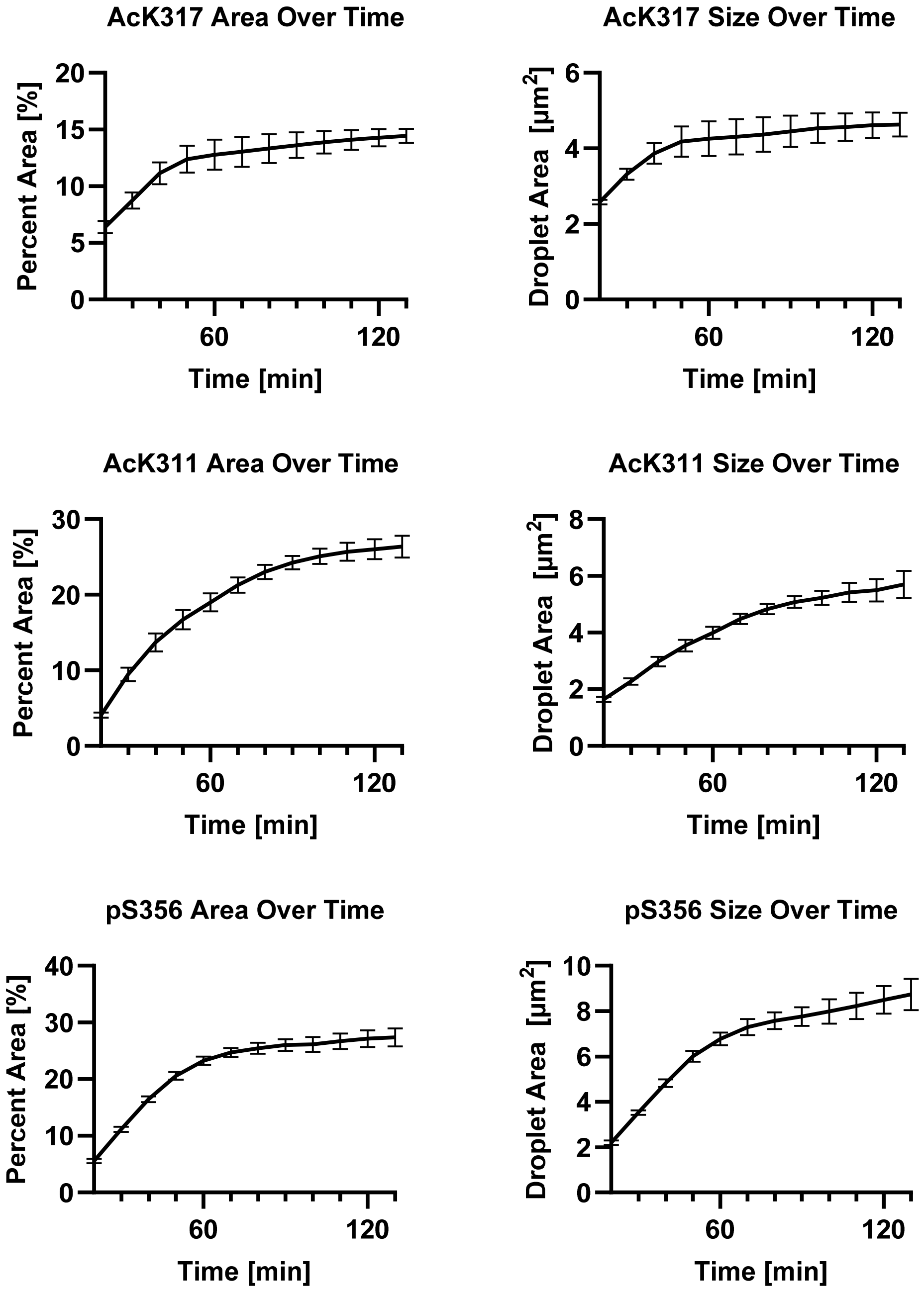

AcK317 Area Over Time
Percent Area [%]
Time [min]
AcK317 Size Over Time
Droplet Area [µm2]
Time [min]
AcK311 Area Over Time
Percent Area [%]
Time [min]
AcK311 Size Over Time
Droplet Area [µm2]
Time [min]
pS356 Area Over Time
Percent Area [%]
Time [min]
pS356 Size Over Time
Droplet Area [µm2]
Time [min]

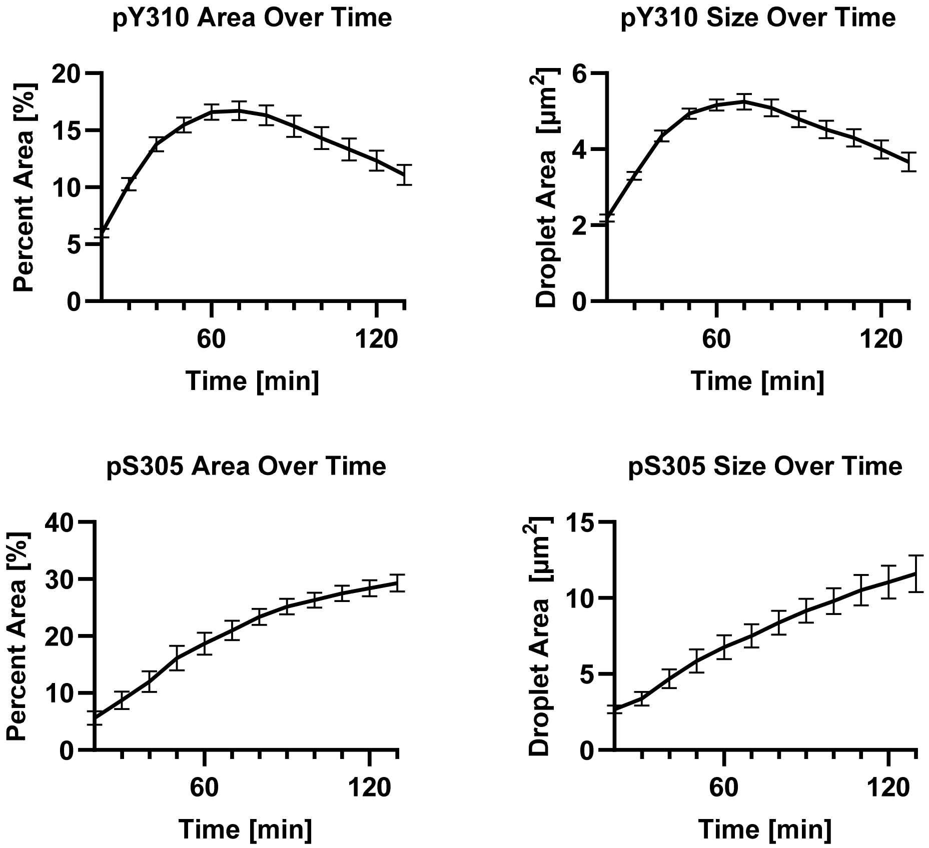
pY310 Area Over Time
Percent Area [%]
20
15
10
5
0
60
120
Time [min]
pY310 Size Over Time
Droplet Area [μm2]
6
4
2
0
60
120
Time [min]
pS305 Area Over Time
Percent Area [%]
40
30
20
10
0
60
120
Time [min]
pS305 Size Over Time
Droplet Area [μm2]
15
10
5
0
60
120
Time [min]

#### 23. FRAP

A 0.65 µL Eppendorf tube was charged with 3 µL of 2x protein stock. Next, 3 µL of 2x poly(U) RNA stock was added, and the mixture was pipetted up and down several times. The sample was transferred to the center of the micro-well in a 35 mm Glass bottom dish with 14 mm micro-well. The edge of the microwell was lined with 25 mM HEPES (~ 12 µL) to provide an evaporation shield, and the microwell was covered with a glass coverslip, and the chamber was sealed with nail polish. For each droplet preparation, one droplet was photobleached after 1 h, and another droplet aws photobleached after 2 h. The experiment is repeated for a total of 5 independent experiments.

**Stock Solutions:**

- 2x protein stock: 60 µM Tau(291-391), 0.6 µM AF488, 10 mM DTT, and 25 mM HEPES (pH 7.4).
- 2x poly U RNA stock: 80 µg/mL poly U RNA in 25 mM HEPES (pH 7.4).

**Final Concentration:**

- 30 µM Tau(291-391), 0.3 µM AF488 (labeled Tau(291-391)), 40 µg/mL poly U RNA, 5 mM DTT, 25 mM HEPES (pH 7.4), 23 ºC.

**FRAP Microscope Settings:**

A Nikon Eclipse Ti Laser Scanning Confocal microscope and 60x oil immersion objective was used for this experiment. Droplets were imaged in the EGFP channel with 3 laser power and 5 detector gain. The pinhole size was 1.2. All imaging fields used Nyquist 0.09 µm and a 1024x1024 field. Droplets were imaged at the bottom of the dish using PFS and autoscaled to avoid saturation. The 488 channel was selected for photobleaching with a laser power of 20. Stimulation, background, and reference ROIs were all circular ROIs with a 1.2 x 1.2 feret.

Timing for FRAP experiments was as follows:

- Acquisition after 2 second intervals over 10 seconds (5 data points).
- Stimulation for 2 seconds.
- Acquisition after 2 second intervals over 1 minute 30 seconds (45 data points).
- Acquisition after 10 second intervals over 7 minutes (42 data points).

The data was analyzed in GraphPad Prism using a double normalization, according to the literature.[11] Replicates were averaged and displayed graphically with error bars representing the standard deviation.

**Mobile Fraction Calculation:**

The mobile fraction is calculated with the equation:

$$MF = \frac{I_\infty - I_c}{I_{c_0} - I_c}$$

MF: Mobile fraction is reported as a percentage.

$I_\infty$: fluorescence intensity at saturation end value at post recovery.

$I_c$: fluorescence intensity at photobleaching.

$I_{c0}$: fluorescence intensity before photobleaching.

#### FRAP Kinetic Plots of Tau(291-391)-Poly(U) RNA Droplets

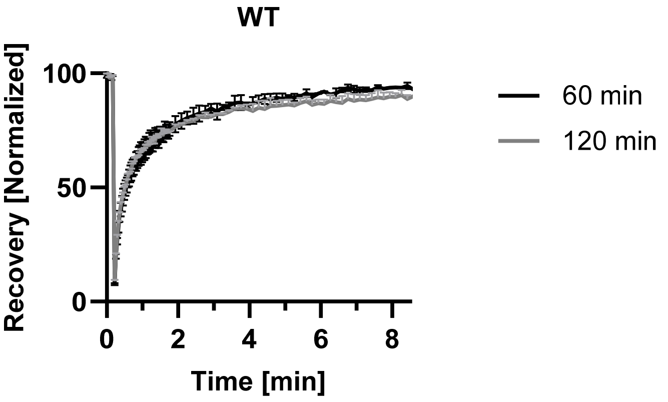

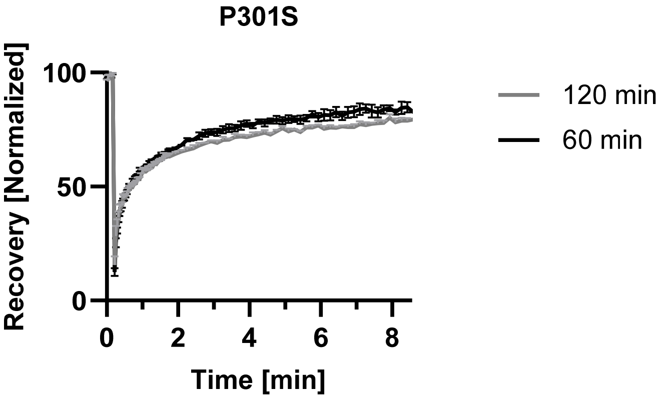

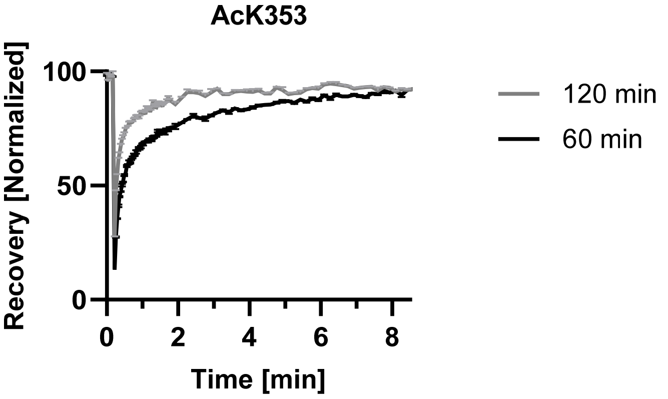

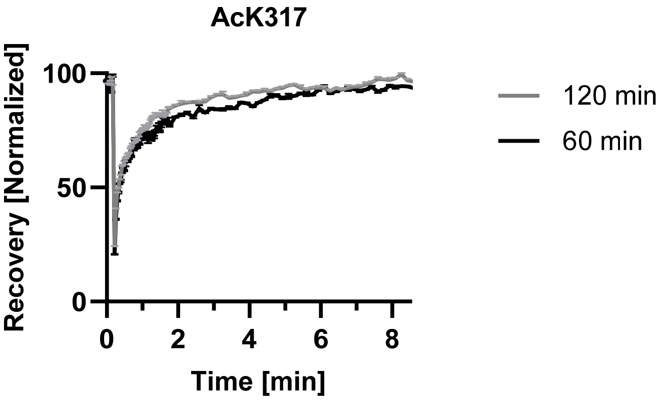
AcK317
Recovery [Normalized]
100
50
0
0
2
4
6
8
Time [min]
120 min
60 min

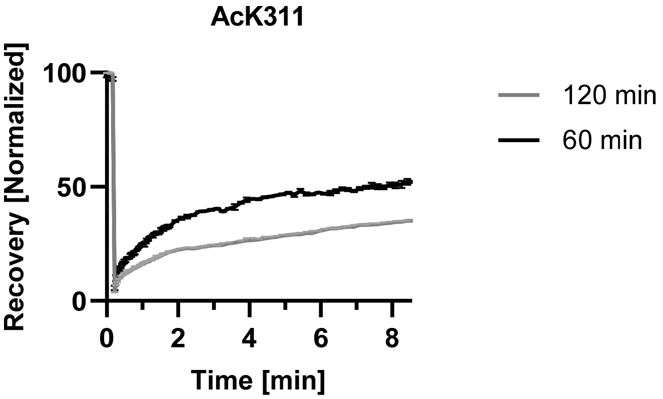
AcK311
Recovery [Normalized]
100
50
0
0
2
4
6
8
Time [min]
120 min
60 min

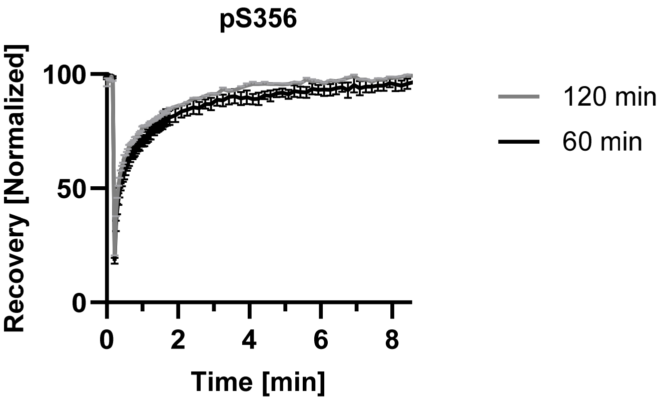
pS356
Recovery [Normalized]
100
50
0
0
2
4
6
8
Time [min]
120 min
60 min

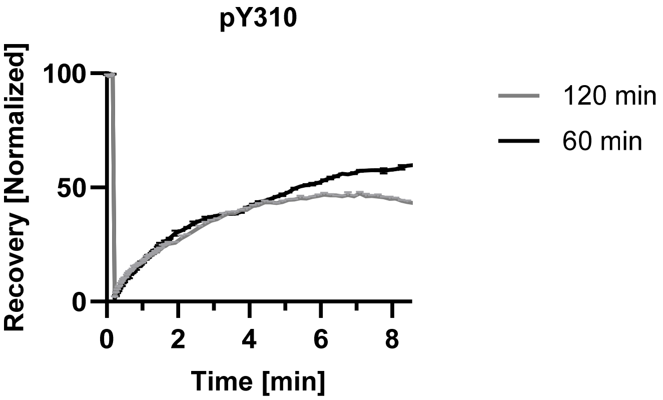

pY310
Recovery [Normalized]
100
50
0
0
2
4
6
8
Time [min]
120 min
60 min

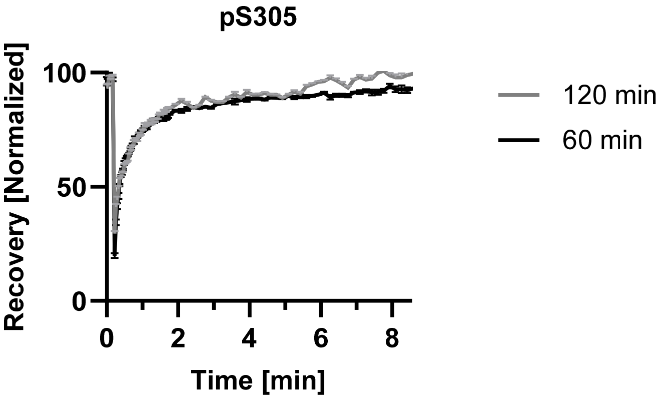

pS305
Recovery [Normalized]
100
50
0
0
2
4
6
8
Time [min]
120 min
60 min
